# Deciphering the Molecular Mechanism of Post-Acute Sequelae of COVID-19 through Comorbidity Network Analysis

**DOI:** 10.1101/2024.01.17.575851

**Authors:** Lue Tian, Ian C.K. Wong, Qingpeng Zhang

## Abstract

**Introduction:** The post-acute sequelae of COVID-19 presents a significant health challenge in the post-pandemic world. Our study aims to analyze longitudinal electronic health records to determine the impact of COVID-19 on disease progression, provide molecular insights into these mechanisms, and identify associated biomarkers.

**Method:** We included 58,710 patients with COVID-19 records from 01/01/2020 to 31/08/2022 and at least one hospital admission before and after the acute phase of COVID-19 (28 days) as the treatment group. A healthy control group of 174,071 individuals was established for comparison using propensity score matching based on pre-existing diseases (before COVID-19). We built a comorbidity network using Pearson correlation coefficient differences between pairs of pre-existing disease and post-infection disease in both groups. Disease-protein mapping and protein-protein interaction network analysis revealed the impact of COVID-19 on disease trajectories through protein interactions in the human body.

**Results:** The disparity in the weight of prevalent disease comorbidity patterns between the treatment and control groups highlights the impact of COVID-19. Certain specific comorbidity patterns show a more pronounced influence by COVID-19. For each comorbidity pattern, overlapping proteins directly associated with pre-existing diseases, post-infection diseases, and COVID-19 help to elucidate the biological mechanism of COVID-19’s impact on each comorbidity pattern. Proteins essential for explaining the biological mechanism can be identified based on their weights.

**Conclusion:** Disease comorbidity associations influenced by COVID-19, as identified through longitudinal electronic health records and disease-protein mapping, can help elucidate the biological mechanisms of COVID-19, discover intervention methods, and decode the molecular basis of comorbidity associations. This analysis can also yield potential biomarkers and corresponding treatments for specific disease patterns.

**Ethical approval:** Ethical approval for this study was granted by the Institutional Review Board of the University of Hong Kong/HA HK West Cluster (UW20-556, UW21-149 and UW21-138).

**RESEARCH IN CONTEXT:** *Evidence before this study:* We searched PubMed for research articles up to Nov 30, 2022, with no language restrictions, using the terms “Post-Acute Sequelae of COVID-19” OR “PASC” OR “Long COVID” AND “comorbidity” OR “multimorbidity” OR “co-morbidity” OR “multi-morbidity”. We found most related papers focus on the comorbidity or multimorbidity patterns among PASC. Some papers focus on the associations between specific diseases and PASC. However, no study investigated the biological mechanism of PASC from the perspective of comorbidity network.

*Added value of this study:* This study investigated the biological mechanism of PASC based on the comorbidity network including the impact of pre-existing diseases (diseases diagnosed within 730 days before COVID-19) on the development of PASC. We classified pairs of pre-existing disease and post-infection disease (new diseases diagnosed in 28 days to 180 days after COVID-19) as comorbidity associations. Through a comparison of the frequency of comorbidity associations in health people group and patients with COVID-19 infection group, we identified comorbidity patterns that are significantly influenced by COVID-19 infection and constructed a comorbidity network comprising of 117 nodes (representing diseases) and 271 edges (representing comorbidity patterns). These comorbidity patterns suggest COVID-19 patients with these pre-existing diseases have higher risk for post-infection diseases. Through the analysis of the Protein-Protein interaction (PPI) network and associations between diseases and proteins, we identified key proteins in the topological distance of each comorbidity pattern and important biological pathways by GO enrichment analysis. These proteins and biological pathways provide insights into the underlying biological mechanism of PASC.

*Implications of all the available evidence:* The identification of elevated-risk comorbidity patterns associated with COVID-19 infection is crucial for the effective allocation of medical resources, ensuring prompt care for those in greatest need. Furthermore, it facilitates the recovery process of patients from COVID-19, offering a roadmap for their path back to health. The key proteins identified in our study have the potential to serve as biomarkers and targets for therapeutic intervention, thereby establishing a foundation for the development of new drugs and the repurposing of existing ones. Further research should focus on drug discovery and the development of drug recommendations for patients with COVID-19 infections.

## INTRODUCTION

The global pandemic of Coronavirus Disease 2019 (COVID-19), caused by severe acute respiratory syndrome coronavirus 2 (SARS-CoV-2) infection, has impacted billions of people, resulting in millions of deaths, and leaving tens of millions suffering from persistent symptoms and signs after the acute phase of COVID-19^1^. This phenomenon, referred to as post-acute sequelae of COVID-19 (PASC), is gradually coming to an end^2,3^. PASC exhibits heterogeneous manifestations and severity^4^, impacting various organ systems including cardiovascular^5,6^, mental^7^, metabolic^8^, and renal systems^9^. However, the underlying biological mechanisms of PASC remain intricate and elusive.

Current research on the potential biological mechanisms of COVID-19 and PASC primarily involves small patient cohorts and concentrates on the relationship between COVID-19 and diseases in specific systems or organs individually^2,10–12^. Investigations involving large patient cohorts and associations between COVID-19 and diseases across multiple organs and systems can aid in uncovering the biological mechanisms of multimorbidity present in PASC and pre-existing diseases influenced by COVID-19. Comorbidity and multimorbidity^13^ refer to the co-occurrence of two or more diseases in an individual. If the frequency of co-occurrence of diseases exceeds the frequency of disease combinations selected by chance, multimorbidity exists among these diseases. Research on multimorbidity associations with PASC has shown that pre-existing multimorbidity may drive PASC^10,14^. The underlying mechanisms of multimorbidity are complex, potentially involving shared genetic or environmental factors or resulting from the treatment or intervention for one disease leading to the development of another^14^. Studies on multimorbidity relations in PASC have considered the influence of demographic factors^15,16^ like sex, age, race, vaccine injection, and patient electronic health records (EHR) diseases, as well as acute infection phase severity. However, the underlying biological mechanisms of these multimorbidity relations influenced by COVID-19 remain unclear.

Our research utilized territory-wide EHR data from Hong Kong Hospital Authority to investigate the impact of COVID-19 on PASC. We employed network-based methods to assess the influence of COVID-19 on multimorbidity across multiple organs and systems in PASC. Individuals with specific pre-existing diseases may have a higher risk of developing certain diseases included in PASC due to the effects of COVID-19. To explore the underlying mechanisms of these particular multimorbidity relations, we incorporated biological knowledge from the Protein-Protein Interaction (PPI) network^17^ and Gene Ontology (GO) enrichment analysis^18^ to identify the most affected and essential proteins that could serve as potential targets for future interventions, such as preventive measures.

## RESULTS

### Overall pipeline

Using propensity score matching, we construct a healthy people group (control group) and a COVID-19 patient group (treatment group) with similar clinical records prior to their respective first COVID-19 record (simulating infection date for healthy individuals) from January 1, 2020 to August 31, 2022. Then, we construct and analyze the comorbidity network as follows:

- For each individual, we first define *pre-existing diseases* as those diagnosed within 730 days before COVID-19, and *post-infection diseases* as new diseases diagnosed in 28 days to 180 days after COVID-19.
- Then, we define the disease pair as a pair of pre-existing disease and a post-infection disease in the treatment group. We identify *comorbidity patterns* as these disease pairs with Pearson correlation coefficient > 0, relative risk > 1 and frequency > 10.
- We then construct a comorbidity network^19^ consisting of these comorbidity patterns and involved diseases. In our comorbidity network, each node represents a disease. The existence of a directed edge from the pre-existing disease to the post-infection disease indicates the comorbidity pattern is significantly more frequent (p-value < 0.05) in the treatment group than in the control group. The difference of Pearson correlation coefficient and relative risk for each comorbidity pattern between treatment group and control group are considered as additional attributes for edges.
- For each comorbidity pattern, as represented by an edge in the comorbidity network, network analysis and GO enrichment analysis were employed for biological pathways discovery by utilizing corresponding disease-associated proteins with COVID-19 associated proteins, and important proteins and GO terms were identified.

### Study cohorts

Before matching, the dataset comprised 58710 individuals with COVID-19 and 488,585 individuals without COVID-19 (refer to Supplementary Table 1). Following the matching process (as detailed in the Methods section), the treatment group contained 58,710 observations, while the control group contained 174,071 observations. The median age for the control group is 68, compared to 66 for the treatment group. The control group was composed of 50.3% males and 49.7% females, whereas the treatment group consisted of 51.1% males and 48.9% females. Over 60% of the individuals in both groups are aged 60 years or older. Further details regarding specific diseases are shown in Supplementary Figure 1. Utilizing the electronic health records of each individual in the two groups, we were able to construct a directed comorbidity network, which illustrates the trajectories of diseases before and after COVID-19.

### Comorbidity Network Analysis

Our comorbidity network comprises 117 nodes and 271 edges, representing the comorbidity patterns significantly influenced by COVID-19. Nodes are categorized into 14 disease groups according to their ICD9 categories. Edges are classified into two groups according to the adjacent nodes: intra-group edges (source node and target node in the same disease group), inter-group edges (source node and target node in different disease groups).

Figure 1 depicts the constructed comorbidity network and disease group classifications. The network is heterogeneously connected, with most disease groups sparsely associated with other disease groups and a few disease groups more closely related to some other disease groups. More specifically, among all groups, disease group (001–139 Infectious and Parasitic Diseases) and disease group (460–519 Diseases of the Respiratory System) have a higher frequency than other disease groups. 079 (Viral and chlamydial infection in diseases classified elsewhere and of unspecified site) and 519 (Other diseases of respiratory system) are two most frequent diseases among all, indicating that COVID-19 has the most significant impact on the respiratory system. Additionally, the neural, gastrointestinal and circulatory systems are also frequently affected by the COVID-19, which is aligns with the literature^20–22^.Diseases such as 401(Essential hypertension), 278 (Overweight, obesity and other hyperalimentation), 780 (General symptoms like Dizziness/vertigo, NOS), 785 (Symptoms involving cardiovascular system), 783 (Symptoms concerning nutrition, metabolism and development) are also more likely to be involved in the comorbidity relationships. (Refer to Supplementary Table 2)

**Figure 1.**
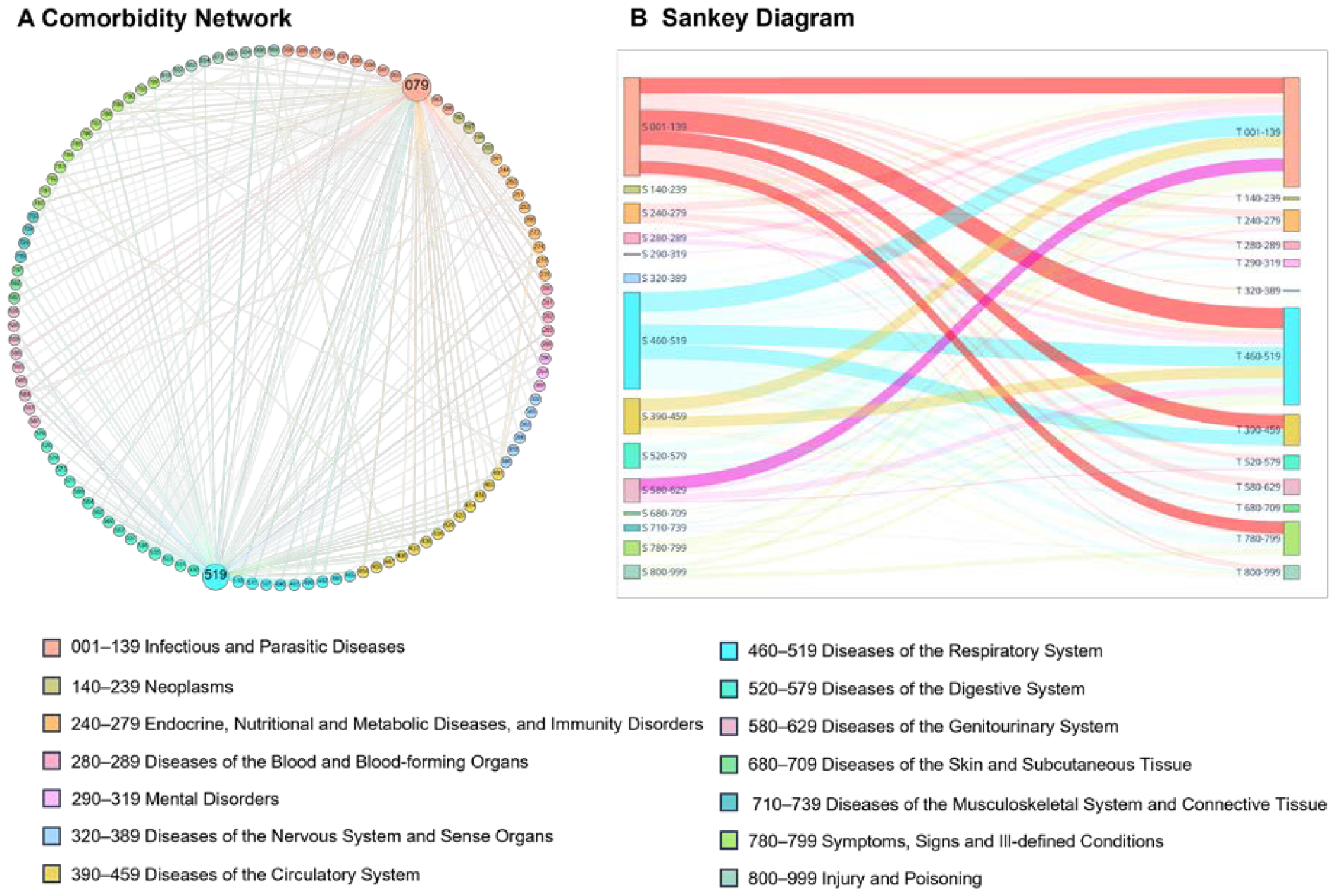
Visualization of the comorbidity network (A) and the associations among disease groups (B). In (A), the size of a node is proportional to its occurrence in our dataset. In (B), the rectangles in the Sankey diagram correspond to ICD 9 disease categories. The left rectangles represent disease groups of pre-existing diseases. The right rectangles represent disease groups of post-infection diseases. The edge linking a pair of rectangles indicates that the comorbidity patterns are significantly more frequent among COVID-19 patients as compared to those in control group. The thickness of an edge is proportional to increased occurrence among COVID-19 patients as compared to those in control group. Please refer to Supplementary Table 3 for more details.

The comorbidity network’s edges suggest more pronounced comorbidity relationships in the treatment group compared to the control group. These relationships imply that patients with a history of respiratory system diseases are at an increased risk of developing further respiratory system diseases due to COVID-19. Additionally, these patients also demonstrate a heightened risk for diseases within the circulatory system, genitourinary system, among others. Patients previously diagnosed with essential hypertension also exhibit a higher risk of developing respiratory system diseases and lipid disorders due to COVID-19. Patients with a history of peptic ulcer (site unspecified) are at an increased risk of developing Gastritis and duodenitis (Refer to Supplementary Table 4).

### Biological Mechanism Explanation

To investigate the biological mechanisms underlying identified comorbidity relations, we utilize the PPI and GO terms associated with the diseases. We hypothesize that for COVID-19 to influence the disease comorbidity patterns of patients, its host factors (genes/proteins) should be localized in the corresponding subnetwork within the human PPI network, either directly targeting the disease-associated genes/proteins or indirectly affecting them through PPIs. Specifically, we hypothesize that these comorbidity patterns identified from the comorbidity network experience a larger reduction in the topological distance in the PPI network caused by the inclusion of proteins associated with COVID-19 as additional proteins associated with pre-existing disease. Similar patterns were previously observed in the relationship between COVID-19 and brain microvascular injury. ^23^

After we eliminate disease pairs without associated protein information, 230 pre-existing disease-post-infection disease pairs, including 95 disease types (79 pre-existing diseases, 57 post-infection diseases), remain. Then, the network distance from pre-existing diseases to post-infection diseases is measured on the PPI network. The network distance is measured as the averaged shortest path length between each protein associated with post-infection disease and proteins associated with pre-existing disease. Next, we measure the change of network distance of pre-existing diseases and post-infection diseases in the PPI network when treating COVID-19 as an additional pre-existing disease. We observed that the network distance between the disease pairs with elevated comorbidity risk after COVID-19 became shorter (mean value 0.649 vs 0.565, p-value=0.049) because of the addition of proteins associated with COVID-19. Please refer to the Method section for two hypothesis testing methods (Z-test within treatment groups Supplementary Table 5, Mann-Whitney U test between treatment and control groups) Further GO analysis of involved proteins reveals that COVID-19 introduced additional mechanistic pathways towards post-infection diseases, effectively increasing the risk of developing post-infection diseases. Please refer to the Supplementary Table 6 for a list of frequent GO terms associated with these proteins.

Among involved proteins, *overlapping* proteins (those associated with both the post-infection disease and pre-existing disease with COVID-19) play the major role in shortening the distance between the disease pair. Please refer to the Supplementary Materials for a list of the most frequent overlapping proteins (Refer to Supplementary table 7). GO enrichment analysis reveals that, in addition to the roles in COVID-19 and following inflammatory response, the expression disorder of these overlapping proteins is leading towards other diseases involving cardiovascular system, urinary system, and respiratory system.

Figure 2 illustrates the interplays between a representative disease comorbidity pair, from 401 (Essential hypertension) to 272 (Disorders of lipoid metabolism), which has an elevated risk because of the COVID-19 infection. We find that the overlapping proteins associated with both the post-infection disease (272) and pre-existing diseases (401 and COVID-19) play an important role in the development of disease comorbidity. There are five additional overlapping proteins because of the COVID-19 infection: NEU1, INHBE, NPC2, AGPS and GLA.^24,25^ Specifically, NEU1 has a significant effect on lipid metabolism and inflammatory processes and is a potential drug target for decreasing atherosclerosis^26,27^. INHBE activates energy expenditure through brown/beige adipocyte activation, and it can be a potential drug target for obesity therapy^28,29^. NPC2 is essential for the pathways involved in glucose and lipid metabolism, helping the egress of lipids from the lysosome^30,31^. AGPS is a ether lipid generating enzyme which is important for the balance of structural and signaling lipids.^32^ GLA is a polyunsaturated fatty acid that can reduce lipid deposition.^33^ The addition of these five overlapping proteins potentially explains the mechanism of how COVID-19 elevated the risk of developing the post-infection disease (272).

**Figure 2.**
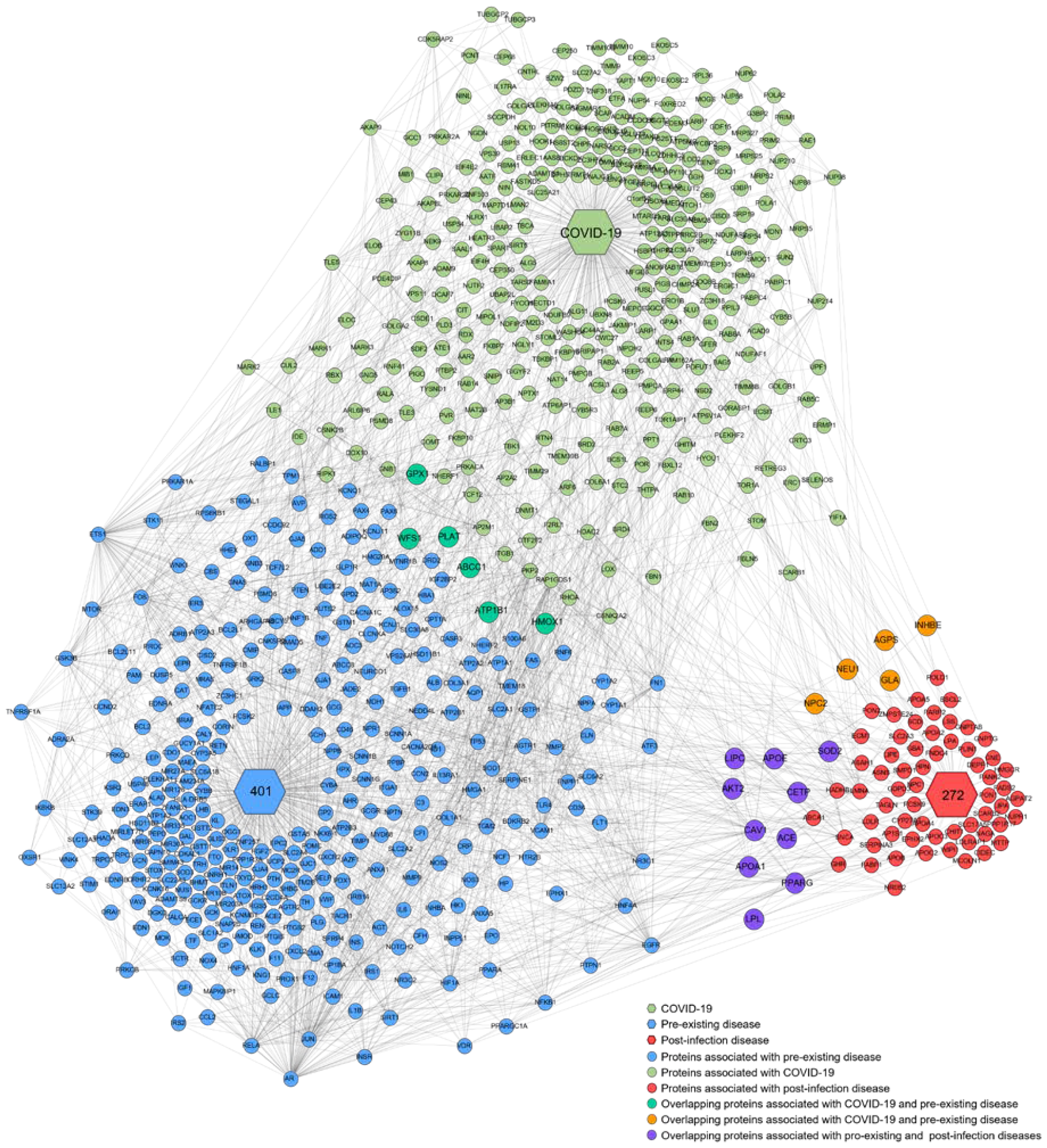
PPIs underlying the comorbidity pattern between ICD 401 and ICD 272. A hexagon represents a disease. A circle represents a protein. An edge between circles represents the existence of protein-protein interaction. An edge between a hexagon and a circle represents the protein associated with the disease.

## DISCUSSION

Our study, leveraging population-based EHR and a wealth of biomedical data, stands as a pioneering quantitative analysis of the complex molecular mechanisms underlying the comorbidity patterns associated with COVID-19. This research is not merely an exploration but a comprehensive examination of the data, aiming to unravel the progression of the disease and the evolution of comorbidity patterns resulting from a COVID-19 infection.

Our findings provide a deeper understanding of the phenomenon known as long COVID, or Post-Acute Sequelae of COVID-19 infection (PASC), a disease characterized by lingering symptoms after recovery from the acute phase of COVID-19 that has been a global concern for healthcare professionals. Our research illuminates the elevated-risk comorbidity patterns associated with PASC, significantly contributing to the existing body of knowledge on this subject.

Central to our study are the key proteins we identified, which play a pivotal role in increasing the risk of these comorbidity patterns. These proteins are not merely markers but potential targets for therapeutic intervention, laying the groundwork for the development of new drugs and the repurposing of existing ones. The ultimate goal is not only to reduce the risk of COVID re-infection but also to prevent the onset of PASC, offering hope to millions of patients worldwide.

The practical applications of our study are extensive. Using the comorbidity patterns, we discovered and the wealth of data from electronic health records, we can identify patients who are at high risk for PASC. This information is crucial for the effective allocation of medical resources, ensuring prompt care for those who need it most. Moreover, it aids in the recovery process of patients from COVID-19, providing a roadmap for their journey back to health. Our study, therefore, stands at the intersection of research and real-world application, contributing to the fight against this global pandemic.

Our study has limitations. First, our analyses were based on the topology of the PPI network. The PPI network serves as a “skeleton” of the biological signaling circuitry in the human body. However, PPI network cannot fully represent the pharmacokinetics and pharmacodynamics (PK/PD) associated with drugs. Future research is needed to incorporate the PK/PD models for a better understanding of the effects of drugs in the human body. Secondly, our population EHR data was obtained from public hospitals in Hong Kong. Although our data is among the most complete for a population, there is inevitable under-reporting problem, especially for young patients. Third, although we have tried our best to build a comprehensive mapping between diseases and proteins in the PPI network. The mapping may still be subject to bias because of the lack of such data. Further biological research is needed to enrich and complement existing databases.

## METHODS

### Study design and population

All electronic datasets included in this research are from the Hong Kong Hospital Authority (HKHA) database. Based on the COVID-19 record (based on rapid antigen test [RAT] or polymerase chain reaction [PCR] test in throat swab, nasopharyngeal aspirate, or deep throat sputum specimens), patients are divided into two groups: treatment and control groups. (Supplementary Figure 2) For the treatment group, the diagnose records within 730 days before COVID-19 and 180 days after COVID-19 are retained. We then exclude all diagnose records within 28 days (acute-phase of infection) after COVID-19. For each individual, diseases appearing before COVID-19 were considered as pre-existing diseases, and new emerging diseases appearing after COVID-19 were considered as post-infection diseases.

For control group, we applied the same procedure to identify the pre-existing and post-infection diseases for each individual by treating the date 180 days before the last record date as the simulated COVID-19 infection date.

To investigate the influence of COVID-19 on PASC, we compared the differences in comorbidity patterns between the treatment group (people with first COVID-19 in 2022) and the control group (people without COVID-19). To ensure a fair comparison, we employed propensity score matching^34^ to select a control group with pre-existing disease records similar to the treatment group. In the matching process, we considered not only individual EHR data denoted by ICD-9 codes from the baseline period, but also demographic data (age, sex) and vaccine information (vaccine number). After matching, we calculated the standardized mean difference (SMD) to quantify the balance for each confounder. An SMD value below 0.2 serves as a threshold to determine whether the confounder is well-balanced.

### Propensity Score Matching

We included all disease related ICD-9 CM codes from pre-existing diseases in both control and treatment groups. Each ICD-9 code serves as a feature in the Propensity Score Matching. Patients in the two groups are further divided into subgroups based on age and sex combinations, such as (0-20 years old, female), (20-40 years old, female), and so on up to (80+ years old, male). The number of vaccinations received prior to COVID-19 is also included as a feature. For each individual in the treatment group, we select the 15 nearest neighbors from the control group based on the processed propensity score, which is obtained by applying a logit function. After propensity score matching, the COVID-19 and control groups include 58,710 and 174,071 patients respectively, from 58,710 in the COVID-19 group and 488,585 in the control group.

### SMD Compute

standardized mean difference (SMD) is an indicator for evaluating the goodness-of-balance of confounders between two groups. For the confounder *X* in the treatment and control groups denoted by *X*_*t*_ and *X*_*c*_. And SMD < 0.2 means this confounder is balanced^2,35,36^.

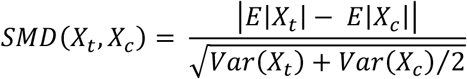

For dichotomous variable *XX*:

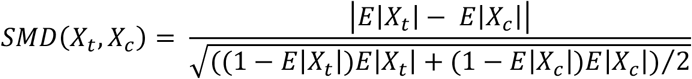

*E*|*X*| is the expectation of dichotomous variable *X*.

### Comorbidity network construction

According to COVID-19 infection date for each individual in treatment group (simulated for control group). we first define pre-existing diseases as those diagnosed before COVID-19, and post-infection diseases as new emerging diseases diagnosed >28 days after COVID-19. We generated disease pairs by selecting the first disease from all pre-existing diseases and the second disease from all post-infection diseases and applied Pearson correlation coefficient and relative risk to quantify the co-occurrence of two diseases composing each disease pair^37^. To reduce bias from data and computation methods, we used both values to identify co-occurring disease pairs combining with chi-square and fisher exact test^37^. We identified disease pairs appearing in the treatment group with Pearson correlation coefficient > 0 and relative risk > 1 as comorbidity patterns. To investigate comorbidity patterns influenced by COVID-19, we selected comorbidity patterns according to the two requirements: (1) comorbidity patterns are significantly more frequent (Fisher exact test or chi-square test, p-value < 0.05) in the treatment group than in the control group. (2) the count number of each comorbidity pattern in the treatment group is at least 10. (3) the Pearson correlation coefficient and relative risk of each comorbidity are larger in the treatment group than in the control group. Selected comorbidity patterns are used for comorbidity network construction. In the comorbidity network, each node represents a disease, and each directed edge from a pre-existing disease to a post-infection disease indicates a comorbidity pattern. The difference of Pearson correlation coefficients and relative risk for each comorbidity pattern between treatment group and control group are considered as additional attributes for edges.

Relative Risk denoted by *RR*_*ij*_ represents the relative risk between disease *i* and disease j. *C*_*ij*_ is the number of co-occurrence incidences of disease *i* and disease j, *I*_*i*_ is the number of incidences of disease *i, I*_*j*_ is the number of incidences of disease j, and *N* is the number of patients included in the dataset (for the control group and treatment group).

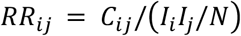

Pearson correlation is another common method to evaluate the strength of diseases’ connection. The Pearson correlation between disease *i* and disease j is denoted by *ϕ*_*i*j_, and the formula is the following:

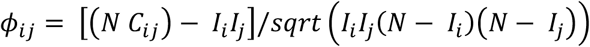

### Protein-Protein Interaction (PPI) network and SARS-CoV-2 human proteins

The protein-protein interaction network used in this study was assembled from 21 public databases by Barabási^35^. The final interactome used in our study contains 18,505 proteins and 327,924 interactions between them. For SARS-CoV-2 human proteins, we used related data detected by Gordon^38^. To quantify the distance between the post-infection and pre-existing diseases, we utilized node-node distances for each protein pair on the largest connected component of the protein-protein interaction network, which contains 18446 nodes and 327868 edges. All proteins are represented by their encoded genes (Entrez ID and Symbol ID).

### ICD code and Gene/Protein association data

Data were derived from the DisGeNET database and the OMIM dataset. These datasets encompass information about proteins and diseases, as well as their interrelationships. Utilizing these datasets, we were able to identify proteins associated with each disease. The distances between diseases were determined based on the distances between their associated proteins. From the DisGeNET database and the OMIM dataset, we assigned associated proteins to 537 diseases, each disease identified by an ICD-9 code and each disease-associated protein identified by its encoded gene (Entrez ID and Symbol ID).

### Distance Measure

Disease-protein relations are utilized to map each disease, denoted by an ICD-9 code, to the Protein-Protein Interaction (PPI) networks. Diseases are represented by protein groups, which are sets of proteins associated with a specific disease. The topological distance between corresponding protein groups in the PPI networks measures the distance between two diseases. A shorter distance between the protein groups indicates a closer relation between the diseases.

For example, if we have a pre-existing disease A and a post-infection disease B, we can map them to the PPI networks and find their corresponding protein groups. Then, we can calculate the distance between them as *d*_*AB*_, where *d*(*a, b*) represents the shortest path from protein a to protein b in the PPI networks.

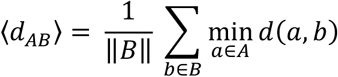

### Testing Method

We employed two statistical tests, the Chi-square test and the Fisher’s exact test, to compare the co-occurrence frequency of disease pairs between the control and treatment groups. These tests can assist us in determining whether a significant association exists between two diseases in the presence or absence of COVID-19.

Among all 537 diseases which have at least one associated protein, for each pre-existing disease, we treated selected comorbidity patterns as positive sample and generated negative sample (disease pairs composed of pre-existing disease and other diseases, which are different from selected comorbidity patterns). Additionally, we utilized the Mann-Whitney U test, to compare the changes of Protein-Protein Interaction (PPI) distance before and after the addition of proteins associated with COVID-19 to proteins associated with pre-existing disease between positive and negative groups. This test can help us evaluate whether a significant difference exists in the PPI distance between the positive and negative sample groups, thereby indicating the impact of COVID-19 on the relationship between two diseases.

### Negative Sample for Z-score Compute

For each pre-existing post-infection disease pair, we conduct a permutation test of 1000 repeats to compute Z-score. We randomly select proteins from the PPI network, and the selected protein groups are required to have similar node degree distribution to pre-existing disease and post-infection disease respectively. Distance differences before and after adding proteins associated with COVID-19 as a part of pre-existing disease associated proteins is also calculated based on the randomly selected protein groups, and the mean value and standardized deviation of the result are used to compute Z-score for each pre-existing post-infection disease pair according to the following formula:

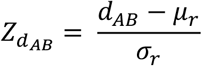

*d*_*AB*_ is the distance change of pre-existing post-infection disease pair before and after adding proteins associated with COVID-19 as a part of pre-existing disease associated proteins. *μ*_*r*_ is the mean value of the distance change from the permutation test, *σ*_*r*_ is the standardized deviation of the distance change from the permutation test.

### GO enrichment Analysis

GO terms describe the functions of gene products across three primary aspects: biological process, molecular function, and cellular component. By conducting GO enrichment analysis, we can pinpoint the GO terms and crucial genes most impacted by COVID-19, thereby enhancing our understanding of the disease’s underlying biology.^18,39–41^. We employed Fisher’s exact test for the enrichment analysis and used the Benjamini-Hochberg procedure to adjust the p-values for multiple testing.^42,43^.

GO Term and Gene Information are from the National Center for Biotechnology Information(NCBI). These datasets include information on GO terms and the relationships between GO terms, genes, and proteins^42^. We can also identify proteins associated with each GO term. By utilizing GO enrichment analysis, we can discover highly influenced GO terms for each disease based on its associated proteins.

### Protein and GO terms Evaluation

The importance of overlapping proteins and GO terms (those associated with both the post-infection disease and pre-existing disease with COVID-19) is the sum of the coefficients of disease pairs they are associated with. We can sort overlapping proteins and GO terms by Relative Risk (RR) and Correlation Coefficient, respectively. The final index of each item is determined by the mean index of those two indices. According to the final index, we can identify important proteins and GO terms.

**Table 1.**
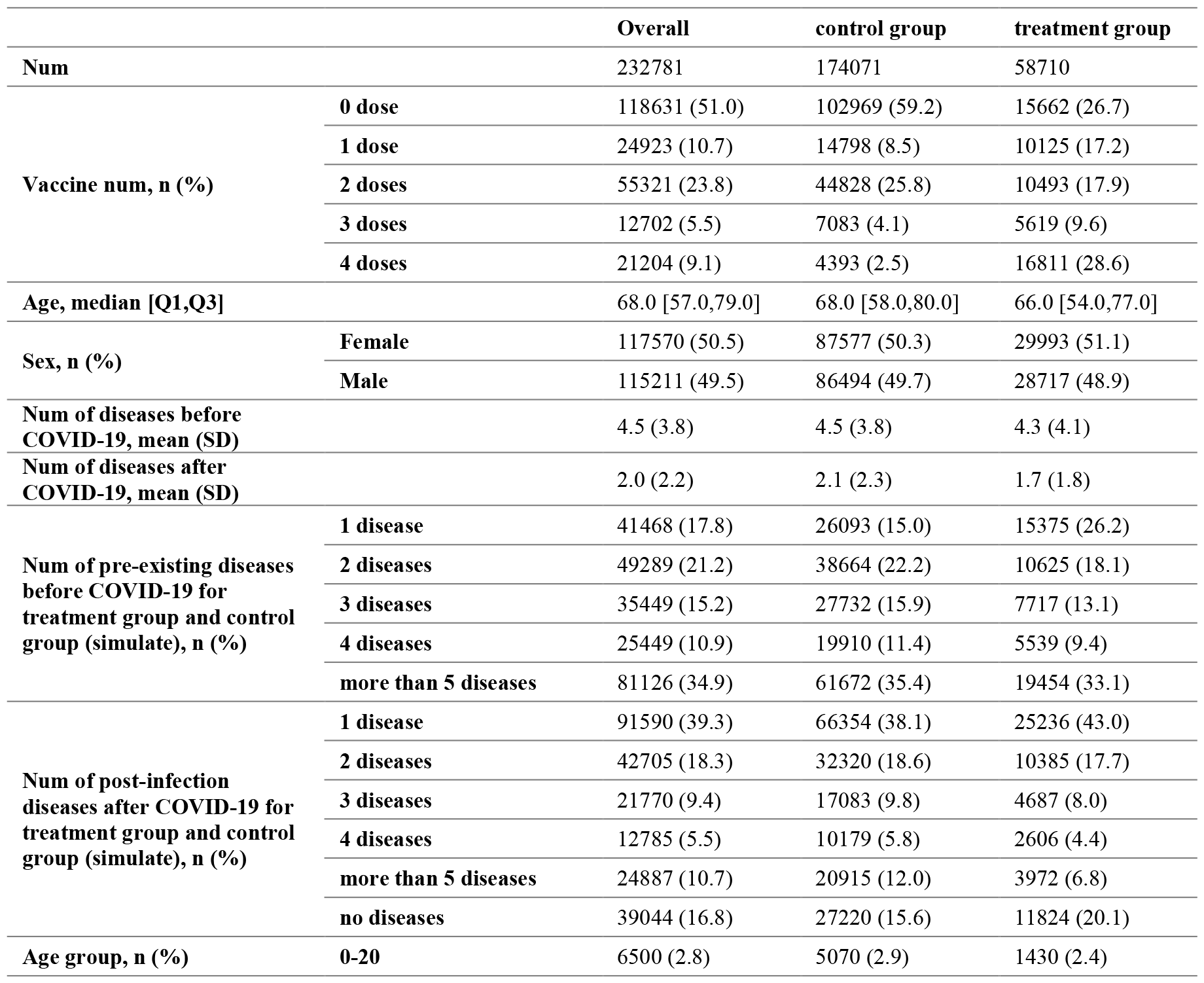
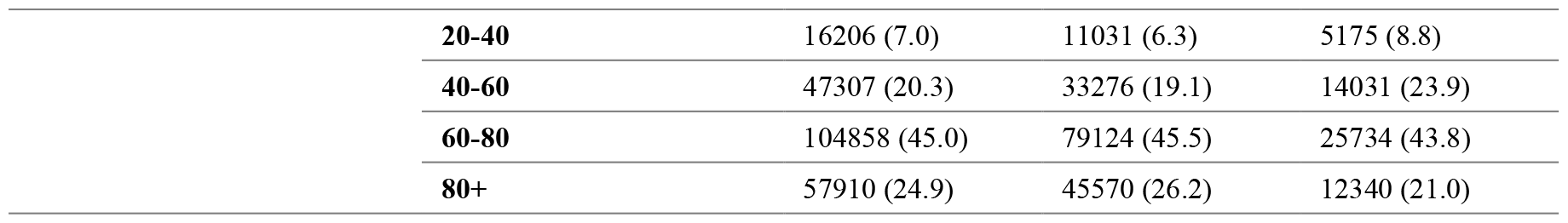
Summary statistics of the dataset.

## References

1. Davis, H. E., McCorkell, L., Vogel, J. M. & Topol, E. J. Long COVID: major findings, mechanisms and recommendations. Nat. Rev. Microbiol. 21, 133–146 (2023).

2. Zhang, H. et al. Data-driven identification of post-acute SARS-CoV-2 infection subphenotypes. Nat. Med. 29, 226–235 (2023).

3. Russell, C. D., Lone, N. I. & Baillie, J. K. Comorbidities, multimorbidity and COVID-19. Nat. Med. 29, 334–343 (2023).

4. Altmann, D. M., Whettlock, E. M., Liu, S., Arachchillage, D. J. & Boyton, R. J. The immunology of long COVID. Nat. Rev. Immunol. 23, 618–634 (2023).

5. Xie, Y., Xu, E., Bowe, B. & Al-Aly, Z. Long-term cardiovascular outcomes of COVID-19. Nat. Med. 28, 583–590 (2022).

6. Raman, B., Bluemke, D. A., Lüscher, T. F. & Neubauer, S. Long COVID: post-acute sequelae of COVID-19 with a cardiovascular focus. Eur. Heart J. 43, 1157–1172 (2022).

7. Xie, Y., Xu, E. & Al-Aly, Z. Risks of mental health outcomes in people with covid-19: cohort study. BMJ 376, e068993 (2022).

8. Xie, Y. & Al-Aly, Z. Risks and burdens of incident diabetes in long COVID: a cohort study. Lancet Diabetes Endocrinol. 10, 311–321 (2022).

9. Bowe, B., Xie, Y., Xu, E. & Al-Aly, Z. Kidney Outcomes in Long COVID. J. Am. Soc. Nephrol. 32, 2851 (2021).

10. Su, Y. et al. Multiple early factors anticipate post-acute COVID-19 sequelae. Cell 185, 881–895.e20 (2022).

11. Brodin, P. et al. Studying severe long COVID to understand post-infectious disorders beyond COVID-19. Nat. Med. 28, 879–882 (2022).

12. Mehandru, S. & Merad, M. Pathological sequelae of long-haul COVID. Nat. Immunol. 23, 194–202 (2022).

13. Lai, F. T. T. et al. Multimorbidity and adverse events of special interest associated with Covid-19 vaccines in Hong Kong. Nat. Commun. 13, 411 (2022).

14. Kuan, V. et al. Identifying and visualising multimorbidity and comorbidity patterns in patients in the English National Health Service: a population-based study. Lancet Digit. Health 5, e16–e27 (2023).

15. Subramanian, A. et al. Symptoms and risk factors for long COVID in non-hospitalized adults. Nat. Med. 28, 1706–1714 (2022).

16. Ayoubkhani, D. et al. Trajectory of long covid symptoms after covid-19 vaccination: community based cohort study. BMJ 377, e069676 (2022).

17. Yang, J., Xu, Z., Wu, W. K. K., Chu, Q. & Zhang, Q. GraphSynergy: a network-inspired deep learning model for anticancer drug combination prediction. J. Am. Med. Inform. Assoc. 28, 2336–2345 (2021).

18. Ashburner, M. et al. Gene Ontology: tool for the unification of biology. Nat. Genet. 25, 25–29 (2000).

19. Xu, Z., Zhang, J., Zhang, Q., Xuan, Q. & Yip, P. S. F. A comorbidity knowledge-aware model for disease prognostic prediction. IEEE Trans. Cybern. 52, 9809–9819 (2021).

20. Borczuk, A. C. & Yantiss, R. K. The pathogenesis of coronavirus-19 disease. J. Biomed. Sci. 29, 87 (2022).

21. van Doorn, A. S., Meijer, B., Frampton, C. M. A., Barclay, M. L. & de Boer, N. K. H. Systematic review with meta-analysis: SARS-CoV-2 stool testing and the potential for faecal-oral transmission. Aliment. Pharmacol. Ther. 52, 1276–1288 (2020).

22. Xiao, F. et al. Evidence for Gastrointestinal Infection of SARS-CoV-2. Gastroenterology 158, 1831–1833.e3 (2020).

23. Zhou, Y. et al. Network medicine links SARS-CoV-2/COVID-19 infection to brain microvascular injury and neuroinflammation in dementia-like cognitive impairment. Alzheimers Res. Ther. 13, 110 (2021).

24. Yang, D. et al. Targeting intracellular Neu1 for coronavirus infection treatment. iScience 26, 106037 (2023).

25. Satu, M. S. et al. Diseasome and comorbidities complexities of SARS-CoV-2 infection with common malignant diseases. Brief. Bioinform. 22, 1415–1429 (2021).

26. White, E. J. et al. Sialidase down-regulation reduces non-HDL cholesterol, inhibits leukocyte transmigration, and attenuates atherosclerosis in ApoE knockout mice. J. Biol. Chem. 293, 14689–14706 (2018).

27. Zhang, C., Chen, J., Liu, Y. & Xu, D. Sialic acid metabolism as a potential therapeutic target of atherosclerosis. Lipids Health Dis. 18, 173 (2019).

28. Jensen-Cody, S. O. & Potthoff, M. J. Hepatokines and metabolism: Deciphering communication from the liver. Mol. Metab. 44, 101138 (2021).

29. Sekiyama, K., Ushiro, Y., Kurisaki, A., Funaba, M. & Hashimoto, O. Activin E enhances insulin sensitivity and thermogenesis by activating brown/beige adipocytes. J. Vet. Med. Sci. 81, 646–652 (2019).

30. Gu, J. et al. The role of lysosomal membrane proteins in glucose and lipid metabolism. FASEB J. 35, e21848 (2021).

31. Sleat, D. E. et al. Genetic evidence for nonredundant functional cooperativity between NPC1 and NPC2 in lipid transport. Proc. Natl. Acad. Sci. 101, 5886–5891 (2004).

32. Benjamin, D. I. et al. Ether lipid generating enzyme AGPS alters the balance of structural and signaling lipids to fuel cancer pathogenicity. Proc. Natl. Acad. Sci. 110, 14912–14917 (2013).

33. Liang, Y. et al. γ-Linolenic Acid Prevents Lipid Metabolism Disorder in Palmitic Acid-Treated Alpha Mouse Liver-12 Cells by Balancing Autophagy and Apoptosis via the LKB1-AMPK-mTOR Pathway. J. Agric. Food Chem. 69, 8257–8267 (2021).

34. Rosenbaum, P. R. & Rubin, D. B. The central role of the propensity score in observational studies for causal effects. Biometrika 70, 41–55 (1983).

35. Morselli Gysi, D. et al. Network medicine framework for identifying drug-repurposing opportunities for COVID-19. Proc. Natl. Acad. Sci. 118, e2025581118 (2021).

36. Zhao, Q.-Y. et al. Propensity score matching with R: conventional methods and new features. Ann. Transl. Med. 9, 812 (2021).

37. Gomez-Cabrero, D. et al. From comorbidities of chronic obstructive pulmonary disease to identification of shared molecular mechanisms by data integration. BMC Bioinformatics 17, 441 (2016).

38. Gordon, D. E. et al. A SARS-CoV-2 protein interaction map reveals targets for drug repurposing. Nature 583, 459–468 (2020).

39. Klopfenstein, D. V. et al. GOATOOLS: A Python library for Gene Ontology analyses. Sci. Rep. 8, 10872 (2018).

40. Yu, G. et al. GOSemSim: an R package for measuring semantic similarity among GO terms and gene products. Bioinformatics 26, 976–978 (2010).

41. Yu, G. Gene Ontology Semantic Similarity Analysis Using GOSemSim. in Stem Cell Transcriptional Networks: Methods and Protocols (ed. Kidder, B. L.) 207–215 (Springer US, 2020). doi:10.1007/978-1-0716-0301-7_11.

42. Rivals, I., Personnaz, L., Taing, L. & Potier, M.-C. Enrichment or depletion of a GO category within a class of genes: which test? Bioinformatics 23, 401–407 (2007).

43. Benjamini, Y. & Hochberg, Y. Controlling the False Discovery Rate: A Practical and Powerful Approach to Multiple Testing. J. R. Stat. Soc. Ser. B Methodol. 57, 289–300 (1995).

