## Appendix for "Deciphering the Molecular Mechanism of Post-Acute Sequelae of COVID-19 through Comorbidity Network Analysis"

Supplementary table 1: Table One (Statistics about Datasets Before Matching)

Supplementary table 2: Pre-existing disease frequency and post-infection disease frequency

Supplementary table 3: Disease Groups association weight

Supplementary table 4: Details about comorbidity patterns

Supplementary table 5: Z-score about PPI distance

Supplementary table 6: Details about overlapped GO functions

Supplementary table 7: Overall importance for overlapped proteins

Supplementary table 8: GO Functions associated with overlapping proteins

Supplementary Figure 1: Plots about PSM Results

Supplementary Figure 2: Inclusion-exclusion cascade
