## Supplementary Figure 1 for "Deciphering the Molecular Mechanism of Post-Acute Sequelae of COVID-19 through Comorbidity Network Analysis"

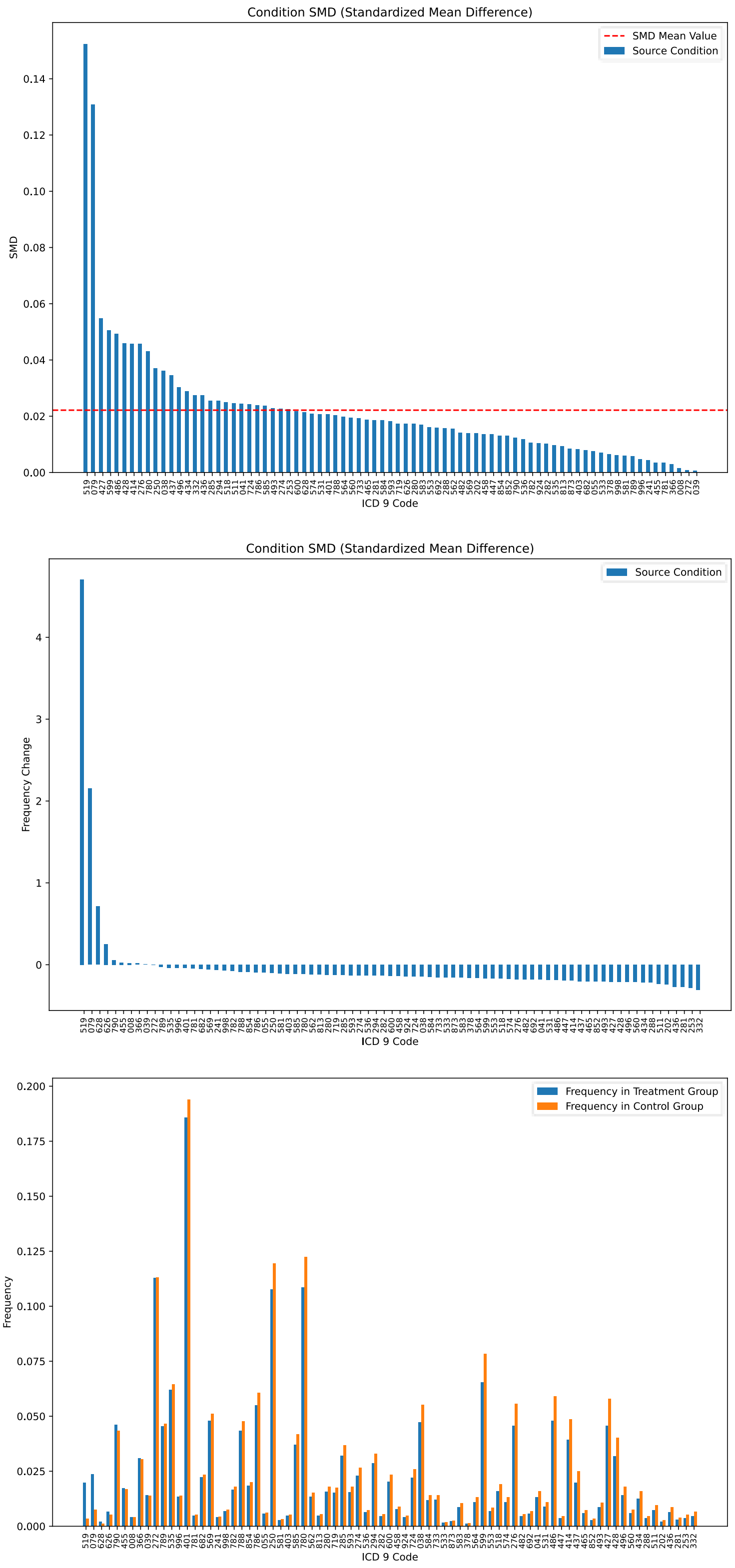


Supplementary Figure 1 illustrates the Standardized Mean Differences (SMD) and frequency of diseases in the treatment and control groups. The upper plot (A) presents a bar plot representing the

SMD result for each disease, with each bar corresponding to a specific disease. The middle plot (B) displays a bar plot of the normalized SMD results across all disease types. The bottom plot (C) display s the frequency of each disease in control group and treatment group.
