## Supplementary Figure 2 for "Deciphering the Molecular Mechanism of Post-Acute Sequelae of COVID-19 through Comorbidity Network Analysis"

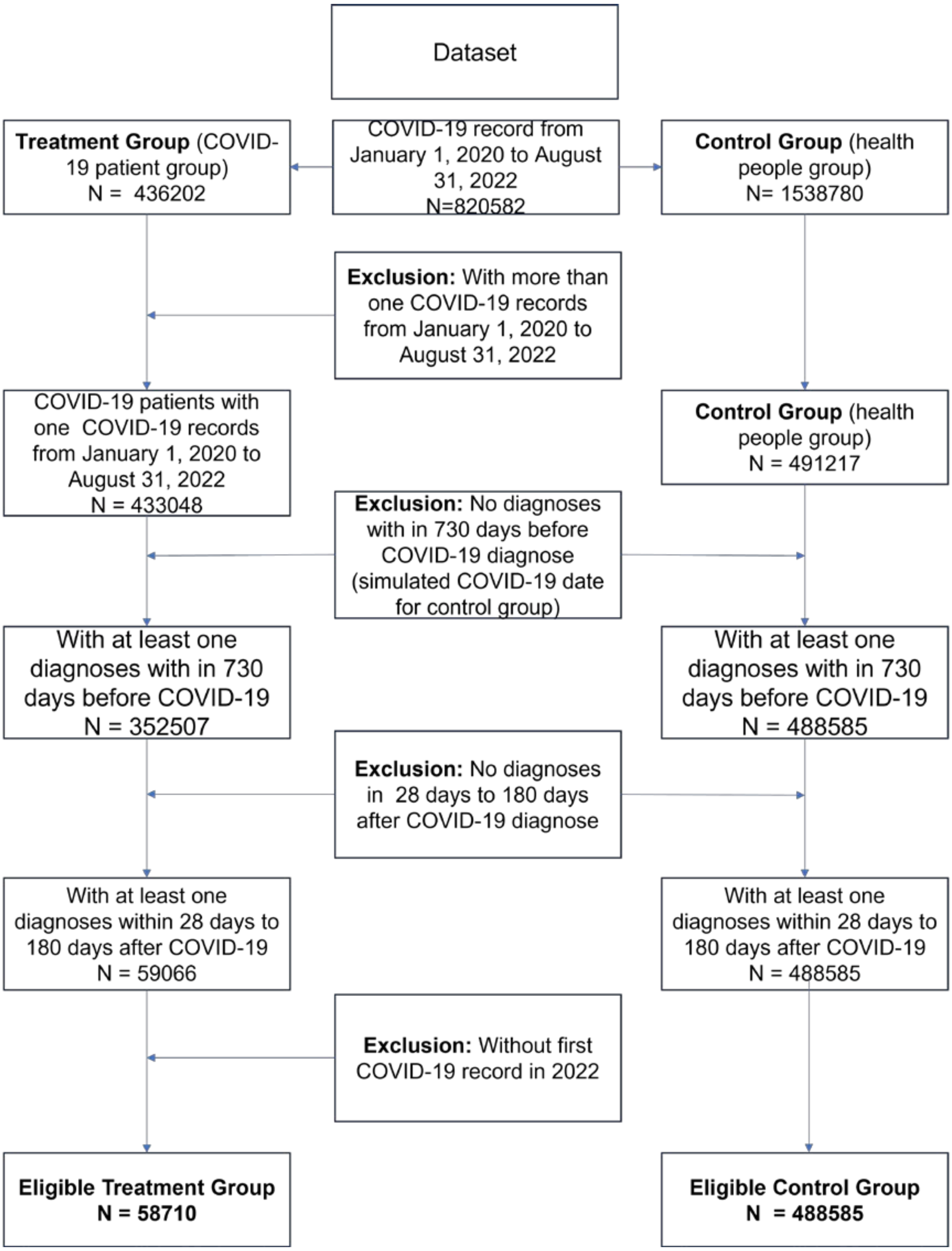


Supplementary Figure 1 Inclusion-exclusion cascade for the dataset Inclusion-exclusion cascades for the dataset
