## Supplementary Table 1 for "Deciphering the Molecular Mechanism of Post-Acute Sequelae of COVID-19 through Comorbidity Network Analysis"

| Table One (Statistics about Datasets Before Matching) | | | | |
|---|---|---|---|---|
| | | **Overall** | **control group** | **treatment group** |
| **Num** | | 547295 | 488585 | 58710 |
| **Vaccine num, n (%)** | **0 dose** | 378728 (69.2) | 363066 (74.3) | 15662 (26.7) |
| | **1 dose** | 28426 (5.2) | 18301 (3.7) | 10125 (17.2) |
| | **2 doses** | 104182 (19.0) | 93689 (19.2) | 10493 (17.9) |
| | **3 doses** | 13933 (2.5) | 8314 (1.7) | 5619 (9.6) |
| | **4 doses** | 22026 (4.0) | 5215 (1.1) | 16811 (28.6) |
| **Age, median [Q1,Q3]** | | 66.0 [54.0,77.0] | 66.0 [54.0,76.0] | 66.0 [54.0,77.0] |
| **Sex, n (%)** | **Female** | 292682 (53.5) | 262689 (53.8) | 29993 (51.1) |
| | **Male** | 254613 (46.5) | 225896 (46.2) | 28717 (48.9) |
| **Number of diseases before index date, mean (SD)** | | 3.3 (3.2) | 3.2 (3.1) | 4.3 (4.1) |
| **Number of new diseases after index date, mean (SD)** | | 2.0 (2.1) | 2.0 (2.2) | 1.7 (1.8) |
| **num of diseases before COVID infections for treatment group and control group (simulate), n (%)** | **1 disease** | 193190 (35.3) | 177815 (36.4) | 15375 (26.2) |
| | **2 diseases** | 115496 (21.1) | 104871 (21.5) | 10625 (18.1) |
| | **3 diseases** | 71466 (13.1) | 63749 (13.0) | 7717 (13.1) |
| | **4 diseases** | 46235 (8.4) | 40696 (8.3) | 5539 (9.4) |
| | **more than 5 diseases** | 120908 (22.1) | 101454 (20.8) | 19454 (33.1) |
| **num of new diseases after COVID infections for treatment group and control group (simulate), n (%)** | **one disease** | 235686 (43.1) | 210450 (43.1) | 25236 (43.0) |
| | **2 diseases** | 104670 (19.1) | 94285 (19.3) | 10385 (17.7) |
| | **3 diseases** | 50045 (9.1) | 45358 (9.3) | 4687 (8.0) |
| | **4 diseases** | 27866 (5.1) | 25260 (5.2) | 2606 (4.4) |
| | **more than 5 diseases** | 52691 (9.6) | 48719 (10.0) | 3972 (6.8) |
| | **no diseases** | 76337 (13.9) | 64513 (13.2) | 11824 (20.1) |

Supplementary table 1 illustrates the statistics of the datasets: control group and treatment group.
