## Supplementary Table 2 for "Deciphering the Molecular Mechanism of Post-Acute Sequelae of COVID-19 through Comorbidity Network Analysis"

| Conditions | Count Num Source | Count Num(Weighted Corr) Source | Count Num(Weighted RR) Source | Count Num Target | Count Num(Weighted Corr) Target | Count Num(Weighted RR) Target | Detail | Detail_extend |
|---|---|---|---|---|---|---|---|---|
| ICD 008 | 1 | 0.011 | 1.093 | 2 | 0.043 | 3.097 | Intestinal infections due to other organisms | 001-139:Infectious And Parasitic Diseases |
| ICD 009 | 0 | 0 | 0 | 1 | 0.008 | 1.367 | Ill-defined intestinal infections | 001-139:Infectious And Parasitic Diseases |
| ICD 011 | 0 | 0 | 0 | 1 | 0.018 | 2.595 | Pulmonary tuberculosis | 001-139:Infectious And Parasitic Diseases |
| ICD 036 | 0 | 0 | 0 | 1 | 0.02 | 3.248 | Meningococcal infection | 001-139:Infectious And Parasitic Diseases |
| ICD 037 | 0 | 0 | 0 | 2 | 0.03 | 4.545 | Tetanus | 001-139:Infectious And Parasitic Diseases |
| ICD 038 | 2 | 0.059 | 2.5 | 2 | 0.076 | 5.186 | Septicemia | 001-139:Infectious And Parasitic Diseases |
| ICD 041 | 1 | 0.013 | 1.247 | 2 | 0.046 | 3.956 | Bacterial infection in conditions classified elsewhere and of unspecified site | 001-139:Infectious And Parasitic Diseases |
| ICD 079 | 56 | 1.13 | 202.939 | 73 | 1.156 | 147.461 | Viral and chlamydial infection in conditions classified elsewhere and of unspecified site | 001-139:Infectious And Parasitic Diseases |
| ICD 093 | 0 | 0 | 0 | 2 | 0.041 | 3.298 | Cardiovascular syphilis | 001-139:Infectious And Parasitic Diseases |
| ICD 096 | 0 | 0 | 0 | 2 | 0.036 | 4.198 | Late syphilis, latent | 001-139:Infectious And Parasitic Diseases |
| ICD 197 | 2 | 0.018 | 6.281 | 2 | 0.017 | 4.788 | Secondary malignant neoplasm of respiratory and digestive systems | 140-239:Neoplasms |
| ICD 198 | 0 | 0 | 0 | 2 | 0.021 | 6.152 | Secondary malignant neoplasm of other specified sites | 140-239:Neoplasms |
| ICD 250 | 3 | 0.059 | 5.146 | 1 | 0.006 | 1.521 | Diabetes mellitus | 240-279:Endocrine, Nutritional And Metabolic Diseases, And Immunity Disorders |
| ICD 251 | 0 | 0 | 0 | 3 | 0.076 | 15.404 | Other disorders of pancreatic internal secretion | 240-279:Endocrine, Nutritional And Metabolic Diseases, And Immunity Disorders |
| ICD 266 | 0 | 0 | 0 | 1 | 0.014 | 3.253 | Deficiency of b-complex components | 240-279:Endocrine, Nutritional And Metabolic Diseases, And Immunity Disorders |
| ICD 272 | 4 | 0.061 | 7.357 | 1 | 0.044 | 1.295 | Disorders of lipoid metabolism | 240-279:Endocrine, Nutritional And Metabolic Diseases, And Immunity Disorders |
| ICD 274 | 2 | 0.058 | 5.141 | 2 | 0.016 | 4.805 | Gout | 240-279:Endocrine, Nutritional And Metabolic Diseases, And Immunity Disorders |
| ICD 276 | 2 | 0.049 | 2.362 | 2 | 0.084 | 7.098 | Disorders of fluid electrolyte and acid-base balance | 240-279:Endocrine, Nutritional And Metabolic Diseases, And Immunity Disorders |
| ICD 278 | 0 | 0 | 0 | 3 | 0.054 | 3.576 | Overweight, obesity and other hyperalimentation | 240-279:Endocrine, Nutritional And Metabolic Diseases, And Immunity Disorders |
| ICD 280 | 1 | 0.01 | 1.396 | 2 | 0.041 | 5.483 | Iron deficiency anemias | 280-289:Diseases Of The Blood And Blood-Forming Organs |
| ICD 285 | 2 | 0.055 | 3.118 | 2 | 0.057 | 5.126 | Other and unspecified anemias | 280-289:Diseases Of The Blood And Blood-Forming Organs |
| ICD 290 | 0 | 0 | 0 | 2 | 0.042 | 23.114 | Dementias | 290-319:Mental Disorders |
| ICD 294 | 1 | 0.019 | 1.016 | 2 | 0.045 | 7.112 | Persistent mental disorders due to conditions classified elsewhere | 290-319:Mental Disorders |
| ICD 309 | 0 | 0 | 0 | 1 | 0.017 | 1.667 | Adjustment reaction | 290-319:Mental Disorders |
| ICD 345 | 0 | 0 | 0 | 1 | 0.01 | 1.41 | Epilepsy and recurrent seizures | 320-389:Diseases Of The Nervous System And Sense Organs |
| ICD 386 | 0 | 0 | 0 | 1 | 0.015 | 2.224 | Vertiginous syndromes and other disorders of vestibular system | 320-389:Diseases Of The Nervous System And Sense Organs |
| ICD 401 | 6 | 0.122 | 9.143 | 2 | 0.018 | 2.765 | Essential hypertension | 390-459:Diseases Of The Circulatory System |
| ICD 410 | 2 | 0.014 | 3.319 | 2 | 0.058 | 12.308 | Acute myocardial infarction | 390-459:Diseases Of The Circulatory System |
| ICD 414 | 2 | 0.031 | 3.49 | 2 | 0.04 | 6.097 | Other forms of chronic ischemic heart disease | 390-459:Diseases Of The Circulatory System |
| ICD 427 | 2 | 0.041 | 2.971 | 2 | 0.039 | 4.059 | Cardiac dysrhythmias | 390-459:Diseases Of The Circulatory System |
| ICD 428 | 2 | 0.061 | 4.188 | 3 | 0.08 | 7.552 | Heart failure | 390-459:Diseases Of The Circulatory System |
| ICD 436 | 2 | 0.014 | 2.646 | 2 | 0.034 | 7.716 | Acute, but ill-defined, cerebrovascular disease | 390-459:Diseases Of The Circulatory System |
| ICD 437 | 3 | 0.058 | 5.804 | 2 | 0.058 | 16.979 | Other and ill-defined cerebrovascular disease | 390-459:Diseases Of The Circulatory System |
| ICD 438 | 0 | 0 | 0 | 2 | 0.046 | 6.959 | Late effects of cerebrovascular disease | 390-459:Diseases Of The Circulatory System |
| ICD 458 | 2 | 0.042 | 4.108 | 2 | 0.043 | 6.52 | Hypotension | 390-459:Diseases Of The Circulatory System |
| ICD 480 | 0 | 0 | 0 | 1 | 0.012 | 2.173 | Viral pneumonia | 460-519:Diseases Of The Respiratory System |
| ICD 482 | 1 | 0.006 | 1.005 | 2 | 0.081 | 5.144 | Other bacterial pneumonia | 460-519:Diseases Of The Respiratory System |
| ICD 486 | 3 | 0.097 | 5.283 | 2 | 0.14 | 7.308 | Pneumonia, organism unspecified | 460-519:Diseases Of The Respiratory System |
| ICD 496 | 2 | 0.061 | 4.781 | 2 | 0.018 | 5.435 | Chronic airway obstruction, not elsewhere classified | 460-519:Diseases Of The Respiratory System |
| ICD 507 | 0 | 0 | 0 | 1 | 0.013 | 1.811 | Pneumonitis due to solids and liquids | 460-519:Diseases Of The Respiratory System |
| ICD 511 | 2 | 0.036 | 5.837 | 2 | 0.031 | 13.393 | Pleurisy | 460-519:Diseases Of The Respiratory System |
| ICD 518 | 2 | 0.036 | 3.331 | 2 | 0.046 | 3.987 | Other diseases of lung | 460-519:Diseases Of The Respiratory System |
| ICD 519 | 50 | 1.027 | 125.883 | 63 | 0.962 | 113.954 | Other diseases of respiratory system | 460-519:Diseases Of The Respiratory System |
| ICD 535 | 4 | 0.04 | 5.894 | 4 | 0.07 | 8.783 | Gastritis and duodenitis | 520-579:Diseases Of The Digestive System |
| ICD 553 | 2 | 0.026 | 6.248 | 1 | 0.011 | 2.842 | Other hernia of abdominal cavity without mention of obstruction or gangrene | 520-579:Diseases Of The Digestive System |
| ICD 564 | 2 | 0.026 | 2.412 | 2 | 0.024 | 6.364 | Functional digestive disorders not elsewhere classified | 520-579:Diseases Of The Digestive System |
| ICD 569 | 2 | 0.023 | 2.922 | 2 | 0.012 | 3.039 | Other disorders of intestine | 520-579:Diseases Of The Digestive System |
| ICD 573 | 2 | 0.008 | 2.424 | 2 | 0.033 | 8.839 | Other disorders of liver | 520-579:Diseases Of The Digestive System |
| ICD 578 | 0 | 0 | 0 | 2 | 0.036 | 7.244 | Gastrointestinal hemorrhage | 520-579:Diseases Of The Digestive System |
| ICD 583 | 2 | 0.014 | 2.552 | 1 | 0.022 | 5.88 | Nephritis and nephropathy not specified as acute or chronic | 580-629:Diseases Of The Genitourinary System |
| ICD 584 | 3 | 0.076 | 6.564 | 2 | 0.046 | 5.977 | Acute kidney failure | 580-629:Diseases Of The Genitourinary System |
| ICD 585 | 2 | 0.095 | 6.045 | 2 | 0.023 | 5.016 | Chronic kidney disease (ckd) | 580-629:Diseases Of The Genitourinary System |
| ICD 599 | 1 | 0.021 | 1.065 | 2 | 0.107 | 5.974 | Other disorders of urethra and urinary tract | 580-629:Diseases Of The Genitourinary System |
| ICD 626 | 1 | 0.01 | 1.584 | 1 | 0.01 | 1.612 | Disorders of menstruation and other abnormal bleeding from female genital tract | 580-629:Diseases Of The Genitourinary System |
| ICD 682 | 1 | 0.012 | 1.199 | 2 | 0.024 | 3.699 | Other cellulitis and abscess | 680-709:Diseases Of The Skin And Subcutaneous Tissue |
| ICD 707 | 0 | 0 | 0 | 2 | 0.073 | 7.861 | Chronic ulcer of skin | 680-709:Diseases Of The Skin And Subcutaneous Tissue |
| ICD 780 | 1 | 0.017 | 1.051 | 3 | 0.064 | 6.975 | General symptoms | 780-799:Symptoms, Signs, And Ill-Defined Conditions |
| ICD 782 | 2 | 0.012 | 2.16 | 1 | 0.007 | 1.482 | Symptoms involving skin and other integumentary tissue | 780-799:Symptoms, Signs, And Ill-Defined Conditions |
| ICD 783 | 0 | 0 | 0 | 5 | 0.102 | 13.737 | Symptoms concerning nutrition metabolism and development | 780-799:Symptoms, Signs, And Ill-Defined Conditions |
| ICD 785 | 1 | 0.003 | 1.111 | 3 | 0.043 | 5.78 | Symptoms involving cardiovascular system | 780-799:Symptoms, Signs, And Ill-Defined Conditions |
| ICD 786 | 1 | 0.009 | 1.165 | 2 | 0.048 | 2.999 | Symptoms involving respiratory system and other chest symptoms | 780-799:Symptoms, Signs, And Ill-Defined Conditions |
| ICD 787 | 0 | 0 | 0 | 2 | 0.032 | 3.565 | Symptoms involving digestive system | 780-799:Symptoms, Signs, And Ill-Defined Conditions |
| ICD 788 | 3 | 0.047 | 4.988 | 3 | 0.061 | 9.752 | Symptoms involving urinary system | 780-799:Symptoms, Signs, And Ill-Defined Conditions |
| ICD 789 | 1 | 0.017 | 1.667 | 3 | 0.03 | 5.886 | Other symptoms involving abdomen and pelvis | 780-799:Symptoms, Signs, And Ill-Defined Conditions |
| ICD 793 | 1 | 0.005 | 1.422 | 1 | 0.014 | 7.035 | Nonspecific (abnormal) findings on radiological and other examination of body structure | 780-799:Symptoms, Signs, And Ill-Defined Conditions |
| ICD 799 | 0 | 0 | 0 | 2 | 0.053 | 5.925 | Other ill-defined and unknown causes of morbidity and mortality | 780-799:Symptoms, Signs, And Ill-Defined Conditions |
| ICD 822 | 0 | 0 | 0 | 1 | 0.018 | 3.448 | Fracture of patella | 800-999:Injury And Poisoning |
| ICD 854 | 2 | 0.031 | 2.485 | 2 | 0.016 | 3.971 | Intracranial injury of other and unspecified nature | 800-999:Injury And Poisoning |
| ICD 883 | 0 | 0 | 0 | 2 | 0.021 | 4.52 | Open wound of finger(s) | 800-999:Injury And Poisoning |
| ICD 924 | 1 | 0.006 | 1.027 | 2 | 0.03 | 4.573 | Contusion of lower limb and of other and unspecified sites | 800-999:Injury And Poisoning |
| ICD 996 | 2 | 0.043 | 3.456 | 3 | 0.087 | 14.703 | Complications peculiar to certain specified procedures | 800-999:Injury And Poisoning |
| ICD 998 | 1 | 0.017 | 3.457 | 1 | 0.016 | 2.981 | Other complications of procedures not elsewhere classified | 800-999:Injury And Poisoning |
| ICD 039 | 2 | 0.093 | 10.743 | 0 | 0 | 0 | Actinomycotic infections | 001-139:Infectious And Parasitic Diseases |
| ICD 055 | 1 | 0.012 | 2.545 | 0 | 0 | 0 | Measles | 001-139:Infectious And Parasitic Diseases |
| ICD 162 | 2 | 0.018 | 6.641 | 0 | 0 | 0 | Malignant neoplasm of trachea bronchus and lung | 140-239:Neoplasms |
| ICD 202 | 2 | 0.064 | 18.769 | 0 | 0 | 0 | Other malignant neoplasms of lymphoid and histiocytic tissue | 140-239:Neoplasms |
| ICD 241 | 1 | 0.019 | 3.445 | 0 | 0 | 0 | Nontoxic nodular goiter | 240-279:Endocrine, Nutritional And Metabolic Diseases, And Immunity Disorders |
| ICD 244 | 2 | 0.014 | 3.18 | 0 | 0 | 0 | Acquired hypothyroidism | 240-279:Endocrine, Nutritional And Metabolic Diseases, And Immunity Disorders |
| ICD 253 | 1 | 0.011 | 1.64 | 0 | 0 | 0 | Disorders of the pituitary gland and its hypothalamic control | 240-279:Endocrine, Nutritional And Metabolic Diseases, And Immunity Disorders |
| ICD 281 | 2 | 0.051 | 5.135 | 0 | 0 | 0 | Other deficiency anemias | 280-289:Diseases Of The Blood And Blood-Forming Organs |
| ICD 282 | 1 | 0.01 | 1.48 | 0 | 0 | 0 | Hereditary hemolytic anemias | 280-289:Diseases Of The Blood And Blood-Forming Organs |
| ICD 288 | 1 | 0.015 | 2.174 | 0 | 0 | 0 | Diseases of white blood cells | 280-289:Diseases Of The Blood And Blood-Forming Organs |
| ICD 332 | 2 | 0.033 | 3.758 | 0 | 0 | 0 | Parkinson's disease | 320-389:Diseases Of The Nervous System And Sense Organs |
| ICD 362 | 2 | 0.011 | 2.946 | 0 | 0 | 0 | Other retinal disorders | 320-389:Diseases Of The Nervous System And Sense Organs |
| ICD 366 | 3 | 0.053 | 5.073 | 0 | 0 | 0 | Cataract | 320-389:Diseases Of The Nervous System And Sense Organs |
| ICD 378 | 1 | 0.019 | 3.968 | 0 | 0 | 0 | Strabismus and other disorders of binocular eye movements | 320-389:Diseases Of The Nervous System And Sense Organs |
| ICD 403 | 2 | 0.029 | 2.548 | 0 | 0 | 0 | Hypertensive chronic kidney disease | 390-459:Diseases Of The Circulatory System |
| ICD 434 | 1 | 0.013 | 1.359 | 0 | 0 | 0 | Occlusion of cerebral arteries | 390-459:Diseases Of The Circulatory System |
| ICD 447 | 1 | 0.029 | 5.414 | 0 | 0 | 0 | Other disorders of arteries and arterioles | 390-459:Diseases Of The Circulatory System |
| ICD 455 | 1 | 0.016 | 2.981 | 0 | 0 | 0 | Hemorrhoids | 390-459:Diseases Of The Circulatory System |
| ICD 465 | 1 | 0.006 | 1.054 | 0 | 0 | 0 | Acute upper respiratory infections of multiple or unspecified sites | 460-519:Diseases Of The Respiratory System |
| ICD 493 | 2 | 0.026 | 3.164 | 0 | 0 | 0 | Asthma | 460-519:Diseases Of The Respiratory System |
| ICD 530 | 2 | 0.009 | 2.365 | 0 | 0 | 0 | Diseases of esophagus | 520-579:Diseases Of The Digestive System |
| ICD 531 | 2 | 0.031 | 5.999 | 0 | 0 | 0 | Gastric ulcer | 520-579:Diseases Of The Digestive System |
| ICD 533 | 1 | 0.044 | 4.072 | 0 | 0 | 0 | Peptic ulcer site unspecified | 520-579:Diseases Of The Digestive System |
| ICD 536 | 2 | 0.026 | 6.12 | 0 | 0 | 0 | Disorders of function of stomach | 520-579:Diseases Of The Digestive System |
| ICD 537 | 2 | 0.023 | 6.46 | 0 | 0 | 0 | Other disorders of stomach and duodenum | 520-579:Diseases Of The Digestive System |
| ICD 560 | 2 | 0.032 | 5.365 | 0 | 0 | 0 | Intestinal obstruction without mention of hernia | 520-579:Diseases Of The Digestive System |
| ICD 562 | 1 | 0.02 | 3.248 | 0 | 0 | 0 | Diverticula of intestine | 520-579:Diseases Of The Digestive System |
| ICD 571 | 1 | 0.003 | 1.236 | 0 | 0 | 0 | Chronic liver disease and cirrhosis | 520-579:Diseases Of The Digestive System |
| ICD 574 | 2 | 0.016 | 2.819 | 0 | 0 | 0 | Cholelithiasis | 520-579:Diseases Of The Digestive System |
| ICD 576 | 1 | 0.005 | 1.281 | 0 | 0 | 0 | Other disorders of biliary tract | 520-579:Diseases Of The Digestive System |
| ICD 581 | 1 | 0.022 | 5.9 | 0 | 0 | 0 | Nephrotic syndrome | 580-629:Diseases Of The Genitourinary System |
| ICD 593 | 2 | 0.014 | 2.507 | 0 | 0 | 0 | Other disorders of kidney and ureter | 580-629:Diseases Of The Genitourinary System |
| ICD 600 | 2 | 0.039 | 4.26 | 0 | 0 | 0 | Hyperplasia of prostate | 580-629:Diseases Of The Genitourinary System |
| ICD 628 | 1 | 0.032 | 2.043 | 0 | 0 | 0 | Female infertility | 580-629:Diseases Of The Genitourinary System |
| ICD 692 | 2 | 0.023 | 3.377 | 0 | 0 | 0 | Contact dermatitis and other eczema | 680-709:Diseases Of The Skin And Subcutaneous Tissue |
| ICD 719 | 3 | 0.044 | 5.525 | 0 | 0 | 0 | Other and unspecified disorders of joint | 710-739:Diseases Of The Musculoskeletal System And Connective Tissue |
| ICD 724 | 2 | 0.013 | 2.138 | 0 | 0 | 0 | Other and unspecified disorders of back | 710-739:Diseases Of The Musculoskeletal System And Connective Tissue |
| ICD 729 | 2 | 0.01 | 2.323 | 0 | 0 | 0 | Other disorders of soft tissues | 710-739:Diseases Of The Musculoskeletal System And Connective Tissue |
| ICD 733 | 2 | 0.02 | 2.965 | 0 | 0 | 0 | Other disorders of bone and cartilage | 710-739:Diseases Of The Musculoskeletal System And Connective Tissue |
| ICD 781 | 2 | 0.026 | 4.367 | 0 | 0 | 0 | Symptoms involving nervous and musculoskeletal systems | 780-799:Symptoms, Signs, And Ill-Defined Conditions |
| ICD 784 | 1 | 0.002 | 1.19 | 0 | 0 | 0 | Symptoms involving head and neck | 780-799:Symptoms, Signs, And Ill-Defined Conditions |
| ICD 790 | 4 | 0.054 | 7.015 | 0 | 0 | 0 | Nonspecific findings on examination of blood | 780-799:Symptoms, Signs, And Ill-Defined Conditions |
| ICD 813 | 1 | 0.02 | 4.012 | 0 | 0 | 0 | Fracture of radius and ulna | 800-999:Injury And Poisoning |
| ICD 852 | 2 | 0.032 | 3.75 | 0 | 0 | 0 | Subarachnoid subdural and extradural hemorrhage following injury | 800-999:Injury And Poisoning |
| ICD 873 | 1 | 0.032 | 4.964 | 0 | 0 | 0 | Other open wound of head | 800-999:Injury And Poisoning |
