## Supplementary Table 3 for "Deciphering the Molecular Mechanism of Post-Acute Sequelae of COVID-19 through Comorbidity Network Analysis"

| source | target | Frequency | Diff_Corr | Diff_RR |
|---|---|---|---|---|
| 001-139:Infectious And Parasitic Diseases | 001-139:Infectious And Parasitic Diseases | 9 | 0.214 | 22.871 |
| 001-139:Infectious And Parasitic Diseases | 140-239:Neoplasms | 2 | 0.024 | 8.703 |
| 001-139:Infectious And Parasitic Diseases | 240-279:Endocrine, Nutritional And Metabolic Diseases, And Immunity Disorders | 4 | 0.083 | 19.448 |
| 001-139:Infectious And Parasitic Diseases | 280-289:Diseases Of The Blood And Blood-Forming Organs | 2 | 0.056 | 6.743 |
| 001-139:Infectious And Parasitic Diseases | 290-319:Mental Disorders | 2 | 0.043 | 22.985 |
| 001-139:Infectious And Parasitic Diseases | 320-389:Diseases Of The Nervous System And Sense Organs | 1 | 0.01 | 1.41 |
| 001-139:Infectious And Parasitic Diseases | 390-459:Diseases Of The Circulatory System | 9 | 0.193 | 34.868 |
| 001-139:Infectious And Parasitic Diseases | 460-519:Diseases Of The Respiratory System | 10 | 0.281 | 37.294 |
| 001-139:Infectious And Parasitic Diseases | 520-579:Diseases Of The Digestive System | 6 | 0.066 | 13.254 |
| 001-139:Infectious And Parasitic Diseases | 580-629:Diseases Of The Genitourinary System | 4 | 0.109 | 14.862 |
| 001-139:Infectious And Parasitic Diseases | 680-709:Diseases Of The Skin And Subcutaneous Tissue | 2 | 0.043 | 5.559 |
| 001-139:Infectious And Parasitic Diseases | 780-799:Symptoms, Signs, And Ill-Defined Conditions | 10 | 0.163 | 25.346 |
| 001-139:Infectious And Parasitic Diseases | 800-999:Injury And Poisoning | 2 | 0.033 | 7.722 |
| 140-239:Neoplasms | 001-139:Infectious And Parasitic Diseases | 3 | 0.052 | 14.976 |
| 140-239:Neoplasms | 460-519:Diseases Of The Respiratory System | 3 | 0.047 | 16.715 |
| 240-279:Endocrine, Nutritional And Metabolic Diseases, And Immunity Disorders | 001-139:Infectious And Parasitic Diseases | 6 | 0.113 | 10.573 |
| 240-279:Endocrine, Nutritional And Metabolic Diseases, And Immunity Disorders | 240-279:Endocrine, Nutritional And Metabolic Diseases, And Immunity Disorders | 1 | 0.017 | 1.151 |
| 240-279:Endocrine, Nutritional And Metabolic Diseases, And Immunity Disorders | 460-519:Diseases Of The Respiratory System | 5 | 0.094 | 7.308 |
| 240-279:Endocrine, Nutritional And Metabolic Diseases, And Immunity Disorders | 780-799:Symptoms, Signs, And Ill-Defined Conditions | 1 | 0.019 | 3.445 |
| 240-279:Endocrine, Nutritional And Metabolic Diseases, And Immunity Disorders | 800-999:Injury And Poisoning | 2 | 0.029 | 5.794 |
| 280-289:Diseases Of The Blood And Blood-Forming Organs | 001-139:Infectious And Parasitic Diseases | 4 | 0.071 | 6.983 |
| 280-289:Diseases Of The Blood And Blood-Forming Organs | 460-519:Diseases Of The Respiratory System | 3 | 0.07 | 6.32 |
| 290-319:Mental Disorders | 001-139:Infectious And Parasitic Diseases | 1 | 0.019 | 1.016 |
| 320-389:Diseases Of The Nervous System And Sense Organs | 001-139:Infectious And Parasitic Diseases | 3 | 0.039 | 5.442 |
| 320-389:Diseases Of The Nervous System And Sense Organs | 390-459:Diseases Of The Circulatory System | 1 | 0.024 | 1.884 |
| 320-389:Diseases Of The Nervous System And Sense Organs | 460-519:Diseases Of The Respiratory System | 3 | 0.034 | 4.452 |
| 320-389:Diseases Of The Nervous System And Sense Organs | 780-799:Symptoms, Signs, And Ill-Defined Conditions | 1 | 0.019 | 3.968 |
| 390-459:Diseases Of The Circulatory System | 001-139:Infectious And Parasitic Diseases | 10 | 0.168 | 17.146 |
| 390-459:Diseases Of The Circulatory System | 240-279:Endocrine, Nutritional And Metabolic Diseases, And Immunity Disorders | 2 | 0.069 | 2.559 |
| 390-459:Diseases Of The Circulatory System | 460-519:Diseases Of The Respiratory System | 10 | 0.153 | 15.068 |
| 390-459:Diseases Of The Circulatory System | 780-799:Symptoms, Signs, And Ill-Defined Conditions | 1 | 0.025 | 2.629 |
| 390-459:Diseases Of The Circulatory System | 800-999:Injury And Poisoning | 3 | 0.055 | 10.57 |
| 460-519:Diseases Of The Respiratory System | 001-139:Infectious And Parasitic Diseases | 12 | 0.26 | 23.617 |
| 460-519:Diseases Of The Respiratory System | 140-239:Neoplasms | 2 | 0.015 | 2.237 |
| 460-519:Diseases Of The Respiratory System | 240-279:Endocrine, Nutritional And Metabolic Diseases, And Immunity Disorders | 4 | 0.074 | 9.499 |
| 460-519:Diseases Of The Respiratory System | 280-289:Diseases Of The Blood And Blood-Forming Organs | 2 | 0.043 | 3.865 |
| 460-519:Diseases Of The Respiratory System | 290-319:Mental Disorders | 2 | 0.043 | 7.24 |
| 460-519:Diseases Of The Respiratory System | 390-459:Diseases Of The Circulatory System | 9 | 0.197 | 34.202 |
| 460-519:Diseases Of The Respiratory System | 460-519:Diseases Of The Respiratory System | 11 | 0.263 | 16.755 |
| 460-519:Diseases Of The Respiratory System | 520-579:Diseases Of The Digestive System | 5 | 0.066 | 18.2 |
| 460-519:Diseases Of The Respiratory System | 580-629:Diseases Of The Genitourinary System | 3 | 0.088 | 7.985 |
| 460-519:Diseases Of The Respiratory System | 680-709:Diseases Of The Skin And Subcutaneous Tissue | 2 | 0.053 | 6 |
| 460-519:Diseases Of The Respiratory System | 780-799:Symptoms, Signs, And Ill-Defined Conditions | 8 | 0.138 | 12.469 |
| 460-519:Diseases Of The Respiratory System | 800-999:Injury And Poisoning | 3 | 0.056 | 8.27 |
| 520-579:Diseases Of The Digestive System | 001-139:Infectious And Parasitic Diseases | 13 | 0.149 | 29.007 |
| 520-579:Diseases Of The Digestive System | 460-519:Diseases Of The Respiratory System | 12 | 0.114 | 22.333 |
| 520-579:Diseases Of The Digestive System | 520-579:Diseases Of The Digestive System | 1 | 0.044 | 4.072 |
| 520-579:Diseases Of The Digestive System | 580-629:Diseases Of The Genitourinary System | 1 | 0.01 | 1.612 |
| 520-579:Diseases Of The Digestive System | 800-999:Injury And Poisoning | 1 | 0.014 | 1.842 |
| 580-629:Diseases Of The Genitourinary System | 001-139:Infectious And Parasitic Diseases | 8 | 0.174 | 19.156 |
| 580-629:Diseases Of The Genitourinary System | 240-279:Endocrine, Nutritional And Metabolic Diseases, And Immunity Disorders | 1 | 0.04 | 3.134 |
| 580-629:Diseases Of The Genitourinary System | 460-519:Diseases Of The Respiratory System | 5 | 0.099 | 8.645 |
| 580-629:Diseases Of The Genitourinary System | 520-579:Diseases Of The Digestive System | 1 | 0.01 | 1.584 |
| 680-709:Diseases Of The Skin And Subcutaneous Tissue | 001-139:Infectious And Parasitic Diseases | 2 | 0.02 | 2.373 |
| 680-709:Diseases Of The Skin And Subcutaneous Tissue | 460-519:Diseases Of The Respiratory System | 1 | 0.015 | 2.203 |
| 710-739:Diseases Of The Musculoskeletal System And Connective Tissue | 001-139:Infectious And Parasitic Diseases | 4 | 0.034 | 5.46 |
| 710-739:Diseases Of The Musculoskeletal System And Connective Tissue | 460-519:Diseases Of The Respiratory System | 4 | 0.031 | 4.645 |
| 710-739:Diseases Of The Musculoskeletal System And Connective Tissue | 780-799:Symptoms, Signs, And Ill-Defined Conditions | 1 | 0.021 | 2.846 |
| 780-799:Symptoms, Signs, And Ill-Defined Conditions | 001-139:Infectious And Parasitic Diseases | 9 | 0.101 | 14.002 |
| 780-799:Symptoms, Signs, And Ill-Defined Conditions | 240-279:Endocrine, Nutritional And Metabolic Diseases, And Immunity Disorders | 1 | 0.011 | 1.161 |
| 780-799:Symptoms, Signs, And Ill-Defined Conditions | 290-319:Mental Disorders | 1 | 0.017 | 1.667 |
| 780-799:Symptoms, Signs, And Ill-Defined Conditions | 320-389:Diseases Of The Nervous System And Sense Organs | 1 | 0.015 | 2.224 |
| 780-799:Symptoms, Signs, And Ill-Defined Conditions | 460-519:Diseases Of The Respiratory System | 5 | 0.048 | 7.081 |
| 800-999:Injury And Poisoning | 001-139:Infectious And Parasitic Diseases | 4 | 0.059 | 6.332 |
| 800-999:Injury And Poisoning | 460-519:Diseases Of The Respiratory System | 3 | 0.053 | 4.386 |
| 800-999:Injury And Poisoning | 780-799:Symptoms, Signs, And Ill-Defined Conditions | 3 | 0.068 | 12.432 |
