## Supplementary Table 4 for "Deciphering the Molecular Mechanism of Post-Acute Sequelae of COVID-19 through Comorbidity Network Analysis"

| edge | edge_num_treat | source | target | source_num_treat | target_num_treat | RR_treat | Correlation_treat | edge_num_control | source_num_control | target_num_control | RR_control | Correlation_control | p_value | Diff_RR | Diff_Corr |
|---|---|---|---|---|---|---|---|---|---|---|---|---|---|---|---|
| ["['401']", '272'] | 1144 | 401 | 272 | 10912 | 3393 | 1.814 | 0.096 | 3141 | 33765 | 11560 | 1.401 | 0.052 | 0.026 | 1.295 | 0.044 |
| ["['780']", '079'] | 196 | 780 | 79 | 6375 | 1002 | 1.801 | 0.037 | 202 | 21298 | 963 | 1.714 | 0.02 | 0 | 1.051 | 0.017 |
| ["['493']", '079'] | 20 | 493 | 79 | 504 | 1002 | 2.325 | 0.016 | 13 | 1884 | 963 | 1.247 | 0.002 | 0 | 1.864 | 0.014 |
| ["['493']", '519'] | 20 | 493 | 519 | 504 | 922 | 2.527 | 0.018 | 15 | 1884 | 713 | 1.944 | 0.006 | 0 | 1.3 | 0.012 |
| ["['401']", '278'] | 308 | 401 | 278 | 10912 | 795 | 2.084 | 0.061 | 670 | 33765 | 2094 | 1.65 | 0.035 | 0 | 1.264 | 0.026 |
| ["['079']", '427'] | 51 | 79 | 427 | 1393 | 1210 | 1.776 | 0.018 | 33 | 1310 | 5788 | 0.758 | -0.004 | 0 | 2.345 | 0.021 |
| ["['079']", '428'] | 44 | 79 | 428 | 1393 | 792 | 2.341 | 0.024 | 15 | 1310 | 3108 | 0.641 | -0.004 | 0 | 3.651 | 0.029 |
| ["['079']", '584'] | 30 | 79 | 584 | 1393 | 528 | 2.395 | 0.021 | 12 | 1310 | 2666 | 0.598 | -0.004 | 0 | 4.004 | 0.025 |
| ["['079']", '799'] | 36 | 79 | 799 | 1393 | 606 | 2.504 | 0.024 | 14 | 1310 | 3026 | 0.615 | -0.004 | 0 | 4.073 | 0.028 |
| ["['519']", '427'] | 45 | 519 | 427 | 1157 | 1210 | 1.887 | 0.018 | 22 | 601 | 5788 | 1.101 | 0.001 | 0 | 1.714 | 0.017 |
| ["['519']", '428'] | 44 | 519 | 428 | 1157 | 792 | 2.819 | 0.03 | 15 | 601 | 3108 | 1.398 | 0.003 | 0 | 2.017 | 0.027 |
| ["['519']", '584'] | 29 | 519 | 584 | 1157 | 528 | 2.787 | 0.024 | 13 | 601 | 2666 | 1.412 | 0.003 | 0 | 1.973 | 0.021 |
| ["['519']", '799'] | 36 | 519 | 799 | 1157 | 606 | 3.014 | 0.029 | 17 | 601 | 3026 | 1.627 | 0.005 | 0 | 1.853 | 0.024 |
| ["['272']", '079'] | 136 | 272 | 79 | 6624 | 1002 | 1.203 | 0.01 | 94 | 19683 | 963 | 0.863 | -0.004 | 0 | 1.394 | 0.013 |
| ["['272']", '519'] | 132 | 272 | 519 | 6624 | 922 | 1.269 | 0.012 | 75 | 19683 | 713 | 0.93 | -0.002 | 0 | 1.364 | 0.014 |
| ["['401']", '079'] | 249 | 401 | 79 | 10912 | 1002 | 1.337 | 0.021 | 208 | 33765 | 963 | 1.114 | 0.004 | 0 | 1.201 | 0.017 |
| ["['401']", '519'] | 244 | 401 | 519 | 10912 | 922 | 1.424 | 0.026 | 190 | 33765 | 713 | 1.374 | 0.012 | 0 | 1.036 | 0.014 |
| ["['038']", '079'] | 124 | 38 | 79 | 2780 | 1002 | 2.613 | 0.047 | 104 | 9632 | 963 | 1.952 | 0.017 | 0 | 1.339 | 0.03 |
| ["['038']", '519'] | 122 | 38 | 519 | 2780 | 922 | 2.794 | 0.051 | 95 | 9632 | 713 | 2.408 | 0.022 | 0 | 1.161 | 0.029 |
| ["['276']", '079'] | 113 | 276 | 79 | 2682 | 1002 | 2.469 | 0.042 | 106 | 9695 | 963 | 1.976 | 0.018 | 0 | 1.249 | 0.025 |
| ["['276']", '519'] | 111 | 276 | 519 | 2682 | 922 | 2.635 | 0.045 | 94 | 9695 | 713 | 2.367 | 0.021 | 0 | 1.113 | 0.024 |
| ["['294']", '079'] | 88 | 294 | 79 | 1678 | 1002 | 3.073 | 0.047 | 96 | 5740 | 963 | 3.023 | 0.028 | 0 | 1.016 | 0.019 |
| ["['584']", '079'] | 29 | 584 | 79 | 701 | 1002 | 2.424 | 0.021 | 19 | 2444 | 963 | 1.405 | 0.004 | 0 | 1.725 | 0.017 |
| ["['584']", '519'] | 30 | 584 | 519 | 701 | 922 | 2.725 | 0.024 | 16 | 2444 | 713 | 1.598 | 0.005 | 0 | 1.705 | 0.019 |
| ["['599']", '079'] | 157 | 599 | 79 | 3840 | 1002 | 2.396 | 0.049 | 170 | 13656 | 963 | 2.25 | 0.027 | 0 | 1.065 | 0.021 |
| ["['600']", '079'] | 40 | 600 | 79 | 1195 | 1002 | 1.961 | 0.018 | 18 | 4099 | 963 | 0.794 | -0.002 | 0 | 2.471 | 0.021 |
| ["['600']", '519'] | 40 | 600 | 519 | 1195 | 922 | 2.131 | 0.021 | 20 | 4099 | 713 | 1.191 | 0.002 | 0 | 1.789 | 0.019 |
| ["['788']", '079'] | 76 | 788 | 79 | 2550 | 1002 | 1.746 | 0.021 | 59 | 8296 | 963 | 1.286 | 0.005 | 0 | 1.358 | 0.016 |
| ["['788']", '519'] | 78 | 788 | 519 | 2550 | 922 | 1.948 | 0.026 | 64 | 8296 | 713 | 1.883 | 0.013 | 0 | 1.034 | 0.013 |
| ["['569']", '079'] | 62 | 569 | 79 | 2820 | 1002 | 1.288 | 0.009 | 41 | 8887 | 963 | 0.834 | -0.003 | 0 | 1.545 | 0.011 |
| ["['569']", '519'] | 62 | 569 | 519 | 2820 | 922 | 1.4 | 0.011 | 37 | 8887 | 713 | 1.016 | 0 | 0 | 1.377 | 0.011 |
| ["['079']", '535'] | 61 | 79 | 535 | 1393 | 2221 | 1.158 | 0.005 | 42 | 1310 | 7332 | 0.761 | -0.004 | 0 | 1.521 | 0.009 |
| ["['519']", '535'] | 50 | 519 | 535 | 1157 | 2221 | 1.142 | 0.004 | 18 | 601 | 7332 | 0.711 | -0.004 | 0 | 1.607 | 0.008 |
| ["['079']", '486'] | 165 | 79 | 486 | 1393 | 2330 | 2.985 | 0.063 | 46 | 1310 | 10312 | 0.593 | -0.009 | 0 | 5.035 | 0.072 |
| ["['079']", '682'] | 23 | 79 | 682 | 1393 | 650 | 1.491 | 0.008 | 15 | 1310 | 2162 | 0.922 | -0.001 | 0 | 1.618 | 0.009 |
| ["['079']", '786'] | 82 | 79 | 786 | 1393 | 1641 | 2.106 | 0.029 | 59 | 1310 | 6107 | 1.284 | 0.005 | 0 | 1.641 | 0.025 |
| ["['519']", '486'] | 170 | 519 | 486 | 1157 | 2330 | 3.702 | 0.078 | 58 | 601 | 10312 | 1.629 | 0.009 | 0 | 2.273 | 0.069 |
| ["['519']", '682'] | 25 | 519 | 682 | 1157 | 650 | 1.952 | 0.014 | 7 | 601 | 2162 | 0.938 | 0 | 0 | 2.081 | 0.015 |
| ["['519']", '786'] | 75 | 519 | 786 | 1157 | 1641 | 2.319 | 0.032 | 36 | 601 | 6107 | 1.707 | 0.008 | 0 | 1.358 | 0.024 |
| ["['272']", '278'] | 225 | 272 | 278 | 6624 | 795 | 2.508 | 0.063 | 516 | 19683 | 2094 | 2.179 | 0.046 | 0.001 | 1.151 | 0.017 |
| ["['535']", '924'] | 21 | 535 | 924 | 3646 | 140 | 2.415 | 0.018 | 35 | 11221 | 414 | 1.311 | 0.004 | 0.05 | 1.842 | 0.014 |
| ["['079']", '410'] | 25 | 79 | 410 | 1393 | 373 | 2.825 | 0.023 | 6 | 1310 | 2089 | 0.382 | -0.006 | 0 | 7.402 | 0.029 |
| ["['519']", '410'] | 25 | 519 | 410 | 1157 | 373 | 3.401 | 0.027 | 5 | 601 | 2089 | 0.693 | -0.002 | 0 | 4.906 | 0.029 |
| ["['079']", '038'] | 83 | 79 | 38 | 1393 | 1418 | 2.467 | 0.036 | 49 | 1310 | 6500 | 1.002 | 0 | 0 | 2.463 | 0.036 |
| ["['519']", '038'] | 78 | 519 | 38 | 1157 | 1418 | 2.791 | 0.04 | 23 | 601 | 6500 | 1.025 | 0 | 0 | 2.724 | 0.04 |
| ["['079']", '290'] | 16 | 79 | 290 | 1393 | 246 | 2.741 | 0.018 | 1 | 1310 | 992 | 0.134 | -0.006 | 0 | 20.465 | 0.023 |
| ["['519']", '290'] | 15 | 519 | 290 | 1157 | 246 | 3.094 | 0.019 | 4 | 601 | 992 | 1.168 | 0.001 | 0 | 2.649 | 0.019 |
| ["['285']", '079'] | 77 | 285 | 79 | 1889 | 1002 | 2.388 | 0.033 | 52 | 6413 | 963 | 1.466 | 0.007 | 0 | 1.63 | 0.027 |
| ["['285']", '519'] | 79 | 285 | 519 | 1889 | 922 | 2.663 | 0.038 | 47 | 6413 | 713 | 1.789 | 0.01 | 0 | 1.488 | 0.028 |
| ["['414']", '079'] | 57 | 414 | 79 | 2309 | 1002 | 1.446 | 0.012 | 36 | 8479 | 963 | 0.767 | -0.004 | 0 | 1.885 | 0.016 |
| ["['414']", '519'] | 57 | 414 | 519 | 2309 | 922 | 1.572 | 0.015 | 34 | 8479 | 713 | 0.979 | 0 | 0 | 1.606 | 0.015 |
| ["['427']", '079'] | 84 | 427 | 79 | 2685 | 1002 | 1.833 | 0.024 | 65 | 10071 | 963 | 1.167 | 0.003 | 0 | 1.571 | 0.021 |
| ["['427']", '519'] | 83 | 427 | 519 | 2685 | 922 | 1.968 | 0.027 | 58 | 10071 | 713 | 1.406 | 0.006 | 0 | 1.4 | 0.02 |
| ["['852']", '079'] | 11 | 852 | 79 | 166 | 1002 | 3.883 | 0.02 | 6 | 620 | 963 | 1.749 | 0.003 | 0.001 | 2.22 | 0.017 |
| ["['852']", '519'] | 11 | 852 | 519 | 166 | 922 | 4.22 | 0.022 | 7 | 620 | 713 | 2.756 | 0.007 | 0.001 | 1.531 | 0.015 |
| ["['250']", '079'] | 171 | 250 | 79 | 6323 | 1002 | 1.585 | 0.027 | 118 | 20788 | 963 | 1.026 | 0.001 | 0 | 1.544 | 0.026 |
| ["['250']", '519'] | 167 | 250 | 519 | 6323 | 922 | 1.682 | 0.03 | 114 | 20788 | 713 | 1.339 | 0.008 | 0 | 1.256 | 0.022 |
| ["['536']", '079'] | 15 | 536 | 79 | 376 | 1002 | 2.337 | 0.014 | 4 | 1286 | 963 | 0.562 | -0.003 | 0 | 4.157 | 0.017 |
| ["['536']", '519'] | 11 | 536 | 519 | 376 | 922 | 1.863 | 0.009 | 5 | 1286 | 713 | 0.949 | 0 | 0 | 1.963 | 0.009 |
| ["['079']", '041'] | 26 | 79 | 41 | 1393 | 564 | 1.943 | 0.014 | 14 | 1310 | 1832 | 1.015 | 0 | 0 | 1.913 | 0.014 |
| ["['535']", '079'] | 77 | 535 | 79 | 3646 | 1002 | 1.237 | 0.008 | 63 | 11221 | 963 | 1.015 | 0 | 0 | 1.219 | 0.008 |
| ["['535']", '519'] | 73 | 535 | 519 | 3646 | 922 | 1.275 | 0.009 | 48 | 11221 | 713 | 1.044 | 0.001 | 0 | 1.221 | 0.008 |
| ["['626']", '535'] | 24 | 626 | 535 | 391 | 2221 | 1.623 | 0.01 | 40 | 927 | 7332 | 1.024 | 0 | 0.034 | 1.584 | 0.01 |
| ["['079']", '438'] | 23 | 79 | 438 | 1393 | 377 | 2.571 | 0.02 | 13 | 1310 | 1919 | 0.9 | -0.001 | 0 | 2.856 | 0.021 |
| ["['079']", '780'] | 111 | 79 | 780 | 1393 | 2786 | 1.679 | 0.024 | 77 | 1310 | 10935 | 0.936 | -0.001 | 0 | 1.795 | 0.025 |
| ["['519']", '438'] | 23 | 519 | 438 | 1157 | 377 | 3.096 | 0.024 | 5 | 601 | 1919 | 0.755 | -0.002 | 0 | 4.102 | 0.025 |
| ["['519']", '780'] | 104 | 519 | 780 | 1157 | 2786 | 1.894 | 0.028 | 59 | 601 | 10935 | 1.563 | 0.009 | 0 | 1.212 | 0.02 |
| ["['782']", '079'] | 27 | 782 | 79 | 972 | 1002 | 1.628 | 0.011 | 25 | 3120 | 963 | 1.448 | 0.005 | 0 | 1.124 | 0.006 |
| ["['782']", '519'] | 26 | 782 | 519 | 972 | 922 | 1.703 | 0.012 | 21 | 3120 | 713 | 1.643 | 0.006 | 0 | 1.037 | 0.006 |
| ["['486']", '924'] | 15 | 486 | 924 | 2814 | 140 | 2.235 | 0.014 | 20 | 10276 | 414 | 0.818 | -0.002 | 0.027 | 2.732 | 0.016 |
| ["['079']", '707'] | 34 | 79 | 707 | 1393 | 397 | 3.61 | 0.034 | 9 | 1310 | 1306 | 0.916 | -0.001 | 0 | 3.942 | 0.034 |
| ["['519']", '707'] | 34 | 519 | 707 | 1157 | 397 | 4.346 | 0.039 | 5 | 601 | 1306 | 1.109 | 0.001 | 0 | 3.919 | 0.039 |
| ["['055']", '079'] | 12 | 55 | 79 | 331 | 1002 | 2.124 | 0.011 | 5 | 1083 | 963 | 0.835 | -0.001 | 0 | 2.545 | 0.012 |
| ["['583']", '079'] | 16 | 583 | 79 | 517 | 1002 | 1.813 | 0.01 | 13 | 1820 | 963 | 1.291 | 0.002 | 0 | 1.404 | 0.008 |
| ["['583']", '519'] | 15 | 583 | 519 | 517 | 922 | 1.847 | 0.01 | 12 | 1820 | 713 | 1.61 | 0.004 | 0.001 | 1.148 | 0.006 |
| ["['585']", '079'] | 101 | 585 | 79 | 2182 | 1002 | 2.712 | 0.044 | 34 | 7276 | 963 | 0.845 | -0.002 | 0 | 3.211 | 0.047 |
| ["['585']", '519'] | 101 | 585 | 519 | 2182 | 922 | 2.947 | 0.048 | 31 | 7276 | 713 | 1.04 | 0.001 | 0 | 2.834 | 0.048 |
| ["['410']", '079'] | 16 | 410 | 79 | 692 | 1002 | 1.355 | 0.005 | 11 | 2584 | 963 | 0.769 | -0.002 | 0 | 1.761 | 0.007 |
| ["['410']", '519'] | 16 | 410 | 519 | 692 | 922 | 1.472 | 0.007 | 10 | 2584 | 713 | 0.945 | 0 | 0 | 1.558 | 0.007 |
| ["['428']", '079'] | 72 | 428 | 79 | 1866 | 1002 | 2.261 | 0.03 | 37 | 7020 | 963 | 0.953 | -0.001 | 0 | 2.373 | 0.031 |
| ["['428']", '519'] | 74 | 428 | 519 | 1866 | 922 | 2.525 | 0.035 | 40 | 7020 | 713 | 1.391 | 0.005 | 0 | 1.815 | 0.03 |
| ["['486']", '079'] | 157 | 486 | 79 | 2814 | 1002 | 3.269 | 0.067 | 129 | 10276 | 963 | 2.269 | 0.024 | 0 | 1.441 | 0.043 |
| ["['486']", '519'] | 161 | 486 | 519 | 2814 | 922 | 3.643 | 0.075 | 138 | 10276 | 713 | 3.279 | 0.037 | 0 | 1.111 | 0.038 |
| ["['079']", '093'] | 29 | 79 | 93 | 1393 | 427 | 2.862 | 0.025 | 26 | 1310 | 2181 | 1.584 | 0.006 | 0 | 1.807 | 0.019 |
| ["['079']", '507'] | 12 | 79 | 507 | 1393 | 162 | 3.122 | 0.017 | 10 | 1310 | 771 | 1.723 | 0.004 | 0.003 | 1.811 | 0.013 |
| ["['079']", '518'] | 39 | 79 | 518 | 1393 | 630 | 2.609 | 0.026 | 23 | 1310 | 3213 | 0.951 | -0.001 | 0 | 2.743 | 0.027 |
| ["['519']", '093'] | 30 | 519 | 93 | 1157 | 427 | 3.565 | 0.031 | 18 | 601 | 2181 | 2.39 | 0.009 | 0 | 1.491 | 0.022 |
| ["['519']", '518'] | 39 | 519 | 518 | 1157 | 630 | 3.141 | 0.032 | 28 | 601 | 3213 | 2.524 | 0.012 | 0 | 1.245 | 0.019 |
| ["['282']", '079'] | 12 | 282 | 79 | 276 | 1002 | 2.548 | 0.014 | 9 | 945 | 963 | 1.722 | 0.004 | 0.002 | 1.48 | 0.01 |
| ["['873']", '783'] | 11 | 873 | 783 | 131 | 555 | 8.883 | 0.036 | 10 | 461 | 2110 | 1.79 | 0.005 | 0.009 | 4.964 | 0.032 |
| ["['079']", '783'] | 31 | 79 | 783 | 1393 | 555 | 2.354 | 0.021 | 33 | 1310 | 2110 | 2.078 | 0.01 | 0 | 1.133 | 0.01 |
| ["['519']", '783'] | 28 | 519 | 783 | 1157 | 555 | 2.56 | 0.022 | 12 | 601 | 2110 | 1.647 | 0.004 | 0 | 1.554 | 0.017 |
| ["['533']", '535'] | 26 | 533 | 535 | 91 | 2221 | 7.553 | 0.051 | 25 | 320 | 7332 | 1.855 | 0.008 | 0 | 4.072 | 0.044 |
| ["['437']", '783'] | 33 | 437 | 783 | 1167 | 555 | 2.991 | 0.028 | 60 | 4351 | 2110 | 1.138 | 0.002 | 0.031 | 2.629 | 0.025 |
| ["['584']", '251'] | 29 | 584 | 251 | 701 | 310 | 7.835 | 0.055 | 41 | 2444 | 1168 | 2.5 | 0.015 | 0.003 | 3.134 | 0.04 |
| ["['401']", '883'] | 13 | 401 | 883 | 10912 | 52 | 1.345 | 0.005 | 12 | 33765 | 100 | 0.619 | -0.004 | 0.004 | 2.174 | 0.009 |
| ["['628']", '041'] | 14 | 628 | 41 | 123 | 564 | 11.848 | 0.049 | 13 | 213 | 1832 | 5.799 | 0.017 | 0.003 | 2.043 | 0.032 |
| ["['366']", '079'] | 51 | 366 | 79 | 1818 | 1002 | 1.644 | 0.015 | 27 | 5300 | 963 | 0.921 | -0.001 | 0 | 1.785 | 0.016 |
| ["['786']", '079'] | 77 | 786 | 79 | 3230 | 1002 | 1.397 | 0.013 | 70 | 10549 | 963 | 1.199 | 0.004 | 0 | 1.165 | 0.009 |
| ["['482']", '079'] | 12 | 482 | 79 | 266 | 1002 | 2.643 | 0.015 | 14 | 962 | 963 | 2.631 | 0.009 | 0.026 | 1.005 | 0.006 |
| ["['790']", '278'] | 91 | 790 | 278 | 2705 | 795 | 2.484 | 0.038 | 195 | 7575 | 2094 | 2.14 | 0.027 | 0.012 | 1.161 | 0.011 |
| ["['437']", '079'] | 41 | 437 | 79 | 1167 | 1002 | 2.059 | 0.02 | 28 | 4351 | 963 | 1.163 | 0.002 | 0 | 1.77 | 0.018 |
| ["['437']", '519'] | 39 | 437 | 519 | 1167 | 922 | 2.128 | 0.02 | 27 | 4351 | 713 | 1.515 | 0.005 | 0 | 1.405 | 0.015 |
| ["['455']", '998'] | 11 | 455 | 998 | 1021 | 214 | 2.956 | 0.016 | 13 | 2949 | 774 | 0.991 | 0 | 0.037 | 2.981 | 0.016 |
| ["['079']", '276'] | 85 | 79 | 276 | 1393 | 1520 | 2.357 | 0.034 | 31 | 1310 | 6562 | 0.628 | -0.006 | 0 | 3.755 | 0.041 |
| ["['519']", '276'] | 84 | 519 | 276 | 1157 | 1520 | 2.804 | 0.042 | 19 | 601 | 6562 | 0.839 | -0.002 | 0 | 3.344 | 0.044 |
| ["['366']", '428'] | 63 | 366 | 428 | 1818 | 792 | 2.569 | 0.033 | 129 | 5300 | 3108 | 1.363 | 0.009 | 0.019 | 1.884 | 0.024 |
| ["['274']", '079'] | 54 | 274 | 79 | 1352 | 1002 | 2.34 | 0.027 | 21 | 4621 | 963 | 0.821 | -0.002 | 0 | 2.849 | 0.029 |
| ["['274']", '519'] | 54 | 274 | 519 | 1352 | 922 | 2.543 | 0.03 | 21 | 4621 | 713 | 1.109 | 0.001 | 0 | 2.292 | 0.029 |
| ["['511']", '079'] | 18 | 511 | 79 | 427 | 1002 | 2.47 | 0.017 | 7 | 1657 | 963 | 0.764 | -0.002 | 0 | 3.235 | 0.018 |
| ["['511']", '519'] | 18 | 511 | 519 | 427 | 922 | 2.684 | 0.018 | 7 | 1657 | 713 | 1.031 | 0 | 0 | 2.603 | 0.018 |
| ["['719']", '079'] | 30 | 719 | 79 | 902 | 1002 | 1.949 | 0.016 | 22 | 3060 | 963 | 1.3 | 0.003 | 0 | 1.5 | 0.013 |
| ["['719']", '519'] | 28 | 719 | 519 | 902 | 922 | 1.977 | 0.015 | 21 | 3060 | 713 | 1.675 | 0.006 | 0 | 1.18 | 0.01 |
| ["['079']", '401'] | 91 | 79 | 401 | 1393 | 3438 | 1.116 | 0.004 | 79 | 1310 | 13282 | 0.79 | -0.005 | 0 | 1.411 | 0.01 |
| ["['519']", '401'] | 78 | 519 | 401 | 1157 | 3438 | 1.151 | 0.005 | 39 | 601 | 13282 | 0.85 | -0.003 | 0 | 1.354 | 0.008 |
| ["['079']", '787'] | 28 | 79 | 787 | 1393 | 584 | 2.021 | 0.016 | 20 | 1310 | 2339 | 1.136 | 0.001 | 0 | 1.778 | 0.015 |
| ["['519']", '787'] | 28 | 519 | 787 | 1157 | 584 | 2.433 | 0.02 | 11 | 601 | 2339 | 1.362 | 0.002 | 0 | 1.786 | 0.018 |
| ["['719']", '785'] | 25 | 719 | 785 | 902 | 699 | 2.328 | 0.018 | 43 | 3060 | 2990 | 0.818 | -0.003 | 0.04 | 2.846 | 0.021 |
| ["['724']", '079'] | 34 | 724 | 79 | 1302 | 1002 | 1.53 | 0.011 | 35 | 4507 | 963 | 1.404 | 0.005 | 0 | 1.09 | 0.006 |
| ["['724']", '519'] | 36 | 724 | 519 | 1302 | 922 | 1.761 | 0.014 | 31 | 4507 | 713 | 1.679 | 0.007 | 0 | 1.048 | 0.007 |
| ["['079']", '599'] | 124 | 79 | 599 | 1393 | 1837 | 2.845 | 0.052 | 45 | 1310 | 6800 | 0.879 | -0.002 | 0 | 3.235 | 0.054 |
| ["['519']", '599'] | 114 | 519 | 599 | 1157 | 1837 | 3.149 | 0.055 | 27 | 601 | 6800 | 1.15 | 0.002 | 0 | 2.738 | 0.053 |
| ["['079']", '482'] | 32 | 79 | 482 | 1393 | 250 | 5.395 | 0.045 | 16 | 1310 | 1306 | 1.628 | 0.005 | 0 | 3.314 | 0.04 |
| ["['519']", '482'] | 34 | 519 | 482 | 1157 | 250 | 6.901 | 0.055 | 17 | 601 | 1306 | 3.77 | 0.014 | 0 | 1.83 | 0.041 |
| ["['272']", '822'] | 13 | 272 | 822 | 6624 | 45 | 2.56 | 0.015 | 11 | 19683 | 131 | 0.743 | -0.003 | 0.002 | 3.448 | 0.018 |
| ["['574']", '079'] | 19 | 574 | 79 | 647 | 1002 | 1.721 | 0.01 | 13 | 2318 | 963 | 1.014 | 0 | 0 | 1.697 | 0.01 |
| ["['574']", '519'] | 18 | 574 | 519 | 647 | 922 | 1.772 | 0.01 | 15 | 2318 | 713 | 1.58 | 0.004 | 0 | 1.121 | 0.006 |
| ["['576']", '079'] | 13 | 576 | 79 | 489 | 1002 | 1.558 | 0.007 | 12 | 1784 | 963 | 1.216 | 0.002 | 0.004 | 1.281 | 0.005 |
| ["['079']", '585'] | 22 | 79 | 585 | 1393 | 670 | 1.384 | 0.006 | 17 | 1310 | 2845 | 0.794 | -0.002 | 0 | 1.743 | 0.009 |
| ["['519']", '585'] | 22 | 519 | 585 | 1157 | 670 | 1.666 | 0.01 | 5 | 601 | 2845 | 0.509 | -0.004 | 0 | 3.273 | 0.014 |
| ["['447']", '996'] | 11 | 447 | 996 | 218 | 420 | 7.053 | 0.031 | 9 | 798 | 1507 | 1.303 | 0.002 | 0.005 | 5.414 | 0.029 |
| ["['079']", '788'] | 47 | 79 | 788 | 1393 | 1258 | 1.575 | 0.013 | 15 | 1310 | 4438 | 0.449 | -0.008 | 0 | 3.506 | 0.021 |
| ["['519']", '788'] | 47 | 519 | 788 | 1157 | 1258 | 1.896 | 0.019 | 13 | 601 | 4438 | 0.848 | -0.001 | 0 | 2.235 | 0.02 |
| ["['496']", '079'] | 44 | 496 | 79 | 828 | 1002 | 3.114 | 0.033 | 18 | 3116 | 963 | 1.044 | 0 | 0 | 2.982 | 0.033 |
| ["['496']", '519'] | 44 | 496 | 519 | 828 | 922 | 3.384 | 0.036 | 24 | 3116 | 713 | 1.88 | 0.008 | 0 | 1.8 | 0.028 |
| ["['530']", '079'] | 21 | 530 | 79 | 976 | 1002 | 1.261 | 0.004 | 18 | 3312 | 963 | 0.982 | 0 | 0 | 1.283 | 0.005 |
| ["['530']", '519'] | 22 | 530 | 519 | 976 | 922 | 1.435 | 0.007 | 18 | 3312 | 713 | 1.327 | 0.003 | 0 | 1.082 | 0.004 |
| ["['280']", '079'] | 28 | 280 | 79 | 922 | 1002 | 1.779 | 0.013 | 22 | 3120 | 963 | 1.275 | 0.003 | 0 | 1.396 | 0.01 |
| ["['531']", '079'] | 18 | 531 | 79 | 519 | 1002 | 2.032 | 0.013 | 6 | 1895 | 963 | 0.572 | -0.003 | 0 | 3.551 | 0.016 |
| ["['531']", '519'] | 18 | 531 | 519 | 519 | 922 | 2.208 | 0.014 | 7 | 1895 | 713 | 0.902 | -0.001 | 0 | 2.449 | 0.015 |
| ["['079']", '553'] | 11 | 79 | 553 | 1393 | 248 | 1.869 | 0.009 | 5 | 1310 | 1010 | 0.658 | -0.002 | 0 | 2.842 | 0.011 |
| ["['241']", '789'] | 15 | 241 | 789 | 246 | 1347 | 2.658 | 0.016 | 16 | 780 | 4629 | 0.771 | -0.003 | 0.006 | 3.445 | 0.019 |
| ["['079']", '854'] | 15 | 79 | 854 | 1393 | 479 | 1.32 | 0.005 | 11 | 1310 | 1746 | 0.837 | -0.001 | 0 | 1.577 | 0.006 |
| ["['519']", '854'] | 15 | 519 | 854 | 1157 | 479 | 1.589 | 0.008 | 4 | 601 | 1746 | 0.664 | -0.002 | 0 | 2.395 | 0.01 |
| ["['790']", '386'] | 13 | 790 | 386 | 2705 | 89 | 3.17 | 0.019 | 17 | 7575 | 274 | 1.426 | 0.004 | 0.038 | 2.224 | 0.015 |
| ["['244']", '079'] | 15 | 244 | 79 | 584 | 1002 | 1.505 | 0.007 | 9 | 2051 | 963 | 0.793 | -0.002 | 0 | 1.897 | 0.008 |
| ["['244']", '519'] | 14 | 244 | 519 | 584 | 922 | 1.526 | 0.007 | 10 | 2051 | 713 | 1.19 | 0.001 | 0 | 1.282 | 0.005 |
| ["['733']", '079'] | 21 | 733 | 79 | 704 | 1002 | 1.748 | 0.011 | 15 | 2470 | 963 | 1.098 | 0.001 | 0 | 1.592 | 0.01 |
| ["['733']", '519'] | 21 | 733 | 519 | 704 | 922 | 1.899 | 0.013 | 14 | 2470 | 713 | 1.384 | 0.003 | 0 | 1.373 | 0.01 |
| ["['785']", '079'] | 24 | 785 | 79 | 1106 | 1002 | 1.271 | 0.005 | 24 | 3791 | 963 | 1.144 | 0.002 | 0 | 1.111 | 0.003 |
| ["['790']", '079'] | 60 | 790 | 79 | 2705 | 1002 | 1.3 | 0.009 | 31 | 7575 | 963 | 0.74 | -0.004 | 0 | 1.757 | 0.013 |
| ["['790']", '519'] | 59 | 790 | 519 | 2705 | 922 | 1.389 | 0.011 | 23 | 7575 | 713 | 0.741 | -0.004 | 0 | 1.874 | 0.014 |
| ["['039']", '079'] | 52 | 39 | 79 | 824 | 1002 | 3.698 | 0.042 | 8 | 2430 | 963 | 0.595 | -0.004 | 0 | 6.214 | 0.046 |
| ["['039']", '519'] | 53 | 39 | 519 | 824 | 922 | 4.096 | 0.047 | 9 | 2430 | 713 | 0.904 | -0.001 | 0 | 4.53 | 0.047 |
| ["['288']", '519'] | 11 | 288 | 519 | 213 | 922 | 3.288 | 0.017 | 5 | 807 | 713 | 1.513 | 0.002 | 0 | 2.174 | 0.015 |
| ["['366']", '519'] | 48 | 366 | 519 | 1818 | 922 | 1.681 | 0.015 | 26 | 5300 | 713 | 1.198 | 0.002 | 0 | 1.404 | 0.013 |
| ["['996']", '079'] | 35 | 996 | 79 | 786 | 1002 | 2.609 | 0.025 | 20 | 2426 | 963 | 1.49 | 0.004 | 0 | 1.751 | 0.02 |
| ["['996']", '519'] | 36 | 996 | 519 | 786 | 922 | 2.916 | 0.028 | 17 | 2426 | 713 | 1.711 | 0.005 | 0 | 1.705 | 0.023 |
| ["['813']", '788'] | 15 | 813 | 788 | 288 | 1258 | 2.431 | 0.015 | 15 | 971 | 4438 | 0.606 | -0.005 | 0.004 | 4.012 | 0.02 |
| ["['401']", '480'] | 11 | 401 | 480 | 10912 | 27 | 2.192 | 0.012 | 9 | 33765 | 46 | 1.009 | 0 | 0.005 | 2.173 | 0.012 |
| ["['041']", '519'] | 29 | 41 | 519 | 771 | 922 | 2.395 | 0.02 | 22 | 2796 | 713 | 1.921 | 0.008 | 0 | 1.247 | 0.013 |
| ["['518']", '079'] | 39 | 518 | 79 | 938 | 1002 | 2.436 | 0.024 | 20 | 3351 | 963 | 1.079 | 0.001 | 0 | 2.258 | 0.023 |
| ["['518']", '519'] | 38 | 518 | 519 | 938 | 922 | 2.58 | 0.025 | 33 | 3351 | 713 | 2.404 | 0.013 | 0 | 1.073 | 0.013 |
| ["['332']", '079'] | 15 | 332 | 79 | 272 | 1002 | 3.231 | 0.02 | 10 | 1165 | 963 | 1.552 | 0.003 | 0 | 2.083 | 0.017 |
| ["['332']", '519'] | 15 | 332 | 519 | 272 | 922 | 3.512 | 0.022 | 10 | 1165 | 713 | 2.096 | 0.006 | 0 | 1.676 | 0.016 |
| ["['079']", '096'] | 19 | 79 | 96 | 1393 | 264 | 3.033 | 0.021 | 19 | 1310 | 1802 | 1.401 | 0.004 | 0.001 | 2.165 | 0.018 |
| ["['519']", '096'] | 17 | 519 | 96 | 1157 | 264 | 3.268 | 0.022 | 10 | 601 | 1802 | 1.607 | 0.004 | 0 | 2.033 | 0.018 |
| ["['079']", '280'] | 29 | 79 | 280 | 1393 | 450 | 2.716 | 0.024 | 10 | 1310 | 1707 | 0.778 | -0.002 | 0 | 3.489 | 0.025 |
| ["['519']", '280'] | 21 | 519 | 280 | 1157 | 450 | 2.368 | 0.017 | 7 | 601 | 1707 | 1.188 | 0.001 | 0 | 1.994 | 0.016 |
| ["['079']", '458'] | 20 | 79 | 458 | 1393 | 300 | 2.81 | 0.02 | 5 | 1310 | 1146 | 0.58 | -0.003 | 0 | 4.847 | 0.023 |
| ["['079']", '569'] | 39 | 79 | 569 | 1393 | 1620 | 1.015 | 0 | 27 | 1310 | 5776 | 0.621 | -0.006 | 0 | 1.633 | 0.006 |
| ["['519']", '458'] | 20 | 519 | 458 | 1157 | 300 | 3.383 | 0.024 | 8 | 601 | 1146 | 2.022 | 0.005 | 0 | 1.673 | 0.019 |
| ["['519']", '569'] | 36 | 519 | 569 | 1157 | 1620 | 1.128 | 0.003 | 16 | 601 | 5776 | 0.802 | -0.002 | 0 | 1.405 | 0.005 |
| ["['079']", '789'] | 43 | 79 | 789 | 1393 | 1347 | 1.345 | 0.008 | 42 | 1310 | 4629 | 1.206 | 0.003 | 0 | 1.116 | 0.005 |
| ["['519']", '250'] | 46 | 519 | 250 | 1157 | 2109 | 1.107 | 0.003 | 19 | 601 | 7564 | 0.728 | -0.003 | 0 | 1.521 | 0.006 |
| ["['519']", '789'] | 33 | 519 | 789 | 1157 | 1347 | 1.243 | 0.005 | 15 | 601 | 4629 | 0.939 | -0.001 | 0 | 1.325 | 0.006 |
| ["['789']", '309'] | 35 | 789 | 309 | 2665 | 261 | 2.954 | 0.028 | 66 | 8114 | 799 | 1.772 | 0.012 | 0.039 | 1.667 | 0.017 |
| ["['079']", '573'] | 18 | 79 | 573 | 1393 | 371 | 2.045 | 0.013 | 7 | 1310 | 1479 | 0.629 | -0.003 | 0 | 3.251 | 0.016 |
| ["['519']", '573'] | 16 | 519 | 573 | 1157 | 371 | 2.188 | 0.013 | 2 | 601 | 1479 | 0.392 | -0.003 | 0 | 5.587 | 0.017 |
| ["['535']", '626'] | 22 | 535 | 626 | 3646 | 205 | 1.728 | 0.011 | 34 | 11221 | 492 | 1.072 | 0.001 | 0.023 | 1.612 | 0.01 |
| ["['573']", '079'] | 15 | 573 | 79 | 692 | 1002 | 1.27 | 0.004 | 14 | 2386 | 963 | 1.061 | 0.001 | 0.002 | 1.197 | 0.003 |
| ["['573']", '519'] | 15 | 573 | 519 | 692 | 922 | 1.38 | 0.005 | 11 | 2386 | 713 | 1.126 | 0.001 | 0 | 1.226 | 0.004 |

| | | | | | | | | | | | | | | | |
|---|---|---|---|---|---|---|---|---|---|---|---|---|---|---|---|
| ["['854']", '079'] | 43 | 854 | 79 | 1078 | 1002 | 2.337 | 0.024 | 34 | 3509 | 963 | 1.751 | 0.008 | 0 | 1.334 | 0.016 |
| ["['854']", '519'] | 42 | 854 | 519 | 1078 | 922 | 2.481 | 0.026 | 31 | 3509 | 713 | 2.157 | 0.011 | 0 | 1.15 | 0.015 |
| ["['079']", '511'] | 16 | 79 | 511 | 1393 | 373 | 1.808 | 0.01 | 2 | 1310 | 1759 | 0.151 | -0.007 | 0 | 11.966 | 0.018 |
| ["['519']", '511'] | 19 | 519 | 511 | 1157 | 373 | 2.585 | 0.018 | 11 | 601 | 1759 | 1.811 | 0.005 | 0 | 1.427 | 0.013 |
| ["['378']", '780'] | 11 | 378 | 780 | 70 | 2786 | 3.312 | 0.018 | 13 | 248 | 10935 | 0.834 | -0.002 | 0.037 | 3.968 | 0.019 |
| ["['560']", '079'] | 15 | 560 | 79 | 345 | 1002 | 2.548 | 0.016 | 7 | 1299 | 963 | 0.974 | 0 | 0 | 2.615 | 0.016 |
| ["['560']", '519'] | 14 | 560 | 519 | 345 | 922 | 2.584 | 0.015 | 5 | 1299 | 713 | 0.94 | 0 | 0 | 2.75 | 0.016 |
| ["['197']", '079'] | 14 | 197 | 79 | 663 | 1002 | 1.237 | 0.003 | 5 | 3171 | 963 | 0.285 | -0.007 | 0 | 4.341 | 0.011 |
| ["['197']", '519'] | 14 | 197 | 519 | 663 | 922 | 1.345 | 0.005 | 9 | 3171 | 713 | 0.693 | -0.003 | 0 | 1.94 | 0.007 |
| ["['281']", '079'] | 15 | 281 | 79 | 171 | 1002 | 5.14 | 0.029 | 8 | 697 | 963 | 2.075 | 0.005 | 0 | 2.477 | 0.024 |
| ["['281']", '519'] | 15 | 281 | 519 | 171 | 922 | 5.586 | 0.031 | 6 | 697 | 713 | 2.102 | 0.004 | 0 | 2.658 | 0.027 |
| ["['593']", '079'] | 23 | 593 | 79 | 914 | 1002 | 1.474 | 0.008 | 19 | 3115 | 963 | 1.103 | 0.001 | 0 | 1.337 | 0.007 |
| ["['593']", '519'] | 25 | 593 | 519 | 914 | 922 | 1.742 | 0.012 | 19 | 3115 | 713 | 1.489 | 0.004 | 0 | 1.17 | 0.008 |
| ["['564']", '079'] | 30 | 564 | 79 | 646 | 1002 | 2.721 | 0.024 | 26 | 2291 | 963 | 2.051 | 0.009 | 0 | 1.326 | 0.015 |
| ["['564']", '519'] | 27 | 564 | 519 | 646 | 922 | 2.661 | 0.022 | 23 | 2291 | 713 | 2.451 | 0.011 | 0 | 1.086 | 0.011 |
| ["['079']", '285'] | 56 | 79 | 285 | 1393 | 1042 | 2.265 | 0.027 | 22 | 1310 | 4200 | 0.696 | -0.004 | 0 | 3.254 | 0.031 |
| ["['079']", '436'] | 12 | 79 | 436 | 1393 | 203 | 2.491 | 0.014 | 7 | 1310 | 931 | 0.999 | 0 | 0 | 2.494 | 0.014 |
| ["['519']", '285'] | 53 | 519 | 285 | 1157 | 1042 | 2.581 | 0.03 | 20 | 601 | 4200 | 1.379 | 0.004 | 0 | 1.871 | 0.027 |
| ["['519']", '436'] | 13 | 519 | 436 | 1157 | 203 | 3.25 | 0.019 | 2 | 601 | 931 | 0.622 | -0.002 | 0 | 5.223 | 0.02 |
| ["['202']", '079'] | 13 | 202 | 79 | 128 | 1002 | 5.951 | 0.03 | 3 | 501 | 963 | 1.082 | 0 | 0 | 5.498 | 0.03 |
| ["['202']", '519'] | 13 | 202 | 519 | 128 | 922 | 6.467 | 0.032 | 1 | 501 | 713 | 0.487 | -0.002 | 0 | 13.271 | 0.034 |
| ["['537']", '079'] | 13 | 537 | 79 | 421 | 1002 | 1.809 | 0.009 | 5 | 1442 | 963 | 0.627 | -0.003 | 0 | 2.887 | 0.012 |
| ["['537']", '519'] | 12 | 537 | 519 | 421 | 922 | 1.815 | 0.009 | 3 | 1442 | 713 | 0.508 | -0.003 | 0 | 3.573 | 0.012 |
| ["['692']", '079'] | 14 | 692 | 79 | 332 | 1002 | 2.471 | 0.015 | 14 | 1202 | 963 | 2.105 | 0.007 | 0.005 | 1.174 | 0.008 |
| ["['692']", '519'] | 14 | 692 | 519 | 332 | 922 | 2.685 | 0.016 | 6 | 1202 | 713 | 1.219 | 0.001 | 0 | 2.203 | 0.015 |
| ["['362']", '079'] | 19 | 362 | 79 | 921 | 1002 | 1.209 | 0.003 | 13 | 3060 | 963 | 0.768 | -0.002 | 0 | 1.574 | 0.006 |
| ["['362']", '519'] | 19 | 362 | 519 | 921 | 922 | 1.314 | 0.005 | 12 | 3060 | 713 | 0.957 | 0 | 0 | 1.372 | 0.005 |
| ["['403']", '079'] | 17 | 403 | 79 | 281 | 1002 | 3.545 | 0.023 | 13 | 936 | 963 | 2.511 | 0.008 | 0 | 1.412 | 0.015 |
| ["['403']", '519'] | 17 | 403 | 519 | 281 | 922 | 3.852 | 0.025 | 13 | 936 | 713 | 3.391 | 0.011 | 0 | 1.136 | 0.014 |
| ["['434']", '079'] | 30 | 434 | 79 | 739 | 1002 | 2.379 | 0.021 | 27 | 2788 | 963 | 1.751 | 0.007 | 0 | 1.359 | 0.013 |
| ["['436']", '079'] | 12 | 436 | 79 | 372 | 1002 | 1.89 | 0.009 | 10 | 1515 | 963 | 1.193 | 0.001 | 0.003 | 1.584 | 0.008 |
| ["['436']", '519'] | 13 | 436 | 519 | 372 | 922 | 2.225 | 0.012 | 13 | 1515 | 713 | 2.095 | 0.007 | 0.007 | 1.062 | 0.006 |
| ["['562']", '036'] | 13 | 562 | 36 | 789 | 265 | 3.65 | 0.021 | 17 | 2660 | 990 | 1.124 | 0.001 | 0.038 | 3.248 | 0.02 |
| ["['079']", '578'] | 18 | 79 | 578 | 1393 | 332 | 2.285 | 0.015 | 9 | 1310 | 1478 | 0.809 | -0.002 | 0 | 2.824 | 0.017 |
| ["['519']", '578'] | 17 | 519 | 578 | 1157 | 332 | 2.598 | 0.017 | 3 | 601 | 1478 | 0.588 | -0.002 | 0 | 4.42 | 0.019 |
| ["['079']", '414'] | 41 | 79 | 414 | 1393 | 1077 | 1.604 | 0.013 | 14 | 1310 | 4193 | 0.444 | -0.008 | 0 | 3.616 | 0.02 |
| ["['519']", '414'] | 40 | 519 | 414 | 1157 | 1077 | 1.885 | 0.017 | 11 | 601 | 4193 | 0.76 | -0.002 | 0 | 2.48 | 0.019 |
| ["['079']", '008'] | 19 | 79 | 8 | 1393 | 144 | 5.561 | 0.035 | 15 | 1310 | 593 | 3.361 | 0.012 | 0 | 1.654 | 0.023 |
| ["['519']", '008'] | 14 | 519 | 8 | 1157 | 144 | 4.933 | 0.028 | 7 | 601 | 593 | 3.419 | 0.008 | 0 | 1.443 | 0.019 |
| ["['079']", '437'] | 27 | 79 | 437 | 1393 | 447 | 2.546 | 0.021 | 6 | 1310 | 1956 | 0.408 | -0.005 | 0 | 6.246 | 0.027 |
| ["['519']", '437'] | 28 | 519 | 437 | 1157 | 447 | 3.179 | 0.027 | 2 | 601 | 1956 | 0.296 | -0.004 | 0 | 10.733 | 0.031 |
| ["['458']", '079'] | 24 | 458 | 79 | 451 | 1002 | 3.118 | 0.025 | 12 | 1552 | 963 | 1.398 | 0.003 | 0 | 2.231 | 0.022 |
| ["['458']", '519'] | 23 | 458 | 519 | 451 | 922 | 3.247 | 0.025 | 11 | 1552 | 713 | 1.73 | 0.004 | 0 | 1.877 | 0.021 |
| ["['079']", '294'] | 33 | 79 | 294 | 1393 | 678 | 2.051 | 0.018 | 17 | 1310 | 2776 | 0.814 | -0.002 | 0 | 2.521 | 0.02 |
| ["['519']", '294'] | 32 | 519 | 294 | 1157 | 678 | 2.395 | 0.021 | 5 | 601 | 2776 | 0.522 | -0.004 | 0 | 4.591 | 0.025 |
| ["['553']", '079'] | 13 | 553 | 79 | 410 | 1002 | 1.858 | 0.009 | 5 | 1459 | 963 | 0.619 | -0.003 | 0 | 2.999 | 0.012 |
| ["['553']", '519'] | 14 | 553 | 519 | 410 | 922 | 2.174 | 0.012 | 4 | 1459 | 713 | 0.669 | -0.002 | 0 | 3.249 | 0.014 |
| ["['079']", '037'] | 23 | 79 | 37 | 1393 | 560 | 1.731 | 0.011 | 10 | 1310 | 2127 | 0.625 | -0.004 | 0 | 2.771 | 0.015 |
| ["['519']", '037'] | 24 | 519 | 37 | 1157 | 560 | 2.175 | 0.016 | 9 | 601 | 2127 | 1.226 | 0.001 | 0 | 1.774 | 0.015 |
| ["['079']", '785'] | 29 | 79 | 785 | 1393 | 699 | 1.749 | 0.013 | 22 | 1310 | 2990 | 0.978 | 0 | 0 | 1.788 | 0.013 |
| ["['519']", '785'] | 26 | 519 | 785 | 1157 | 699 | 1.887 | 0.014 | 17 | 601 | 2990 | 1.647 | 0.005 | 0 | 1.146 | 0.009 |
| ["['079']", '793'] | 11 | 79 | 793 | 1393 | 245 | 1.892 | 0.009 | 2 | 1310 | 988 | 0.269 | -0.005 | 0 | 7.035 | 0.014 |
| ["['253']", '079'] | 11 | 253 | 79 | 224 | 1002 | 2.877 | 0.015 | 9 | 927 | 963 | 1.755 | 0.004 | 0.005 | 1.64 | 0.011 |
| ["['682']", '079'] | 45 | 682 | 79 | 1311 | 1002 | 2.011 | 0.02 | 38 | 4095 | 963 | 1.677 | 0.008 | 0 | 1.199 | 0.012 |
| ["['079']", '274'] | 16 | 79 | 274 | 1393 | 563 | 1.198 | 0.003 | 8 | 1310 | 2026 | 0.525 | -0.004 | 0 | 2.283 | 0.008 |
| ["['079']", '996'] | 27 | 79 | 996 | 1393 | 420 | 2.709 | 0.023 | 5 | 1310 | 1507 | 0.441 | -0.005 | 0 | 6.146 | 0.027 |
| ["['519']", '274'] | 16 | 519 | 274 | 1157 | 563 | 1.442 | 0.006 | 4 | 601 | 2026 | 0.572 | -0.003 | 0 | 2.522 | 0.009 |
| ["['519']", '996'] | 30 | 519 | 996 | 1157 | 420 | 3.625 | 0.032 | 6 | 601 | 1507 | 1.153 | 0.001 | 0 | 3.143 | 0.031 |
| ["['250']", '883'] | 11 | 250 | 883 | 6323 | 52 | 1.964 | 0.01 | 10 | 20788 | 100 | 0.837 | -0.001 | 0.009 | 2.346 | 0.011 |
| ["['519']", '197'] | 17 | 519 | 197 | 1157 | 442 | 1.952 | 0.012 | 14 | 601 | 2221 | 1.826 | 0.006 | 0 | 1.069 | 0.006 |
| ["['079']", '564'] | 15 | 79 | 564 | 1393 | 289 | 2.188 | 0.013 | 17 | 1310 | 1221 | 1.85 | 0.006 | 0.009 | 1.182 | 0.007 |
| ["['729']", '079'] | 13 | 729 | 79 | 445 | 1002 | 1.712 | 0.008 | 11 | 1485 | 963 | 1.339 | 0.002 | 0.002 | 1.278 | 0.006 |
| ["['729']", '519'] | 12 | 729 | 519 | 445 | 922 | 1.717 | 0.008 | 10 | 1485 | 713 | 1.644 | 0.004 | 0.003 | 1.044 | 0.004 |
| ["['571']", '519'] | 15 | 571 | 519 | 830 | 922 | 1.151 | 0.002 | 10 | 2623 | 713 | 0.931 | -0.001 | 0 | 1.236 | 0.003 |
| ["['998']", '783'] | 11 | 998 | 783 | 408 | 555 | 2.852 | 0.015 | 13 | 1300 | 2110 | 0.825 | -0.002 | 0.037 | 3.457 | 0.017 |
| ["['079']", '345'] | 11 | 79 | 345 | 1393 | 157 | 2.953 | 0.016 | 9 | 1310 | 571 | 2.094 | 0.005 | 0.005 | 1.41 | 0.01 |
| ["['924']", '079'] | 11 | 924 | 79 | 248 | 1002 | 2.599 | 0.014 | 12 | 857 | 963 | 2.531 | 0.008 | 0.024 | 1.027 | 0.006 |
| ["['784']", '519'] | 14 | 784 | 519 | 790 | 922 | 1.128 | 0.002 | 10 | 2574 | 713 | 0.948 | 0 | 0 | 1.19 | 0.002 |
| ["['788']", '011'] | 15 | 788 | 11 | 2550 | 104 | 3.321 | 0.021 | 20 | 8296 | 328 | 1.279 | 0.003 | 0.027 | 2.595 | 0.018 |
| ["['079']", '782'] | 18 | 79 | 782 | 1393 | 498 | 1.523 | 0.008 | 15 | 1310 | 1939 | 1.028 | 0 | 0 | 1.482 | 0.007 |
| ["['079']", '583'] | 12 | 79 | 583 | 1393 | 145 | 3.488 | 0.019 | 3 | 1310 | 672 | 0.593 | -0.002 | 0 | 5.88 | 0.022 |
| ["['581']", '079'] | 11 | 581 | 79 | 165 | 1002 | 3.906 | 0.02 | 2 | 546 | 963 | 0.662 | -0.001 | 0 | 5.9 | 0.022 |
| ["['781']", '079'] | 12 | 781 | 79 | 290 | 1002 | 2.425 | 0.013 | 5 | 902 | 963 | 1.002 | 0 | 0 | 2.42 | 0.013 |
| ["['781']", '519'] | 12 | 781 | 519 | 290 | 922 | 2.635 | 0.015 | 5 | 902 | 713 | 1.353 | 0.002 | 0 | 1.947 | 0.013 |
| ["['079']", '251'] | 17 | 79 | 251 | 1393 | 310 | 2.311 | 0.015 | 2 | 1310 | 1168 | 0.228 | -0.006 | 0 | 10.158 | 0.02 |
| ["['519']", '251'] | 16 | 519 | 251 | 1157 | 310 | 2.619 | 0.017 | 5 | 601 | 1168 | 1.24 | 0.001 | 0 | 2.112 | 0.016 |
| ["['793']", '079'] | 14 | 793 | 79 | 598 | 1002 | 1.372 | 0.005 | 11 | 2061 | 963 | 0.965 | 0 | 0.001 | 1.422 | 0.005 |
| ["['465']", '519'] | 13 | 465 | 519 | 347 | 922 | 2.386 | 0.013 | 12 | 1294 | 713 | 2.264 | 0.007 | 0.004 | 1.054 | 0.006 |
| ["['162']", '079'] | 14 | 162 | 79 | 617 | 1002 | 1.329 | 0.004 | 4 | 2794 | 963 | 0.259 | -0.007 | 0 | 5.138 | 0.012 |
| ["['162']", '519'] | 14 | 162 | 519 | 617 | 922 | 1.445 | 0.006 | 11 | 2794 | 713 | 0.961 | 0 | 0.001 | 1.503 | 0.006 |
| ["['519']", '564'] | 14 | 519 | 564 | 1157 | 289 | 2.458 | 0.015 | 2 | 601 | 1221 | 0.474 | -0.003 | 0 | 5.181 | 0.017 |
| ["['079']", '197'] | 14 | 79 | 197 | 1393 | 442 | 1.335 | 0.005 | 6 | 1310 | 2221 | 0.359 | -0.006 | 0 | 3.719 | 0.011 |
| ["['079']", '198'] | 11 | 79 | 198 | 1393 | 280 | 1.656 | 0.007 | 4 | 1310 | 1600 | 0.332 | -0.006 | 0 | 4.984 | 0.013 |
| ["['519']", '198'] | 14 | 519 | 198 | 1157 | 280 | 2.537 | 0.015 | 12 | 601 | 1600 | 2.172 | 0.007 | 0.002 | 1.168 | 0.008 |
| ["['008']", '519'] | 13 | 8 | 519 | 250 | 922 | 3.311 | 0.019 | 9 | 725 | 713 | 3.031 | 0.008 | 0.001 | 1.093 | 0.011 |
| ["['519']", '009'] | 11 | 519 | 9 | 1157 | 267 | 2.091 | 0.01 | 5 | 601 | 947 | 1.529 | 0.002 | 0 | 1.367 | 0.008 |
| ["['079']", '266'] | 11 | 79 | 266 | 1393 | 192 | 2.415 | 0.013 | 4 | 1310 | 716 | 0.742 | -0.001 | 0 | 3.253 | 0.014 |
| ["['079']", '496'] | 11 | 79 | 496 | 1393 | 262 | 1.77 | 0.008 | 3 | 1310 | 990 | 0.403 | -0.004 | 0 | 4.395 | 0.012 |
| ["['519']", '496'] | 11 | 519 | 496 | 1157 | 262 | 2.13 | 0.011 | 7 | 601 | 990 | 2.048 | 0.005 | 0.001 | 1.04 | 0.006 |
