## Supplementary Table 5 for "Deciphering the Molecular Mechanism of Post-Acute Sequelae of COVID-19 through Comorbidity Network Analysis"

| source | target | edge | source_protein | target_protein | source_protein_add | distance | distance with COVID | bg_mean | bg_std | Diff_repeat | Diff_outcome1 | z_score | p_value |
|---|---|---|---|---|---|---|---|---|---|---|---|---|---|
| ICD 038 | ICD 079 | 038_079 | 6, 3929, 4049, 4282, 4 | [1636] | 5, 5624, 5714, 5817, 5 | 2 | 2 | 0.262 | 0.489 | , 0.0, 1.0, 0.0, 0.0, 0.0 | 0 | -0.535 | 0.24 |
| ICD 038 | ICD 519 | 038_519 | 6, 3929, 4049, 4282, 4 | [2778, 5173, 7356] | 5, 5624, 5714, 5817, 5 | 1.667 | 1.333 | 0.426 | 0.295 | 6667, 0.33333333333 | 0.333 | -0.316 | 0.362 |
| ICD 041 | ICD 519 | 041_519 | [5243, 5803] | [2778, 5173, 7356] | 6923, 6938, 7088, 709 | 2.667 | 1.333 | 1.096 | 0.35 | 6666663, 0.66666666 | 1.333 | 0.679 | 0.106 |
| ICD 055 | ICD 079 | 055_079 | [4179] | [1636] | 6938, 7088, 7090, 745 | 3 | 2 | 1.202 | 0.611 | , 1.0, 1.0, 2.0, 1.0, 2.0 | 1 | -0.33 | 0.262 |
| ICD 079 | ICD 038 | 079_038 | [1636] | 6, 3929, 4049, 4282, 4 | 6938, 7088, 7090, 745 | 2.667 | 1.333 | 1.551 | 0.222 | 67, 1.4583333333333 | 1.333 | -0.981 | 0.816 |
| ICD 079 | ICD 041 | 079_041 | [1636] | [5243, 5803] | 6938, 7088, 7090, 745 | 3 | 1 | 1.517 | 0.498 | , 1.5, 1.0, 1.5, 1.5, 0.5 | 2 | 0.97 | 0.059 |
| ICD 079 | ICD 197 | 079_197 | [1636] | 6648, 6722, 6750, 70 | 6938, 7088, 7090, 745 | 2.672 | 1.328 | 1.567 | 0.2 | 9253731347, 1.716417 | 1.343 | -1.116 | 0.861 |
| ICD 079 | ICD 198 | 079_198 | [1636] | 6648, 6722, 6750, 70 | 6938, 7088, 7090, 745 | 2.672 | 1.328 | 1.568 | 0.193 | 28358207, 1.6417910 | 1.343 | -1.162 | 0.871 |
| ICD 079 | ICD 251 | 079_251 | [1636] | 3767, 5265, 5465, 66 | 6938, 7088, 7090, 745 | 2.824 | 1.353 | 1.534 | 0.228 | 235294117647056, 1.1 | 1.471 | -0.28 | 0.577 |
| ICD 079 | ICD 266 | 079_266 | [1636] | , 3141, 4524, 4548, 5 | 6938, 7088, 7090, 745 | 2.8 | 1.8 | 1.51 | 0.329 | 7, 1.6, 1.2000000000 | 1 | -1.553 | 0.946 |
| ICD 079 | ICD 274 | 079_274 | [1636] | 443, 7498, 9429, 229 | 6938, 7088, 7090, 745 | 2.909 | 1.727 | 1.489 | 0.259 | , 1.454545454545454 | 1.182 | -1.186 | 0.863 |
| ICD 079 | ICD 276 | 079_276 | [1636] | 2, 4780, 4879, 5019, 5 | 6938, 7088, 7090, 745 | 2.722 | 1.389 | 1.48 | 0.23 | 5556, 1.50000000000 | 1.333 | -0.636 | 0.709 |
| ICD 079 | ICD 280 | 079_280 | [1636] | , 6521, 6647, 7037, 7 | 6938, 7088, 7090, 745 | 2.75 | 1.417 | 1.542 | 0.241 | 33333, 1.4166666666 | 1.333 | -0.864 | 0.766 |
| ICD 079 | ICD 285 | 079_285 | [1636] | 5, 6006, 6007, 6223, | 6938, 7088, 7090, 745 | 2.729 | 1.458 | 1.501 | 0.191 | 4067796607, 1.64406 | 1.271 | -1.203 | 0.864 |
| ICD 079 | ICD 290 | 079_290 | [1636] | 4200, 4205, 4353, 45 | 6938, 7088, 7090, 745 | 2.759 | 1.387 | 1.531 | 0.19 | 904522613067, 1.723 | 1.372 | -0.841 | 0.843 |
| ICD 079 | ICD 294 | 079_294 | [1636] | , 5243, 5533, 5578, 5 | 6938, 7088, 7090, 745 | 2.741 | 1.385 | 1.531 | 0.189 | 706293706293708, 1.4 | 1.357 | -0.922 | 0.854 |
| ICD 079 | ICD 345 | 079_345 | [1636] | 4803, 4842, 4843, 48 | 6938, 7088, 7090, 745 | 2.81 | 1.473 | 1.52 | 0.186 | 36283184, 1.8982300 | 1.336 | -0.987 | 0.865 |
| ICD 079 | ICD 401 | 079_401 | [1636] | 40, 2778, 2784, 2796, | 6938, 7088, 7090, 745 | 2.737 | 1.413 | 1.518 | 0.182 | 014749, 1.49262536 | 1.324 | -1.067 | 0.86 |
| ICD 079 | ICD 410 | 079_410 | [1636] | 17, 3726, 3764, 3791, | 6938, 7088, 7090, 745 | 2.713 | 1.315 | 1.565 | 0.183 | 685082872928, 1.558 | 1.398 | -0.915 | 0.857 |
| ICD 079 | ICD 414 | 079_414 | [1636] | 3306, 3313, 3383, 339 | 6938, 7088, 7090, 745 | 2.715 | 1.361 | 1.552 | 0.175 | 36546184738954, 1.5 | 1.353 | -1.135 | 0.886 |
| ICD 079 | ICD 427 | 079_427 | [1636] | 31, 6336, 6401, 6442, | 6938, 7088, 7090, 745 | 2.777 | 1.394 | 1.506 | 0.194 | 42553192, 1.7180851 | 1.383 | -0.634 | 0.791 |
| ICD 079 | ICD 428 | 079_428 | [1636] | 5294, 5327, 5349, 54 | 6938, 7088, 7090, 745 | 2.738 | 1.326 | 1.508 | 0.191 | 63829792, 1.5886524 | 1.411 | -0.506 | 0.766 |
| ICD 079 | ICD 436 | 079_436 | [1636] | [1906, 5328] | 6938, 7088, 7090, 745 | 2.5 | 1.5 | 1.445 | 0.511 | , 1.5, 1.0, 2.0, 1.5, 1.0 | 1 | -0.871 | 0.642 |
| ICD 079 | ICD 437 | 079_437 | [1636] | 3119, 3123, 4763, 511 | 6938, 7088, 7090, 745 | 2.667 | 1.278 | 1.531 | 0.233 | 44444444444442, 1.5 | 1.389 | -0.612 | 0.743 |
| ICD 079 | ICD 438 | 079_438 | [1636] | 906, 4864, 5328, 544 | 6938, 7088, 7090, 745 | 2.5 | 1.5 | 1.446 | 0.367 | .25, 1.5, 1.5, 1.25, 1.5 | 1 | -1.217 | 0.822 |
| ICD 079 | ICD 458 | 079_458 | [1636] | 3, 3558, 3569, 3630, 3 | 6938, 7088, 7090, 745 | 2.709 | 1.345 | 1.469 | 0.194 | 0, 1.6, 1.50909090909 | 1.364 | -0.545 | 0.764 |
| ICD 079 | ICD 482 | 079_482 | [1636] | , 5173, 5594, 5595, 71 | 6938, 7088, 7090, 745 | 2.364 | 1.364 | 1.532 | 0.257 | 181, 1.545454545454 | 1 | -2.068 | 0.964 |
| ICD 079 | ICD 486 | 079_486 | [1636] | [2778, 5173, 7356] | 6938, 7088, 7090, 745 | 2 | 1.333 | 1.477 | 0.388 | 333333333333333, 1. | 0.667 | -2.089 | 0.957 |
| ICD 079 | ICD 496 | 079_496 | [1636] | 12, 4318, 4323, 4843 | 6938, 7088, 7090, 745 | 2.714 | 1.257 | 1.526 | 0.194 | 7142857142857146, 1. | 1.457 | -0.354 | 0.652 |
| ICD 079 | ICD 507 | 079_507 | [1636] | [2778, 5173, 7356] | 6938, 7088, 7090, 745 | 2 | 1.333 | 1.463 | 0.403 | .333333333333333, 1 | 0.667 | -1.977 | 0.951 |
| ICD 079 | ICD 511 | 079_511 | [1636] | 3, 3569, 3586, 5173, 6 | 6938, 7088, 7090, 745 | 2.556 | 1.222 | 1.545 | 0.262 | 4, 1.888888888888888 | 1.333 | -0.805 | 0.741 |
| ICD 079 | ICD 518 | 079_518 | [1636] | 4615, 5173, 6299, 7 | 6938, 7088, 7090, 745 | 2.636 | 1.364 | 1.533 | 0.247 | 363636363636365, 1. | 1.273 | -1.054 | 0.872 |
| ICD 079 | ICD 535 | 079_535 | [1636] | [3553] | 6938, 7088, 7090, 745 | 3 | 1 | 1.604 | 0.574 | , 1.0, 1.0, 2.0, 2.0, 2.0 | 2 | 0.69 | 0.006 |
| ICD 079 | ICD 564 | 079_564 | [1636] | 8, 5443, 5444, 7067, 7 | 6938, 7088, 7090, 745 | 2.889 | 1.556 | 1.536 | 0.267 | 3333333333333, 1.88 | 1.333 | -0.76 | 0.75 |
| ICD 079 | ICD 569 | 079_569 | [1636] | 2, 4843, 4846, 10105, | 6938, 7088, 7090, 745 | 2.5 | 1.333 | 1.553 | 0.241 | 3333333333333, 1.16 | 1.167 | -1.603 | 0.911 |
| ICD 079 | ICD 573 | 079_573 | [1636] | 43, 4864, 5054, 5168, | 6938, 7088, 7090, 745 | 2.783 | 1.434 | 1.541 | 0.189 | 4096385542168, 1.68 | 1.349 | -1.018 | 0.857 |
| ICD 079 | ICD 578 | 079_578 | [1636] | 596, 4057, 5649, 888 | 6938, 7088, 7090, 745 | 2.889 | 1.667 | 1.502 | 0.281 | 1.111111111111111, | 1.222 | -0.994 | 0.803 |
| ICD 079 | ICD 583 | 079_583 | [1636] | [977] | 6938, 7088, 7090, 745 | 3 | 1 | 1.35 | 0.646 | , 1.0, 1.0, 0.0, 1.0, 1.0 | 2 | 1.005 | 0.043 |
| ICD 079 | ICD 584 | 079_584 | [1636] | 6, 4353, 4524, 4780, 4 | 6938, 7088, 7090, 745 | 2.707 | 1.457 | 1.545 | 0.187 | 98, 1.5869565217391 | 1.25 | -1.576 | 0.895 |
| ICD 079 | ICD 585 | 079_585 | [1636] | , 4790, 4803, 5054, 5 | 6938, 7088, 7090, 745 | 2.673 | 1.345 | 1.51 | 0.193 | 3635, 1.79999999999 | 1.327 | -0.948 | 0.841 |
| ICD 079 | ICD 599 | 079_599 | [1636] | 10, 4627, 4843, 5328, | 6938, 7088, 7090, 745 | 2.6 | 1.6 | 1.475 | 0.262 | 1.3, 1.9, 0.899999999 | 1 | -1.811 | 0.956 |
| ICD 079 | ICD 780 | 079_780 | [1636] | 204, 4211, 4282, 4504 | 6938, 7088, 7090, 745 | 2.842 | 1.432 | 1.498 | 0.187 | 918032787, 1.972677 | 1.41 | -0.468 | 0.732 |
| ICD 079 | ICD 782 | 079_782 | [1636] | 30, 3827, 4014, 4598, | 6938, 7088, 7090, 745 | 2.81 | 1.524 | 1.526 | 0.204 | 7142857146, 1.66666 | 1.286 | -1.178 | 0.865 |
| ICD 079 | ICD 783 | 079_783 | [1636] | , 4137, 4160, 4852, 4 | 6938, 7088, 7090, 745 | 3 | 1.5 | 1.506 | 0.19 | 1.2166666666666668 | 1.5 | -0.032 | 0.544 |
| ICD 079 | ICD 785 | 079_785 | [1636] | 60, 3630, 3827, 3952 | 6938, 7088, 7090, 745 | 2.852 | 1.407 | 1.494 | 0.209 | 85185185185186, 1.5 | 1.444 | -0.238 | 0.607 |
| ICD 079 | ICD 786 | 079_786 | [1636] | 3350, 3565, 3827, 48 | 6938, 7088, 7090, 745 | 2.769 | 1.154 | 1.501 | 0.244 | 307692, 1.538461538 | 1.615 | 0.471 | 0.256 |
| ICD 079 | ICD 787 | 079_787 | [1636] | 057, 4645, 5697, 655 | 6938, 7088, 7090, 745 | 2.941 | 1.647 | 1.443 | 0.225 | 470588235294, 1.529 | 1.294 | -0.664 | 0.733 |
| ICD 079 | ICD 788 | 079_788 | [1636] | 906, 3952, 5020, 557 | 6938, 7088, 7090, 745 | 2.5 | 1.333 | 1.418 | 0.315 | 3333333333, 1.83333 | 1.167 | -0.796 | 0.694 |
| ICD 079 | ICD 789 | 079_789 | [1636] | 93, 5184, 5465, 5663, | 6938, 7088, 7090, 745 | 2.694 | 1.444 | 1.567 | 0.192 | 1.6666666666666665 | 1.25 | -1.654 | 0.905 |
| ICD 079 | ICD 799 | 079_799 | [1636] | 4353, 4830, 4842, 48 | 6938, 7088, 7090, 745 | 2.64 | 1.34 | 1.529 | 0.205 | 0000000002, 1.44, 1.4 | 1.3 | -1.12 | 0.848 |
| ICD 079 | ICD 996 | 079_996 | [1636] | [4683] | 6938, 7088, 7090, 745 | 2 | 2 | 1.77 | 0.558 | , 2.0, 2.0, 2.0, 2.0, 1.0 | 0 | -3.172 | 0.985 |
| ICD 162 | ICD 079 | 162_079 | 547, 2597, 2618, 269 | [1636] | 042, 2044, 2046, 206 | 1 | 1 | 0.129 | 0.347 | , 0.0, 0.0, 0.0, 0.0, 0.0 | 0 | -0.372 | 0.125 |
| ICD 162 | ICD 519 | 162_519 | 547, 2597, 2618, 269 | [2778, 5173, 7356] | 042, 2044, 2046, 206 | 1.667 | 1.333 | 0.09 | 0.173 | .0, 0.3333333333333 | 0.333 | 1.404 | 0.033 |
| ICD 197 | ICD 079 | 197_079 | 6648, 6722, 6750, 70 | [1636] | 5318, 5327, 5329, 53 | 2 | 2 | 0.244 | 0.461 | , 0.0, 1.0, 2.0, 0.0, 0.0 | 0 | -0.529 | 0.231 |
| ICD 197 | ICD 519 | 197_519 | 6648, 6722, 6750, 70 | [2778, 5173, 7356] | 5318, 5327, 5329, 53 | 1.667 | 1.333 | 0.28 | 0.269 | 3333333326, 0.33333 | 0.333 | 0.197 | 0.181 |
| ICD 202 | ICD 079 | 202_079 | , 5743, 5819, 5894, 59 | [1636] | , 3123, 3162, 3281, 32 | 2 | 2 | 0.231 | 0.44 | , 0.0, 0.0, 1.0, 0.0, 0.0 | 0 | -0.525 | 0.223 |
| ICD 202 | ICD 519 | 202_519 | , 5743, 5819, 5894, 59 | [2778, 5173, 7356] | , 3123, 3162, 3281, 32 | 1.667 | 1.333 | 0.15 | 0.217 | , 0.0, 0.0, 0.0, 0.33333 | 0.333 | 0.846 | 0.074 |
| ICD 241 | ICD 789 | 241_789 | 693, 6347, 6926, 725 | 93, 5184, 5465, 5663, | 729, 6731, 6902, 692 | 2.181 | 1.444 | 0.742 | 0.066 | , 0.805555555555555 | 0.736 | -0.082 | 0.491 |
| ICD 244 | ICD 079 | 244_079 | 7068, 7080, 7201, 72 | [1636] | 046, 6347, 6388, 652 | 2 | 2 | 0.435 | 0.612 | , 0.0, 1.0, 2.0, 0.0, 1.0 | 0 | -0.711 | 0.373 |
| ICD 244 | ICD 519 | 244_519 | 7068, 7080, 7201, 72 | [2778, 5173, 7356] | 046, 6347, 6388, 652 | 2.333 | 1.333 | 0.541 | 0.322 | 666666666663, 0.333 | 1 | 1.426 | 0.06 |
| ICD 250 | ICD 079 | 250_079 | , 3375, 3383, 3397, 34 | [1636] | , 2717, 2740, 2778, 27 | 0 | 0 | 0.179 | 0.394 | , 0.0, 1.0, 0.0, 0.0, 0.0 | 0 | -0.454 | 0.175 |
| ICD 250 | ICD 519 | 250_519 | , 3375, 3383, 3397, 34 | [2778, 5173, 7356] | , 2717, 2740, 2778, 27 | 1 | 0.667 | 0.136 | 0.203 | 33333333326, 0.3333 | 0.333 | 0.973 | 0.232 |
| ICD 253 | ICD 079 | 253_079 | 6870, 7466, 8820, 882 | [1636] | 4, 5443, 5447, 5449, 5 | 2 | 2 | 0.32 | 0.518 | , 0.0, 1.0, 1.0, 0.0, 0.0 | 0 | -0.618 | 0.296 |
| ICD 272 | ICD 079 | 272_079 | 346, 5424, 5444, 544 | [1636] | 58, 4864, 4884, 4927, | 0 | 0 | 0.281 | 0.486 | , 1.0, 1.0, 0.0, 0.0, 1.0 | 0 | -0.578 | 0.264 |
| ICD 272 | ICD 278 | 272_278 | 346, 5424, 5444, 544 | 023, 4160, 4199, 431 | 58, 4864, 4884, 4927, | 1.734 | 1.379 | 0.401 | 0.039 | 44335, 0.4088669950 | 0.355 | -1.182 | 0.906 |
| ICD 272 | ICD 519 | 272_519 | 346, 5424, 5444, 544 | [2778, 5173, 7356] | 58, 4864, 4884, 4927, | 2 | 1.333 | 0.368 | 0.28 | 3333333326, 0.3333333 | 0.667 | 1.068 | 0.067 |
| ICD 274 | ICD 079 | 274_079 | 443, 7498, 9429, 2290 | [1636] | 731, 6902, 6921, 692 | 2 | 2 | 0.733 | 0.678 | , 1.0, 1.0, 1.0, 0.0, 2.0 | 0 | -1.081 | 0.603 |
| ICD 274 | ICD 519 | 274_519 | 443, 7498, 9429, 2290 | [2778, 5173, 7356] | 731, 6902, 6921, 692 | 2.333 | 1.333 | 0.772 | 0.337 | 0.9999999999999998 | 1 | 0.678 | 0.119 |
| ICD 276 | ICD 079 | 276_079 | 2, 4780, 4879, 5019, 5 | [1636] | 78, 5898, 5910, 5962, | 1 | 1 | 0.405 | 0.584 | , 0.0, 0.0, 1.0, 1.0, 0.0 | 0 | -0.693 | 0.358 |
| ICD 276 | ICD 519 | 276_519 | 2, 4780, 4879, 5019, 5 | [2778, 5173, 7356] | 78, 5898, 5910, 5962, | 1.667 | 1.333 | 0.511 | 0.328 | 333333326, 0.666666 | 0.333 | -0.541 | 0.47 |
| ICD 280 | ICD 079 | 280_079 | , 6521, 6647, 7037, 7 | [1636] | 28, 6729, 6731, 6902, | 2 | 2 | 0.551 | 0.654 | , 0.0, 0.0, 0.0, 0.0, 1.0 | 0 | -0.842 | 0.464 |
| ICD 281 | ICD 079 | 281_079 | 48, 6208, 6521, 7372, | [1636] | , 6521, 6728, 6729, 67 | 2 | 2 | 0.532 | 0.64 | , 0.0, 0.0, 0.0, 1.0, 1.0 | 0 | -0.831 | 0.454 |
| ICD 281 | ICD 519 | 281_519 | 48, 6208, 6521, 7372, | [2778, 5173, 7356] | , 6521, 6728, 6729, 67 | 1.667 | 1 | 0.609 | 0.332 | 666667, 0.333333333 | 0.667 | 0.173 | 0.224 |
| ICD 282 | ICD 079 | 282_079 | 40, 3934, 4524, 4548, | [1636] | 2, 5447, 5538, 5557, 5 | 2 | 2 | 0.353 | 0.539 | , 0.0, 0.0, 0.0, 0.0, 1.0 | 0 | -0.655 | 0.322 |
| ICD 285 | ICD 079 | 285_079 | 5, 6006, 6007, 6223, | [1636] | 47, 5538, 5557, 5558, | 1 | 1 | 0.268 | 0.474 | , 0.0, 0.0, 0.0, 0.0, 1.0 | 0 | -0.566 | 0.255 |
| ICD 285 | ICD 519 | 285_519 | 5, 6006, 6007, 6223, | [2778, 5173, 7356] | 47, 5538, 5557, 5558, | 2 | 1.333 | 0.375 | 0.281 | 33333335, 0.6666666 | 0.667 | 1.04 | 0.065 |
| ICD 288 | ICD 519 | 288_519 | 897, 6023, 6208, 6521 | [2778, 5173, 7356] | 58, 4884, 4893, 4927, | 1.667 | 1.333 | 0.284 | 0.258 | 666666666666667, 0 | 0.333 | 0.191 | 0.178 |
| ICD 294 | ICD 079 | 294_079 | , 5243, 5533, 5578, 5 | [1636] | 3480, 3481, 3482, 361 | 1 | 1 | 0.205 | 0.428 | , 0.0, 0.0, 0.0, 0.0, 0.0 | 0 | -0.479 | 0.195 |
| ICD 332 | ICD 079 | 332_079 | 803, 4842, 4843, 492 | [1636] | 688, 3761, 4015, 409 | 2 | 2 | 0.235 | 0.438 | , 1.0, 0.0, 1.0, 0.0, 0.0 | 0 | -0.536 | 0.229 |
| ICD 332 | ICD 519 | 332_519 | 803, 4842, 4843, 492 | [2778, 5173, 7356] | 688, 3761, 4015, 409 | 1 | 1 | 0.26 | 0.261 | .6666666666666667, | 0 | -0.995 | 0.589 |
| ICD 362 | ICD 079 | 362_079 | 74, 7275, 7287, 7399, | [1636] | 1, 4117, 4139, 4140, 4 | 1 | 1 | 0.219 | 0.426 | , 1.0, 0.0, 0.0, 0.0, 0.0 | 0 | -0.514 | 0.214 |
| ICD 362 | ICD 519 | 362_519 | 74, 7275, 7287, 7399, | [2778, 5173, 7356] | 1, 4117, 4139, 4140, 4 | 2 | 1.333 | 0.249 | 0.263 | , 1.0, 0.0, 0.0, 0.6666 | 0.667 | 1.586 | 0.03 |
| ICD 366 | ICD 079 | 366_079 | 9, 3982, 4094, 4284, 4 | [1636] | 16, 5308, 5309, 5318, | 1 | 1 | 0.284 | 0.494 | , 0.0, 0.0, 0.0, 0.0, 0.0 | 0 | -0.575 | 0.264 |
| ICD 366 | ICD 428 | 366_428 | 9, 3982, 4094, 4284, 4 | 5294, 5327, 5349, 54 | 16, 5308, 5309, 5318, | 1.837 | 1.305 | 0.456 | 0.045 | 205673756, 0.553191 | 0.532 | 1.689 | 0.038 |
| ICD 366 | ICD 519 | 366_519 | 9, 3982, 4094, 4284, 4 | [2778, 5173, 7356] | 16, 5308, 5309, 5318, | 1.667 | 1.333 | 0.397 | 0.284 | 999999998, 0.0, 0.33 | 0.333 | -0.223 | 0.305 |
| ICD 378 | ICD 780 | 378_780 | 864, 5428, 6647, 664 | 204, 4211, 4282, 4504 | 5976, 6046, 6388, 66 | 1.973 | 1.388 | 0.545 | 0.042 | 551912568306, 0.546 | 0.585 | 0.932 | 0.16 |
| ICD 401 | ICD 079 | 401_079 | 40, 2778, 2784, 2796, | [1636] | 040, 2052, 2056, 2108 | 0 | 0 | 0.178 | 0.401 | , 0.0, 0.0, 0.0, 1.0, 0.0 | 0 | -0.444 | 0.171 |
| ICD 401 | ICD 272 | 401_272 | 40, 2778, 2784, 2796, | 346, 5424, 5444, 544 | 040, 2052, 2056, 2108 | 1.211 | 1 | 0.156 | 0.044 | 64, 0.197368421052 | 0.211 | 1.235 | 0.085 |
| ICD 401 | ICD 278 | 401_278 | 40, 2778, 2784, 2796, | 023, 4160, 4199, 431 | 040, 2052, 2056, 2108 | 1.138 | 1.034 | 0.151 | 0.029 | 576354679802956, 0. | 0.103 | -1.622 | 0.947 |
| ICD 401 | ICD 480 | 401_480 | 40, 2778, 2784, 2796, | 636, 2778, 5173, 735 | 040, 2052, 2056, 2108 | 0.75 | 0.5 | 0.137 | 0.178 | .0, 0.25, 0.0, 0.0, 0.25 | 0.25 | 0.634 | 0.09 |
| ICD 401 | ICD 519 | 401_519 | 40, 2778, 2784, 2796, | [2778, 5173, 7356] | 040, 2052, 2056, 2108 | 1 | 0.667 | 0.126 | 0.196 | 3333333335, 0.0, 0.0, | 0.333 | 1.056 | 0.219 |
| ICD 403 | ICD 079 | 403_079 | , 3569, 3579, 3630, 36 | [1636] | , 2697, 2701, 2702, 27 | 0 | 0 | 0.208 | 0.416 | , 0.0, 0.0, 1.0, 0.0, 0.0 | 0 | -0.5 | 0.204 |
| ICD 403 | ICD 519 | 403_519 | , 3569, 3579, 3630, 36 | [2778, 5173, 7356] | , 2697, 2701, 2702, 27 | 1 | 0.667 | 0.17 | 0.233 | 3333335, 0.0, 0.66666 | 0.333 | 0.698 | 0.258 |
| ICD 410 | ICD 079 | 410_079 | 17, 3726, 3764, 3791, | [1636] | 804, 2876, 2920, 293 | 1 | 1 | 0.191 | 0.401 | , 0.0, 1.0, 1.0, 0.0, 0.0 | 0 | -0.476 | 0.188 |
| ICD 410 | ICD 519 | 410_519 | 17, 3726, 3764, 3791, | [2778, 5173, 7356] | 804, 2876, 2920, 293 | 1.333 | 1 | 0.164 | 0.23 | 3326, 0.0, 0.0, 0.0, 0.0 | 0.333 | 0.738 | 0.251 |
| ICD 414 | ICD 079 | 414_079 | 3306, 3313, 3383, 339 | [1636] | , 2671, 2677, 2690, 26 | 0 | 0 | 0.2 | 0.42 | , 1.0, 0.0, 0.0, 0.0, 0.0 | 0 | -0.476 | 0.192 |
| ICD 414 | ICD 519 | 414_519 | 3306, 3313, 3383, 339 | [2778, 5173, 7356] | , 2671, 2677, 2690, 26 | 1.333 | 1 | 0.149 | 0.212 | 3337, 0.0, 0.0, 0.0, 0.3 | 0.333 | 0.87 | 0.267 |
| ICD 427 | ICD 079 | 427_079 | 31, 6336, 6401, 6442, | [1636] | , 3688, 3741, 3752, 3 | 0 | 0 | 0.209 | 0.417 | , 1.0, 0.0, 1.0, 0.0, 0.0 | 0 | -0.502 | 0.205 |
| ICD 427 | ICD 519 | 427_519 | 31, 6336, 6401, 6442, | [2778, 5173, 7356] | , 3688, 3741, 3752, 3 | 1.667 | 1.333 | 0.201 | 0.241 | 3333326, 0.33333333 | 0.333 | 0.547 | 0.103 |
| ICD 428 | ICD 079 | 428_079 | 5294, 5327, 5349, 544 | [1636] | 3357, 3416, 3458, 355 | 0 | 0 | 0.216 | 0.431 | , 0.0, 0.0, 0.0, 0.0, 0.0 | 0 | -0.501 | 0.208 |
| ICD 428 | ICD 519 | 428_519 | 5294, 5327, 5349, 544 | [2778, 5173, 7356] | 3357, 3416, 3458, 355 | 1.667 | 1.333 | 0.249 | 0.258 | .0, 0.3333333333333 | 0.333 | 0.325 | 0.16 |
| ICD 434 | ICD 079 | 434_079 | 4018, 4321, 4524, 48 | [1636] | 715, 4758, 4843, 4846 | 0 | 0 | 0.255 | 0.452 | , 1.0, 0.0, 0.0, 1.0, 0.0 | 0 | -0.564 | 0.248 |
| ICD 436 | ICD 079 | 436_079 | [1906, 5328] | [1636] | 6923, 6938, 7088, 709 | 2 | 2 | 1.06 | 0.606 | , 2.0, 1.0, 2.0, 1.0, 1.0 | 0 | -1.75 | 0.848 |
| ICD 436 | ICD 519 | 436_519 | [1906, 5328] | [2778, 5173, 7356] | 6923, 6938, 7088, 709 | 2.667 | 1.333 | 1.109 | 0.357 | 999999999999998, 0 | 1.333 | 0.63 | 0.117 |
| ICD 437 | ICD 079 | 437_079 | 3119, 3123, 4763, 511 | [1636] | , 5962, 5976, 6046, 63 | 2 | 2 | 0.523 | 0.643 | , 0.0, 0.0, 2.0, 1.0, 0.0 | 0 | -0.813 | 0.443 |
| ICD 437 | ICD 519 | 437_519 | 3119, 3123, 4763, 511 | [2778, 5173, 7356] | , 5962, 5976, 6046, 63 | 1.667 | 1 | 0.604 | 0.321 | 0.3333333333333332 | 0.667 | 0.195 | 0.228 |
| ICD 437 | ICD 783 | 437_783 | 3119, 3123, 4763, 511, | 4137, 4160, 4852, 4 | , 5962, 5976, 6046, 63 | 2.167 | 1.483 | 0.607 | 0.075 | 6666666666665, 0.64 | 0.683 | 1.019 | 0.171 |
| ICD 447 | ICD 996 | 447_996 | 543, 4883, 5444, 562 | [4683] | 6902, 6921, 6923, 693 | 2 | 2 | 1.068 | 0.502 | , 1.0, 1.0, 1.0, 1.0, 1.0 | 0 | -2.129 | 0.907 |
| ICD 458 | ICD 079 | 458_079 | 3, 3558, 3569, 3630, 3 | [1636] | 4879, 4884, 4927, 492 | 0 | 0 | 0.284 | 0.483 | , 0.0, 0.0, 0.0, 0.0, 1.0 | 0 | -0.588 | 0.269 |
| ICD 458 | ICD 519 | 458_519 | 3, 3558, 3569, 3630, 3 | [2778, 5173, 7356] | 4879, 4884, 4927, 492 | 1 | 1 | 0.407 | 0.297 | , 0.333333333333333 | 0 | -1.374 | 0.799 |
| ICD 465 | ICD 519 | 465_519 | [2778, 5173, 7356] | [2778, 5173, 7356] | 6923, 6938, 7088, 709 | 0 | 0 | 1.01 | 0.341 | 1.0, 1.3333333333333 | 0 | -2.96 | 0.994 |
| ICD 482 | ICD 079 | 482_079 | , 5173, 5594, 5595, 71 | [1636] | 6728, 6729, 6731, 69 | 2 | 2 | 0.421 | 0.588 | , 1.0, 0.0, 0.0, 1.0, 0.0 | 0 | -0.716 | 0.37 |
| ICD 486 | ICD 079 | 486_079 | [2778, 5173, 7356] | [1636] | 6923, 6938, 7088, 709 | 2 | 2 | 1.019 | 0.638 | , 1.0, 0.0, 1.0, 1.0, 1.0 | 0 | -1.597 | 0.812 |
| ICD 486 | ICD 519 | 486_519 | [2778, 5173, 7356] | [2778, 5173, 7356] | 6923, 6938, 7088, 709 | 0 | 0 | 1.021 | 0.352 | 9998, 0.66666666666 | 0 | -2.897 | 0.995 |
| ICD 493 | ICD 079 | 493_079 | 843, 4852, 5142, 5144 | [1636] | , 4363, 4589, 4602, 47 | 2 | 2 | 0.231 | 0.451 | , 0.0, 0.0, 0.0, 0.0, 0.0 | 0 | -0.512 | 0.218 |
| ICD 493 | ICD 519 | 493_519 | 843, 4852, 5142, 5144 | [2778, 5173, 7356] | , 4363, 4589, 4602, 47 | 0 | 0 | 0.318 | 0.266 | , 0.333333333333333 | 0 | -1.198 | 0.696 |
| ICD 496 | ICD 079 | 496_079 | 12, 4318, 4323, 4843, | [1636] | 5566, 5576, 5577, 571 | 2 | 2 | 0.285 | 0.488 | , 0.0, 0.0, 0.0, 0.0, 1.0 | 0 | -0.584 | 0.268 |
| ICD 496 | ICD 519 | 496_519 | 12, 4318, 4323, 4843, | [2778, 5173, 7356] | 5566, 5576, 5577, 571 | 0 | 0 | 0.395 | 0.293 | 666666666667, 0.333 | 0 | -1.347 | 0.792 |
| ICD 511 | ICD 079 | 511_079 | 3, 3569, 3586, 5173, 6 | [1636] | 6728, 6729, 6731, 69 | 2 | 2 | 0.619 | 0.67 | , 0.0, 1.0, 1.0, 2.0, 1.0 | 0 | -0.925 | 0.517 |
| ICD 511 | ICD 519 | 511_519 | 3, 3569, 3586, 5173, 6 | [2778, 5173, 7356] | 6728, 6729, 6731, 69 | 0 | 0 | 0.682 | 0.358 | 3326, 0.99999999999 | 0 | -1.907 | 0.944 |
| ICD 518 | ICD 079 | 518_079 | 4615, 5173, 6299, 7 | [1636] | 6728, 6729, 6731, 69 | 2 | 2 | 0.577 | 0.668 | , 0.0, 0.0, 1.0, 0.0, 0.0 | 0 | -0.863 | 0.478 |
| ICD 518 | ICD 519 | 518_519 | 4615, 5173, 6299, 7 | [2778, 5173, 7356] | 6728, 6729, 6731, 69 | 0 | 0 | 0.65 | 0.336 | 666666667, 0.999999 | 0 | -1.936 | 0.943 |
| ICD 519 | ICD 038 | 519_038 | [2778, 5173, 7356] | 6, 3929, 4049, 4282, 4 | 6923, 6938, 7088, 709 | 2.167 | 1.333 | 1.04 | 0.158 | 6666666665, 0.95833 | 0.833 | -1.306 | 0.885 |
| ICD 519 | ICD 197 | 519_197 | [2778, 5173, 7356] | 6648, 6722, 6750, 70 | 6923, 6938, 7088, 709 | 2.299 | 1.328 | 1.062 | 0.124 | 74626868, 1.0746268 | 0.97 | -0.739 | 0.759 |
| ICD 519 | ICD 198 | 519_198 | [2778, 5173, 7356] | 6648, 6722, 6750, 70 | 6923, 6938, 7088, 709 | 2.299 | 1.328 | 1.064 | 0.127 | 268656718, 0.955223 | 0.97 | -0.739 | 0.776 |
| ICD 519 | ICD 250 | 519_250 | [2778, 5173, 7356] | , 3375, 3383, 3397, 34 | 6923, 6938, 7088, 709 | 2.327 | 1.433 | 1.04 | 0.101 | 887820512820513, 1 | 0.894 | -1.449 | 0.975 |
| ICD 519 | ICD 251 | 519_251 | [2778, 5173, 7356] | 3767, 5265, 5465, 66 | 6923, 6938, 7088, 709 | 2.353 | 1.353 | 1.047 | 0.18 | 351, 0.882352941176 | 1 | -0.262 | 0.537 |
| ICD 519 | ICD 274 | 519_274 | [2778, 5173, 7356] | 443, 7498, 9429, 229 | 6923, 6938, 7088, 709 | 2.545 | 1.727 | 1.036 | 0.23 | 63636363638, 1.0, 0.7 | 0.818 | -0.945 | 0.794 |
| ICD 519 | ICD 276 | 519_276 | [2778, 5173, 7356] | 2, 4780, 4879, 5019, 5 | 6923, 6938, 7088, 709 | 2.389 | 1.444 | 1.011 | 0.183 | 77775, 0.9444444444 | 0.944 | -0.364 | 0.619 |
| ICD 519 | ICD 280 | 519_280 | [2778, 5173, 7356] | , 6521, 6647, 7037, 7 | 6923, 6938, 7088, 709 | 2.25 | 1.417 | 1.052 | 0.205 | , 1.083333333333333 | 0.833 | -1.07 | 0.809 |
| ICD 519 | ICD 285 | 519_285 | [2778, 5173, 7356] | 5, 6006, 6007, 6223, | 6923, 6938, 7088, 709 | 2.424 | 1.475 | 1.037 | 0.132 | 6, 0.81355932203389 | 0.949 | -0.67 | 0.723 |
| ICD 519 | ICD 290 | 519_290 | [2778, 5173, 7356] | 4200, 4205, 4353, 45 | 6923, 6938, 7088, 709 | 2.317 | 1.397 | 1.045 | 0.111 | 768844218, 0.984924 | 0.92 | -1.122 | 0.916 |
| ICD 519 | ICD 294 | 519_294 | [2778, 5173, 7356] | , 5243, 5533, 5578, 5 | 6923, 6938, 7088, 709 | 2.322 | 1.385 | 1.037 | 0.111 | 048951048952, 0.818 | 0.937 | -0.901 | 0.826 |
| ICD 519 | ICD 401 | 519_401 | [2778, 5173, 7356] | 40, 2778, 2784, 2796, | 6923, 6938, 7088, 709 | 2.342 | 1.422 | 1.043 | 0.108 | 12979351032452, 0.9 | 0.92 | -1.136 | 0.938 |
| ICD 519 | ICD 410 | 519_410 | [2778, 5173, 7356] | 17, 3726, 3764, 3791, | 6923, 6938, 7088, 709 | 2.221 | 1.32 | 1.063 | 0.123 | 79005524861875, 0.9 | 0.901 | -1.317 | 0.975 |
| ICD 519 | ICD 414 | 519_414 | [2778, 5173, 7356] | 3306, 3313, 3383, 339 | 6923, 6938, 7088, 709 | 2.269 | 1.373 | 1.057 | 0.114 | 678714859439, 1.361 | 0.896 | -1.418 | 0.979 |
| ICD 519 | ICD 427 | 519_427 | [2778, 5173, 7356] | 31, 6336, 6401, 6442, | 6923, 6938, 7088, 709 | 2.367 | 1.404 | 1.035 | 0.107 | 2978723404, 1.03191 | 0.963 | -0.674 | 0.744 |
| ICD 519 | ICD 428 | 519_428 | [2778, 5173, 7356] | 5294, 5327, 5349, 54 | 6923, 6938, 7088, 709 | 2.362 | 1.348 | 1.043 | 0.126 | 29078014184396, 0.7 | 1.014 | -0.229 | 0.497 |
| ICD 519 | ICD 436 | 519_436 | [2778, 5173, 7356] | [1906, 5328] | 6923, 6938, 7088, 709 | 2.5 | 1.5 | 1.042 | 0.534 | , 1.0, 0.5, 1.0, 1.5, 1.0 | 1 | -0.08 | 0.343 |
| ICD 519 | ICD 437 | 519_437 | [2778, 5173, 7356] | 3119, 3123, 4763, 511 | 6923, 6938, 7088, 709 | 2.222 | 1.278 | 1.049 | 0.195 | 8, 0.77777777777777 | 0.944 | -0.539 | 0.644 |
| ICD 519 | ICD 438 | 519_438 | [2778, 5173, 7356] | 906, 4864, 5328, 544 | 6923, 6938, 7088, 709 | 2.25 | 1.5 | 1.002 | 0.358 | 1.0, 1.0, 1.0, 0.75, 1.2 | 0.75 | -0.703 | 0.633 |
| ICD 519 | ICD 458 | 519_458 | [2778, 5173, 7356] | 3, 3558, 3569, 3630, 3 | 6923, 6938, 7088, 709 | 2.236 | 1.382 | 1.022 | 0.14 | 2, 1.218181818181817 | 0.855 | -1.191 | 0.902 |
| ICD 519 | ICD 482 | 519_482 | [2778, 5173, 7356] | , 5173, 5594, 5595, 71 | 6923, 6938, 7088, 709 | 1.818 | 1 | 1.033 | 0.197 | 36363636365, 1.0, 1. | 0.818 | -1.086 | 0.91 |
| ICD 519 | ICD 486 | 519_486 | [2778, 5173, 7356] | [2778, 5173, 7356] | 6923, 6938, 7088, 709 | 0 | 0 | 1.013 | 0.353 | 98, 0.9999999999999 | 0 | -2.866 | 0.993 |
| ICD 519 | ICD 496 | 519_496 | [2778, 5173, 7356] | 12, 4318, 4323, 4843 | 6923, 6938, 7088, 709 | 2.171 | 1.143 | 1.035 | 0.15 | 71428572, 0.9428571 | 1.029 | -0.043 | 0.454 |
| ICD 519 | ICD 511 | 519_511 | [2778, 5173, 7356] | 3, 3569, 3586, 5173, 6 | 6923, 6938, 7088, 709 | 1.556 | 0.778 | 1.056 | 0.234 | 111111111111112, 1. | 0.778 | -1.191 | 0.904 |
| ICD 519 | ICD 518 | 519_518 | [2778, 5173, 7356] | 4615, 5173, 6299, 7 | 6923, 6938, 7088, 709 | 1.818 | 1 | 1.039 | 0.201 | 090909090909092, 0. | 0.818 | -1.094 | 0.916 |
| ICD 519 | ICD 535 | 519_535 | [2778, 5173, 7356] | [3553] | 6923, 6938, 7088, 709 | 3 | 1 | 1.044 | 0.582 | , 2.0, 1.0, 1.0, 2.0, 1.0 | 2 | 1.643 | 0 |
| ICD 519 | ICD 564 | 519_564 | [2778, 5173, 7356] | 8, 5443, 5444, 7067, 7 | 6923, 6938, 7088, 709 | 2.444 | 1.556 | 1.057 | 0.233 | 1111111111114, 1.22 | 0.889 | -0.719 | 0.667 |
| ICD 519 | ICD 569 | 519_569 | [2778, 5173, 7356] | 2, 4843, 4846, 10105, | 6923, 6938, 7088, 709 | 2.333 | 1.333 | 1.056 | 0.208 | 66666663, 1.0833333 | 1 | -0.268 | 0.559 |
| ICD 519 | ICD 573 | 519_573 | [2778, 5173, 7356] | 43, 4864, 5054, 5168, | 6923, 6938, 7088, 709 | 2.325 | 1.434 | 1.045 | 0.125 | 16867469886, 0.9397 | 0.892 | -1.229 | 0.921 |
| ICD 519 | ICD 578 | 519_578 | [2778, 5173, 7356] | 596, 4057, 5649, 888 | 6923, 6938, 7088, 709 | 2.333 | 1.667 | 1.028 | 0.216 | 1111111111, 1.44444 | 0.667 | -1.674 | 0.957 |
| ICD 519 | ICD 584 | 519_584 | [2778, 5173, 7356] | 6, 4353, 4524, 4780, 4 | 6923, 6938, 7088, 709 | 2.38 | 1.446 | 1.041 | 0.122 | , 0.945652173913043 | 0.935 | -0.873 | 0.806 |

| | | | | | | | | | | | | | |
|---|---|---|---|---|---|---|---|---|---|---|---|---|---|
| ICD 519 | ICD 585 | 519_585 | [2778, 5173, 7356] | , 4790, 4803, 5054, 50 | 6923, 6938, 7088, 709 | 2.182 | 1.382 | 1.035 | 0.132 | 1.1636363636363636, | 0.8 | -1.779 | 0.987 |
| ICD 519 | ICD 599 | 519_599 | [2778, 5173, 7356] | 10, 4627, 4843, 5328, | 6923, 6938, 7088, 709 | 2.4 | 1.6 | 1.014 | 0.235 | 99999, 1.40000000000 | 0.8 | -0.912 | 0.803 |
| ICD 519 | ICD 780 | 519_780 | [2778, 5173, 7356] | 204, 4211, 4282, 4504 | 6923, 6938, 7088, 709 | 2.383 | 1.432 | 1.027 | 0.12 | 01639344262295, 1.1 | 0.951 | -0.636 | 0.724 |
| ICD 519 | ICD 783 | 519_783 | [2778, 5173, 7356] | , 4137, 4160, 4852, 49 | 6923, 6938, 7088, 709 | 2.417 | 1.5 | 1.022 | 0.126 | 9, 1.15, 0.9333333333 | 0.917 | -0.839 | 0.798 |
| ICD 519 | ICD 785 | 519_785 | [2778, 5173, 7356] | 360, 3630, 3827, 3952, | 6923, 6938, 7088, 709 | 2.222 | 1.37 | 1.036 | 0.174 | 4074074074074074, 1. | 0.852 | -1.055 | 0.836 |
| ICD 519 | ICD 786 | 519_786 | [2778, 5173, 7356] | 3350, 3565, 3827, 48 | 6923, 6938, 7088, 709 | 2.308 | 1.308 | 1.055 | 0.203 | 3077, 1.307692307692 | 1 | -0.271 | 0.658 |
| ICD 519 | ICD 787 | 519_787 | [2778, 5173, 7356] | 057, 4645, 5697, 6550 | 6923, 6938, 7088, 709 | 2.765 | 1.647 | 1.006 | 0.197 | 0.8235294117647058, | 1.118 | 0.565 | 0.317 |
| ICD 519 | ICD 788 | 519_788 | [2778, 5173, 7356] | 906, 3952, 5020, 5578 | 6923, 6938, 7088, 709 | 2.667 | 1.333 | 0.989 | 0.285 | 99999999999998, 1.1 | 1.333 | 1.211 | 0.063 |
| ICD 519 | ICD 789 | 519_789 | [2778, 5173, 7356] | 93, 5184, 5465, 5663, | 6923, 6938, 7088, 709 | 2.347 | 1.444 | 1.059 | 0.124 | 777781, 1.2916666666 | 0.903 | -1.259 | 0.912 |
| ICD 519 | ICD 799 | 519_799 | [2778, 5173, 7356] | 4353, 4830, 4842, 48 | 6923, 6938, 7088, 709 | 2.2 | 1.3 | 1.041 | 0.134 | 1.1, 1.06, 1.06, 1.2200 | 0.9 | -1.051 | 0.856 |
| ICD 519 | ICD 996 | 519_996 | [2778, 5173, 7356] | [4683] | 6923, 6938, 7088, 709 | 2 | 2 | 1.217 | 0.576 | , 2.0, 2.0, 0.0, 2.0, 1.0 | 0 | -2.111 | 0.926 |
| ICD 530 | ICD 079 | 530_079 | 58, 5743, 5970, 6548, | [1636] | 5862, 5878, 5898, 5910 | 2 | 2 | 0.363 | 0.545 | , 0.0, 0.0, 1.0, 0.0, 0.0 | 0 | -0.665 | 0.33 |
| ICD 530 | ICD 519 | 530_519 | 58, 5743, 5970, 6548, | [2778, 5173, 7356] | 5862, 5878, 5898, 5910 | 2 | 1.333 | 0.44 | 0.3 | 5666663, 0.333333333 | 0.667 | 0.755 | 0.098 |
| ICD 531 | ICD 079 | 531_079 | 53, 3952, 4318, 4843, | [1636] | 78, 5898, 5910, 5962, | 2 | 2 | 0.538 | 0.632 | , 0.0, 1.0, 0.0, 0.0, 0.0 | 0 | -0.852 | 0.466 |
| ICD 531 | ICD 519 | 531_519 | 53, 3952, 4318, 4843, | [2778, 5173, 7356] | 78, 5898, 5910, 5962, | 1.667 | 1.333 | 0.581 | 0.311 | .33333333333333326, | 0.333 | -0.798 | 0.575 |
| ICD 533 | ICD 535 | 533_535 | [1906, 2520, 3553] | [3553] | 6921, 6923, 6938, 708 | 0 | 0 | 0.948 | 0.538 | , 1.0, 1.0, 0.0, 1.0, 1.0 | 0 | -1.762 | 0.828 |
| ICD 535 | ICD 079 | 535_079 | [3553] | [1636] | 6938, 7088, 7090, 745 | 3 | 2 | 1.055 | 0.615 | , 1.0, 2.0, 0.0, 1.0, 2.0 | 1 | -0.089 | 0.2 |
| ICD 535 | ICD 519 | 535_519 | [3553] | [2778, 5173, 7356] | 6938, 7088, 7090, 745 | 3 | 1.333 | 1.103 | 0.353 | , 0.999999999999999 | 1.667 | 1.596 | 0.021 |
| ICD 535 | ICD 626 | 535_626 | [3553] | 586, 1588, 2488, 544 | 6938, 7088, 7090, 745 | 2.667 | 1.333 | 1.069 | 0.286 | 565, 0.833333333333 | 1.333 | 0.924 | 0.11 |
| ICD 536 | ICD 079 | 536_079 | [3162] | [1636] | 7088, 7090, 7458, 746 | 3 | 2 | 1.092 | 0.613 | , 1.0, 2.0, 1.0, 1.0, 1.0 | 1 | -0.15 | 0.214 |
| ICD 536 | ICD 519 | 536_519 | [3162] | [2778, 5173, 7356] | 7088, 7090, 7458, 746 | 2.333 | 1.333 | 1.095 | 0.368 | 66666667, 1.33333333 | 1 | -0.257 | 0.452 |
| ICD 560 | ICD 079 | 560_079 | 6550, 79827, 115019 | [1636] | 6938, 7088, 7090, 745 | 3 | 2 | 1.178 | 0.589 | , 1.0, 2.0, 1.0, 1.0, 1.0 | 1 | -0.302 | 0.25 |
| ICD 560 | ICD 519 | 560_519 | 6550, 79827, 115019 | [2778, 5173, 7356] | 6938, 7088, 7090, 745 | 2.333 | 1.333 | 1.202 | 0.357 | , 1.333333333333333 | 1 | -0.567 | 0.603 |
| ICD 564 | ICD 079 | 564_079 | 8, 5443, 5444, 7067, 7 | [1636] | 6731, 6902, 6921, 69 | 2 | 2 | 0.614 | 0.664 | , 0.0, 0.0, 2.0, 0.0, 1.0 | 0 | -0.924 | 0.517 |
| ICD 564 | ICD 519 | 564_519 | 8, 5443, 5444, 7067, 7 | [2778, 5173, 7356] | 6731, 6902, 6921, 69 | 2.333 | 1.333 | 0.723 | 0.353 | 998, 0.333333333333 | 1 | 0.785 | 0.114 |
| ICD 569 | ICD 079 | 569_079 | 2, 4843, 4846, 10105, | [1636] | 6728, 6729, 6731, 690 | 2 | 2 | 0.628 | 0.684 | , 0.0, 1.0, 2.0, 0.0, 0.0 | 0 | -0.918 | 0.515 |
| ICD 569 | ICD 519 | 569_519 | 2, 4843, 4846, 10105, | [2778, 5173, 7356] | 6728, 6729, 6731, 690 | 2.333 | 1.333 | 0.661 | 0.353 | 3333333326, 0.333333 | 1 | 0.96 | 0.1 |
| ICD 571 | ICD 519 | 571_519 | 33, 4288, 4313, 4353, | [2778, 5173, 7356] | 4087, 4088, 4139, 414 | 1.333 | 1 | 0.23 | 0.248 | 66666666667, 0.6666 | 0.333 | 0.416 | 0.301 |
| ICD 573 | ICD 079 | 573_079 | 43, 4864, 5054, 5168, | [1636] | 53, 4524, 4715, 4758, | 2 | 2 | 0.263 | 0.463 | , 1.0, 2.0, 0.0, 0.0, 0.0 | 0 | -0.568 | 0.253 |
| ICD 573 | ICD 519 | 573_519 | 43, 4864, 5054, 5168, | [2778, 5173, 7356] | 53, 4524, 4715, 4758, | 1.333 | 1 | 0.31 | 0.266 | 3333333333326, 0.0, 0 | 0.333 | 0.089 | 0.339 |
| ICD 574 | ICD 079 | 574_079 | [5244, 374569] | [1636] | 6938, 7088, 7090, 745 | 2 | 2 | 1.241 | 0.598 | , 3.0, 1.0, 1.0, 2.0, 1.0 | 0 | -2.076 | 0.939 |
| ICD 574 | ICD 519 | 574_519 | [5244, 374569] | [2778, 5173, 7356] | 6938, 7088, 7090, 745 | 3 | 1.333 | 1.327 | 0.366 | .9999999999999998, | 1.667 | 0.93 | 0.069 |
| ICD 576 | ICD 079 | 576_079 | 042, 7076, 7097, 7124 | [1636] | 34, 4927, 4928, 5046, | 1 | 1 | 0.254 | 0.456 | , 1.0, 0.0, 0.0, 0.0, 0.0 | 0 | -0.557 | 0.245 |
| ICD 581 | ICD 079 | 581_079 | 7018, 7040, 7225, 749 | [1636] | 47, 5538, 5557, 5558, | 2 | 2 | 0.275 | 0.471 | , 1.0, 1.0, 0.0, 0.0, 0.0 | 0 | -0.584 | 0.264 |
| ICD 583 | ICD 079 | 583_079 | [977] | [1636] | 938, 7088, 7090, 7458 | 3 | 2 | 1.283 | 0.624 | , 3.0, 2.0, 2.0, 2.0, 0.0 | 1 | -0.454 | 0.301 |
| ICD 583 | ICD 519 | 583_519 | [977] | [2778, 5173, 7356] | 938, 7088, 7090, 7458 | 2.667 | 1.333 | 1.401 | 0.372 | 33333, 1.66666666666 | 1.333 | -0.181 | 0.371 |
| ICD 584 | ICD 079 | 584_079 | 6, 4353, 4524, 4780, 4 | [1636] | 3934, 4015, 4036, 413 | 1 | 1 | 0.257 | 0.46 | , 1.0, 1.0, 0.0, 0.0, 0.0 | 0 | -0.559 | 0.247 |
| ICD 584 | ICD 251 | 584_251 | 6, 4353, 4524, 4780, 4 | 3767, 5265, 5465, 66 | 3934, 4015, 4036, 413 | 1.294 | 1.118 | 0.342 | 0.12 | 53, 0.2352941176470 | 0.176 | -1.387 | 0.872 |
| ICD 584 | ICD 519 | 584_519 | 6, 4353, 4524, 4780, 4 | [2778, 5173, 7356] | 3934, 4015, 4036, 413 | 1.667 | 1 | 0.282 | 0.266 | 3333326, 0.333333333 | 0.667 | 1.446 | 0.035 |
| ICD 585 | ICD 079 | 585_079 | , 4790, 4803, 5054, 50 | [1636] | 5318, 5327, 5352, 542 | 0 | 0 | 0.247 | 0.45 | , 0.0, 0.0, 1.0, 1.0, 0.0 | 0 | -0.549 | 0.239 |
| ICD 585 | ICD 519 | 585_519 | , 4790, 4803, 5054, 50 | [2778, 5173, 7356] | 5318, 5327, 5352, 542 | 1.667 | 1.333 | 0.331 | 0.278 | 3333333333333326, 0 | 0.333 | 0.008 | 0.236 |
| ICD 593 | ICD 079 | 593_079 | 092, 6928, 7148, 2577 | [1636] | 6923, 6928, 6938, 708 | 2 | 2 | 0.612 | 0.666 | , 2.0, 1.0, 1.0, 1.0, 2.0 | 0 | -0.919 | 0.511 |
| ICD 593 | ICD 519 | 593_519 | 092, 6928, 7148, 2577 | [2778, 5173, 7356] | 6923, 6928, 6938, 708 | 2.667 | 1.333 | 0.683 | 0.356 | 333333333333326, 0.6 | 1.333 | 1.824 | 0.016 |
| ICD 599 | ICD 079 | 599_079 | 10, 4627, 4843, 5328, | [1636] | 6388, 6728, 6729, 67 | 1 | 1 | 0.559 | 0.667 | , 0.0, 1.0, 1.0, 0.0, 0.0 | 0 | -0.838 | 0.465 |
| ICD 600 | ICD 079 | 600_079 | 4, 367, 2252, 5617, 67 | [1636] | 31, 6902, 6921, 6923, | 2 | 2 | 0.629 | 0.659 | , 1.0, 1.0, 1.0, 1.0, 0.0 | 0 | -0.955 | 0.532 |
| ICD 600 | ICD 519 | 600_519 | 4, 367, 2252, 5617, 67 | [2778, 5173, 7356] | 31, 6902, 6921, 6923, | 2.333 | 1.333 | 0.701 | 0.352 | 3333333326, 0.99999 | 1 | 0.848 | 0.091 |
| ICD 626 | ICD 535 | 626_535 | 586, 1588, 2488, 544 | [3553] | 6731, 6902, 6921, 692 | 2 | 1 | 0.984 | 0.573 | , 1.0, 2.0, 1.0, 0.0, 0.0 | 1 | 0.028 | 0.156 |
| ICD 628 | ICD 041 | 628_041 | 3953, 3972, 3976, 452 | [5243, 5803] | 5594, 5595, 5617, 571 | 1.5 | 1 | 0.462 | 0.398 | , 1.0, 0.5, 0.0, 1.0, 1.0 | 0.5 | 0.094 | 0.231 |
| ICD 692 | ICD 079 | 692_079 | 597, 4598, 9733, 5443 | [1636] | 6921, 6923, 6938, 708 | 2 | 2 | 0.778 | 0.684 | , 0.0, 1.0, 1.0, 0.0, 1.0 | 0 | -1.138 | 0.634 |
| ICD 692 | ICD 519 | 692_519 | 597, 4598, 9733, 5443 | [2778, 5173, 7356] | 6921, 6923, 6938, 708 | 2 | 1.333 | 0.826 | 0.364 | 3333333326, 1.333333 | 0.667 | -0.438 | 0.471 |
| ICD 719 | ICD 079 | 719_079 | [1440, 4057, 4598] | [1636] | 6921, 6923, 6938, 708 | 3 | 2 | 1.035 | 0.6 | , 1.0, 2.0, 1.0, 2.0, 1.0 | 1 | -0.058 | 0.187 |
| ICD 719 | ICD 519 | 719_519 | [1440, 4057, 4598] | [2778, 5173, 7356] | 6921, 6923, 6938, 708 | 2.667 | 1.333 | 1.041 | 0.344 | , 0.666666666666666 | 1.333 | 0.85 | 0.067 |
| ICD 719 | ICD 785 | 719_785 | [1440, 4057, 4598] | 360, 3630, 3827, 3952, | 6921, 6923, 6938, 708 | 2.519 | 1.407 | 1.073 | 0.159 | 000000000000002, 1.0 | 1.111 | 0.242 | 0.375 |
| ICD 724 | ICD 079 | 724_079 | 3791, 4148, 5327, 639 | [1636] | 962, 5976, 6046, 6388 | 2 | 2 | 0.453 | 0.598 | , 0.0, 0.0, 0.0, 1.0, 0.0 | 0 | -0.757 | 0.399 |
| ICD 724 | ICD 519 | 724_519 | 3791, 4148, 5327, 639 | [2778, 5173, 7356] | 962, 5976, 6046, 6388 | 1.667 | 1.333 | 0.567 | 0.339 | 0, 0.666666666666666 | 0.333 | -0.689 | 0.544 |
| ICD 729 | ICD 079 | 729_079 | 7290, 7353, 8214, 822 | [1636] | 6388, 6728, 6729, 673 | 2 | 2 | 0.362 | 0.532 | , 0.0, 0.0, 0.0, 2.0, 1.0 | 0 | -0.68 | 0.336 |
| ICD 729 | ICD 519 | 729_519 | 7290, 7353, 8214, 822 | [2778, 5173, 7356] | 6388, 6728, 6729, 673 | 2.333 | 1.333 | 0.477 | 0.31 | 66666666666663, 0.33 | 1 | 1.689 | 0.05 |
| ICD 733 | ICD 079 | 733_079 | 5315, 5341, 5443, 568 | [1636] | 4140, 4218, 4240, 428 | 2 | 2 | 0.223 | 0.426 | , 0.0, 1.0, 1.0, 0.0, 0.0 | 0 | -0.524 | 0.219 |
| ICD 733 | ICD 519 | 733_519 | 5315, 5341, 5443, 568 | [2778, 5173, 7356] | 4140, 4218, 4240, 428 | 1.667 | 1.333 | 0.282 | 0.272 | 3333333333333326, 0. | 0.333 | 0.19 | 0.186 |
| ICD 780 | ICD 079 | 780_079 | 204, 4211, 4282, 4504 | [1636] | 2892, 2897, 2925, 296 | 1 | 1 | 0.215 | 0.425 | , 0.0, 0.0, 0.0, 0.0, 0.0 | 0 | -0.505 | 0.209 |
| ICD 781 | ICD 079 | 781_079 | , 4626, 4864, 5443, 63 | [1636] | 817, 5861, 5862, 5878 | 2 | 2 | 0.399 | 0.588 | , 0.0, 1.0, 0.0, 1.0, 0.0 | 0 | -0.678 | 0.347 |
| ICD 781 | ICD 519 | 781_519 | , 4626, 4864, 5443, 63 | [2778, 5173, 7356] | 817, 5861, 5862, 5878 | 2 | 1.333 | 0.527 | 0.318 | 26, 0.9999999999999 | 0.667 | 0.44 | 0.151 |
| ICD 782 | ICD 079 | 782_079 | 30, 3827, 4014, 4598, | [1636] | 538, 5557, 5558, 5566 | 2 | 2 | 0.336 | 0.525 | , 0.0, 0.0, 0.0, 0.0, 1.0 | 0 | -0.64 | 0.311 |
| ICD 782 | ICD 519 | 782_519 | 30, 3827, 4014, 4598, | [2778, 5173, 7356] | 538, 5557, 5558, 5566 | 1.667 | 1.333 | 0.448 | 0.303 | .33333333333333326, | 0.333 | -0.38 | 0.382 |
| ICD 784 | ICD 519 | 784_519 | 3569, 3897, 4864, 502 | [2778, 5173, 7356] | 046, 6388, 6728, 6729, | 2 | 1.333 | 0.732 | 0.346 | 65, 0.9999999999999 | 0.667 | -0.19 | 0.366 |
| ICD 785 | ICD 079 | 785_079 | 360, 3630, 3827, 3952, | [1636] | 4, 5817, 5861, 5862, 5 | 2 | 2 | 0.404 | 0.575 | , 1.0, 0.0, 0.0, 0.0, 1.0 | 0 | -0.702 | 0.36 |
| ICD 786 | ICD 079 | 786_079 | 3350, 3565, 3827, 48 | [1636] | 046, 6388, 6728, 6729 | 0 | 0 | 0.68 | 0.69 | , 1.0, 1.0, 1.0, 1.0, 1.0 | 0 | -0.986 | 0.557 |
| ICD 788 | ICD 011 | 788_011 | 906, 3952, 5020, 5578 | 47, 6556, 7097, 1115 | 731, 6902, 6921, 692 | 2 | 1.462 | 0.69 | 0.169 | .7692307692307692, | 0.538 | -0.899 | 0.82 |
| ICD 788 | ICD 079 | 788_079 | 906, 3952, 5020, 5578 | [1636] | 731, 6902, 6921, 692 | 2 | 2 | 0.636 | 0.669 | , 0.0, 1.0, 0.0, 1.0, 0.0 | 0 | -0.95 | 0.532 |
| ICD 788 | ICD 519 | 788_519 | 906, 3952, 5020, 5578 | [2778, 5173, 7356] | 731, 6902, 6921, 692 | 2.667 | 1.333 | 0.685 | 0.347 | 3333333333304, 0.999 | 1.333 | 1.869 | 0.013 |
| ICD 789 | ICD 309 | 789_309 | 93, 5184, 5465, 5663, | [1815] | , 4846, 4851, 4884, 48 | 2 | 2 | 0.394 | 0.574 | , 0.0, 0.0, 1.0, 1.0, 1.0 | 0 | -0.687 | 0.349 |
| ICD 790 | ICD 079 | 790_079 | 0, 5327, 7369, 56606, | [1636] | 921, 6923, 6938, 7088 | 2 | 2 | 0.77 | 0.681 | , 1.0, 0.0, 1.0, 0.0, 0.0 | 0 | -1.131 | 0.631 |
| ICD 790 | ICD 278 | 790_278 | 0, 5327, 7369, 56606, | 4023, 4160, 4199, 431 | 921, 6923, 6938, 7088 | 2.256 | 1.498 | 0.802 | 0.058 | 56650246308, 0.8128 | 0.759 | -0.753 | 0.767 |
| ICD 790 | ICD 386 | 790_386 | 0, 5327, 7369, 56606, | [1837, 5579, 84908] | 921, 6923, 6938, 7088 | 2 | 1 | 0.894 | 0.312 | 99999999998, 0.9999 | 1 | 0.338 | 0.17 |
| ICD 790 | ICD 519 | 790_519 | 0, 5327, 7369, 56606, | [2778, 5173, 7356] | 921, 6923, 6938, 7088 | 2 | 1.333 | 0.828 | 0.357 | 6663, 0.33333333333 | 0.667 | -0.451 | 0.489 |
| ICD 996 | ICD 079 | 996_079 | [4683] | [1636] | 6938, 7088, 7090, 745 | 2 | 2 | 1.043 | 0.637 | , 1.0, 1.0, 0.0, 0.0, 2.0 | 0 | -1.638 | 0.83 |
| ICD 996 | ICD 519 | 996_519 | [4683] | [2778, 5173, 7356] | 6938, 7088, 7090, 745 | 2.667 | 1.333 | 1.081 | 0.37 | , 0.999999999999999 | 1.333 | 0.682 | 0.121 |
