## Supplementary Table 6 for "Deciphering the Molecular Mechanism of Post-Acute Sequelae of COVID-19 through Comorbidity Network Analysis"

| ONTOLOGY | ID | Description | pvalue | p.adjust | qvalue | geneID | Count |
|---|---|---|---|---|---|---|---|
| BP | GO:0019915 | lipid storage | 0 | 0 | 0 | 7124/3553/335/857/10577/23411/79068/5465/948/1401/3569/3952 | 18 |
| BP | GO:1905952 | regulation of lipid localization | 0 | 0 | 0 | 124/3553/208/335/857/1071/23411/9370/5465/948/1401/3569/395 | 19 |
| BP | GO:0006869 | lipid transport | 0 | 0 | 0 | /335/857/1071/10577/3990/5743/23411/9370/5465/948/3952/5443/ | 24 |
| BP | GO:1901653 | cellular response to peptide | 0 | 0 | 0 | /2671/208/857/3416/5770/23411/5573/26291/9370/7351/948/2641/ | 22 |
| BP | GO:0019216 | regulation of lipid metabolic process | 0 | 0 | 0 | 53/208/335/857/10577/5743/23411/6647/26291/9370/5465/2784/36 | 21 |
| BP | GO:0016042 | lipid catabolic process | 0 | 0 | 0 | /3630/7124/3553/4758/34/208/3990/26291/9370/5465/3290/3667/ | 20 |
| BP | GO:0031667 | response to nutrient levels | 0 | 0 | 0 | 48/4023/7124/1312/34/6581/5743/23411/26291/9370/7351/4968/54 | 23 |
| BP | GO:0062012 | regulation of small molecule metabolic process | 0 | 0 | 0 | 4/3553/34/208/335/857/5743/23411/6647/9370/5465/3953/2641/27 | 20 |
| BP | GO:0045834 | positive regulation of lipid metabolic process | 0 | 0 | 0 | 48/3630/7124/3553/208/335/5743/26291/9370/5465/3667/949/355 | 15 |
| BP | GO:0001659 | temperature homeostasis | 0 | 0 | 0 | 124/3553/857/5743/79068/26291/9370/7351/3953/948/3952/2834 | 16 |
| BP | GO:0010883 | regulation of lipid storage | 0 | 0 | 0 | 5468/4023/7124/23411/79068/5465/948/1401/3569/3952/949 | 11 |
| BP | GO:0030301 | cholesterol transport | 0 | 0 | 0 | 3/348/335/857/1071/10577/3990/23411/9370/948/3952/949/4864/3 | 14 |
| BP | GO:0008203 | cholesterol metabolic process | 0 | 0 | 0 | 335/1071/10577/3990/6647/3953/2784/3952/949/8431/4864/3949/ | 14 |
| BP | GO:0015918 | sterol transport | 0 | 0 | 0 | 3/348/335/857/1071/10577/3990/23411/9370/948/3952/949/4864/3 | 14 |
| BP | GO:0003018 | vascular process in circulatory system | 0 | 0 | 0 | 2150/2876/348/3630/7124/857/6581/5743/6647/3953/948/1401/395 | 17 |
| BP | GO:0050727 | regulation of inflammatory response | 0 | 0 | 0 | /4023/3630/7124/3553/335/5743/6647/9370/7133/5465/196/3569/5 | 20 |
| BP | GO:1902652 | secondary alcohol metabolic process | 0 | 0 | 0 | 335/1071/10577/3990/6647/3953/2784/3952/949/8431/4864/3949/ | 14 |
| BP | GO:0043434 | response to peptide hormone | 0 | 0 | 0 | 208/857/3416/5743/5770/23411/5573/26291/9370/7351/5465/2641 | 20 |
| BP | GO:0008202 | steroid metabolic process | 0 | 0 | 0 | 071/10577/3990/23411/6647/3953/2784/3290/3952/949/3552/8431/ | 18 |
| BP | GO:0016125 | sterol metabolic process | 0 | 0 | 0 | 335/1071/10577/3990/6647/3953/2784/3952/949/8431/4864/3949/ | 14 |
| BP | GO:0045444 | fat cell differentiation | 0 | 0 | 0 | 5468/4023/3630/7124/208/5743/23411/79068/9370/2784/3569/395 | 16 |
| BP | GO:0051235 | maintenance of location | 0 | 0 | 0 | 7124/3553/335/857/10577/23411/79068/5465/948/1401/3569/3952 | 18 |
| BP | GO:0042593 | glucose homeostasis | 0 | 0 | 0 | 468/3630/23411/26291/9370/7351/3953/2641/3383/3569/3667/395 | 16 |
| BP | GO:0033500 | carbohydrate homeostasis | 0 | 0 | 0 | 468/3630/23411/26291/9370/7351/3953/2641/3383/3569/3667/395 | 16 |
| BP | GO:0015850 | organic hydroxy compound transport | 0 | 0 | 0 | 5/857/1071/10577/3990/6581/23411/9370/948/3952/5443/949/843 | 17 |
| BP | GO:0071375 | cellular response to peptide hormone stimulus | 0 | 0 | 0 | 30/3553/208/857/3416/5770/23411/5573/26291/9370/7351/2641/36 | 17 |
| BP | GO:1905954 | positive regulation of lipid localization | 0 | 0 | 0 | 5468/348/4023/3553/335/857/1071/23411/9370/948/949/3552 | 12 |
| BP | GO:0035296 | regulation of tube diameter | 0 | 0 | 0 | 636/6648/2150/2876/348/3630/7124/857/5743/6647/1401/3952/15 | 13 |
| BP | GO:0045598 | regulation of fat cell differentiation | 0 | 0 | 0 | 3/5468/4023/3630/7124/5743/23411/79068/9370/2784/3569/3952/5 | 13 |
| BP | GO:0097746 | blood vessel diameter maintenance | 0 | 0 | 0 | 636/6648/2150/2876/348/3630/7124/857/5743/6647/1401/3952/15 | 13 |
| BP | GO:0035150 | regulation of tube size | 0 | 0 | 0 | 636/6648/2150/2876/348/3630/7124/857/5743/6647/1401/3952/15 | 13 |
| BP | GO:0062013 | positive regulation of small molecule metabolic process | 0 | 0 | 0 | 468/348/3630/7124/3553/208/335/5743/23411/9370/5465/2641/36 | 13 |
| BP | GO:0009743 | response to carbohydrate | 0 | 0 | 0 | 8/2876/4023/7124/3553/5743/26291/9370/7351/2641/3383/3952/3 | 15 |
| BP | GO:0032368 | regulation of lipid transport | 0 | 0 | 0 | 468/348/3553/208/335/857/1071/23411/9370/5465/3952/5443/355 | 13 |
| BP | GO:0006909 | phagocytosis | 0 | 0 | 0 | 150/3688/7124/3553/335/6347/6647/9370/3953/948/1401/3952/94 | 15 |
| BP | GO:0097006 | regulation of plasma lipoprotein particle levels | 0 | 0 | 0 | 3162/348/4023/335/1071/3990/26291/9370/948/949/3949 | 11 |
| BP | GO:0006979 | response to oxidative stress | 0 | 0 | 0 | 3066/2876/348/10516/5743/23411/6647/9370/7351/4968/948/3569 | 18 |
| BP | GO:0032768 | regulation of monooxygenase activity | 0 | 0 | 0 | 2717/348/3630/7124/3553/3952/949/3552/5447 | 9 |
| BP | GO:0000302 | response to reactive oxygen species | 0 | 0 | 0 | 7356/6648/3066/2876/348/10516/23411/6647/7351/948/3569/4318 | 14 |
| BP | GO:0034284 | response to monosaccharide | 0 | 0 | 0 | 648/2876/4023/7124/5743/26291/9370/7351/2641/3383/3952/355 | 14 |
| BP | GO:0042060 | wound healing | 0 | 0 | 0 | 7/2150/2876/5468/348/3688/3630/7124/857/5265/5465/948/3569/ | 18 |
| BP | GO:0042632 | cholesterol homeostasis | 0 | 0 | 0 | 348/4023/335/857/1071/10577/3990/23411/949/4864/3949 | 11 |
| BP | GO:0033344 | cholesterol efflux | 0 | 0 | 0 | 5468/348/335/857/1071/10577/23411/9370/949/4864 | 10 |
| BP | GO:0055092 | sterol homeostasis | 0 | 0 | 0 | 348/4023/335/857/1071/10577/3990/23411/949/4864/3949 | 11 |
| BP | GO:0002237 | response to molecule of bacterial origin | 0 | 0 | 0 | 648/3066/7124/3553/2671/5743/6347/7133/196/948/3569/5054/43 | 17 |
| BP | GO:0010906 | regulation of glucose metabolic process | 0 | 0 | 0 | 3630/34/208/23411/9370/5465/3953/2641/2784/3667/3952 | 11 |
| BP | GO:0055088 | lipid homeostasis | 0 | 0 | 0 | 468/348/4023/3630/335/857/1071/10577/3990/23411/949/4864/39 | 13 |
| BP | GO:0043255 | regulation of carbohydrate biosynthetic process | 0 | 0 | 0 | 3630/34/208/23411/9370/5465/3953/2641/3667/3952/5167 | 11 |
| BP | GO:0015718 | monocarboxylic acid transport | 0 | 0 | 0 | 636/5468/348/7124/3553/208/6581/5743/5465/948/3952/3552/843 | 13 |
| BP | GO:0006641 | triglyceride metabolic process | 0 | 0 | 0 | 2876/348/4023/857/1071/3990/23411/26291/2784/949/3949 | 11 |
| BP | GO:0010827 | regulation of glucose transmembrane transport | 0 | 0 | 0 | 1636/3630/7124/3553/208/26291/9370/3667/3952/5167 | 10 |
| BP | GO:0034381 | plasma lipoprotein particle clearance | 0 | 0 | 0 | 3162/348/335/3990/26291/9370/948/949/3949 | 9 |
| BP | GO:0008217 | regulation of blood pressure | 0 | 0 | 0 | 36/6648/2150/5468/7124/5743/6647/9370/5465/2784/3952/5443/1 | 13 |
| BP | GO:0006109 | regulation of carbohydrate metabolic process | 0 | 0 | 0 | 30/34/208/23411/9370/5465/3953/2641/2784/3667/3952/5443/51 | 13 |
| BP | GO:0006631 | fatty acid metabolic process | 0 | 0 | 0 | 5468/4023/3630/3553/34/208/857/3990/5743/23411/9370/5465/94 | 17 |
| BP | GO:0051051 | negative regulation of transport | 0 | 0 | 0 | 0/348/3630/7124/3553/208/857/5743/9370/7351/7133/948/3667/3 | 18 |
| BP | GO:0032496 | response to lipopolysaccharide | 0 | 0 | 0 | /6648/3066/7124/3553/2671/5743/6347/7133/948/3569/5054/431 | 16 |
| BP | GO:2001234 | negative regulation of apoptotic signaling pathway | 0 | 0 | 0 | 648/2876/3630/7124/3553/5743/5770/23411/3383/5054/4318/746 | 14 |
| BP | GO:0046324 | regulation of glucose import | 0 | 0 | 0 | 1636/3630/7124/208/26291/9370/3667/3952/5167 | 9 |
| BP | GO:0055094 | response to lipoprotein particle | 0 | 0 | 0 | 5468/348/4023/3688/26291/948/4864/3949 | 8 |
| BP | GO:0046942 | carboxylic acid transport | 0 | 0 | 0 | 3/348/3688/7124/3553/208/6581/5743/7351/5465/948/3952/3552/ | 16 |
| BP | GO:0015849 | organic acid transport | 0 | 0 | 0 | 3/348/3688/7124/3553/208/6581/5743/7351/5465/948/3952/3552/ | 16 |
| BP | GO:0050994 | regulation of lipid catabolic process | 0 | 0 | 0 | 3630/7124/3553/208/26291/5465/3667/949/3949 | 9 |
| BP | GO:0032102 | negative regulation of response to external stimulus | 0 | 0 | 0 | 6/5468/348/3630/7124/2671/335/6347/6647/9370/7133/5465/196/3 | 18 |
| BP | GO:0046883 | regulation of hormone secretion | 0 | 0 | 0 | 5468/3630/7124/3553/5573/9370/7351/2641/3569/3667/3952/5443 | 14 |
| BP | GO:0010878 | cholesterol storage | 0 | 0 | 0 | 5468/4023/10577/5465/948/949/4864 | 7 |
| BP | GO:0071402 | cellular response to lipoprotein particle stimulus | 0 | 0 | 0 | 5468/348/4023/3688/26291/948/4864/3949 | 8 |
| BP | GO:0050708 | regulation of protein secretion | 0 | 0 | 0 | /5468/348/3630/7124/3553/5573/7351/2641/3569/3667/3952/3552/ | 14 |
| BP | GO:0006639 | acylglycerol metabolic process | 0 | 0 | 0 | 2876/348/4023/857/1071/3990/23411/26291/2784/949/3949 | 11 |
| BP | GO:0006638 | neutral lipid metabolic process | 0 | 0 | 0 | 2876/348/4023/857/1071/3990/23411/26291/2784/949/3949 | 11 |
| BP | GO:0019217 | regulation of fatty acid metabolic process | 0 | 0 | 0 | 5468/3630/3553/208/857/5743/23411/9370/5465/3667 | 10 |
| BP | GO:0044242 | cellular lipid catabolic process | 0 | 0 | 0 | 717/4023/4758/34/208/3990/26291/9370/5465/3667/3952/949/394 | 13 |
| BP | GO:0120162 | positive regulation of cold-induced thermogenesis | 0 | 0 | 0 | 2778/857/26291/9370/7351/3953/948/3952/283455/153 | 10 |
| BP | GO:0042311 | vasodilation | 0 | 0 | 0 | 6648/2150/2876/348/3630/7124/6647/153 | 8 |
| BP | GO:0006953 | acute-phase response | 0 | 0 | 0 | 3630/7124/3553/5265/5743/1401/3569/3552 | 8 |
| BP | GO:2001233 | regulation of apoptotic signaling pathway | 0 | 0 | 0 | 2876/3630/7124/3553/857/5743/5770/23411/6647/3383/5054/431 | 16 |
| BP | GO:0070482 | response to oxygen levels | 0 | 0 | 0 | 2/5327/6648/3066/5468/7124/857/5743/23411/9370/7351/5465/3 | 15 |
| BP | GO:0050999 | regulation of nitric-oxide synthase activity | 0 | 0 | 0 | 2717/348/3630/3553/3952/949/3552 | 7 |
| BP | GO:0046323 | glucose import | 0 | 0 | 0 | 1636/3630/7124/208/26291/9370/3667/3952/5167 | 9 |
| BP | GO:0046890 | regulation of lipid biosynthetic process | 0 | 0 | 0 | 348/3630/7124/3553/5743/23411/6647/5465/3952/949/3552/3949 | 12 |
| BP | GO:0030100 | regulation of endocytosis | 0 | 0 | 0 | 0/348/7124/57534/3553/335/857/5770/6347/6647/9370/948/5054/ | 14 |
| BP | GO:0006006 | glucose metabolic process | 0 | 0 | 0 | 3630/7124/34/208/23411/9370/5465/3953/2641/2784/3667/3952 | 12 |
| BP | GO:0006111 | regulation of gluconeogenesis | 0 | 0 | 0 | 3630/34/23411/9370/5465/3953/2641/3952 | 8 |
| BP | GO:0001666 | response to hypoxia | 0 | 0 | 0 | 3162/5327/6648/5468/7124/857/5743/23411/9370/7351/5465/3952 | 14 |
| BP | GO:1904035 | regulation of epithelial cell apoptotic process | 0 | 0 | 0 | 3162/7124/6347/26291/5465/3383/3569/5054/7466/4864 | 10 |
| BP | GO:0010743 | regulation of macrophage derived foam cell differentiation | 0 | 0 | 0 | 5468/4023/1071/9370/5465/948/1401 | 7 |
| BP | GO:0030730 | sequestering of triglyceride | 0 | 0 | 0 | 5468/4023/7124/3553/5465/5167 | 6 |
| BP | GO:0051341 | regulation of oxidoreductase activity | 0 | 0 | 0 | 2717/348/3630/7124/3553/3952/949/3552/5447 | 9 |
| BP | GO:0015908 | fatty acid transport | 0 | 0 | 0 | 1636/5468/348/3553/208/5743/5465/948/3952/3552 | 10 |
| BP | GO:1904659 | glucose transmembrane transport | 0 | 0 | 0 | 1636/3630/7124/3553/208/26291/9370/3667/3952/5167 | 10 |
| BP | GO:0046879 | hormone secretion | 0 | 0 | 0 | 5468/3630/7124/3553/5573/9370/7351/2641/3569/3667/3952/5443 | 14 |
| BP | GO:0009410 | response to xenobiotic stimulus | 0 | 0 | 0 | 6648/3066/2876/7124/3553/5743/6647/26291/9370/4968/196/4363 | 16 |
| BP | GO:0008645 | hexose transmembrane transport | 0 | 0 | 0 | 1636/3630/7124/3553/208/26291/9370/3667/3952/5167 | 10 |
| BP | GO:0032869 | cellular response to insulin stimulus | 0 | 0 | 0 | 5468/3630/3553/208/3416/5770/23411/9370/7351/3667/3952/5167 | 12 |
| BP | GO:0032370 | positive regulation of lipid transport | 0 | 0 | 0 | 5468/348/3553/335/857/1071/23411/9370/3552 | 9 |
| BP | GO:0036293 | response to decreased oxygen levels | 0 | 0 | 0 | 3162/5327/6648/5468/7124/857/5743/23411/9370/7351/5465/3952 | 14 |
| BP | GO:0015711 | organic anion transport | 0 | 0 | 0 | 3/348/3688/7124/3553/208/6581/5743/7351/5465/948/3952/3552/ | 16 |
| BP | GO:0015749 | monosaccharide transmembrane transport | 0 | 0 | 0 | 1636/3630/7124/3553/208/26291/9370/3667/3952/5167 | 10 |
| BP | GO:0009914 | hormone transport | 0 | 0 | 0 | 5468/3630/7124/3553/5573/9370/7351/2641/3569/3667/3952/5443 | 14 |
| BP | GO:0071216 | cellular response to biotic stimulus | 0 | 0 | 0 | 124/3553/2671/6347/7133/196/948/3569/5054/4318/7466/949/355 | 13 |
| BP | GO:0046889 | positive regulation of lipid biosynthetic process | 0 | 0 | 0 | 348/3630/7124/3553/5743/5465/949/3552/3949 | 9 |
| BP | GO:0032373 | positive regulation of sterol transport | 0 | 0 | 0 | 5468/348/335/857/1071/23411/9370 | 7 |
| BP | GO:0032376 | positive regulation of cholesterol transport | 0 | 0 | 0 | 5468/348/335/857/1071/23411/9370 | 7 |
| BP | GO:0071404 | cellular response to low-density lipoprotein particle stimulus | 0 | 0 | 0 | 5468/4023/3688/26291/948/4864/3949 | 7 |
| BP | GO:0032868 | response to insulin | 0 | 0 | 0 | 58/3630/3553/208/3416/5770/23411/9370/7351/5465/3667/3952/5 | 13 |
| BP | GO:0048660 | regulation of smooth muscle cell proliferation | 0 | 0 | 0 | 3162/6648/1786/5468/348/7124/9510/5743/9370/3569/4318 | 11 |

| | | | | | | | |
|---|---|---|---|---|---|---|---|
| BP | GO:0051048 | negative regulation of secretion | 0 | 0 | 0 | 7879/2150/348/3630/7124/3553/9370/7351/7133/3667/3952 | 11 |
| BP | GO:0010888 | negative regulation of lipid storage | 0 | 0 | 0 | 5468/7124/5465/1401/3569/3952 | 6 |
| BP | GO:0098856 | intestinal lipid absorption | 0 | 0 | 0 | 335/948/3952/949/4864/3949 | 6 |
| BP | GO:0010742 | macrophage derived foam cell differentiation | 0 | 0 | 0 | 5468/4023/1071/9370/5465/948/1401 | 7 |
| BP | GO:0048659 | smooth muscle cell proliferation | 0 | 0 | 0 | 3162/6648/1786/5468/348/7124/9510/5743/9370/3569/4318 | 11 |
| BP | GO:0045923 | positive regulation of fatty acid metabolic process | 0 | 0 | 0 | 5468/3553/208/5743/9370/5465/3667 | 7 |
| BP | GO:0090077 | foam cell differentiation | 0 | 0 | 0 | 5468/4023/1071/9370/5465/948/1401 | 7 |
| BP | GO:0150077 | regulation of neuroinflammatory response | 0 | 0 | 0 | 7124/3553/5743/7133/3569/4318/3949 | 7 |
| BP | GO:0043410 | positive regulation of MAPK cascade | 0 | 0 | 0 | /348/3630/7124/3553/5770/6347/6647/26291/948/2641/3383/3569/ | 16 |
| BP | GO:0034219 | carbohydrate transmembrane transport | 0 | 0 | 0 | 1636/3630/7124/3553/208/26291/9370/3667/3952/5167 | 10 |
| BP | GO:0043086 | negative regulation of catalytic activity | 0 | 0 | 0 | /2876/5468/348/3630/7124/3553/857/5265/5770/23411/5573/9370/ | 16 |
| BP | GO:0050764 | regulation of phagocytosis | 0 | 0 | 0 | 2150/7124/3553/335/6347/6647/9370/948/949 | 9 |
| BP | GO:0019318 | hexose metabolic process | 0 | 0 | 0 | 3630/7124/34/208/23411/9370/5465/3953/2641/2784/3667/3952 | 12 |
| BP | GO:0023061 | signal release | 0 | 0 | 0 | 3630/7124/3553/5743/5573/9370/7351/2641/3569/3667/3952/5443 | 16 |
| BP | GO:0055090 | acylglycerol homeostasis | 0 | 0 | 0 | 348/4023/335/1071/3990/23411/949 | 7 |
| BP | GO:0070328 | triglyceride homeostasis | 0 | 0 | 0 | 348/4023/335/1071/3990/23411/949 | 7 |
| BP | GO:0072593 | reactive oxygen species metabolic process | 0 | 0 | 0 | 6648/2150/2876/10516/3630/7124/6647/7351/5465/948/1401/3952 | 12 |
| BP | GO:0034370 | triglyceride-rich lipoprotein particle remodeling | 0 | 0 | 0 | 348/4023/335/1071/3990 | 5 |
| BP | GO:0034372 | very-low-density lipoprotein particle remodeling | 0 | 0 | 0 | 348/4023/335/1071/3990 | 5 |
| BP | GO:0071219 | cellular response to molecule of bacterial origin | 0 | 0 | 0 | 7124/3553/2671/6347/7133/196/948/3569/5054/4318/949/3552 | 12 |
| BP | GO:1900076 | regulation of cellular response to insulin stimulus | 0 | 0 | 0 | 5468/3630/3553/5770/23411/3667/3952/5167 | 8 |
| BP | GO:0019218 | regulation of steroid metabolic process | 0 | 0 | 0 | 348/7124/335/23411/6647/2784/3952/3552/3949 | 9 |
| BP | GO:0010565 | regulation of cellular ketone metabolic process | 0 | 0 | 0 | 5468/3630/3553/208/857/5743/23411/9370/5465/3667 | 10 |
| BP | GO:1904019 | epithelial cell apoptotic process | 0 | 0 | 0 | 3162/7124/6347/26291/5465/3383/3569/5054/7466/4864 | 10 |
| BP | GO:0001819 | positive regulation of cytokine production | 0 | 0 | 0 | 2150/4023/3630/7124/3553/5743/23411/6647/9370/948/3569/3952 | 16 |
| BP | GO:0006809 | nitric oxide biosynthetic process | 0 | 0 | 0 | 6648/2717/7124/3553/857/5743/948/1727 | 8 |
| BP | GO:2000377 | regulation of reactive oxygen species metabolic process | 0 | 0 | 0 | 6648/2150/10516/3630/7124/6647/5465/948/1401/3952 | 10 |
| BP | GO:0033002 | muscle cell proliferation | 0 | 0 | 0 | 3162/6648/1786/5468/348/7124/9510/5743/5573/9370/3569/4318 | 12 |
| BP | GO:0090276 | regulation of peptide hormone secretion | 0 | 0 | 0 | 2150/3630/7124/3553/5573/7351/2641/3569/3667/3952/8431 | 11 |
| BP | GO:0009306 | protein secretion | 0 | 0 | 0 | /5468/348/3630/7124/3553/5573/7351/2641/3569/3667/3952/3552/ | 14 |
| BP | GO:0035592 | establishment of protein localization to extracellular region | 0 | 0 | 0 | /5468/348/3630/7124/3553/5573/7351/2641/3569/3667/3952/3552/ | 14 |
| BP | GO:0120161 | regulation of cold-induced thermogenesis | 0 | 0 | 0 | 2778/857/26291/9370/7351/3953/948/3952/283455/153 | 10 |
| BP | GO:0010875 | positive regulation of cholesterol efflux | 0 | 0 | 0 | 5468/348/335/857/23411/9370 | 6 |
| BP | GO:0051222 | positive regulation of protein transport | 0 | 0 | 0 | 16/2150/5468/3630/7124/3553/208/5743/5573/2641/3952/3552/84 | 13 |
| BP | GO:0106106 | cold-induced thermogenesis | 0 | 0 | 0 | 2778/857/26291/9370/7351/3953/948/3952/283455/153 | 10 |
| BP | GO:0002791 | regulation of peptide secretion | 0 | 0 | 0 | 2150/3630/7124/3553/5573/7351/2641/3569/3667/3952/8431 | 11 |
| BP | GO:0009746 | response to hexose | 0 | 0 | 0 | 1636/2876/4023/7124/5743/26291/9370/7351/2641/3383/8431 | 11 |
| BP | GO:0006066 | alcohol metabolic process | 0 | 0 | 0 | 335/1071/10577/3990/6647/3953/2784/3952/949/8431/4864/3949/ | 14 |
| BP | GO:1903531 | negative regulation of secretion by cell | 0 | 0 | 0 | 7879/2150/348/3630/3553/9370/7351/7133/3667/3952 | 10 |
| BP | GO:0090087 | regulation of peptide transport | 0 | 0 | 0 | 2150/3630/7124/3553/5573/7351/2641/3569/3667/3952/8431 | 11 |
| BP | GO:0071692 | protein localization to extracellular region | 0 | 0 | 0 | /5468/348/3630/7124/3553/5573/7351/2641/3569/3667/3952/3552/ | 14 |
| BP | GO:0050728 | negative regulation of inflammatory response | 0 | 0 | 0 | 2876/5468/348/3630/335/6647/9370/7133/5465/196/3949 | 11 |
| BP | GO:0044042 | glucan metabolic process | 0 | 0 | 0 | 3630/34/208/3953/3667/5443/5167/4864 | 8 |
| BP | GO:0046209 | nitric oxide metabolic process | 0 | 0 | 0 | 6648/2717/7124/3553/857/5743/948/1727 | 8 |
| BP | GO:0097009 | energy homeostasis | 0 | 0 | 0 | 9518/348/129787/23411/3953/948/3952/153 | 8 |
| BP | GO:0045807 | positive regulation of endocytosis | 0 | 0 | 0 | 2150/348/7124/57534/3553/335/6347/6647/948/5054 | 10 |
| BP | GO:0062197 | cellular response to chemical stress | 0 | 0 | 0 | 52/6648/3066/2876/10516/857/5743/23411/6647/948/3569/4318/4: | 13 |
| BP | GO:2001057 | reactive nitrogen species metabolic process | 0 | 0 | 0 | 6648/2717/7124/3553/857/5743/948/1727 | 8 |
| BP | GO:0010745 | negative regulation of macrophage derived foam cell differentiation | 0 | 0 | 0 | 5468/1071/9370/5465/1401 | 5 |
| BP | GO:0032770 | positive regulation of monooxygenase activity | 0 | 0 | 0 | 348/3630/7124/3553/949/5447 | 6 |
| BP | GO:0032371 | regulation of sterol transport | 0 | 0 | 0 | 5468/348/335/857/1071/23411/9370/3952 | 8 |
| BP | GO:0032374 | regulation of cholesterol transport | 0 | 0 | 0 | 5468/348/335/857/1071/23411/9370/3952 | 8 |
| BP | GO:0005996 | monosaccharide metabolic process | 0 | 0 | 0 | 3630/7124/34/208/23411/9370/5465/3953/2641/2784/3667/3952 | 12 |
| BP | GO:0031348 | negative regulation of defense response | 0 | 0 | 0 | 150/2876/5468/348/3630/2671/335/6647/9370/7133/5465/196/394 | 13 |
| BP | GO:0008643 | carbohydrate transport | 0 | 0 | 0 | 1636/3630/7124/3553/208/26291/9370/3667/3952/5167 | 10 |
| BP | GO:1904951 | positive regulation of establishment of protein localization | 0 | 0 | 0 | 16/2150/5468/3630/7124/3553/208/5743/5573/2641/3952/3552/84 | 13 |
| BP | GO:0150076 | neuroinflammatory response | 0 | 0 | 0 | 3688/7124/3553/5743/7133/3569/4318/3949 | 8 |
| BP | GO:0031331 | positive regulation of cellular catabolic process | 0 | 0 | 0 | 48/3630/7124/3553/208/5770/23411/79068/26291/7133/5465/356 | 14 |
| BP | GO:0016051 | carbohydrate biosynthetic process | 0 | 0 | 0 | 3630/34/208/23411/9370/5465/3953/2641/3667/3952/5167 | 11 |
| BP | GO:0045471 | response to ethanol | 0 | 0 | 0 | 6648/3066/7124/6647/9370/4968/5465/3952/8431 | 9 |
| BP | GO:0010874 | regulation of cholesterol efflux | 0 | 0 | 0 | 5468/348/335/857/1071/23411/9370 | 7 |
| BP | GO:0050873 | brown fat cell differentiation | 0 | 0 | 0 | 3630/5743/23411/79068/9370/3952/153 | 7 |
| BP | GO:0050796 | regulation of insulin secretion | 0 | 0 | 0 | 2150/7124/3553/5573/7351/2641/3569/3667/3952/8431 | 10 |
| BP | GO:1990845 | adaptive thermogenesis | 0 | 0 | 0 | 2778/857/26291/9370/7351/3953/948/3952/283455/153 | 10 |
| BP | GO:0070542 | response to fatty acid | 0 | 0 | 0 | 4023/5743/9370/7351/948/3667/3949 | 7 |
| BP | GO:0008286 | insulin receptor signaling pathway | 0 | 0 | 0 | 3630/3553/208/3416/5770/23411/3667/3952/5167 | 9 |
| BP | GO:0034375 | high-density lipoprotein particle remodeling | 0 | 0 | 0 | 348/335/1071/3990/949 | 5 |
| BP | GO:0009266 | response to temperature stimulus | 0 | 0 | 0 | 3162/3066/34/5743/23411/6647/7351/153/3552/5346 | 10 |
| BP | GO:0034368 | protein-lipid complex remodeling | 0 | 0 | 0 | 348/4023/335/1071/3990/949 | 6 |
| BP | GO:0034369 | plasma lipoprotein particle remodeling | 0 | 0 | 0 | 348/4023/335/1071/3990/949 | 6 |
| BP | GO:0006112 | energy reserve metabolic process | 0 | 0 | 0 | 3630/34/208/3953/3667/3952/5443/5167 | 8 |
| BP | GO:0097237 | cellular response to toxic substance | 0 | 0 | 0 | 6648/2876/348/10516/7124/2671/5743/6647/948 | 9 |
| BP | GO:0045428 | regulation of nitric oxide biosynthetic process | 0 | 0 | 0 | 6648/2717/7124/3553/857/5743/948 | 7 |
| BP | GO:0071222 | cellular response to lipopolysaccharide | 0 | 0 | 0 | 7124/3553/2671/6347/7133/948/3569/5054/4318/949/3552 | 11 |
| BP | GO:0006094 | gluconeogenesis | 0 | 0 | 0 | 3630/34/23411/9370/5465/3953/2641/3952 | 8 |
| BP | GO:0034383 | low-density lipoprotein particle clearance | 0 | 0 | 0 | 3162/26291/9370/948/949/3949 | 6 |
| BP | GO:0031649 | heat generation | 0 | 0 | 0 | 7124/3553/5743/153/3552 | 5 |
| BP | GO:0080164 | regulation of nitric oxide metabolic process | 0 | 0 | 0 | 6648/2717/7124/3553/857/5743/948 | 7 |
| BP | GO:0034367 | protein-containing complex remodeling | 0 | 0 | 0 | 348/4023/335/1071/3990/949 | 6 |
| BP | GO:0019319 | hexose biosynthetic process | 0 | 0 | 0 | 3630/34/23411/9370/5465/3953/2641/3952 | 8 |
| BP | GO:0043534 | blood vessel endothelial cell migration | 0 | 0 | 0 | 3162/2876/5468/348/3688/7124/335/5743/23411/949 | 10 |
| BP | GO:0030299 | intestinal cholesterol absorption | 0 | 0 | 0 | 335/948/3952/4864/3949 | 5 |
| BP | GO:0032757 | positive regulation of interleukin-8 production | 0 | 0 | 0 | 2150/7124/3553/9370/3569/3952/5054 | 7 |
| BP | GO:0030072 | peptide hormone secretion | 0 | 0 | 0 | 2150/3630/7124/3553/5573/7351/2641/3569/3667/3952/8431 | 11 |
| BP | GO:0010885 | regulation of cholesterol storage | 0 | 0 | 0 | 5468/4023/5465/948/949 | 5 |
| BP | GO:2001242 | regulation of intrinsic apoptotic signaling pathway | 0 | 0 | 0 | 6648/2876/3630/857/5743/5770/23411/6647/4318/7466 | 10 |
| BP | GO:0071901 | negative regulation of protein serine/threonine kinase activity | 0 | 0 | 0 | 5577/5468/348/3553/5770/23411/5573/9370 | 8 |
| BP | GO:0002790 | peptide secretion | 0 | 0 | 0 | 2150/3630/7124/3553/5573/7351/2641/3569/3667/3952/8431 | 11 |
| BP | GO:1905953 | negative regulation of lipid localization | 0 | 0 | 0 | 5468/7124/208/5465/1401/3569/3952 | 7 |
| BP | GO:0032755 | positive regulation of interleukin-6 production | 0 | 0 | 0 | 2150/4023/7124/3553/948/3569/3952/3552 | 8 |
| BP | GO:0005976 | polysaccharide metabolic process | 0 | 0 | 0 | 3630/34/208/3953/3667/5443/5167/4864 | 8 |
| BP | GO:0046364 | monosaccharide biosynthetic process | 0 | 0 | 0 | 3630/34/23411/9370/5465/3953/2641/3952 | 8 |
| BP | GO:0043691 | reverse cholesterol transport | 0 | 0 | 0 | 348/335/1071/3990/949 | 5 |
| BP | GO:0032677 | regulation of interleukin-8 production | 0 | 0 | 0 | 2150/7124/3553/9370/1401/3569/3952/5054 | 8 |
| BP | GO:0070372 | regulation of ERK1 and ERK2 cascade | 0 | 0 | 0 | 2150/348/7124/3553/5770/6347/26291/9370/948/2641/3383/3552 | 12 |
| BP | GO:0032637 | interleukin-8 production | 0 | 0 | 0 | 2150/7124/3553/9370/1401/3569/3952/5054 | 8 |
| BP | GO:0097305 | response to alcohol | 0 | 0 | 0 | 1636/6648/3066/7124/6647/9370/4968/5465/196/3952/8431 | 11 |
| BP | GO:0046626 | regulation of insulin receptor signaling pathway | 0 | 0 | 0 | 3630/3553/5770/23411/3667/3952/5167 | 7 |
| BP | GO:0050892 | intestinal absorption | 0 | 0 | 0 | 335/948/3952/949/4864/3949 | 6 |
| BP | GO:0034599 | cellular response to oxidative stress | 0 | 0 | 0 | 3162/6648/3066/2876/10516/23411/6647/948/3569/4318/4363 | 11 |
| BP | GO:0030073 | insulin secretion | 0 | 0 | 0 | 2150/7124/3553/5573/7351/2641/3569/3667/3952/8431 | 10 |
| BP | GO:0010829 | negative regulation of glucose transmembrane transport | 0 | 0 | 0 | 1636/7124/3553/3952/5167 | 5 |
| BP | GO:0097193 | intrinsic apoptotic signaling pathway | 0 | 0 | 0 | 6648/2876/3630/7124/857/5743/5770/23411/6647/7133/4318/7466 | 12 |
| BP | GO:0002697 | regulation of immune effector process | 0 | 0 | 0 | 150/3630/7124/3553/2671/335/23411/7133/196/948/3383/3569/39: | 13 |
| BP | GO:0002526 | acute inflammatory response | 0 | 0 | 0 | 3630/7124/3553/5265/5743/1401/3569/3552 | 8 |
| BP | GO:0001935 | endothelial cell proliferation | 0 | 0 | 0 | 3162/5468/348/7124/335/857/6347/23411/3952/949 | 10 |
| BP | GO:0044403 | biological process involved in symbiotic interaction | 0 | 0 | 0 | 7879/2150/2876/348/3688/857/3416/5465/3383/949/4864/3949 | 12 |
| BP | GO:0007623 | circadian rhythm | 0 | 0 | 0 | 3066/5468/7124/23411/9370/5465/196/3952/153/8431 | 10 |
| BP | GO:0014823 | response to activity | 0 | 0 | 0 | 6648/7124/26291/9370/2641/3569/3952 | 7 |

| | | | | | | | |
|---|---|---|---|---|---|---|---|
| BP | GO:0032722 | positive regulation of chemokine production | 0 | 0 | 0 | 3162/2150/4023/7124/3553/9370/3569 | 7 |
| BP | GO:0050766 | positive regulation of phagocytosis | 0 | 0 | 0 | 2150/7124/3553/335/6347/6647/948 | 7 |
| BP | GO:0015833 | peptide transport | 0 | 0 | 0 | 2150/3630/7124/3553/5573/7351/2641/3569/3667/3952/8431 | 11 |
| BP | GO:0044241 | lipid digestion | 0 | 0 | 0 | 335/948/3952/4864/3949 | 5 |
| BP | GO:0007584 | response to nutrient | 0 | 0 | 0 | 6648/2876/5468/4023/5743/9370/4968/3952/3552 | 9 |
| BP | GO:0090207 | regulation of triglyceride metabolic process | 0 | 0 | 0 | 348/23411/26291/2784/949/3949 | 6 |
| BP | GO:1903829 | positive regulation of protein localization | 0 | 0 | 0 | '2150/5468/3688/3630/7124/3553/208/5743/5573/2641/3952/3552 | 14 |
| BP | GO:0010907 | positive regulation of glucose metabolic process | 0 | 0 | 0 | 3630/208/23411/5465/2641/3667 | 6 |
| BP | GO:0051090 | regulation of DNA-binding transcription factor activity | 0 | 0 | 0 | 62/3066/5468/3630/7124/3553/857/23411/948/3569/5167/7466/84 | 13 |
| BP | GO:2001243 | negative regulation of intrinsic apoptotic signaling pathway | 0 | 0 | 0 | 6648/2876/3630/5743/5770/23411/4318/7466 | 8 |
| BP | GO:0070371 | ERK1 and ERK2 cascade | 0 | 0 | 0 | 2150/348/7124/3553/5770/6347/26291/9370/948/2641/3383/3552 | 12 |
| BP | GO:0005977 | glycogen metabolic process | 0 | 0 | 0 | 3630/34/208/3953/3667/5443/5167 | 7 |
| BP | GO:0032890 | regulation of organic acid transport | 0 | 0 | 0 | 3688/7124/3553/208/5465/3952/3552 | 7 |
| BP | GO:0046628 | positive regulation of insulin receptor signaling pathway | 0 | 0 | 0 | 3630/5770/23411/3667/3952 | 5 |
| BP | GO:0051353 | positive regulation of oxidoreductase activity | 0 | 0 | 0 | 348/3630/7124/3553/949/5447 | 6 |
| BP | GO:0051402 | neuron apoptotic process | 0 | 0 | 0 | 6648/2876/348/7124/10280/208/6347/23411/6647/7351/7466 | 11 |
| BP | GO:0090335 | regulation of brown fat cell differentiation | 0 | 0 | 0 | 3630/5743/23411/79068/3952 | 5 |
| BP | GO:0002718 | regulation of cytokine production involved in immune response | 0 | 0 | 0 | 2150/7124/3553/335/23411/7133/948/3569 | 8 |
| BP | GO:0002367 | cytokine production involved in immune response | 0 | 0 | 0 | 2150/7124/3553/335/23411/7133/948/3569 | 8 |
| BP | GO:0042886 | amide transport | 0 | 0 | 0 | 2150/3688/3630/7124/3553/5573/7351/2641/3569/3667/3952/8431 | 12 |
| BP | GO:1900078 | positive regulation of cellular response to insulin stimulus | 0 | 0 | 0 | 3630/5770/23411/3667/3952 | 5 |
| BP | GO:0071827 | plasma lipoprotein particle organization | 0 | 0 | 0 | 348/4023/335/1071/3990/949 | 6 |
| BP | GO:0042180 | cellular ketone metabolic process | 0 | 0 | 0 | 5468/3630/3553/208/857/5743/23411/9370/5465/3667 | 10 |
| BP | GO:0001660 | fever generation | 0 | 0 | 0 | 7124/3553/5743/3552 | 4 |
| BP | GO:0050996 | positive regulation of lipid catabolic process | 0 | 0 | 0 | 3553/208/26291/5465/3667 | 5 |
| BP | GO:0048511 | rhythmic process | 0 | 0 | 0 | 3066/5468/7124/9510/23411/9370/5465/196/3952/153/8431 | 11 |
| BP | GO:0071825 | protein-lipid complex organization | 0 | 0 | 0 | 348/4023/335/1071/3990/949 | 6 |
| BP | GO:0051224 | negative regulation of protein transport | 0 | 0 | 0 | 2150/348/3630/3553/9370/7351/948/3667 | 8 |
| BP | GO:0000303 | response to superoxide | 0 | 0 | 0 | 6648/10516/6647/7351/948 | 5 |
| BP | GO:0090594 | inflammatory response to wounding | 0 | 0 | 0 | 3162/5468/7124/3569/3552 | 5 |
| BP | GO:0006469 | negative regulation of protein kinase activity | 0 | 0 | 0 | 5577/5468/348/3553/857/5770/23411/5573/9370 | 9 |
| BP | GO:0000305 | response to oxygen radical | 0 | 0 | 0 | 6648/10516/6647/7351/948 | 5 |
| BP | GO:0061041 | regulation of wound healing | 0 | 0 | 0 | 5327/2150/348/3688/7124/857/948/5054 | 8 |
| BP | GO:1904950 | negative regulation of establishment of protein localization | 0 | 0 | 0 | 2150/348/3630/3553/9370/7351/948/3667 | 8 |
| BP | GO:0003012 | muscle system process | 0 | 0 | 0 | 162/3066/5468/7124/3553/857/5318/5743/6647/7133/5465/3952/48 | 13 |
| BP | GO:1904705 | regulation of vascular associated smooth muscle cell proliferation | 0 | 0 | 0 | 6648/1786/5468/7124/9510/9370/4318 | 7 |
| BP | GO:0060761 | negative regulation of response to cytokine stimulus | 0 | 0 | 0 | 166/2150/5468/335/857/9370/3569 | 7 |
| BP | GO:0090322 | regulation of superoxide metabolic process | 0 | 0 | 0 | 2150/10516/6647/948/1401 | 5 |
| BP | GO:0043467 | regulation of generation of precursor metabolites and energy | 0 | 0 | 0 | 3630/7124/208/3416/5465/3667/5443/5167 | 8 |
| BP | GO:1990874 | vascular associated smooth muscle cell proliferation | 0 | 0 | 0 | 6648/1786/5468/7124/9510/9370/4318 | 7 |
| BP | GO:0046486 | glycerolipid metabolic process | 0 | 0 | 0 | 2876/348/4023/335/857/1071/3990/23411/26291/2784/949/3949 | 12 |
| BP | GO:0009749 | response to glucose | 0 | 0 | 0 | 1636/2876/4023/26291/9370/7351/2641/3383/8431 | 9 |
| BP | GO:0033673 | negative regulation of kinase activity | 0 | 0 | 0 | 5577/5468/348/3553/857/5770/23411/5573/9370 | 9 |
| BP | GO:0019058 | viral life cycle | 0 | 0 | 0 | 7879/348/3688/7124/857/3416/6347/3383/949/4864/3949 | 11 |
| BP | GO:0048661 | positive regulation of smooth muscle cell proliferation | 0 | 0 | 0 | 3162/1786/7124/9510/5743/3569/4318 | 7 |
| BP | GO:0034113 | heterotypic cell-cell adhesion | 0 | 0 | 0 | 3688/7124/3553/335/5318/9370 | 6 |
| BP | GO:0045599 | negative regulation of fat cell differentiation | 0 | 0 | 0 | 6648/7124/23411/9370/3569/5167 | 6 |
| BP | GO:0034763 | negative regulation of transmembrane transport | 0 | 0 | 0 | 1636/7124/3553/208/857/3952/4318/5167 | 8 |
| BP | GO:0009636 | response to toxic substance | 0 | 0 | 0 | 6648/2876/348/10516/7124/2671/5743/6647/196/948 | 10 |
| BP | GO:0098869 | cellular oxidant detoxification | 0 | 0 | 0 | 6648/2876/348/10516/5743/6647/948 | 7 |
| BP | GO:2001237 | negative regulation of extrinsic apoptotic signaling pathway | 0 | 0 | 0 | 3162/2876/7124/3553/3383/5054/3552 | 7 |
| BP | GO:0042542 | response to hydrogen peroxide | 0 | 0 | 0 | 3162/6648/3066/2876/23411/6647/3569 | 7 |
| BP | GO:0071715 | icosanoid transport | 0 | 0 | 0 | 1636/3553/5743/3952/3552/4363 | 6 |
| BP | GO:0031100 | animal organ regeneration | 0 | 0 | 0 | 1636/7124/2671/7351/3569/8431 | 6 |
| BP | GO:1904036 | negative regulation of epithelial cell apoptotic process | 0 | 0 | 0 | 26291/5465/3383/5054/7466/4864 | 6 |
| BP | GO:2000479 | regulation of cAMP-dependent protein kinase activity | 0 | 0 | 0 | 5577/23411/5573/9370 | 4 |
| BP | GO:0015914 | phospholipid transport | 0 | 0 | 0 | 348/335/1071/10577/949/4363/3949 | 7 |
| BP | GO:0015980 | energy derivation by oxidation of organic compounds | 0 | 0 | 0 | 6648/3630/7124/34/208/3416/3953/3667/3952/5443/5167 | 11 |
| BP | GO:0032642 | regulation of chemokine production | 0 | 0 | 0 | 3162/2150/4023/7124/3553/9370/3569 | 7 |
| BP | GO:0050729 | positive regulation of inflammatory response | 0 | 0 | 0 | 4023/7124/3553/5743/3569/5054/4363/3949 | 8 |
| BP | GO:0032602 | chemokine production | 0 | 0 | 0 | 3162/2150/4023/7124/3553/9370/3569 | 7 |
| BP | GO:0034614 | cellular response to reactive oxygen species | 0 | 0 | 0 | 6648/3066/10516/23411/6647/948/3569/4318 | 8 |
| BP | GO:0002532 | production of molecular mediator involved in inflammatory response | 0 | 0 | 0 | 3630/7124/5465/948/3569/3952/5054 | 7 |
| BP | GO:0007631 | feeding behavior | 0 | 0 | 0 | 1636/9518/3630/129787/3953/2641/3952 | 7 |
| BP | GO:0045732 | positive regulation of protein catabolic process | 0 | 0 | 0 | 7879/348/7124/3553/857/3416/7133/7466/3949 | 9 |
| BP | GO:0046888 | negative regulation of hormone secretion | 0 | 0 | 0 | 2150/3553/9370/7351/3667/3952 | 6 |
| BP | GO:0007159 | leukocyte cell-cell adhesion | 0 | 0 | 0 | 7356/3688/7124/3553/857/6347/5573/5465/3383/3569/3952/3552 | 12 |
| BP | GO:0051701 | biological process involved in interaction with host | 0 | 0 | 0 | 7879/3688/857/3416/5465/3383/949/4864/3949 | 9 |
| BP | GO:0046325 | negative regulation of glucose import | 0 | 0 | 0 | 1636/7124/3952/5167 | 4 |
| BP | GO:0051044 | positive regulation of membrane protein ectodomain proteolysis | 0 | 0 | 0 | 348/7124/3553/7133 | 4 |
| BP | GO:0046326 | positive regulation of glucose import | 0 | 0 | 0 | 3630/208/26291/9370/3667 | 5 |
| BP | GO:0030193 | regulation of blood coagulation | 0 | 0 | 0 | 5327/2150/348/857/948/5054 | 6 |
| BP | GO:2000379 | positive regulation of reactive oxygen species metabolic process | 0 | 0 | 0 | 6648/2150/6647/948/1401/3952 | 6 |
| BP | GO:2001236 | regulation of extrinsic apoptotic signaling pathway | 0 | 0 | 0 | 3162/2876/7124/3553/857/3383/5054/3552 | 8 |
| BP | GO:0045765 | regulation of angiogenesis | 0 | 0 | 0 | 3162/5468/3688/7124/3553/9510/23411/3569/3952/5054/3552 | 11 |
| BP | GO:0070661 | leukocyte proliferation | 0 | 0 | 0 | 1636/7356/2150/3553/5573/7133/196/1401/3569/3952/3552 | 11 |
| BP | GO:0045429 | positive regulation of nitric oxide biosynthetic process | 0 | 0 | 0 | 6648/7124/3553/5743/948 | 5 |
| BP | GO:0070873 | regulation of glycogen metabolic process | 0 | 0 | 0 | 3630/208/3667/5443/5167 | 5 |
| BP | GO:0010469 | regulation of signaling receptor activity | 0 | 0 | 0 | 3066/5468/3688/7124/6347/5465/5054 | 7 |
| BP | GO:0034764 | positive regulation of transmembrane transport | 0 | 0 | 0 | 3688/3630/208/857/6347/26291/9370/3667/481 | 9 |
| BP | GO:1900046 | regulation of hemostasis | 0 | 0 | 0 | 5327/2150/348/857/948/5054 | 6 |
| BP | GO:0070374 | positive regulation of ERK1 and ERK2 cascade | 0 | 0 | 0 | 2150/348/7124/6347/26291/948/2641/3383/3552 | 9 |
| BP | GO:0032310 | prostaglandin secretion | 0 | 0 | 0 | 3553/5743/3952/3552 | 4 |
| BP | GO:1901342 | regulation of vasculature development | 0 | 0 | 0 | 3162/5468/3688/7124/3553/9510/23411/3569/3952/5054/3552 | 11 |
| BP | GO:0016032 | viral process | 0 | 0 | 0 | 7879/348/3688/7124/857/3416/6347/5465/3383/949/4864/3949 | 12 |
| BP | GO:0043542 | endothelial cell migration | 0 | 0 | 0 | 3162/2876/5468/348/3688/7124/335/5743/23411/949 | 10 |
| BP | GO:0097242 | amyloid-beta clearance | 0 | 0 | 0 | 348/7124/3416/948/3949 | 5 |
| BP | GO:1904407 | positive regulation of nitric oxide metabolic process | 0 | 0 | 0 | 6648/7124/3553/5743/948 | 5 |
| BP | GO:0006801 | superoxide metabolic process | 0 | 0 | 0 | 6648/2150/10516/6647/948/1401 | 6 |
| BP | GO:0031400 | negative regulation of protein modification process | 0 | 0 | 0 | 5577/3066/5468/348/7124/3553/857/5770/23411/5573/9370/5167 | 12 |
| BP | GO:1903532 | positive regulation of secretion by cell | 0 | 0 | 0 | 7879/2150/5468/3630/3553/5573/2641/3952/3552/8431 | 10 |
| BP | GO:0007596 | blood coagulation | 0 | 0 | 0 | 2778/5327/2150/348/857/5265/948/3569/5054 | 9 |
| BP | GO:0062014 | negative regulation of small molecule metabolic process | 0 | 0 | 0 | 348/3630/23411/6647/9370/5465/3953 | 7 |
| BP | GO:0032732 | positive regulation of interleukin-1 production | 0 | 0 | 0 | 3066/2150/4023/7124/948/3569 | 6 |
| BP | GO:0050818 | regulation of coagulation | 0 | 0 | 0 | 5327/2150/348/857/948/5054 | 6 |
| BP | GO:0043491 | phosphatidylinositol 3-kinase/protein kinase B signal transduction | 0 | 0 | 0 | 9518/2150/2876/3688/3630/7124/23411/5465/3667/3952 | 10 |
| BP | GO:0051348 | negative regulation of transferase activity | 0 | 0 | 0 | 5577/5468/348/3553/857/5770/23411/5573/9370 | 9 |
| BP | GO:0034384 | high-density lipoprotein particle clearance | 0 | 0 | 0 | 348/335/949/3949 | 4 |
| BP | GO:0042176 | regulation of protein catabolic process | 0 | 0 | 0 | 7879/2876/348/3630/7124/3553/857/3416/7133/7466/3949 | 11 |
| BP | GO:0048732 | gland development | 0 | 0 | 0 | 6648/2876/7124/34/2671/208/335/857/6647/7351/3569/4864 | 12 |
| BP | GO:0031952 | regulation of protein autophosphorylation | 0 | 0 | 0 | 1636/3630/857/9370/5167 | 5 |
| BP | GO:0090181 | regulation of cholesterol metabolic process | 0 | 0 | 0 | 348/335/6647/2784/3949 | 5 |
| BP | GO:0050709 | negative regulation of protein secretion | 0 | 0 | 0 | 2150/348/3630/3553/7351/3667 | 6 |
| BP | GO:0050731 | positive regulation of peptidyl-tyrosine phosphorylation | 0 | 0 | 0 | 1636/3066/7124/5770/9370/948/3569/3952 | 8 |
| BP | GO:1902074 | response to salt | 0 | 0 | 0 | 6648/3066/7124/3553/857/5743/9370/948/3952/5167/3552 | 11 |
| BP | GO:0050817 | coagulation | 0 | 0 | 0 | 2778/5327/2150/348/857/5265/948/3569/5054 | 9 |
| BP | GO:0097191 | extrinsic apoptotic signaling pathway | 0 | 0 | 0 | 3162/2876/7124/3553/857/7133/3383/5054/3552 | 9 |
| BP | GO:1990748 | cellular detoxification | 0 | 0 | 0 | 6648/2876/348/10516/5743/6647/948 | 7 |
| BP | GO:0051897 | egulation of phosphatidylinositol 3-kinase/protein kinase B signal tra | 0 | 0 | 0 | 9518/2150/2876/3688/3630/7124/23411/3952 | 8 |

| | | | | | | | |
|---|---|---|---|---|---|---|---|
| BP | GO:0006635 | fatty acid beta-oxidation | 0 | 0 | 0 | 34/208/9370/5465/3667/3952 | 6 |
| BP | GO:0007599 | hemostasis | 0 | 0 | 0 | 2778/5327/2150/348/857/5265/948/3569/5054 | 9 |
| BP | GO:0050730 | regulation of peptidyl-tyrosine phosphorylation | 0 | 0 | 0 | 1636/3066/7124/857/5770/9370/948/3569/3952 | 9 |
| BP | GO:0010631 | epithelial cell migration | 0 | 0 | 0 | 3162/2876/5468/348/3688/7124/335/5743/23411/4318/949 | 11 |
| BP | GO:0043523 | regulation of neuron apoptotic process | 0 | 0 | 0 | 6648/348/7124/10280/6347/23411/6647/7351/7466 | 9 |
| BP | GO:1903034 | regulation of response to wounding | 0 | 0 | 0 | 5327/2150/348/3688/7124/857/948/5054 | 8 |
| BP | GO:0090132 | epithelium migration | 0 | 0 | 0 | 3162/2876/5468/348/3688/7124/335/5743/23411/4318/949 | 11 |
| BP | GO:0045936 | negative regulation of phosphate metabolic process | 0 | 0 | 0 | 5577/5468/348/7124/3553/857/5770/23411/5573/9370/5167 | 11 |
| BP | GO:0010563 | negative regulation of phosphorus metabolic process | 0 | 0 | 0 | 5577/5468/348/7124/3553/857/5770/23411/5573/9370/5167 | 11 |
| BP | GO:0001933 | negative regulation of protein phosphorylation | 0 | 0 | 0 | 5577/5468/348/3553/857/5770/23411/5573/9370/5167 | 10 |
| BP | GO:0010828 | positive regulation of glucose transmembrane transport | 0 | 0 | 0 | 3630/208/26291/9370/3667 | 5 |
| BP | GO:0090130 | tissue migration | 0 | 0 | 0 | 3162/2876/5468/348/3688/7124/335/5743/23411/4318/949 | 11 |
| BP | GO:0032098 | regulation of appetite | 0 | 0 | 0 | 6581/5465/3952/5443 | 4 |
| BP | GO:0032635 | interleukin-6 production | 0 | 0 | 0 | 2150/4023/7124/3553/948/3569/3952/3552 | 8 |
| BP | GO:0032675 | regulation of interleukin-6 production | 0 | 0 | 0 | 2150/4023/7124/3553/948/3569/3952/3552 | 8 |
| BP | GO:0051047 | positive regulation of secretion | 0 | 0 | 0 | 7879/2150/5468/3630/3553/5573/2641/3952/3552/8431 | 10 |
| BP | GO:0045913 | positive regulation of carbohydrate metabolic process | 0 | 0 | 0 | 3630/208/23411/5465/2641/3667 | 6 |
| BP | GO:0050810 | regulation of steroid biosynthetic process | 0 | 0 | 0 | 348/7124/23411/6647/3952/3552 | 6 |
| BP | GO:0032944 | regulation of mononuclear cell proliferation | 0 | 0 | 0 | 7356/3553/5573/7133/196/1401/3569/3952/3552 | 9 |
| BP | GO:0001936 | regulation of endothelial cell proliferation | 0 | 0 | 0 | 3162/5468/348/7124/857/6347/23411/3952 | 8 |
| BP | GO:0005978 | glycogen biosynthetic process | 0 | 0 | 0 | 3630/34/208/3667/5167 | 5 |
| BP | GO:0009250 | glucan biosynthetic process | 0 | 0 | 0 | 3630/34/208/3667/5167 | 5 |
| BP | GO:0032309 | icosanoid secretion | 0 | 0 | 0 | 1636/3553/5743/3952/3552 | 5 |
| BP | GO:0032881 | regulation of polysaccharide metabolic process | 0 | 0 | 0 | 3630/208/3667/5443/5167 | 5 |
| BP | GO:1902175 | gulation of oxidative stress-induced intrinsic apoptotic signaling pathv | 0 | 0 | 0 | 6648/2876/3630/23411/6647 | 5 |
| BP | GO:0032943 | mononuclear cell proliferation | 0 | 0 | 0 | 1636/7356/3553/5573/7133/196/1401/3569/3952/3552 | 10 |
| BP | GO:0060759 | regulation of response to cytokine stimulus | 0 | 0 | 0 | 166/2150/5468/335/857/5770/9370/3569 | 8 |
| BP | GO:0140353 | lipid export from cell | 0 | 0 | 0 | 3553/5743/3952/5443/3552 | 5 |
| BP | GO:0006694 | steroid biosynthetic process | 0 | 0 | 0 | 348/7124/335/23411/6647/3952/3552/1727 | 8 |
| BP | GO:0048771 | tissue remodeling | 0 | 0 | 0 | 1636/7879/5468/857/3953/3569/3952/3552 | 8 |
| BP | GO:0032612 | interleukin-1 production | 0 | 0 | 0 | 3066/2150/4023/7124/335/948/3569 | 7 |
| BP | GO:0032652 | regulation of interleukin-1 production | 0 | 0 | 0 | 3066/2150/4023/7124/335/948/3569 | 7 |
| BP | GO:0051896 | ation of phosphatidylinositol 3-kinase/protein kinase B signal transdu | 0 | 0 | 0 | 9518/2150/2876/3688/3630/7124/23411/5465/3952 | 9 |
| BP | GO:0002703 | regulation of leukocyte mediated immunity | 0 | 0 | 0 | 2150/7124/3553/2671/7133/196/3383/3569/3952 | 9 |
| BP | GO:0007565 | female pregnancy | 0 | 0 | 0 | 1636/7356/3553/5743/6647/7351/3952/4318 | 8 |
| BP | GO:0032640 | tumor necrosis factor production | 0 | 0 | 0 | 3066/4023/9370/948/3569/3952/5443/3552 | 8 |
| BP | GO:0032680 | regulation of tumor necrosis factor production | 0 | 0 | 0 | 3066/4023/9370/948/3569/3952/5443/3552 | 8 |
| BP | GO:0050900 | leukocyte migration | 0 | 0 | 0 | 3162/2150/3688/7124/3553/6347/3383/3569/3952/5054/3552 | 11 |
| BP | GO:0045776 | negative regulation of blood pressure | 0 | 0 | 0 | 6648/7124/9370/5465/153 | 5 |
| BP | GO:0050673 | epithelial cell proliferation | 0 | 0 | 0 | 3162/2876/5468/348/7124/335/857/6347/23411/3569/3952/949 | 12 |
| BP | GO:0044000 | movement in host | 0 | 0 | 0 | 7879/3688/857/3416/3383/949/4864/3949 | 8 |
| BP | GO:0033273 | response to vitamin | 0 | 0 | 0 | 6648/2876/5743/4968/3952/3552 | 6 |
| BP | GO:0002673 | regulation of acute inflammatory response | 0 | 0 | 0 | 3630/7124/3553/5743/3569 | 5 |
| BP | GO:0002700 | regulation of production of molecular mediator of immune response | 0 | 0 | 0 | 2150/7124/3553/335/23411/7133/948/3569 | 8 |
| BP | GO:0071706 | tumor necrosis factor superfamily cytokine production | 0 | 0 | 0 | 3066/4023/9370/948/3569/3952/5443/3552 | 8 |
| BP | GO:1903555 | regulation of tumor necrosis factor superfamily cytokine production | 0 | 0 | 0 | 3066/4023/9370/948/3569/3952/5443/3552 | 8 |
| BP | GO:0019430 | removal of superoxide radicals | 0 | 0 | 0 | 6648/10516/6647/948 | 4 |
| BP | GO:0042326 | negative regulation of phosphorylation | 0 | 0 | 0 | 5577/5468/348/3553/857/5770/23411/5573/9370/5167 | 10 |
| BP | GO:0048545 | response to steroid hormone | 0 | 0 | 0 | 1636/7356/7124/857/5743/23411/9370/5465/3569/4864 | 10 |
| BP | GO:0001960 | negative regulation of cytokine-mediated signaling pathway | 0 | 0 | 0 | 2150/5468/335/857/9370/3569 | 6 |
| BP | GO:0022407 | regulation of cell-cell adhesion | 0 | 0 | 0 | 7356/7124/3553/335/857/6347/5573/9370/5465/3569/3952/3552 | 12 |
| BP | GO:0034612 | response to tumor necrosis factor | 0 | 0 | 0 | 2150/7124/2671/335/5743/6347/23411/9370/7133 | 9 |
| BP | GO:0035094 | response to nicotine | 0 | 0 | 0 | 3162/3066/2876/7124/5465 | 5 |
| BP | GO:0043407 | negative regulation of MAP kinase activity | 0 | 0 | 0 | 5468/348/3553/5770/9370 | 5 |
| BP | GO:0034114 | regulation of heterotypic cell-cell adhesion | 0 | 0 | 0 | 7124/3553/335/9370 | 4 |
| BP | GO:0051043 | regulation of membrane protein ectodomain proteolysis | 0 | 0 | 0 | 348/7124/3553/7133 | 4 |
| BP | GO:0051347 | positive regulation of transferase activity | 0 | 0 | 0 | 1636/348/3630/7124/3553/335/5770/23411/9370/3667/3952 | 11 |
| BP | GO:0071326 | cellular response to monosaccharide stimulus | 0 | 0 | 0 | 1636/2876/26291/7351/2641/3383/3952 | 7 |
| BP | GO:0050886 | endocrine process | 0 | 0 | 0 | 1636/2150/5468/3553/3952/5443 | 6 |
| BP | GO:0042063 | gliogenesis | 0 | 0 | 0 | 3066/7124/3553/208/6347/6647/7133/3953/3569/3949 | 10 |
| BP | GO:1904645 | response to amyloid-beta | 0 | 0 | 0 | 7124/948/3383/4318/4363 | 5 |
| BP | GO:0001889 | liver development | 0 | 0 | 0 | 6648/7124/34/2671/7351/3569/4864 | 7 |
| BP | GO:0048143 | astrocyte activation | 0 | 0 | 0 | 7124/3553/3569/3949 | 4 |
| BP | GO:0071450 | cellular response to oxygen radical | 0 | 0 | 0 | 6648/10516/6647/948 | 4 |
| BP | GO:0071451 | cellular response to superoxide | 0 | 0 | 0 | 6648/10516/6647/948 | 4 |
| BP | GO:0070663 | regulation of leukocyte proliferation | 0 | 0 | 0 | 7356/3553/5573/7133/196/1401/3569/3952/3552 | 9 |
| BP | GO:0050714 | positive regulation of protein secretion | 0 | 0 | 0 | 2150/5468/3630/5573/2641/3552/8431 | 7 |
| BP | GO:0061008 | hepaticobiliary system development | 0 | 0 | 0 | 6648/7124/34/2671/7351/3569/4864 | 7 |
| BP | GO:0044703 | multi-organism reproductive process | 0 | 0 | 0 | 1636/7356/3553/5743/6647/7351/3952/4318 | 8 |
| BP | GO:0045912 | negative regulation of carbohydrate metabolic process | 0 | 0 | 0 | 3630/9370/5465/3953/5167 | 5 |
| BP | GO:0018108 | peptidyl-tyrosine phosphorylation | 0 | 0 | 0 | 1636/3066/7124/857/5770/9370/948/3569/3952 | 9 |
| BP | GO:0060612 | adipose tissue development | 0 | 0 | 0 | 5468/4023/23411/79068/3952 | 5 |
| BP | GO:0045940 | positive regulation of steroid metabolic process | 0 | 0 | 0 | 348/7124/335/3552 | 4 |
| BP | GO:0018212 | peptidyl-tyrosine modification | 0 | 0 | 0 | 1636/3066/7124/857/5770/9370/948/3569/3952 | 9 |
| BP | GO:0071322 | cellular response to carbohydrate stimulus | 0 | 0 | 0 | 1636/2876/26291/7351/2641/3383/3952 | 7 |
| BP | GO:1903409 | reactive oxygen species biosynthetic process | 0 | 0 | 0 | 6648/3630/6647/5465/948 | 5 |
| BP | GO:0015748 | organophosphate ester transport | 0 | 0 | 0 | 348/335/1071/10577/949/4363/3949 | 7 |
| BP | GO:0002675 | positive regulation of acute inflammatory response | 0 | 0 | 0 | 7124/3553/5743/3569 | 4 |
| BP | GO:0015732 | prostaglandin transport | 0 | 0 | 0 | 3553/5743/3952/3552 | 4 |
| BP | GO:0046686 | response to cadmium ion | 0 | 0 | 0 | 3162/6648/4968/4318/4864 | 5 |
| BP | GO:0071900 | regulation of protein serine/threonine kinase activity | 0 | 0 | 0 | 5577/5468/348/7124/3553/5770/23411/5573/9370 | 9 |
| BP | GO:0044706 | multi-multicellular organism process | 0 | 0 | 0 | 1636/7356/3553/5743/6647/7351/3952/4318 | 8 |
| BP | GO:0007566 | embryo implantation | 0 | 0 | 0 | 7356/3553/5743/6647/4318 | 5 |
| BP | GO:0009408 | response to heat | 0 | 0 | 0 | 3162/3066/5743/23411/6647/3552 | 6 |
| BP | GO:0022600 | digestive system process | 0 | 0 | 0 | 335/948/3952/949/4864/3949 | 6 |
| BP | GO:0046718 | viral entry into host cell | 0 | 0 | 0 | 3688/857/3416/3383/949/4864/3949 | 7 |
| BP | GO:1903828 | negative regulation of protein localization | 0 | 0 | 0 | 2150/348/3630/3553/9370/7351/948/3667 | 8 |
| BP | GO:0050878 | regulation of body fluid levels | 0 | 0 | 0 | 2778/5327/2150/348/857/5265/948/3569/5054/7466 | 10 |
| BP | GO:0097421 | liver regeneration | 0 | 0 | 0 | 7124/2671/7351/3569 | 4 |
| BP | GO:0001678 | intracellular glucose homeostasis | 0 | 0 | 0 | 1636/2876/23411/26291/7351/2641/3383 | 7 |
| BP | GO:2000351 | regulation of endothelial cell apoptotic process | 0 | 0 | 0 | 7124/6347/26291/3383/5054 | 5 |
| BP | GO:0098754 | detoxification | 0 | 0 | 0 | 6648/2876/348/10516/5743/6647/948 | 7 |
| BP | GO:0032731 | positive regulation of interleukin-1 beta production | 0 | 0 | 0 | 2150/4023/7124/948/3569 | 5 |
| BP | GO:0033210 | leptin-mediated signaling pathway | 0 | 0 | 0 | 23411/3953/3952 | 3 |
| BP | GO:1903596 | regulation of gap junction assembly | 0 | 0 | 0 | 1636/3553/857 | 3 |
| BP | GO:1904179 | positive regulation of adipose tissue development | 0 | 0 | 0 | 5468/4023/23411 | 3 |
| BP | GO:0009062 | fatty acid catabolic process | 0 | 0 | 0 | 34/208/9370/5465/3667/3952 | 6 |
| BP | GO:0032760 | positive regulation of tumor necrosis factor production | 0 | 0 | 0 | 3066/4023/948/3569/3952/3552 | 6 |
| BP | GO:0010575 | positive regulation of vascular endothelial growth factor production | 0 | 0 | 0 | 3553/5743/3569/3552 | 4 |
| BP | GO:0019395 | fatty acid oxidation | 0 | 0 | 0 | 34/208/9370/5465/3667/3952 | 6 |
| BP | GO:0008631 | intrinsic apoptotic signaling pathway in response to oxidative stress | 0 | 0 | 0 | 6648/2876/3630/23411/6647 | 5 |
| BP | GO:0010573 | vascular endothelial growth factor production | 0 | 0 | 0 | 7124/3553/5743/3569/3552 | 5 |
| BP | GO:0033157 | regulation of intracellular protein transport | 0 | 0 | 0 | 5116/3553/208/5743/5770/9370/948/3952 | 8 |
| BP | GO:0043524 | negative regulation of neuron apoptotic process | 0 | 0 | 0 | 6648/348/6347/23411/6647/7351/7466 | 7 |
| BP | GO:0044409 | entry into host | 0 | 0 | 0 | 3688/857/3416/3383/949/4864/3949 | 7 |
| BP | GO:0001885 | endothelial cell development | 0 | 0 | 0 | 2150/2876/7124/3553/3383 | 5 |
| BP | GO:0005979 | regulation of glycogen biosynthetic process | 0 | 0 | 0 | 3630/208/3667/5167 | 4 |
| BP | GO:0010962 | regulation of glucan biosynthetic process | 0 | 0 | 0 | 3630/208/3667/5167 | 4 |
| BP | GO:1902042 | regulation of extrinsic apoptotic signaling pathway via death domain | 0 | 0 | 0 | 3162/2876/3383/5054 | 4 |

| | | | | | | | |
|---|---|---|---|---|---|---|---|
| BP | GO:2000191 | regulation of fatty acid transport | 0 | 0 | 0 | 3553/208/5465/3552 | 4 |
| BP | GO:0046503 | glycerolipid catabolic process | 0 | 0 | 0 | 4023/3990/26291/949/3949 | 5 |
| BP | GO:0014013 | regulation of gliogenesis | 0 | 0 | 0 | 3066/7124/3553/7133/3569/3949 | 6 |
| BP | GO:1903557 | itive regulation of tumor necrosis factor superfamily cytokine product | 0 | 0 | 0 | 3066/4023/948/3569/3952/3552 | 6 |
| BP | GO:0050863 | regulation of T cell activation | 0 | 0 | 0 | 7356/3553/857/6347/5573/6647/7133/3569/3952/3552 | 10 |
| BP | GO:0002685 | regulation of leukocyte migration | 0 | 0 | 0 | 3162/2150/7124/6347/3383/3569/5054/3552 | 8 |
| BP | GO:1903037 | regulation of leukocyte cell-cell adhesion | 0 | 0 | 0 | 7356/7124/3553/857/6347/5573/5465/3569/3952/3552 | 10 |
| BP | GO:0032611 | interleukin-1 beta production | 0 | 0 | 0 | 2150/4023/7124/335/948/3569 | 6 |
| BP | GO:0032651 | regulation of interleukin-1 beta production | 0 | 0 | 0 | 2150/4023/7124/335/948/3569 | 6 |
| BP | GO:0034391 | regulation of smooth muscle cell apoptotic process | 0 | 0 | 0 | 6648/1786/5468/23411 | 4 |
| BP | GO:0000271 | polysaccharide biosynthetic process | 0 | 0 | 0 | 3630/34/208/3667/5167 | 5 |
| BP | GO:0034605 | cellular response to heat | 0 | 0 | 0 | 3162/3066/5743/23411/3552 | 5 |
| BP | GO:0031953 | negative regulation of protein autophosphorylation | 0 | 0 | 0 | 857/9370/5167 | 3 |
| BP | GO:0032000 | positive regulation of fatty acid beta-oxidation | 0 | 0 | 0 | 208/5465/3667 | 3 |
| BP | GO:0045833 | negative regulation of lipid metabolic process | 0 | 0 | 0 | 348/3630/7124/3553/23411/6647 | 6 |
| BP | GO:0015909 | long-chain fatty acid transport | 0 | 0 | 0 | 1636/5468/348/208/948 | 5 |
| BP | GO:0072577 | endothelial cell apoptotic process | 0 | 0 | 0 | 7124/6347/26291/3383/5054 | 5 |
| BP | GO:0034390 | smooth muscle cell apoptotic process | 0 | 0 | 0 | 6648/1786/5468/23411 | 4 |
| BP | GO:0034440 | lipid oxidation | 0 | 0 | 0 | 34/208/9370/5465/3667/3952 | 6 |
| BP | GO:0001959 | regulation of cytokine-mediated signaling pathway | 0 | 0 | 0 | 2150/5468/335/857/5770/9370/3569 | 7 |
| BP | GO:0050670 | regulation of lymphocyte proliferation | 0 | 0 | 0 | 7356/3553/5573/7133/196/3569/3952/3552 | 8 |
| BP | GO:0071356 | cellular response to tumor necrosis factor | 0 | 0 | 0 | 2150/7124/2671/335/6347/23411/9370/7133 | 8 |
| BP | GO:0031349 | positive regulation of defense response | 0 | 0 | 0 | 2150/4023/7124/3553/857/5743/948/3569/5054/4363/3949 | 11 |
| BP | GO:1902176 | e regulation of oxidative stress-induced intrinsic apoptotic signaling p | 0 | 0 | 0 | 6648/2876/3630/23411 | 4 |
| BP | GO:0032922 | circadian regulation of gene expression | 0 | 0 | 0 | 3066/23411/5465/196/8431 | 5 |
| BP | GO:0048662 | negative regulation of smooth muscle cell proliferation | 0 | 0 | 0 | 3162/6648/5468/348/9370 | 5 |
| BP | GO:0031650 | regulation of heat generation | 0 | 0 | 0 | 7124/3553/5743 | 3 |
| BP | GO:0071377 | cellular response to glucagon stimulus | 0 | 0 | 0 | 5573/26291/2641 | 3 |
| BP | GO:0010959 | regulation of metal ion transport | 0 | 0 | 0 | 1636/3688/857/55676/5318/5743/6347/2641/7466/481 | 10 |
| BP | GO:0043409 | negative regulation of MAPK cascade | 0 | 0 | 0 | 2150/5468/348/3553/857/5770/9370 | 7 |
| BP | GO:0002021 | response to dietary excess | 0 | 0 | 0 | 9518/348/3952/153 | 4 |
| BP | GO:0060047 | heart contraction | 0 | 0 | 0 | 1636/2876/7124/857/5318/6647/153/481 | 8 |
| BP | GO:0050804 | modulation of chemical synaptic transmission | 0 | 0 | 0 | 1636/5327/5577/348/3688/3630/7124/3553/5743/6347/9370 | 11 |
| BP | GO:0099177 | regulation of trans-synaptic signaling | 0 | 0 | 0 | 1636/5327/5577/348/3688/3630/7124/3553/5743/6347/9370 | 11 |
| BP | GO:0014015 | positive regulation of gliogenesis | 0 | 0 | 0 | 3066/7124/3553/7133/3569 | 5 |
| BP | GO:1905898 | positive regulation of response to endoplasmic reticulum stress | 0 | 0 | 0 | 857/5770/23411/7466 | 4 |
| BP | GO:0090257 | regulation of muscle system process | 0 | 0 | 0 | 5468/857/5318/5743/6647/7133/5465/481 | 8 |
| BP | GO:0021782 | glial cell development | 0 | 0 | 0 | 7124/3553/208/6647/3569/3949 | 6 |
| BP | GO:0001667 | ameboidal-type cell migration | 0 | 0 | 0 | 3162/2876/5468/348/3688/7124/335/5743/23411/4318/949 | 11 |
| BP | GO:0042129 | regulation of T cell proliferation | 0 | 0 | 0 | 7356/3553/5573/7133/3569/3952/3552 | 7 |
| BP | GO:0045123 | cellular extravasation | 0 | 0 | 0 | 3688/7124/6347/3383/3952 | 5 |
| BP | GO:0050795 | regulation of behavior | 0 | 0 | 0 | 3066/348/3630/3953/153 | 5 |
| BP | GO:0042755 | eating behavior | 0 | 0 | 0 | 1636/9518/129787/3952 | 4 |
| BP | GO:0031392 | regulation of prostaglandin biosynthetic process | 0 | 0 | 0 | 3553/5743/23411 | 3 |
| BP | GO:0010001 | glial cell differentiation | 0 | 0 | 0 | 3066/7124/3553/208/6647/7133/3569/3949 | 8 |
| BP | GO:0046470 | phosphatidylcholine metabolic process | 0 | 0 | 0 | 335/1071/3990/949/3949 | 5 |
| BP | GO:0003015 | heart process | 0 | 0 | 0 | 1636/2876/7124/857/5318/6647/153/481 | 8 |
| BP | GO:0051384 | response to glucocorticoid | 0 | 0 | 0 | 1636/7356/7124/5743/9370/3569 | 6 |
| BP | GO:0001937 | negative regulation of endothelial cell proliferation | 0 | 0 | 0 | 5468/348/7124/857/6347 | 5 |
| BP | GO:0002706 | regulation of lymphocyte mediated immunity | 0 | 0 | 0 | 7124/3553/2671/7133/196/3569/3952 | 7 |
| BP | GO:0097696 | receptor signaling pathway via STAT | 0 | 0 | 0 | 3066/5468/7124/857/6347/3569/3952 | 7 |
| BP | GO:1903522 | regulation of blood circulation | 0 | 0 | 0 | 1636/7124/857/5318/5743/3952/153/481 | 8 |
| BP | GO:0032885 | regulation of polysaccharide biosynthetic process | 0 | 0 | 0 | 3630/208/3667/5167 | 4 |
| BP | GO:0033674 | positive regulation of kinase activity | 0 | 0 | 0 | 1636/3630/7124/3553/5770/23411/9370/3667/3952 | 9 |
| BP | GO:2001140 | positive regulation of phospholipid transport | 0 | 0 | 0 | 348/335/1071 | 3 |
| BP | GO:0042509 | regulation of tyrosine phosphorylation of STAT protein | 0 | 0 | 0 | 3066/7124/857/3569/3952 | 5 |
| BP | GO:0031099 | regeneration | 0 | 0 | 0 | 1636/2876/7124/2671/7351/3569/8431 | 7 |
| BP | GO:0045088 | regulation of innate immune response | 0 | 0 | 0 | 2150/5468/348/3630/7124/2671/857/5770/948/3952 | 10 |
| BP | GO:0072329 | monocarboxylic acid catabolic process | 0 | 0 | 0 | 34/208/9370/5465/3667/3952 | 6 |
| BP | GO:0043122 | regulation of canonical NF-kappaB signal transduction | 0 | 0 | 0 | 3162/2150/7124/3553/23411/9370/948/3552 | 8 |
| BP | GO:0042554 | superoxide anion generation | 0 | 0 | 0 | 6648/2150/6647/1401 | 4 |
| BP | GO:0046627 | negative regulation of insulin receptor signaling pathway | 0 | 0 | 0 | 3553/5770/3667/5167 | 4 |
| BP | GO:0002720 | ositive regulation of cytokine production involved in immune respons | 0 | 0 | 0 | 2150/3553/23411/948/3569 | 5 |
| BP | GO:0071496 | cellular response to external stimulus | 0 | 0 | 0 | 166/4023/1312/3553/5743/23411/7351/5465/3952 | 9 |
| BP | GO:0043123 | positive regulation of canonical NF-kappaB signal transduction | 0 | 0 | 0 | 3162/2150/7124/3553/9370/948/3552 | 7 |
| BP | GO:0010889 | regulation of sequestering of triglyceride | 0 | 0 | 0 | 5468/4023/5465 | 3 |
| BP | GO:0030728 | ovulation | 0 | 0 | 0 | 9510/23411/3952 | 3 |
| BP | GO:0045725 | positive regulation of glycogen biosynthetic process | 0 | 0 | 0 | 3630/208/3667 | 3 |
| BP | GO:0046321 | positive regulation of fatty acid oxidation | 0 | 0 | 0 | 208/5465/3667 | 3 |
| BP | GO:2001138 | regulation of phospholipid transport | 0 | 0 | 0 | 348/335/1071 | 3 |
| BP | GO:2001279 | regulation of unsaturated fatty acid biosynthetic process | 0 | 0 | 0 | 3553/5743/23411 | 3 |
| BP | GO:0007586 | digestion | 0 | 0 | 0 | 335/948/3952/949/4864/3949 | 6 |
| BP | GO:0007260 | tyrosine phosphorylation of STAT protein | 0 | 0 | 0 | 3066/7124/857/3569/3952 | 5 |
| BP | GO:1900077 | negative regulation of cellular response to insulin stimulus | 0 | 0 | 0 | 3553/5770/3667/5167 | 4 |
| BP | GO:0045862 | positive regulation of proteolysis | 0 | 0 | 0 | 3066/5468/348/7124/3553/857/23411/7133/7466 | 9 |
| BP | GO:0019935 | cyclic-nucleotide-mediated signaling | 0 | 0 | 0 | 348/3630/196/948/5443 | 5 |
| BP | GO:0042310 | vasoconstriction | 0 | 0 | 0 | 1636/857/5743/1401/3952 | 5 |
| BP | GO:0032409 | regulation of transporter activity | 0 | 0 | 0 | 5468/3688/3630/857/6347/9370/4318/481 | 8 |
| BP | GO:0071333 | cellular response to glucose stimulus | 0 | 0 | 0 | 1636/2876/26291/7351/2641/3383 | 6 |
| BP | GO:0042391 | regulation of membrane potential | 0 | 0 | 0 | 6648/7124/208/857/5318/6647/7351/948/153/481 | 10 |
| BP | GO:0030522 | intracellular receptor signaling pathway | 0 | 0 | 0 | 2150/5468/7124/857/23411/5465/196/948/3952 | 9 |
| BP | GO:1990778 | protein localization to cell periphery | 0 | 0 | 0 | 7879/3688/3630/7124/208/857/5318/9370/481 | 9 |
| BP | GO:0014002 | astrocyte development | 0 | 0 | 0 | 7124/3553/3569/3949 | 4 |
| BP | GO:1904646 | cellular response to amyloid-beta | 0 | 0 | 0 | 7124/948/3383/4363 | 4 |
| BP | GO:0071331 | cellular response to hexose stimulus | 0 | 0 | 0 | 1636/2876/26291/7351/2641/3383 | 6 |
| BP | GO:0032930 | positive regulation of superoxide anion generation | 0 | 0 | 0 | 2150/6647/1401 | 3 |
| BP | GO:0034374 | low-density lipoprotein particle remodeling | 0 | 0 | 0 | 348/1071/3990 | 3 |
| BP | GO:0034638 | phosphatidylcholine catabolic process | 0 | 0 | 0 | 3990/949/3949 | 3 |
| BP | GO:1904177 | regulation of adipose tissue development | 0 | 0 | 0 | 5468/4023/23411 | 3 |
| BP | GO:0006509 | membrane protein ectodomain proteolysis | 0 | 0 | 0 | 348/7124/3553/7133 | 4 |
| BP | GO:0009409 | response to cold | 0 | 0 | 0 | 34/7351/153/5346 | 4 |
| BP | GO:0045840 | positive regulation of mitotic nuclear division | 0 | 0 | 0 | 3630/7124/3553/3552 | 4 |
| BP | GO:1903426 | regulation of reactive oxygen species biosynthetic process | 0 | 0 | 0 | 6648/3630/5465/948 | 4 |
| BP | GO:0008625 | extrinsic apoptotic signaling pathway via death domain receptors | 0 | 0 | 0 | 3162/2876/7124/3383/5054 | 5 |
| BP | GO:0010038 | response to metal ion | 0 | 0 | 0 | 3162/6648/857/5743/7351/4968/4318/3552/4864 | 9 |
| BP | GO:0046887 | positive regulation of hormone secretion | 0 | 0 | 0 | 5468/3630/5573/2641/3952/8431 | 6 |
| BP | GO:0010232 | vascular transport | 0 | 0 | 0 | 348/6581/3953/948/4363 | 5 |
| BP | GO:0150104 | transport across blood-brain barrier | 0 | 0 | 0 | 348/6581/3953/948/4363 | 5 |
| BP | GO:0090278 | negative regulation of peptide hormone secretion | 0 | 0 | 0 | 2150/7351/3667/3952 | 4 |
| BP | GO:0043393 | regulation of protein binding | 0 | 0 | 0 | 166/348/857/3416/9370/4318 | 6 |
| BP | GO:0043405 | regulation of MAP kinase activity | 0 | 0 | 0 | 5468/348/7124/3553/5770/9370 | 6 |
| BP | GO:0045860 | positive regulation of protein kinase activity | 0 | 0 | 0 | 1636/3630/7124/3553/5770/23411/9370/3952 | 8 |
| BP | GO:0014074 | response to purine-containing compound | 0 | 0 | 0 | 3066/3553/5743/9370/196/5167 | 6 |
| BP | GO:0044320 | cellular response to leptin stimulus | 0 | 0 | 0 | 23411/3953/3952 | 3 |
| BP | GO:0055089 | fatty acid homeostasis | 0 | 0 | 0 | 348/3630/23411 | 3 |
| BP | GO:0070875 | positive regulation of glycogen metabolic process | 0 | 0 | 0 | 3630/208/3667 | 3 |
| BP | GO:0002792 | negative regulation of peptide secretion | 0 | 0 | 0 | 2150/7351/3667/3952 | 4 |
| BP | GO:0001508 | action potential | 0 | 0 | 0 | 7124/857/5318/6647/948/481 | 6 |
| BP | GO:0003044 | ulation of systemic arterial blood pressure mediated by a chemical sig | 0 | 0 | 0 | 1636/6648/2150/153 | 4 |

| | | | | | | | |
|---|---|---|---|---|---|---|---|
| BP | GO:0042098 | T cell proliferation | 0 | 0 | 0 | 7356/3553/5573/7133/3569/3952/3552 | 7 |
| BP | GO:0002064 | epithelial cell development | 0 | 0 | 0 | 2150/2876/7124/3553/9370/3383/3552 | 7 |
| BP | GO:0043270 | positive regulation of monoatomic ion transport | 0 | 0 | 0 | 857/5318/6347/2641/3952/7466/481 | 7 |
| BP | GO:0072659 | protein localization to plasma membrane | 0 | 0.001 | 0 | 7879/3688/3630/7124/208/5318/9370/481 | 8 |
| BP | GO:0016264 | gap junction assembly | 0 | 0.001 | 0 | 1636/3553/857 | 3 |
| BP | GO:0035234 | ectopic germ cell programmed cell death | 0 | 0.001 | 0 | 3553/6647/3552 | 3 |
| BP | GO:0051000 | positive regulation of nitric-oxide synthase activity | 0 | 0.001 | 0 | 348/3630/949 | 3 |
| BP | GO:0150078 | positive regulation of neuroinflammatory response | 0 | 0.001 | 0 | 7124/3553/3569 | 3 |
| BP | GO:1905288 | vascular associated smooth muscle cell apoptotic process | 0 | 0.001 | 0 | 6648/1786/5468 | 3 |
| BP | GO:1905459 | egulation of vascular associated smooth muscle cell apoptotic proces | 0 | 0.001 | 0 | 6648/1786/5468 | 3 |
| BP | GO:0002687 | positive regulation of leukocyte migration | 0 | 0.001 | 0 | 2150/7124/3383/3569/5054/3552 | 6 |
| BP | GO:0031960 | response to corticosteroid | 0 | 0.001 | 0 | 1636/7356/7124/5743/9370/3569 | 6 |
| BP | GO:0002443 | leukocyte mediated immunity | 0 | 0.001 | 0 | 1636/2150/7124/3553/2671/7133/196/3383/3569/3952 | 10 |
| BP | GO:0061028 | establishment of endothelial barrier | 0 | 0.001 | 0 | 2150/7124/3553/3383 | 4 |
| BP | GO:0010632 | regulation of epithelial cell migration | 0 | 0.001 | 0 | 3162/5468/348/7124/5743/23411/4318/949 | 8 |
| BP | GO:0003073 | regulation of systemic arterial blood pressure | 0 | 0.001 | 0 | 1636/6648/2150/7124/153 | 5 |
| BP | GO:1902041 | lation of extrinsic apoptotic signaling pathway via death domain recep | 0 | 0.001 | 0 | 3162/2876/3383/5054 | 4 |
| BP | GO:0032928 | regulation of superoxide anion generation | 0 | 0.001 | 0 | 2150/6647/1401 | 3 |
| BP | GO:0045721 | negative regulation of gluconeogenesis | 0 | 0.001 | 0 | 3630/9370/3953 | 3 |
| BP | GO:0043535 | regulation of blood vessel endothelial cell migration | 0 | 0.001 | 0 | 3162/5468/348/7124/5743/23411 | 6 |
| BP | GO:0014909 | smooth muscle cell migration | 0 | 0.001 | 0 | 1636/5327/9510/9370/5054 | 5 |
| BP | GO:0007249 | canonical NF-kappaB signal transduction | 0 | 0.001 | 0 | 3162/2150/7124/3553/23411/9370/948/3552 | 8 |
| BP | GO:1905039 | carboxylic acid transmembrane transport | 0 | 0.001 | 0 | 3688/7124/208/7351/948/4363 | 6 |
| BP | GO:1903825 | organic acid transmembrane transport | 0 | 0.001 | 0 | 3688/7124/208/7351/948/4363 | 6 |
| BP | GO:1904707 | sitive regulation of vascular associated smooth muscle cell proliferati | 0 | 0.001 | 0 | 1786/7124/9510/4318 | 4 |
| BP | GO:0031998 | regulation of fatty acid beta-oxidation | 0 | 0.001 | 0 | 208/5465/3667 | 3 |
| BP | GO:0055093 | response to hyperoxia | 0 | 0.001 | 0 | 6648/3066/857 | 3 |
| BP | GO:0060353 | regulation of cell adhesion molecule production | 0 | 0.001 | 0 | 3553/335/857 | 3 |
| BP | GO:0072567 | chemokine (C-X-C motif) ligand 2 production | 0 | 0.001 | 0 | 2150/4023/7124 | 3 |
| BP | GO:2000341 | regulation of chemokine (C-X-C motif) ligand 2 production | 0 | 0.001 | 0 | 2150/4023/7124 | 3 |
| BP | GO:0003300 | cardiac muscle hypertrophy | 0 | 0.001 | 0 | 3066/5468/7133/5465/3952 | 5 |
| BP | GO:0046651 | lymphocyte proliferation | 0 | 0.001 | 0 | 7356/3553/5573/7133/196/3569/3952/3552 | 8 |
| BP | GO:0010594 | regulation of endothelial cell migration | 0 | 0.001 | 0 | 3162/5468/348/7124/5743/23411/949 | 7 |
| BP | GO:0019932 | second-messenger-mediated signaling | 0 | 0.001 | 0 | 348/3630/7124/196/948/283455/5443/481 | 8 |
| BP | GO:0120009 | intermembrane lipid transfer | 0 | 0.001 | 0 | 348/335/1071/10577 | 4 |
| BP | GO:0044321 | response to leptin | 0 | 0.001 | 0 | 23411/3953/3952 | 3 |
| BP | GO:0045722 | positive regulation of gluconeogenesis | 0 | 0.001 | 0 | 23411/5465/2641 | 3 |
| BP | GO:0014897 | striated muscle hypertrophy | 0 | 0.001 | 0 | 3066/5468/7133/5465/3952 | 5 |
| BP | GO:0052547 | regulation of peptidase activity | 0 | 0.001 | 0 | 2876/5468/7124/857/5265/23411/5054/4318 | 8 |
| BP | GO:0002440 | production of molecular mediator of immune response | 0 | 0.001 | 0 | 2150/7124/3553/335/23411/7133/948/3569 | 8 |
| BP | GO:0002573 | myeloid leukocyte differentiation | 0 | 0.001 | 0 | 2150/5468/7124/23411/9370/7351/4318 | 7 |
| BP | GO:0045861 | negative regulation of proteolysis | 0 | 0.001 | 0 | 5327/2876/3630/7124/5265/5054/4318 | 7 |
| BP | GO:0014896 | muscle hypertrophy | 0 | 0.001 | 0 | 3066/5468/7133/5465/3952 | 5 |
| BP | GO:0033138 | positive regulation of peptidyl-serine phosphorylation | 0 | 0.001 | 0 | 7124/857/5743/2641/3569 | 5 |
| BP | GO:0060986 | endocrine hormone secretion | 0 | 0.001 | 0 | 5468/3553/3952/5443 | 4 |
| BP | GO:0003085 | negative regulation of systemic arterial blood pressure | 0 | 0.001 | 0 | 6648/7124/153 | 3 |
| BP | GO:0010866 | regulation of triglyceride biosynthetic process | 0 | 0.001 | 0 | 23411/949/3949 | 3 |
| BP | GO:0033762 | response to glucagon | 0 | 0.001 | 0 | 5573/26291/2641 | 3 |
| BP | GO:0090208 | positive regulation of triglyceride metabolic process | 0 | 0.001 | 0 | 26291/949/3949 | 3 |
| BP | GO:0050678 | regulation of epithelial cell proliferation | 0 | 0.001 | 0 | 3162/2876/5468/348/7124/857/6347/23411/3952 | 9 |
| BP | GO:0061900 | glial cell activation | 0 | 0.001 | 0 | 7124/3553/3569/3949 | 4 |
| BP | GO:0030198 | extracellular matrix organization | 0 | 0.001 | 0 | 3688/10516/7124/857/9510/7133/3569/4318 | 8 |
| BP | GO:0043062 | extracellular structure organization | 0 | 0.001 | 0.001 | 3688/10516/7124/857/9510/7133/3569/4318 | 8 |
| BP | GO:0045229 | external encapsulating structure organization | 0 | 0.001 | 0.001 | 3688/10516/7124/857/9510/7133/3569/4318 | 8 |
| BP | GO:0050769 | positive regulation of neurogenesis | 0 | 0.001 | 0.001 | 1636/3066/3688/7124/3553/7133/3569 | 7 |
| BP | GO:0061756 | leukocyte adhesion to vascular endothelial cell | 0 | 0.001 | 0.001 | 3688/7124/3569/3952 | 4 |
| BP | GO:0060252 | positive regulation of glial cell proliferation | 0 | 0.001 | 0.001 | 7124/3553/3569 | 3 |
| BP | GO:0060352 | cell adhesion molecule production | 0 | 0.001 | 0.001 | 3553/335/857 | 3 |
| BP | GO:0030856 | regulation of epithelial cell differentiation | 0 | 0.001 | 0.001 | 7124/3553/857/5054/4318/3552 | 6 |
| BP | GO:0051091 | positive regulation of DNA-binding transcription factor activity | 0 | 0.001 | 0.001 | 5468/3630/7124/3553/857/948/3569 | 7 |
| BP | GO:0010574 | regulation of vascular endothelial growth factor production | 0 | 0.001 | 0.001 | 3553/5743/3569/3552 | 4 |
| BP | GO:0014009 | glial cell proliferation | 0 | 0.001 | 0.001 | 7124/3553/3953/3569 | 4 |
| BP | GO:0033619 | membrane protein proteolysis | 0 | 0.001 | 0.001 | 348/7124/3553/7133 | 4 |
| BP | GO:0098900 | regulation of action potential | 0 | 0.001 | 0.001 | 7124/857/5318/948 | 4 |
| BP | GO:0014812 | muscle cell migration | 0 | 0.001 | 0.001 | 1636/5327/9510/9370/5054 | 5 |
| BP | GO:0042692 | muscle cell differentiation | 0 | 0.001 | 0.001 | 9518/6648/2876/1786/3688/34/23411/5573/5465 | 9 |
| BP | GO:0007259 | receptor signaling pathway via JAK-STAT | 0 | 0.001 | 0.001 | 3066/7124/857/6347/3569/3952 | 6 |
| BP | GO:0006695 | cholesterol biosynthetic process | 0 | 0.001 | 0.001 | 348/335/6647/1727 | 4 |
| BP | GO:0051785 | positive regulation of nuclear division | 0 | 0.001 | 0.001 | 3630/7124/3553/3552 | 4 |
| BP | GO:1902653 | secondary alcohol biosynthetic process | 0 | 0.001 | 0.001 | 348/335/6647/1727 | 4 |
| BP | GO:0033209 | tumor necrosis factor-mediated signaling pathway | 0 | 0.001 | 0.001 | 2150/7124/335/9370/7133 | 5 |
| BP | GO:0032891 | negative regulation of organic acid transport | 0 | 0.001 | 0.001 | 7124/208/3952 | 3 |
| BP | GO:0034138 | toll-like receptor 3 signaling pathway | 0 | 0.001 | 0.001 | 2150/7124/857 | 3 |
| BP | GO:0035357 | peroxisome proliferator activated receptor signaling pathway | 0 | 0.001 | 0.001 | 5468/23411/3952 | 3 |
| BP | GO:2000209 | regulation of anoikis | 0 | 0.001 | 0.001 | 166/3688/857 | 3 |
| BP | GO:0032092 | positive regulation of protein binding | 0 | 0.001 | 0.001 | 348/857/3416/4318 | 4 |
| BP | GO:0051924 | regulation of calcium ion transport | 0 | 0.001 | 0.001 | 1636/857/5743/6347/2641/7466/481 | 7 |
| BP | GO:1901617 | organic hydroxy compound biosynthetic process | 0 | 0.001 | 0.001 | 348/7124/335/23411/6647/3952/1727 | 7 |
| BP | GO:0032386 | regulation of intracellular transport | 0 | 0.001 | 0.001 | 5116/3553/208/5743/5770/9370/948/3952 | 8 |
| BP | GO:0090303 | positive regulation of wound healing | 0 | 0.001 | 0.001 | 5327/3688/948/5054 | 4 |
| BP | GO:0006706 | steroid catabolic process | 0 | 0.001 | 0.001 | 348/3290/949 | 3 |
| BP | GO:0010623 | programmed cell death involved in cell development | 0 | 0.001 | 0.001 | 3553/6647/3552 | 3 |
| BP | GO:0086064 | ll communication by electrical coupling involved in cardiac conductio | 0 | 0.001 | 0.001 | 857/5318/481 | 3 |
| BP | GO:0040014 | regulation of multicellular organism growth | 0 | 0.001 | 0.001 | 9518/6647/79068/153 | 4 |
| BP | GO:0006898 | receptor-mediated endocytosis | 0 | 0.001 | 0.001 | 348/3688/857/9370/948/5054/3949 | 7 |
| BP | GO:0019229 | regulation of vasoconstriction | 0 | 0.001 | 0.001 | 1636/857/5743/3952 | 4 |
| BP | GO:0030225 | macrophage differentiation | 0 | 0.001 | 0.001 | 23411/9370/7351/4318 | 4 |
| BP | GO:0043500 | muscle adaptation | 0 | 0.001 | 0.001 | 3162/5468/3553/7133/5465 | 5 |
| BP | GO:0050830 | defense response to Gram-positive bacterium | 0 | 0.001 | 0.001 | 7124/3553/948/1401/3569 | 5 |
| BP | GO:0051928 | positive regulation of calcium ion transport | 0 | 0.001 | 0.001 | 857/6347/2641/7466/481 | 5 |
| BP | GO:0050435 | amyloid-beta metabolic process | 0 | 0.001 | 0.001 | 1636/348/7124/3416 | 4 |
| BP | GO:0045766 | positive regulation of angiogenesis | 0 | 0.001 | 0.001 | 3162/3688/3553/23411/5054/3552 | 6 |
| BP | GO:1904018 | positive regulation of vasculature development | 0 | 0.001 | 0.001 | 3162/3688/3553/23411/5054/3552 | 6 |
| BP | GO:1904892 | regulation of receptor signaling pathway via STAT | 0 | 0.001 | 0.001 | 5468/7124/857/3569/3952 | 5 |
| BP | GO:0051098 | regulation of binding | 0 | 0.001 | 0.001 | 166/5468/348/857/3416/9370/4318 | 7 |
| BP | GO:0010804 | egative regulation of tumor necrosis factor-mediated signaling pathwa | 0 | 0.001 | 0.001 | 2150/335/9370 | 3 |
| BP | GO:0010884 | positive regulation of lipid storage | 0 | 0.001 | 0.001 | 4023/948/949 | 3 |
| BP | GO:0036296 | response to increased oxygen levels | 0 | 0.001 | 0.001 | 6648/3066/857 | 3 |
| BP | GO:0050995 | negative regulation of lipid catabolic process | 0 | 0.001 | 0.001 | 3630/7124/3553 | 3 |
| BP | GO:2000272 | negative regulation of signaling receptor activity | 0 | 0.001 | 0.001 | 5468/7124/5465 | 3 |
| BP | GO:0042531 | positive regulation of tyrosine phosphorylation of STAT protein | 0 | 0.002 | 0.001 | 3066/7124/3569/3952 | 4 |
| BP | GO:0009895 | negative regulation of catabolic process | 0 | 0.002 | 0.001 | 3162/3630/7124/3553/5465/3953/3952/4864 | 8 |
| BP | GO:0016126 | sterol biosynthetic process | 0 | 0.002 | 0.001 | 348/335/6647/1727 | 4 |
| BP | GO:0032874 | positive regulation of stress-activated MAPK cascade | 0 | 0.002 | 0.001 | 2150/7124/3553/3952/3552 | 5 |
| BP | GO:0019433 | triglyceride catabolic process | 0 | 0.002 | 0.001 | 4023/3990/26291 | 3 |
| BP | GO:0030194 | positive regulation of blood coagulation | 0 | 0.002 | 0.001 | 5327/948/5054 | 3 |
| BP | GO:1900048 | positive regulation of hemostasis | 0 | 0.002 | 0.001 | 5327/948/5054 | 3 |
| BP | GO:0045446 | endothelial cell differentiation | 0 | 0.002 | 0.001 | 2150/2876/7124/3553/3383 | 5 |
| BP | GO:0070304 | positive regulation of stress-activated protein kinase signaling cascad | 0 | 0.002 | 0.001 | 2150/7124/3553/3952/3552 | 5 |

| | | | | | | | |
|---|---|---|---|---|---|---|---|
| BP | GO:0030258 | lipid modification | 0 | 0.002 | 0.001 | 34/208/9370/5465/3667/3952 | 6 |
| BP | GO:0071466 | cellular response to xenobiotic stimulus | 0 | 0.002 | 0.001 | 3553/26291/9370/196/4363/5447 | 6 |
| BP | GO:0002698 | negative regulation of immune effector process | 0 | 0.002 | 0.001 | 3630/7124/2671/335/196 | 5 |
| BP | GO:0006690 | icosanoid metabolic process | 0 | 0.002 | 0.001 | 2876/3553/5743/23411/4363 | 5 |
| BP | GO:0006910 | phagocytosis, recognition | 0 | 0.002 | 0.001 | 948/1401/949 | 3 |
| BP | GO:0071897 | DNA biosynthetic process | 0 | 0.002 | 0.001 | 7124/5422/2671/23411/9370/5428 | 6 |
| BP | GO:0046683 | response to organophosphorus | 0 | 0.002 | 0.001 | 3553/5743/9370/196/5167 | 5 |
| BP | GO:0051099 | positive regulation of binding | 0 | 0.002 | 0.001 | 5468/348/857/3416/4318 | 5 |
| BP | GO:1903039 | positive regulation of leukocyte cell-cell adhesion | 0 | 0.002 | 0.001 | 7124/3553/857/6347/3569/3952/3552 | 7 |
| BP | GO:0045600 | positive regulation of fat cell differentiation | 0 | 0.002 | 0.001 | 5468/4023/3630/5743 | 4 |
| BP | GO:0007369 | gastrulation | 0 | 0.002 | 0.001 | 3688/1291/335/5573/9370/4318 | 6 |
| BP | GO:0010951 | negative regulation of endopeptidase activity | 0 | 0.002 | 0.001 | 2876/7124/5265/5054/4318 | 5 |
| BP | GO:0007263 | nitric oxide mediated signal transduction | 0 | 0.002 | 0.001 | 348/3630/948 | 3 |
| BP | GO:0050820 | positive regulation of coagulation | 0 | 0.002 | 0.001 | 5327/948/5054 | 3 |
| BP | GO:0002456 | T cell mediated immunity | 0 | 0.002 | 0.001 | 3553/7133/196/3383/3569 | 5 |
| BP | GO:0002449 | lymphocyte mediated immunity | 0 | 0.002 | 0.001 | 7124/3553/2671/7133/196/3383/3569/3952 | 8 |
| BP | GO:0051881 | regulation of mitochondrial membrane potential | 0 | 0.002 | 0.001 | 6648/208/6647/7351 | 4 |
| BP | GO:0031668 | cellular response to extracellular stimulus | 0 | 0.002 | 0.001 | 166/4023/1312/23411/7351/5465/3952 | 7 |
| BP | GO:2000273 | positive regulation of signaling receptor activity | 0 | 0.002 | 0.001 | 3066/3688/6347 | 3 |
| BP | GO:0030336 | negative regulation of cell migration | 0 | 0.002 | 0.001 | 3162/3066/5468/348/7124/6347/9370/5054 | 8 |
| BP | GO:0044282 | small molecule catabolic process | 0 | 0.002 | 0.001 | 348/34/208/9370/5465/3667/3952/949 | 8 |
| BP | GO:0061045 | negative regulation of wound healing | 0 | 0.002 | 0.001 | 5327/348/7124/5054 | 4 |
| BP | GO:0001516 | prostaglandin biosynthetic process | 0.001 | 0.002 | 0.001 | 3553/5743/23411 | 3 |
| BP | GO:0010644 | cell communication by electrical coupling | 0.001 | 0.002 | 0.001 | 857/5318/481 | 3 |
| BP | GO:0032367 | intracellular cholesterol transport | 0.001 | 0.002 | 0.001 | 10577/4864/3949 | 3 |
| BP | GO:0045907 | positive regulation of vasoconstriction | 0.001 | 0.002 | 0.001 | 1636/857/5743 | 3 |
| BP | GO:0046457 | prostanoid biosynthetic process | 0.001 | 0.002 | 0.001 | 3553/5743/23411 | 3 |
| BP | GO:0060055 | angiogenesis involved in wound healing | 0.001 | 0.002 | 0.001 | 2876/7124/5054 | 3 |
| BP | GO:1901889 | negative regulation of cell junction assembly | 0.001 | 0.002 | 0.001 | 1636/7124/3553 | 3 |
| BP | GO:0052548 | regulation of endopeptidase activity | 0.001 | 0.002 | 0.001 | 2876/5468/7124/5265/23411/5054/4318 | 7 |
| BP | GO:0042157 | lipoprotein metabolic process | 0.001 | 0.002 | 0.001 | 348/335/5465/3952/3949 | 5 |
| BP | GO:0002696 | positive regulation of leukocyte activation | 0.001 | 0.003 | 0.001 | 2150/7124/3553/857/6347/3569/3952/3552 | 8 |
| BP | GO:0002702 | itive regulation of production of molecular mediator of immune respo | 0.001 | 0.003 | 0.001 | 2150/3553/23411/948/3569 | 5 |
| BP | GO:0002819 | regulation of adaptive immune response | 0.001 | 0.003 | 0.001 | 7124/3553/23411/7133/196/3569 | 6 |
| BP | GO:0051962 | positive regulation of nervous system development | 0.001 | 0.003 | 0.001 | 1636/3066/3688/7124/3553/7133/3569 | 7 |
| BP | GO:1903036 | positive regulation of response to wounding | 0.001 | 0.003 | 0.001 | 5327/3688/948/5054 | 4 |
| BP | GO:0001818 | negative regulation of cytokine production | 0.001 | 0.003 | 0.001 | 7356/2150/7124/335/9370/5465/3569/5443 | 8 |
| BP | GO:0019934 | cGMP-mediated signaling | 0.001 | 0.003 | 0.001 | 348/3630/948 | 3 |
| BP | GO:0033198 | response to ATP | 0.001 | 0.003 | 0.001 | 3553/5743/5167 | 3 |
| BP | GO:0090279 | regulation of calcium ion import | 0.001 | 0.003 | 0.001 | 1636/6347/2641 | 3 |
| BP | GO:1902235 | n of endoplasmic reticulum stress-induced intrinsic apoptotic signaling | 0.001 | 0.003 | 0.001 | 5770/23411/7466 | 3 |
| BP | GO:0010466 | negative regulation of peptidase activity | 0.001 | 0.003 | 0.001 | 2876/7124/5265/5054/4318 | 5 |
| BP | GO:1903131 | mononuclear cell differentiation | 0.001 | 0.003 | 0.001 | 2150/5468/3688/3553/6647/3953/3569/3952/3552 | 9 |
| BP | GO:0045785 | positive regulation of cell adhesion | 0.001 | 0.003 | 0.001 | 7124/3553/335/857/6347/948/3569/3952/3552 | 9 |
| BP | GO:0033135 | regulation of peptidyl-serine phosphorylation | 0.001 | 0.003 | 0.001 | 7124/857/5743/2641/3569 | 5 |
| BP | GO:0045685 | regulation of glial cell differentiation | 0.001 | 0.003 | 0.001 | 3066/7133/3569/3949 | 4 |
| BP | GO:0043276 | anoikis | 0.001 | 0.003 | 0.001 | 166/3688/857 | 3 |
| BP | GO:0046320 | regulation of fatty acid oxidation | 0.001 | 0.003 | 0.001 | 208/5465/3667 | 3 |
| BP | GO:0046475 | glycerophospholipid catabolic process | 0.001 | 0.003 | 0.001 | 3990/949/3949 | 3 |
| BP | GO:0050901 | leukocyte tethering or rolling | 0.001 | 0.003 | 0.001 | 3688/7124/3952 | 3 |
| BP | GO:0071398 | cellular response to fatty acid | 0.001 | 0.003 | 0.001 | 4023/3667/3949 | 3 |
| BP | GO:0071902 | positive regulation of protein serine/threonine kinase activity | 0.001 | 0.003 | 0.001 | 7124/3553/5770/23411/9370 | 5 |
| BP | GO:2000146 | negative regulation of cell motility | 0.001 | 0.003 | 0.001 | 3162/3066/5468/348/7124/6347/9370/5054 | 8 |
| BP | GO:0045216 | cell-cell junction organization | 0.001 | 0.003 | 0.002 | 1636/2150/7124/3553/857/5318 | 6 |
| BP | GO:0050767 | regulation of neurogenesis | 0.001 | 0.003 | 0.002 | 1636/3066/3688/7124/3553/7133/3569/3949 | 8 |
| BP | GO:0002446 | neutrophil mediated immunity | 0.001 | 0.003 | 0.002 | 1636/2150/3569 | 3 |
| BP | GO:0010614 | negative regulation of cardiac muscle hypertrophy | 0.001 | 0.003 | 0.002 | 5468/7133/5465 | 3 |
| BP | GO:0032366 | intracellular sterol transport | 0.001 | 0.003 | 0.002 | 10577/4864/3949 | 3 |
| BP | GO:1901099 | negative regulation of signal transduction in absence of ligand | 0.001 | 0.003 | 0.002 | 7124/3553/3552 | 3 |
| BP | GO:1903725 | regulation of phospholipid metabolic process | 0.001 | 0.003 | 0.002 | 2784/949/3949 | 3 |
| BP | GO:2001240 | ive regulation of extrinsic apoptotic signaling pathway in absence of l | 0.001 | 0.003 | 0.002 | 7124/3553/3552 | 3 |
| BP | GO:0002534 | cytokine production involved in inflammatory response | 0.001 | 0.003 | 0.002 | 7124/5465/3569/3952 | 4 |
| BP | GO:1900015 | regulation of cytokine production involved in inflammatory response | 0.001 | 0.003 | 0.002 | 7124/5465/3569/3952 | 4 |
| BP | GO:0003158 | endothelium development | 0.001 | 0.003 | 0.002 | 2150/2876/7124/3553/3383 | 5 |
| BP | GO:1905897 | regulation of response to endoplasmic reticulum stress | 0.001 | 0.003 | 0.002 | 857/5770/23411/7466 | 4 |
| BP | GO:0050867 | positive regulation of cell activation | 0.001 | 0.003 | 0.002 | 2150/7124/3553/857/6347/3569/3952/3552 | 8 |
| BP | GO:2000352 | negative regulation of endothelial cell apoptotic process | 0.001 | 0.003 | 0.002 | 26291/3383/5054 | 3 |
| BP | GO:0046165 | alcohol biosynthetic process | 0.001 | 0.003 | 0.002 | 348/335/6647/3952/1727 | 5 |
| BP | GO:0035023 | regulation of Rho protein signal transduction | 0.001 | 0.003 | 0.002 | 2150/348/3688/335 | 4 |
| BP | GO:0046434 | organophosphate catabolic process | 0.001 | 0.003 | 0.002 | 3990/4968/5167/949/3949 | 5 |
| BP | GO:0010660 | regulation of muscle cell apoptotic process | 0.001 | 0.003 | 0.002 | 6648/1786/5468/23411 | 4 |
| BP | GO:0014741 | negative regulation of muscle hypertrophy | 0.001 | 0.003 | 0.002 | 5468/7133/5465 | 3 |
| BP | GO:0071276 | cellular response to cadmium ion | 0.001 | 0.003 | 0.002 | 3162/4968/4318 | 3 |
| BP | GO:0034103 | regulation of tissue remodeling | 0.001 | 0.003 | 0.002 | 5468/3953/3569/3952 | 4 |
| BP | GO:0002369 | T cell cytokine production | 0.001 | 0.004 | 0.002 | 3553/7133/3569 | 3 |
| BP | GO:0002724 | regulation of T cell cytokine production | 0.001 | 0.004 | 0.002 | 3553/7133/3569 | 3 |
| BP | GO:0035633 | maintenance of blood-brain barrier | 0.001 | 0.004 | 0.002 | 3688/5743/3569 | 3 |
| BP | GO:0046461 | neutral lipid catabolic process | 0.001 | 0.004 | 0.002 | 4023/3990/26291 | 3 |
| BP | GO:0046464 | acylglycerol catabolic process | 0.001 | 0.004 | 0.002 | 4023/3990/26291 | 3 |
| BP | GO:0002832 | negative regulation of response to biotic stimulus | 0.001 | 0.004 | 0.002 | 2150/5468/3630/2671/196 | 5 |
| BP | GO:0051092 | positive regulation of NF-kappaB transcription factor activity | 0.001 | 0.004 | 0.002 | 3630/7124/3553/857/948 | 5 |
| BP | GO:0048708 | astrocyte differentiation | 0.001 | 0.004 | 0.002 | 7124/3553/3569/3949 | 4 |
| BP | GO:0046330 | positive regulation of JNK cascade | 0.001 | 0.004 | 0.002 | 2150/7124/3553/3552 | 4 |
| BP | GO:0044060 | regulation of endocrine process | 0.001 | 0.004 | 0.002 | 2150/3952/5443 | 3 |
| BP | GO:0050890 | cognition | 0.001 | 0.004 | 0.002 | 5577/348/3688/3630/7124/5743/3949 | 7 |
| BP | GO:0014910 | regulation of smooth muscle cell migration | 0.001 | 0.004 | 0.002 | 1636/9510/9370/5054 | 4 |
| BP | GO:0046849 | bone remodeling | 0.001 | 0.004 | 0.002 | 7879/3953/3569/3952 | 4 |
| BP | GO:0033028 | myeloid cell apoptotic process | 0.001 | 0.004 | 0.002 | 23411/9370/3569 | 3 |
| BP | GO:0046676 | negative regulation of insulin secretion | 0.001 | 0.004 | 0.002 | 2150/7351/3667 | 3 |
| BP | GO:0008406 | gonad development | 0.001 | 0.004 | 0.002 | 1636/9510/23411/6647/3952/3552 | 6 |
| BP | GO:0010507 | negative regulation of autophagy | 0.001 | 0.004 | 0.002 | 3162/3953/3952/4864 | 4 |
| BP | GO:0010657 | muscle cell apoptotic process | 0.001 | 0.004 | 0.002 | 6648/1786/5468/23411 | 4 |
| BP | GO:0022409 | positive regulation of cell-cell adhesion | 0.001 | 0.004 | 0.002 | 7124/3553/857/6347/3569/3952/3552 | 7 |
| BP | GO:0007043 | cell-cell junction assembly | 0.001 | 0.004 | 0.002 | 1636/7124/3553/857/5318 | 5 |
| BP | GO:0015911 | long-chain fatty acid import across plasma membrane | 0.001 | 0.004 | 0.002 | 208/948 | 2 |
| BP | GO:0032096 | negative regulation of response to food | 0.001 | 0.004 | 0.002 | 5465/3952 | 2 |
| BP | GO:0032099 | negative regulation of appetite | 0.001 | 0.004 | 0.002 | 5465/3952 | 2 |
| BP | GO:0032105 | negative regulation of response to extracellular stimulus | 0.001 | 0.004 | 0.002 | 5465/3952 | 2 |
| BP | GO:0032108 | negative regulation of response to nutrient levels | 0.001 | 0.004 | 0.002 | 5465/3952 | 2 |
| BP | GO:0033034 | positive regulation of myeloid cell apoptotic process | 0.001 | 0.004 | 0.002 | 23411/9370 | 2 |
| BP | GO:0045541 | negative regulation of cholesterol biosynthetic process | 0.001 | 0.004 | 0.002 | 348/6647 | 2 |
| BP | GO:0070874 | negative regulation of glycogen metabolic process | 0.001 | 0.004 | 0.002 | 3630/5167 | 2 |
| BP | GO:0097050 | type B pancreatic cell apoptotic process | 0.001 | 0.004 | 0.002 | 3569/7466 | 2 |
| BP | GO:0106119 | negative regulation of sterol biosynthetic process | 0.001 | 0.004 | 0.002 | 348/6647 | 2 |
| BP | GO:0150172 | regulation of phosphatidylcholine metabolic process | 0.001 | 0.004 | 0.002 | 949/3949 | 2 |
| BP | GO:1903044 | protein localization to membrane raft | 0.001 | 0.004 | 0.002 | 857/481 | 2 |
| BP | GO:1904729 | regulation of intestinal lipid absorption | 0.001 | 0.004 | 0.002 | 335/3952 | 2 |
| BP | GO:1905918 | regulation of CoA-transferase activity | 0.001 | 0.004 | 0.002 | 348/335 | 2 |
| BP | GO:2000121 | regulation of removal of superoxide radicals | 0.001 | 0.004 | 0.002 | 10516/948 | 2 |

| | | | | | | | |
|---|---|---|---|---|---|---|---|
| BP | GO:0060251 | regulation of glial cell proliferation | 0.001 | 0.004 | 0.002 | 7124/3553/3569 | 3 |
| BP | GO:0090316 | positive regulation of intracellular protein transport | 0.001 | 0.004 | 0.002 | 5116/3553/208/5743/3952 | 5 |
| BP | GO:0040013 | negative regulation of locomotion | 0.001 | 0.004 | 0.002 | 3162/3066/5468/348/7124/6347/9370/5054 | 8 |
| BP | GO:0045137 | development of primary sexual characteristics | 0.001 | 0.005 | 0.002 | 1636/9510/23411/6647/3952/3552 | 6 |
| BP | GO:0048638 | regulation of developmental growth | 0.001 | 0.005 | 0.002 | 9518/348/6647/79068/5465/3952/153 | 7 |
| BP | GO:0022898 | regulation of transmembrane transporter activity | 0.001 | 0.005 | 0.002 | 3688/3630/857/6347/4318/481 | 6 |
| BP | GO:1904706 | gative regulation of vascular associated smooth muscle cell proliferati | 0.001 | 0.005 | 0.002 | 6648/5468/9370 | 3 |
| BP | GO:1990000 | amyloid fibril formation | 0.001 | 0.005 | 0.002 | 348/948/3949 | 3 |
| BP | GO:0042742 | defense response to bacterium | 0.001 | 0.005 | 0.003 | 2150/7124/3553/948/1401/3569/5054 | 7 |
| BP | GO:0009314 | response to radiation | 0.001 | 0.005 | 0.003 | 6648/2876/3688/208/23411/4968/4318/3552 | 8 |
| BP | GO:0010712 | regulation of collagen metabolic process | 0.001 | 0.005 | 0.003 | 3066/3688/3569 | 3 |
| BP | GO:0019432 | triglyceride biosynthetic process | 0.001 | 0.005 | 0.003 | 23411/949/3949 | 3 |
| BP | GO:0051955 | regulation of amino acid transport | 0.001 | 0.005 | 0.003 | 3688/7124/3952 | 3 |
| BP | GO:1904037 | positive regulation of epithelial cell apoptotic process | 0.001 | 0.005 | 0.003 | 3162/6347/3569 | 3 |
| BP | GO:0043281 | ation of cysteine-type endopeptidase activity involved in apoptotic pro | 0.001 | 0.005 | 0.003 | 2876/5468/7124/23411/4318 | 5 |
| BP | GO:0001910 | regulation of leukocyte mediated cytotoxicity | 0.001 | 0.005 | 0.003 | 2150/2671/3383/3952 | 4 |
| BP | GO:0010887 | negative regulation of cholesterol storage | 0.001 | 0.005 | 0.003 | 5468/5465 | 2 |
| BP | GO:0034115 | negative regulation of heterotypic cell-cell adhesion | 0.001 | 0.005 | 0.003 | 335/9370 | 2 |
| BP | GO:0034139 | regulation of toll-like receptor 3 signaling pathway | 0.001 | 0.005 | 0.003 | 2150/857 | 2 |
| BP | GO:0042167 | heme catabolic process | 0.001 | 0.005 | 0.003 | 3162/4363 | 2 |
| BP | GO:0042447 | hormone catabolic process | 0.001 | 0.005 | 0.003 | 1636/3416 | 2 |
| BP | GO:0046149 | pigment catabolic process | 0.001 | 0.005 | 0.003 | 3162/4363 | 2 |
| BP | GO:0051593 | response to folic acid | 0.001 | 0.005 | 0.003 | 2876/4968 | 2 |
| BP | GO:0070587 | regulation of cell-cell adhesion involved in gastrulation | 0.001 | 0.005 | 0.003 | 335/9370 | 2 |
| BP | GO:0090206 | negative regulation of cholesterol metabolic process | 0.001 | 0.005 | 0.003 | 348/6647 | 2 |
| BP | GO:1901984 | negative regulation of protein acetylation | 0.001 | 0.005 | 0.003 | 3066/23411 | 2 |
| BP | GO:1905461 | ive regulation of vascular associated smooth muscle cell apoptotic pro | 0.001 | 0.005 | 0.003 | 6648/5468 | 2 |
| BP | GO:2000343 | positive regulation of chemokine (C-X-C motif) ligand 2 production | 0.001 | 0.005 | 0.003 | 4023/7124 | 2 |
| BP | GO:2000644 | regulation of receptor catabolic process | 0.001 | 0.005 | 0.003 | 348/5770 | 2 |
| BP | GO:1903035 | negative regulation of response to wounding | 0.001 | 0.005 | 0.003 | 5327/348/7124/5054 | 4 |
| BP | GO:0051150 | regulation of smooth muscle cell differentiation | 0.001 | 0.006 | 0.003 | 6648/1786/23411 | 3 |
| BP | GO:0140894 | endolysosomal toll-like receptor signaling pathway | 0.001 | 0.006 | 0.003 | 2150/7124/857 | 3 |
| BP | GO:0006633 | fatty acid biosynthetic process | 0.001 | 0.006 | 0.003 | 4023/3553/3990/5743/23411 | 5 |
| BP | GO:0002709 | regulation of T cell mediated immunity | 0.001 | 0.006 | 0.003 | 3553/7133/196/3569 | 4 |
| BP | GO:0031669 | cellular response to nutrient levels | 0.001 | 0.006 | 0.003 | 4023/1312/23411/7351/5465/3952 | 6 |
| BP | GO:0050870 | positive regulation of T cell activation | 0.001 | 0.006 | 0.003 | 3553/857/6347/3569/3952/3552 | 6 |
| BP | GO:0016054 | organic acid catabolic process | 0.001 | 0.006 | 0.003 | 34/208/9370/5465/3667/3952 | 6 |
| BP | GO:0046395 | carboxylic acid catabolic process | 0.001 | 0.006 | 0.003 | 34/208/9370/5465/3667/3952 | 6 |
| BP | GO:0010720 | positive regulation of cell development | 0.001 | 0.006 | 0.003 | 1636/3066/3688/7124/3553/9370/7133/3569 | 8 |
| BP | GO:0032892 | positive regulation of organic acid transport | 0.001 | 0.006 | 0.003 | 3688/3553/3552 | 3 |
| BP | GO:2001239 | egulation of extrinsic apoptotic signaling pathway in absence of ligan | 0.001 | 0.006 | 0.003 | 7124/3553/3552 | 3 |
| BP | GO:0006937 | regulation of muscle contraction | 0.001 | 0.006 | 0.003 | 857/5318/5743/6647/481 | 5 |
| BP | GO:0033083 | regulation of immature T cell proliferation | 0.002 | 0.006 | 0.003 | 3553/3552 | 2 |
| BP | GO:0033084 | regulation of immature T cell proliferation in thymus | 0.002 | 0.006 | 0.003 | 3553/3552 | 2 |
| BP | GO:0040015 | negative regulation of multicellular organism growth | 0.002 | 0.006 | 0.003 | 9518/153 | 2 |
| BP | GO:0070586 | cell-cell adhesion involved in gastrulation | 0.002 | 0.006 | 0.003 | 335/9370 | 2 |
| BP | GO:0090677 | reversible differentiation | 0.002 | 0.006 | 0.003 | 6648/1786 | 2 |
| BP | GO:0098911 | regulation of ventricular cardiac muscle cell action potential | 0.002 | 0.006 | 0.003 | 857/5318 | 2 |
| BP | GO:1904478 | regulation of intestinal absorption | 0.002 | 0.006 | 0.003 | 335/3952 | 2 |
| BP | GO:0008585 | female gonad development | 0.002 | 0.006 | 0.003 | 9510/23411/6647/3952 | 4 |
| BP | GO:0002027 | regulation of heart rate | 0.002 | 0.006 | 0.003 | 7124/857/5318/153 | 4 |
| BP | GO:0008630 | intrinsic apoptotic signaling pathway in response to DNA damage | 0.002 | 0.006 | 0.003 | 6648/7124/23411/7133 | 4 |
| BP | GO:0050680 | negative regulation of epithelial cell proliferation | 0.002 | 0.006 | 0.003 | 5468/348/7124/857/6347 | 5 |
| BP | GO:0050806 | positive regulation of synaptic transmission | 0.002 | 0.006 | 0.003 | 348/3630/7124/5743/6347 | 5 |
| BP | GO:0071453 | cellular response to oxygen levels | 0.002 | 0.006 | 0.003 | 3162/5468/857/5743/23411 | 5 |
| BP | GO:0030195 | negative regulation of blood coagulation | 0.002 | 0.006 | 0.003 | 5327/348/5054 | 3 |
| BP | GO:0043277 | apoptotic cell clearance | 0.002 | 0.006 | 0.003 | 6347/948/949 | 3 |
| BP | GO:0070509 | calcium ion import | 0.002 | 0.006 | 0.003 | 1636/6347/2641 | 3 |
| BP | GO:0010634 | positive regulation of epithelial cell migration | 0.002 | 0.007 | 0.004 | 3162/5743/23411/4318/949 | 5 |
| BP | GO:0042102 | positive regulation of T cell proliferation | 0.002 | 0.007 | 0.004 | 3553/3569/3952/3552 | 4 |
| BP | GO:0046545 | development of primary female sexual characteristics | 0.002 | 0.007 | 0.004 | 9510/23411/6647/3952 | 4 |
| BP | GO:0035265 | organ growth | 0.002 | 0.007 | 0.004 | 5573/6647/5465/3953/3952 | 5 |
| BP | GO:0001774 | microglial cell activation | 0.002 | 0.007 | 0.004 | 7124/3569/3949 | 3 |
| BP | GO:0046427 | positive regulation of receptor signaling pathway via JAK-STAT | 0.002 | 0.007 | 0.004 | 7124/3569/3952 | 3 |
| BP | GO:0046850 | regulation of bone remodeling | 0.002 | 0.007 | 0.004 | 3953/3569/3952 | 3 |
| BP | GO:1900047 | negative regulation of hemostasis | 0.002 | 0.007 | 0.004 | 5327/348/5054 | 3 |
| BP | GO:0006787 | porphyrin-containing compound catabolic process | 0.002 | 0.007 | 0.004 | 3162/4363 | 2 |
| BP | GO:0032306 | regulation of prostaglandin secretion | 0.002 | 0.007 | 0.004 | 3553/3552 | 2 |
| BP | GO:0032308 | positive regulation of prostaglandin secretion | 0.002 | 0.007 | 0.004 | 3553/3552 | 2 |
| BP | GO:0033015 | tetrapyrrole catabolic process | 0.002 | 0.007 | 0.004 | 3162/4363 | 2 |
| BP | GO:0033079 | immature T cell proliferation | 0.002 | 0.007 | 0.004 | 3553/3552 | 2 |
| BP | GO:0033080 | immature T cell proliferation in thymus | 0.002 | 0.007 | 0.004 | 3553/3552 | 2 |
| BP | GO:0034380 | high-density lipoprotein particle assembly | 0.002 | 0.007 | 0.004 | 348/335 | 2 |
| BP | GO:0044406 | adhesion of symbiont to host | 0.002 | 0.007 | 0.004 | 3383/949 | 2 |
| BP | GO:0045986 | negative regulation of smooth muscle contraction | 0.002 | 0.007 | 0.004 | 5743/6647 | 2 |
| BP | GO:0051918 | negative regulation of fibrinolysis | 0.002 | 0.007 | 0.004 | 5327/5054 | 2 |
| BP | GO:0060100 | positive regulation of phagocytosis, engulfment | 0.002 | 0.007 | 0.004 | 2150/948 | 2 |
| BP | GO:0072683 | T cell extravasation | 0.002 | 0.007 | 0.004 | 6347/3383 | 2 |
| BP | GO:1902950 | regulation of dendritic spine maintenance | 0.002 | 0.007 | 0.004 | 348/3630 | 2 |
| BP | GO:1905155 | positive regulation of membrane invagination | 0.002 | 0.007 | 0.004 | 2150/948 | 2 |
| BP | GO:1905907 | negative regulation of amyloid fibril formation | 0.002 | 0.007 | 0.004 | 348/3949 | 2 |
| BP | GO:0032963 | collagen metabolic process | 0.002 | 0.007 | 0.004 | 3066/3688/3569/4318 | 4 |
| BP | GO:0046425 | regulation of receptor signaling pathway via JAK-STAT | 0.002 | 0.007 | 0.004 | 7124/857/3569/3952 | 4 |
| BP | GO:0006693 | prostaglandin metabolic process | 0.002 | 0.007 | 0.004 | 3553/5743/23411 | 3 |
| BP | GO:0008206 | bile acid metabolic process | 0.002 | 0.007 | 0.004 | 23411/3952/4864 | 3 |
| BP | GO:0022602 | ovulation cycle process | 0.002 | 0.007 | 0.004 | 9510/23411/3952 | 3 |
| BP | GO:0006692 | prostanoid metabolic process | 0.002 | 0.007 | 0.004 | 3553/5743/23411 | 3 |
| BP | GO:0010332 | response to gamma radiation | 0.002 | 0.007 | 0.004 | 6648/2876/3552 | 3 |
| BP | GO:0032365 | intracellular lipid transport | 0.002 | 0.007 | 0.004 | 10577/4864/3949 | 3 |
| BP | GO:0046460 | neutral lipid biosynthetic process | 0.002 | 0.007 | 0.004 | 23411/949/3949 | 3 |
| BP | GO:0046463 | acylglycerol biosynthetic process | 0.002 | 0.007 | 0.004 | 23411/949/3949 | 3 |
| BP | GO:0051960 | regulation of nervous system development | 0.002 | 0.007 | 0.004 | 1636/3066/3688/7124/3553/7133/3569/3949 | 8 |
| BP | GO:0046822 | regulation of nucleocytoplasmic transport | 0.002 | 0.007 | 0.004 | 3553/5743/948/3952 | 4 |
| BP | GO:0090277 | positive regulation of peptide hormone secretion | 0.002 | 0.007 | 0.004 | 3630/5573/2641/8431 | 4 |
| BP | GO:0010867 | positive regulation of triglyceride biosynthetic process | 0.002 | 0.008 | 0.004 | 949/3949 | 2 |
| BP | GO:0032095 | regulation of response to food | 0.002 | 0.008 | 0.004 | 5465/3952 | 2 |
| BP | GO:0033700 | phospholipid efflux | 0.002 | 0.008 | 0.004 | 348/335 | 2 |
| BP | GO:0034116 | positive regulation of heterotypic cell-cell adhesion | 0.002 | 0.008 | 0.004 | 7124/3553 | 2 |
| BP | GO:0046479 | glycosphingolipid catabolic process | 0.002 | 0.008 | 0.004 | 2717/4758 | 2 |
| BP | GO:0061043 | regulation of vascular wound healing | 0.002 | 0.008 | 0.004 | 7124/5054 | 2 |
| BP | GO:1901031 | regulation of response to reactive oxygen species | 0.002 | 0.008 | 0.004 | 10516/948 | 2 |
| BP | GO:0002699 | positive regulation of immune effector process | 0.002 | 0.008 | 0.004 | 2150/7124/3553/23411/948/3569 | 6 |
| BP | GO:0002269 | leukocyte activation involved in inflammatory response | 0.002 | 0.008 | 0.004 | 7124/3569/3949 | 3 |
| BP | GO:0006636 | unsaturated fatty acid biosynthetic process | 0.002 | 0.008 | 0.004 | 3553/5743/23411 | 3 |
| BP | GO:0043154 | egulation of cysteine-type endopeptidase activity involved in apoptot | 0.002 | 0.008 | 0.004 | 2876/7124/4318 | 3 |
| BP | GO:0050819 | negative regulation of coagulation | 0.002 | 0.008 | 0.004 | 5327/348/5054 | 3 |
| BP | GO:0051180 | vitamin transport | 0.002 | 0.008 | 0.004 | 335/949/4363 | 3 |
| BP | GO:0002793 | positive regulation of peptide secretion | 0.002 | 0.008 | 0.004 | 3630/5573/2641/8431 | 4 |
| BP | GO:0030512 | ve regulation of transforming growth factor beta receptor signaling pa | 0.002 | 0.008 | 0.004 | 3066/5468/23411/5465 | 4 |

| | | | | | | | |
|---|---|---|---|---|---|---|---|
| BP | GO:0031341 | regulation of cell killing | 0.002 | 0.008 | 0.004 | 2150/2671/3383/3952 | 4 |
| BP | GO:0032872 | regulation of stress-activated MAPK cascade | 0.002 | 0.008 | 0.004 | 2150/7124/3553/3952/3552 | 5 |
| BP | GO:1904894 | positive regulation of receptor signaling pathway via STAT | 0.002 | 0.008 | 0.004 | 7124/3569/3952 | 3 |
| BP | GO:0032355 | response to estradiol | 0.002 | 0.008 | 0.004 | 2876/5743/4968/3952 | 4 |
| BP | GO:0009395 | phospholipid catabolic process | 0.002 | 0.008 | 0.004 | 3990/949/3949 | 3 |
| BP | GO:0042743 | hydrogen peroxide metabolic process | 0.002 | 0.008 | 0.004 | 6648/2876/6647 | 3 |
| BP | GO:0050805 | negative regulation of synaptic transmission | 0.002 | 0.008 | 0.004 | 3553/5743/9370 | 3 |
| BP | GO:0070302 | regulation of stress-activated protein kinase signaling cascade | 0.002 | 0.008 | 0.004 | 2150/7124/3553/3952/3552 | 5 |
| BP | GO:0035358 | gulation of peroxisome proliferator activated receptor signaling pathw | 0.002 | 0.009 | 0.005 | 23411/3952 | 2 |
| BP | GO:0042159 | lipoprotein catabolic process | 0.002 | 0.009 | 0.005 | 348/3949 | 2 |
| BP | GO:0060099 | regulation of phagocytosis, engulfment | 0.002 | 0.009 | 0.005 | 2150/948 | 2 |
| BP | GO:0071639 | positive regulation of monocyte chemotactic protein-1 production | 0.002 | 0.009 | 0.005 | 3553/9370 | 2 |
| BP | GO:0090205 | positive regulation of cholesterol metabolic process | 0.002 | 0.009 | 0.005 | 348/335 | 2 |
| BP | GO:0090209 | negative regulation of triglyceride metabolic process | 0.002 | 0.009 | 0.005 | 348/23411 | 2 |
| BP | GO:0090280 | positive regulation of calcium ion import | 0.002 | 0.009 | 0.005 | 6347/2641 | 2 |
| BP | GO:1900102 | negative regulation of endoplasmic reticulum unfolded protein respons | 0.002 | 0.009 | 0.005 | 5770/7466 | 2 |
| BP | GO:1904294 | positive regulation of ERAD pathway | 0.002 | 0.009 | 0.005 | 857/7466 | 2 |
| BP | GO:1905153 | regulation of membrane invagination | 0.002 | 0.009 | 0.005 | 2150/948 | 2 |
| BP | GO:0006644 | phospholipid metabolic process | 0.002 | 0.009 | 0.005 | 4023/335/1071/3990/2784/949/3949 | 7 |
| BP | GO:0033559 | unsaturated fatty acid metabolic process | 0.002 | 0.009 | 0.005 | 2876/3553/5743/23411 | 4 |
| BP | GO:0043473 | pigmentation | 0.002 | 0.009 | 0.005 | 6648/348/5443/5167 | 4 |
| BP | GO:0042304 | regulation of fatty acid biosynthetic process | 0.002 | 0.009 | 0.005 | 3553/5743/23411 | 3 |
| BP | GO:0002822 | nse based on somatic recombination of immune receptors built from i | 0.003 | 0.009 | 0.005 | 7124/3553/7133/196/3569 | 5 |
| BP | GO:0001836 | release of cytochrome c from mitochondria | 0.003 | 0.009 | 0.005 | 6648/2876/4318 | 3 |
| BP | GO:0030857 | negative regulation of epithelial cell differentiation | 0.003 | 0.009 | 0.005 | 857/4318/3552 | 3 |
| BP | GO:0061097 | regulation of protein tyrosine kinase activity | 0.003 | 0.009 | 0.005 | 1636/857/5770 | 3 |
| BP | GO:0007088 | regulation of mitotic nuclear division | 0.003 | 0.009 | 0.005 | 3630/7124/3553/3552 | 4 |
| BP | GO:2000116 | regulation of cysteine-type endopeptidase activity | 0.003 | 0.009 | 0.005 | 2876/5468/7124/23411/4318 | 5 |
| BP | GO:0010958 | regulation of amino acid import across plasma membrane | 0.003 | 0.009 | 0.005 | 3688/7124 | 2 |
| BP | GO:0034393 | positive regulation of smooth muscle cell apoptotic process | 0.003 | 0.009 | 0.005 | 6648/5468 | 2 |
| BP | GO:0045019 | negative regulation of nitric oxide biosynthetic process | 0.003 | 0.009 | 0.005 | 2717/857 | 2 |
| BP | GO:0050665 | hydrogen peroxide biosynthetic process | 0.003 | 0.009 | 0.005 | 6648/6647 | 2 |
| BP | GO:0051956 | negative regulation of amino acid transport | 0.003 | 0.009 | 0.005 | 7124/3952 | 2 |
| BP | GO:0055091 | phospholipid homeostasis | 0.003 | 0.009 | 0.005 | 335/1071 | 2 |
| BP | GO:0090336 | positive regulation of brown fat cell differentiation | 0.003 | 0.009 | 0.005 | 3630/5743 | 2 |
| BP | GO:1903789 | regulation of amino acid transmembrane transport | 0.003 | 0.009 | 0.005 | 3688/7124 | 2 |
| BP | GO:1904406 | negative regulation of nitric oxide metabolic process | 0.003 | 0.009 | 0.005 | 2717/857 | 2 |
| BP | GO:1901016 | regulation of potassium ion transmembrane transporter activity | 0.003 | 0.01 | 0.005 | 3688/857/481 | 3 |
| BP | GO:1901616 | organic hydroxy compound catabolic process | 0.003 | 0.01 | 0.005 | 348/1312/949 | 3 |
| BP | GO:0032609 | type II interferon production | 0.003 | 0.01 | 0.005 | 7356/2150/7124/3553 | 4 |
| BP | GO:0032649 | regulation of type II interferon production | 0.003 | 0.01 | 0.005 | 7356/2150/7124/3553 | 4 |
| BP | GO:0010721 | negative regulation of cell development | 0.003 | 0.01 | 0.005 | 7124/3553/9370/3569/3552/3949 | 6 |
| BP | GO:0071560 | cellular response to transforming growth factor beta stimulus | 0.003 | 0.01 | 0.005 | 9518/3066/5468/857/23411/5465 | 6 |
| BP | GO:0046660 | female sex differentiation | 0.003 | 0.01 | 0.005 | 9510/23411/6647/3952 | 4 |
| BP | GO:0006749 | glutathione metabolic process | 0.003 | 0.01 | 0.005 | 6648/2876/6647 | 3 |
| BP | GO:0042306 | regulation of protein import into nucleus | 0.003 | 0.01 | 0.005 | 5743/948/3952 | 3 |
| BP | GO:0048008 | platelet-derived growth factor receptor signaling pathway | 0.003 | 0.01 | 0.005 | 5327/5770/9370 | 3 |
| BP | GO:0086065 | cell communication involved in cardiac conduction | 0.003 | 0.01 | 0.005 | 857/5318/481 | 3 |
| BP | GO:0007548 | sex differentiation | 0.003 | 0.01 | 0.005 | 1636/9510/23411/6647/3952/3552 | 6 |
| BP | GO:0046777 | protein autophosphorylation | 0.003 | 0.01 | 0.005 | 1636/3630/857/9370/5167 | 5 |
| BP | GO:0071248 | cellular response to metal ion | 0.003 | 0.01 | 0.005 | 3162/5743/7351/4968/4318 | 5 |
| BP | GO:0010718 | positive regulation of epithelial to mesenchymal transition | 0.003 | 0.01 | 0.006 | 3066/3553/3569 | 3 |
| BP | GO:0032388 | positive regulation of intracellular transport | 0.003 | 0.01 | 0.006 | 5116/3553/208/5743/3952 | 5 |
| BP | GO:0001991 | ılation of systemic arterial blood pressure by circulatory renin-angiote | 0.003 | 0.01 | 0.006 | 1636/2150 | 2 |
| BP | GO:0019377 | glycolipid catabolic process | 0.003 | 0.01 | 0.006 | 2717/4758 | 2 |
| BP | GO:0032488 | Cdc42 protein signal transduction | 0.003 | 0.01 | 0.006 | 348/335 | 2 |
| BP | GO:0036166 | phenotypic switching | 0.003 | 0.01 | 0.006 | 6648/1786 | 2 |
| BP | GO:0044539 | long-chain fatty acid import into cell | 0.003 | 0.01 | 0.006 | 208/948 | 2 |
| BP | GO:0050872 | white fat cell differentiation | 0.003 | 0.01 | 0.006 | 5468/23411 | 2 |
| BP | GO:0098712 | L-glutamate import across plasma membrane | 0.003 | 0.01 | 0.006 | 3688/7124 | 2 |
| BP | GO:1900120 | regulation of receptor binding | 0.003 | 0.01 | 0.006 | 9370/4318 | 2 |
| BP | GO:1901550 | regulation of endothelial cell development | 0.003 | 0.01 | 0.006 | 7124/3553 | 2 |
| BP | GO:1903140 | regulation of establishment of endothelial barrier | 0.003 | 0.01 | 0.006 | 7124/3553 | 2 |
| BP | GO:1905906 | regulation of amyloid fibril formation | 0.003 | 0.01 | 0.006 | 348/3949 | 2 |
| BP | GO:0071559 | response to transforming growth factor beta | 0.003 | 0.011 | 0.006 | 9518/3066/5468/857/23411/5465 | 6 |
| BP | GO:0010803 | regulation of tumor necrosis factor-mediated signaling pathway | 0.003 | 0.011 | 0.006 | 2150/335/9370 | 3 |
| BP | GO:0018105 | peptidyl-serine phosphorylation | 0.003 | 0.011 | 0.006 | 7124/208/857/5743/2641/3569 | 6 |
| BP | GO:0006940 | regulation of smooth muscle contraction | 0.003 | 0.011 | 0.006 | 857/5743/6647 | 3 |
| BP | GO:0046456 | icosanoid biosynthetic process | 0.003 | 0.011 | 0.006 | 3553/5743/23411 | 3 |
| BP | GO:0022408 | negative regulation of cell-cell adhesion | 0.003 | 0.011 | 0.006 | 7356/335/5573/9370/5465 | 5 |
| BP | GO:0050777 | negative regulation of immune response | 0.003 | 0.011 | 0.006 | 2876/5468/3630/2671/196 | 5 |
| BP | GO:0002688 | regulation of leukocyte chemotaxis | 0.003 | 0.011 | 0.006 | 2150/6347/3569/5054 | 4 |
| BP | GO:0043200 | response to amino acid | 0.003 | 0.011 | 0.006 | 1786/1291/7124/26291 | 4 |
| BP | GO:0010042 | response to manganese ion | 0.003 | 0.011 | 0.006 | 6648/5743 | 2 |
| BP | GO:0010755 | regulation of plasminogen activation | 0.003 | 0.011 | 0.006 | 5327/5054 | 2 |
| BP | GO:0033194 | response to hydroperoxide | 0.003 | 0.011 | 0.006 | 2876/948 | 2 |
| BP | GO:0038083 | peptidyl-tyrosine autophosphorylation | 0.003 | 0.011 | 0.006 | 1636/857 | 2 |
| BP | GO:0043217 | myelin maintenance | 0.003 | 0.011 | 0.006 | 208/6647 | 2 |
| BP | GO:0051917 | regulation of fibrinolysis | 0.003 | 0.011 | 0.006 | 5327/5054 | 2 |
| BP | GO:0150079 | negative regulation of neuroinflammatory response | 0.003 | 0.011 | 0.006 | 7133/3949 | 2 |
| BP | GO:2000811 | negative regulation of anoikis | 0.003 | 0.011 | 0.006 | 3688/857 | 2 |
| BP | GO:0071674 | mononuclear cell migration | 0.003 | 0.012 | 0.006 | 7124/6347/3383/3569/5054 | 5 |
| BP | GO:0046824 | positive regulation of nucleocytoplasmic transport | 0.003 | 0.012 | 0.006 | 3553/5743/3952 | 3 |
| BP | GO:0008016 | regulation of heart contraction | 0.004 | 0.012 | 0.006 | 7124/857/5318/153/481 | 5 |
| BP | GO:0031330 | negative regulation of cellular catabolic process | 0.004 | 0.012 | 0.006 | 3162/3630/3953/3952/4864 | 5 |
| BP | GO:0001704 | formation of primary germ layer | 0.004 | 0.012 | 0.006 | 3688/1291/5573/4318 | 4 |
| BP | GO:0002821 | positive regulation of adaptive immune response | 0.004 | 0.013 | 0.007 | 7124/3553/23411/3569 | 4 |
| BP | GO:0010595 | positive regulation of endothelial cell migration | 0.004 | 0.013 | 0.007 | 3162/5743/23411/949 | 4 |
| BP | GO:0062208 | positive regulation of pattern recognition receptor signaling pathway | 0.004 | 0.013 | 0.007 | 2150/857/948 | 3 |
| BP | GO:2001244 | positive regulation of intrinsic apoptotic signaling pathway | 0.004 | 0.013 | 0.007 | 857/23411/6647 | 3 |
| BP | GO:0048608 | reproductive structure development | 0.004 | 0.013 | 0.007 | 1636/9510/23411/6647/3952/3552 | 6 |
| BP | GO:0051146 | striated muscle cell differentiation | 0.004 | 0.013 | 0.007 | 9518/2876/3688/34/5573/5465 | 6 |
| BP | GO:0098739 | import across plasma membrane | 0.004 | 0.013 | 0.007 | 3688/7124/208/948/481 | 5 |
| BP | GO:0010744 | positive regulation of macrophage derived foam cell differentiation | 0.004 | 0.013 | 0.007 | 4023/948 | 2 |
| BP | GO:0032305 | positive regulation of icosanoid secretion | 0.004 | 0.013 | 0.007 | 3553/3552 | 2 |
| BP | GO:0035743 | CD4-positive, alpha-beta T cell cytokine production | 0.004 | 0.013 | 0.007 | 3553/3569 | 2 |
| BP | GO:0035821 | modulation of process of another organism | 0.004 | 0.013 | 0.007 | 2150/5465 | 2 |
| BP | GO:0045346 | regulation of MHC class II biosynthetic process | 0.004 | 0.013 | 0.007 | 3066/23411 | 2 |
| BP | GO:1902931 | negative regulation of alcohol biosynthetic process | 0.004 | 0.013 | 0.007 | 348/6647 | 2 |
| BP | GO:1903427 | negative regulation of reactive oxygen species biosynthetic process | 0.004 | 0.013 | 0.007 | 3630/5465 | 2 |
| BP | GO:1903428 | positive regulation of reactive oxygen species biosynthetic process | 0.004 | 0.013 | 0.007 | 6648/948 | 2 |
| BP | GO:0030217 | T cell differentiation | 0.004 | 0.013 | 0.007 | 3553/6647/3953/3569/3952/3552 | 6 |
| BP | GO:0043903 | regulation of biological process involved in symbiotic interaction | 0.004 | 0.013 | 0.007 | 2150/857/5465 | 3 |
| BP | GO:0070059 | sic apoptotic signaling pathway in response to endoplasmic reticulum | 0.004 | 0.013 | 0.007 | 5770/23411/7466 | 3 |
| BP | GO:2000117 | negative regulation of cysteine-type endopeptidase activity | 0.004 | 0.013 | 0.007 | 2876/7124/4318 | 3 |
| BP | GO:0018209 | peptidyl-serine modification | 0.004 | 0.013 | 0.007 | 7124/208/857/5743/2641/3569 | 6 |
| BP | GO:0046328 | regulation of JNK cascade | 0.004 | 0.013 | 0.007 | 2150/7124/3553/3552 | 4 |
| BP | GO:0061458 | reproductive system development | 0.004 | 0.013 | 0.007 | 1636/9510/23411/6647/3952/3552 | 6 |
| BP | GO:0043271 | negative regulation of monoatomic ion transport | 0.004 | 0.013 | 0.007 | 1636/857/5743/4318 | 4 |

| | | | | | | | |
|---|---|---|---|---|---|---|---|
| BP | GO:0032303 | regulation of icosanoid secretion | 0.004 | 0.014 | 0.007 | 3553/3552 | 2 |
| BP | GO:0042953 | lipoprotein transport | 0.004 | 0.014 | 0.007 | 5468/948 | 2 |
| BP | GO:0045342 | MHC class II biosynthetic process | 0.004 | 0.014 | 0.007 | 3066/23411 | 2 |
| BP | GO:0097709 | connective tissue replacement | 0.004 | 0.014 | 0.007 | 5468/3552 | 2 |
| BP | GO:0140354 | lipid import into cell | 0.004 | 0.014 | 0.007 | 208/948 | 2 |
| BP | GO:1900221 | regulation of amyloid-beta clearance | 0.004 | 0.014 | 0.007 | 348/7124 | 2 |
| BP | GO:1902001 | fatty acid transmembrane transport | 0.004 | 0.014 | 0.007 | 208/948 | 2 |
| BP | GO:1902236 | ation of endoplasmic reticulum stress-induced intrinsic apoptotic sign | 0.004 | 0.014 | 0.007 | 5770/7466 | 2 |
| BP | GO:1903978 | regulation of microglial cell activation | 0.004 | 0.014 | 0.007 | 3569/3949 | 2 |
| BP | GO:0007162 | negative regulation of cell adhesion | 0.004 | 0.014 | 0.007 | 7356/335/5573/9370/5465/5054 | 6 |
| BP | GO:0042594 | response to starvation | 0.004 | 0.014 | 0.007 | 1312/34/23411/7351/5465 | 5 |
| BP | GO:0051346 | negative regulation of hydrolase activity | 0.004 | 0.014 | 0.007 | 2876/7124/5265/5054/4318 | 5 |
| BP | GO:0010611 | regulation of cardiac muscle hypertrophy | 0.004 | 0.014 | 0.007 | 5468/7133/5465 | 3 |
| BP | GO:0038034 | signal transduction in absence of ligand | 0.004 | 0.014 | 0.007 | 7124/3553/3552 | 3 |
| BP | GO:0070301 | cellular response to hydrogen peroxide | 0.004 | 0.014 | 0.007 | 3066/23411/3569 | 3 |
| BP | GO:0071300 | cellular response to retinoic acid | 0.004 | 0.014 | 0.007 | 3066/7124/3952 | 3 |
| BP | GO:0097192 | extrinsic apoptotic signaling pathway in absence of ligand | 0.004 | 0.014 | 0.007 | 7124/3553/3552 | 3 |
| BP | GO:0051055 | negative regulation of lipid biosynthetic process | 0.004 | 0.014 | 0.008 | 348/23411/6647 | 3 |
| BP | GO:0007266 | Rho protein signal transduction | 0.005 | 0.015 | 0.008 | 2150/348/3688/335 | 4 |
| BP | GO:0009416 | response to light stimulus | 0.005 | 0.015 | 0.008 | 2876/3688/208/23411/4968/4318 | 6 |
| BP | GO:0010288 | response to lead ion | 0.005 | 0.015 | 0.008 | 5743/7351 | 2 |
| BP | GO:0034104 | negative regulation of tissue remodeling | 0.005 | 0.015 | 0.008 | 5468/3569 | 2 |
| BP | GO:0044872 | lipoprotein localization | 0.005 | 0.015 | 0.008 | 5468/948 | 2 |
| BP | GO:0046514 | ceramide catabolic process | 0.005 | 0.015 | 0.008 | 2717/4758 | 2 |
| BP | GO:0051152 | positive regulation of smooth muscle cell differentiation | 0.005 | 0.015 | 0.008 | 6648/23411 | 2 |
| BP | GO:0071605 | monocyte chemotactic protein-1 production | 0.005 | 0.015 | 0.008 | 3553/9370 | 2 |
| BP | GO:0071637 | regulation of monocyte chemotactic protein-1 production | 0.005 | 0.015 | 0.008 | 3553/9370 | 2 |
| BP | GO:1904292 | regulation of ERAD pathway | 0.005 | 0.015 | 0.008 | 857/7466 | 2 |
| BP | GO:0010212 | response to ionizing radiation | 0.005 | 0.015 | 0.008 | 6648/2876/23411/3552 | 4 |
| BP | GO:0009311 | oligosaccharide metabolic process | 0.005 | 0.015 | 0.008 | 2717/4758/4864 | 3 |
| BP | GO:0014743 | regulation of muscle hypertrophy | 0.005 | 0.015 | 0.008 | 5468/7133/5465 | 3 |
| BP | GO:1902893 | regulation of miRNA transcription | 0.005 | 0.015 | 0.008 | 5468/7124/5465 | 3 |
| BP | GO:0001909 | leukocyte mediated cytotoxicity | 0.005 | 0.015 | 0.008 | 2150/2671/3383/3952 | 4 |
| BP | GO:0070555 | response to interleukin-1 | 0.005 | 0.015 | 0.008 | 166/3553/6347/3569 | 4 |
| BP | GO:0072330 | monocarboxylic acid biosynthetic process | 0.005 | 0.016 | 0.008 | 4023/3553/3990/5743/23411 | 5 |
| BP | GO:0030098 | lymphocyte differentiation | 0.005 | 0.016 | 0.008 | 3688/3553/6647/3953/3569/3952/3552 | 7 |
| BP | GO:0001101 | response to acid chemical | 0.005 | 0.016 | 0.008 | 1786/1291/7124/26291 | 4 |
| BP | GO:0002548 | monocyte chemotaxis | 0.005 | 0.016 | 0.008 | 6347/3569/5054 | 3 |
| BP | GO:0042698 | ovulation cycle | 0.005 | 0.016 | 0.008 | 9510/23411/3952 | 3 |
| BP | GO:0061614 | miRNA transcription | 0.005 | 0.016 | 0.008 | 5468/7124/5465 | 3 |
| BP | GO:1902475 | L-alpha-amino acid transmembrane transport | 0.005 | 0.016 | 0.008 | 3688/7124/7351 | 3 |
| BP | GO:0030099 | myeloid cell differentiation | 0.005 | 0.016 | 0.008 | 2150/5468/7124/23411/9370/7351/4318 | 7 |
| BP | GO:0006883 | intracellular sodium ion homeostasis | 0.005 | 0.016 | 0.008 | 3552/481 | 2 |
| BP | GO:0010988 | regulation of low-density lipoprotein particle clearance | 0.005 | 0.016 | 0.008 | 26291/3949 | 2 |
| BP | GO:0097062 | dendritic spine maintenance | 0.005 | 0.016 | 0.008 | 348/3630 | 2 |
| BP | GO:0098801 | regulation of renal system process | 0.005 | 0.016 | 0.008 | 2150/9370 | 2 |
| BP | GO:0060326 | cell chemotaxis | 0.005 | 0.016 | 0.009 | 2150/3553/6347/3569/5054/4363 | 6 |
| BP | GO:0060537 | muscle tissue development | 0.005 | 0.016 | 0.009 | 2876/3688/34/857/5318/5573/5465 | 7 |
| BP | GO:0043954 | cellular component maintenance | 0.005 | 0.016 | 0.009 | 2150/348/3630 | 3 |
| BP | GO:0007179 | transforming growth factor beta receptor signaling pathway | 0.005 | 0.016 | 0.009 | 9518/3066/5468/23411/5465 | 5 |
| BP | GO:0071241 | cellular response to inorganic substance | 0.005 | 0.016 | 0.009 | 3162/5743/7351/4968/4318 | 5 |
| BP | GO:1900180 | regulation of protein localization to nucleus | 0.005 | 0.016 | 0.009 | 3630/5743/948/3952 | 4 |
| BP | GO:0051403 | stress-activated MAPK cascade | 0.005 | 0.017 | 0.009 | 2150/7124/3553/3952/3552 | 5 |
| BP | GO:0043112 | receptor metabolic process | 0.005 | 0.017 | 0.009 | 348/5770/3949 | 3 |
| BP | GO:0140962 | multicellular organismal-level chemical homeostasis | 0.005 | 0.017 | 0.009 | 3162/6648/7466 | 3 |
| BP | GO:0032412 | regulation of monoatomic ion transmembrane transporter activity | 0.005 | 0.017 | 0.009 | 3688/857/6347/4318/481 | 5 |
| BP | GO:0051251 | positive regulation of lymphocyte activation | 0.006 | 0.017 | 0.009 | 3553/857/6347/3569/3952/3552 | 6 |
| BP | GO:0030534 | adult behavior | 0.006 | 0.017 | 0.009 | 3066/5465/3952/4864 | 4 |
| BP | GO:0042552 | myelination | 0.006 | 0.017 | 0.009 | 7124/208/6647/7133 | 4 |
| BP | GO:0050671 | positive regulation of lymphocyte proliferation | 0.006 | 0.017 | 0.009 | 3553/3569/3952/3552 | 4 |
| BP | GO:2001235 | positive regulation of apoptotic signaling pathway | 0.006 | 0.017 | 0.009 | 7124/857/23411/6647 | 4 |
| BP | GO:0010893 | positive regulation of steroid biosynthetic process | 0.006 | 0.017 | 0.009 | 7124/3552 | 2 |
| BP | GO:0016575 | histone deacetylation | 0.006 | 0.017 | 0.009 | 3066/23411 | 2 |
| BP | GO:0043171 | peptide catabolic process | 0.006 | 0.017 | 0.009 | 1636/3416 | 2 |
| BP | GO:0045932 | negative regulation of muscle contraction | 0.006 | 0.017 | 0.009 | 5743/6647 | 2 |
| BP | GO:0061042 | vascular wound healing | 0.006 | 0.017 | 0.009 | 7124/5054 | 2 |
| BP | GO:0072574 | hepatocyte proliferation | 0.006 | 0.017 | 0.009 | 7124/3569 | 2 |
| BP | GO:0072575 | epithelial cell proliferation involved in liver morphogenesis | 0.006 | 0.017 | 0.009 | 7124/3569 | 2 |
| BP | GO:2000193 | positive regulation of fatty acid transport | 0.006 | 0.017 | 0.009 | 3553/3552 | 2 |
| BP | GO:0002753 | cytosolic pattern recognition receptor signaling pathway | 0.006 | 0.017 | 0.009 | 2150/7124/857/948 | 4 |
| BP | GO:0051783 | regulation of nuclear division | 0.006 | 0.017 | 0.009 | 3630/7124/3553/3552 | 4 |
| BP | GO:0007272 | ensheathment of neurons | 0.006 | 0.018 | 0.009 | 7124/208/6647/7133 | 4 |
| BP | GO:0008366 | axon ensheathment | 0.006 | 0.018 | 0.009 | 7124/208/6647/7133 | 4 |
| BP | GO:0035264 | multicellular organism growth | 0.006 | 0.018 | 0.009 | 9518/6647/79068/153 | 4 |
| BP | GO:0071456 | cellular response to hypoxia | 0.006 | 0.018 | 0.009 | 3162/5468/5743/23411 | 4 |
| BP | GO:0030968 | endoplasmic reticulum unfolded protein response | 0.006 | 0.018 | 0.009 | 5770/26291/7466 | 3 |
| BP | GO:0032946 | positive regulation of mononuclear cell proliferation | 0.006 | 0.018 | 0.01 | 3553/3569/3952/3552 | 4 |
| BP | GO:0001503 | ossification | 0.006 | 0.018 | 0.01 | 5468/1291/7124/5743/3569/3952/5167 | 7 |
| BP | GO:0002922 | positive regulation of humoral immune response | 0.006 | 0.018 | 0.01 | 7124/3553 | 2 |
| BP | GO:0003081 | regulation of systemic arterial blood pressure by renin-angiotensin | 0.006 | 0.018 | 0.01 | 1636/2150 | 2 |
| BP | GO:0006925 | inflammatory cell apoptotic process | 0.006 | 0.018 | 0.01 | 23411/3569 | 2 |
| BP | GO:0045723 | positive regulation of fatty acid biosynthetic process | 0.006 | 0.018 | 0.01 | 3553/5743 | 2 |
| BP | GO:0046716 | muscle cell cellular homeostasis | 0.006 | 0.018 | 0.01 | 857/6647 | 2 |
| BP | GO:0060907 | positive regulation of macrophage cytokine production | 0.006 | 0.018 | 0.01 | 23411/948 | 2 |
| BP | GO:0006816 | calcium ion transport | 0.006 | 0.018 | 0.01 | 1636/857/5743/6347/2641/7466/481 | 7 |
| BP | GO:0050768 | negative regulation of neurogenesis | 0.006 | 0.018 | 0.01 | 7124/3553/3569/3949 | 4 |
| BP | GO:0043536 | positive regulation of blood vessel endothelial cell migration | 0.006 | 0.018 | 0.01 | 3162/5743/23411 | 3 |
| BP | GO:0051145 | smooth muscle cell differentiation | 0.006 | 0.018 | 0.01 | 6648/1786/23411 | 3 |
| BP | GO:0086001 | cardiac muscle cell action potential | 0.006 | 0.018 | 0.01 | 857/5318/481 | 3 |
| BP | GO:2000242 | negative regulation of reproductive process | 0.006 | 0.018 | 0.01 | 5327/6647/3552 | 3 |
| BP | GO:0031098 | stress-activated protein kinase signaling cascade | 0.006 | 0.018 | 0.01 | 2150/7124/3553/3952/3552 | 5 |
| BP | GO:0006865 | amino acid transport | 0.006 | 0.019 | 0.01 | 3688/7124/7351/3952 | 4 |
| BP | GO:0016525 | negative regulation of angiogenesis | 0.006 | 0.019 | 0.01 | 5468/7124/9510/5054 | 4 |
| BP | GO:0032091 | negative regulation of protein binding | 0.006 | 0.019 | 0.01 | 166/857/9370 | 3 |
| BP | GO:0032729 | positive regulation of type II interferon production | 0.006 | 0.019 | 0.01 | 2150/7124/3553 | 3 |
| BP | GO:0001890 | placenta development | 0.006 | 0.019 | 0.01 | 5468/5743/6647/3952 | 4 |
| BP | GO:0002705 | positive regulation of leukocyte mediated immunity | 0.006 | 0.019 | 0.01 | 2150/7124/3553/3569 | 4 |
| BP | GO:0030595 | leukocyte chemotaxis | 0.006 | 0.019 | 0.01 | 2150/3553/6347/3569/5054 | 5 |
| BP | GO:2000181 | negative regulation of blood vessel morphogenesis | 0.007 | 0.019 | 0.01 | 5468/7124/9510/5054 | 4 |
| BP | GO:0055117 | regulation of cardiac muscle contraction | 0.007 | 0.019 | 0.01 | 857/5318/481 | 3 |
| BP | GO:0045540 | regulation of cholesterol biosynthetic process | 0.007 | 0.019 | 0.01 | 348/6647 | 2 |
| BP | GO:0060396 | growth hormone receptor signaling pathway | 0.007 | 0.019 | 0.01 | 9518/5770 | 2 |
| BP | GO:0060586 | multicellular organismal-level iron ion homeostasis | 0.007 | 0.019 | 0.01 | 3162/6648 | 2 |
| BP | GO:0072576 | liver morphogenesis | 0.007 | 0.019 | 0.01 | 7124/3569 | 2 |
| BP | GO:0106118 | regulation of sterol biosynthetic process | 0.007 | 0.019 | 0.01 | 348/6647 | 2 |
| BP | GO:1902894 | negative regulation of miRNA transcription | 0.007 | 0.019 | 0.01 | 5468/5465 | 2 |
| BP | GO:1905063 | regulation of vascular associated smooth muscle cell differentiation | 0.007 | 0.019 | 0.01 | 6648/1786 | 2 |
| BP | GO:1905563 | negative regulation of vascular endothelial cell proliferation | 0.007 | 0.019 | 0.01 | 5468/6347 | 2 |
| BP | GO:0097529 | myeloid leukocyte migration | 0.007 | 0.019 | 0.01 | 3553/6347/3569/5054/3552 | 5 |

| | | | | | | | |
|---|---|---|---|---|---|---|---|
| BP | GO:1901343 | negative regulation of vasculature development | 0.007 | 0.02 | 0.01 | 5468/7124/9510/5054 | 4 |
| BP | GO:0051966 | regulation of synaptic transmission, glutamatergic | 0.007 | 0.02 | 0.011 | 7124/5743/6347 | 3 |
| BP | GO:0051961 | negative regulation of nervous system development | 0.007 | 0.02 | 0.011 | 7124/3553/3569/3949 | 4 |
| BP | GO:0043537 | negative regulation of blood vessel endothelial cell migration | 0.007 | 0.021 | 0.011 | 5468/348/7124 | 3 |
| BP | GO:0002026 | regulation of the force of heart contraction | 0.007 | 0.021 | 0.011 | 857/153 | 2 |
| BP | GO:0002726 | positive regulation of T cell cytokine production | 0.007 | 0.021 | 0.011 | 3553/3569 | 2 |
| BP | GO:0031579 | membrane raft organization | 0.007 | 0.021 | 0.011 | 857/4864 | 2 |
| BP | GO:0031954 | positive regulation of protein autophosphorylation | 0.007 | 0.021 | 0.011 | 1636/3630 | 2 |
| BP | GO:0060259 | regulation of feeding behavior | 0.007 | 0.021 | 0.011 | 3630/3953 | 2 |
| BP | GO:0071378 | cellular response to growth hormone stimulus | 0.007 | 0.021 | 0.011 | 9518/5770 | 2 |
| BP | GO:1900407 | regulation of cellular response to oxidative stress | 0.007 | 0.021 | 0.011 | 10516/948 | 2 |
| BP | GO:1904996 | positive regulation of leukocyte adhesion to vascular endothelial cell | 0.007 | 0.021 | 0.011 | 7124/3569 | 2 |
| BP | GO:2000629 | negative regulation of miRNA metabolic process | 0.007 | 0.021 | 0.011 | 5468/5465 | 2 |
| BP | GO:0048738 | cardiac muscle tissue development | 0.007 | 0.021 | 0.011 | 3688/34/5318/5573/5465 | 5 |
| BP | GO:0006936 | muscle contraction | 0.007 | 0.021 | 0.011 | 7124/857/5318/5743/6647/481 | 6 |
| BP | GO:0007189 | enylate cyclase-activating G protein-coupled receptor signaling pathw | 0.007 | 0.022 | 0.011 | 2778/5573/2641/153 | 4 |
| BP | GO:0036294 | cellular response to decreased oxygen levels | 0.007 | 0.022 | 0.011 | 3162/5468/5743/23411 | 4 |
| BP | GO:0031639 | plasminogen activation | 0.008 | 0.022 | 0.012 | 5327/5054 | 2 |
| BP | GO:0034377 | plasma lipoprotein particle assembly | 0.008 | 0.022 | 0.012 | 348/335 | 2 |
| BP | GO:0042730 | fibrinolysis | 0.008 | 0.022 | 0.012 | 5327/5054 | 2 |
| BP | GO:0048714 | positive regulation of oligodendrocyte differentiation | 0.008 | 0.022 | 0.012 | 3066/7133 | 2 |
| BP | GO:1901875 | positive regulation of post-translational protein modification | 0.008 | 0.022 | 0.012 | 857/23411/2641/7466 | 4 |
| BP | GO:0017015 | gulation of transforming growth factor beta receptor signaling pathwa | 0.008 | 0.022 | 0.012 | 3066/5468/23411/5465 | 4 |
| BP | GO:0006935 | chemotaxis | 0.008 | 0.023 | 0.012 | 2150/3553/335/6347/3569/5054/4363 | 7 |
| BP | GO:0031589 | cell-substrate adhesion | 0.008 | 0.023 | 0.012 | 3688/10516/335/5318/948/5054 | 6 |
| BP | GO:0006606 | protein import into nucleus | 0.008 | 0.023 | 0.012 | 4928/5743/948/3952 | 4 |
| BP | GO:0090287 | regulation of cellular response to growth factor stimulus | 0.008 | 0.023 | 0.012 | 3066/5468/3553/5770/23411/5465 | 6 |
| BP | GO:0032024 | positive regulation of insulin secretion | 0.008 | 0.023 | 0.012 | 5573/2641/8431 | 3 |
| BP | GO:0042330 | taxis | 0.008 | 0.023 | 0.012 | 2150/3553/335/6347/3569/5054/4363 | 7 |
| BP | GO:0051147 | regulation of muscle cell differentiation | 0.008 | 0.023 | 0.012 | 6648/1786/23411/5465 | 4 |
| BP | GO:0022011 | myelination in peripheral nervous system | 0.008 | 0.023 | 0.012 | 208/6647 | 2 |
| BP | GO:0032104 | regulation of response to extracellular stimulus | 0.008 | 0.023 | 0.012 | 5465/3952 | 2 |
| BP | GO:0032107 | regulation of response to nutrient levels | 0.008 | 0.023 | 0.012 | 5465/3952 | 2 |
| BP | GO:0032292 | peripheral nervous system axon ensheathment | 0.008 | 0.023 | 0.012 | 208/6647 | 2 |
| BP | GO:0032801 | receptor catabolic process | 0.008 | 0.023 | 0.012 | 348/5770 | 2 |
| BP | GO:0051953 | negative regulation of amine transport | 0.008 | 0.023 | 0.012 | 7124/3952 | 2 |
| BP | GO:2000756 | regulation of peptidyl-lysine acetylation | 0.008 | 0.023 | 0.012 | 3066/23411 | 2 |
| BP | GO:0071230 | cellular response to amino acid stimulus | 0.008 | 0.024 | 0.013 | 1786/1291/7124 | 3 |
| BP | GO:0007254 | JNK cascade | 0.008 | 0.024 | 0.013 | 2150/7124/3553/3552 | 4 |
| BP | GO:0016052 | carbohydrate catabolic process | 0.008 | 0.024 | 0.013 | 3630/4758/7351/5465 | 4 |
| BP | GO:1903844 | gulation of cellular response to transforming growth factor beta stimul | 0.008 | 0.024 | 0.013 | 3066/5468/23411/5465 | 4 |
| BP | GO:1901379 | regulation of potassium ion transmembrane transport | 0.009 | 0.024 | 0.013 | 3688/857/481 | 3 |
| BP | GO:2000628 | regulation of miRNA metabolic process | 0.009 | 0.024 | 0.013 | 5468/7124/5465 | 3 |
| BP | GO:0015813 | L-glutamate transmembrane transport | 0.009 | 0.025 | 0.013 | 3688/7124 | 2 |
| BP | GO:0051354 | negative regulation of oxidoreductase activity | 0.009 | 0.025 | 0.013 | 2717/3630 | 2 |
| BP | GO:0098901 | regulation of cardiac muscle cell action potential | 0.009 | 0.025 | 0.013 | 857/5318 | 2 |
| BP | GO:0033077 | T cell differentiation in thymus | 0.009 | 0.025 | 0.013 | 3553/6647/3552 | 3 |
| BP | GO:0071868 | cellular response to monoamine stimulus | 0.009 | 0.025 | 0.013 | 2778/3066/9370 | 3 |
| BP | GO:0071870 | cellular response to catecholamine stimulus | 0.009 | 0.025 | 0.013 | 2778/3066/9370 | 3 |
| BP | GO:0051170 | import into nucleus | 0.009 | 0.025 | 0.013 | 4928/5743/948/3952 | 4 |
| BP | GO:0070665 | positive regulation of leukocyte proliferation | 0.009 | 0.025 | 0.013 | 3553/3569/3952/3552 | 4 |
| BP | GO:0043433 | negative regulation of DNA-binding transcription factor activity | 0.009 | 0.025 | 0.013 | 3066/23411/7466/8431 | 4 |
| BP | GO:0033627 | cell adhesion mediated by integrin | 0.009 | 0.026 | 0.014 | 3688/3383/5054 | 3 |
| BP | GO:0043406 | positive regulation of MAP kinase activity | 0.009 | 0.026 | 0.014 | 7124/3553/5770 | 3 |
| BP | GO:0090068 | positive regulation of cell cycle process | 0.009 | 0.026 | 0.014 | 3630/7124/3553/9510/3552 | 5 |
| BP | GO:0010894 | negative regulation of steroid biosynthetic process | 0.009 | 0.026 | 0.014 | 348/6647 | 2 |
| BP | GO:0010955 | negative regulation of protein processing | 0.009 | 0.026 | 0.014 | 5327/5054 | 2 |
| BP | GO:0061098 | positive regulation of protein tyrosine kinase activity | 0.009 | 0.026 | 0.014 | 1636/5770 | 2 |
| BP | GO:0065005 | protein-lipid complex assembly | 0.009 | 0.026 | 0.014 | 348/335 | 2 |
| BP | GO:1900101 | regulation of endoplasmic reticulum unfolded protein response | 0.009 | 0.026 | 0.014 | 5770/7466 | 2 |
| BP | GO:1903318 | negative regulation of protein maturation | 0.009 | 0.026 | 0.014 | 5327/5054 | 2 |
| BP | GO:0006096 | glycolytic process | 0.009 | 0.026 | 0.014 | 3630/7351/5465 | 3 |
| BP | GO:0032945 | negative regulation of mononuclear cell proliferation | 0.009 | 0.026 | 0.014 | 7356/5573/1401 | 3 |
| BP | GO:0034620 | cellular response to unfolded protein | 0.009 | 0.026 | 0.014 | 5770/26291/7466 | 3 |
| BP | GO:0050829 | defense response to Gram-negative bacterium | 0.009 | 0.026 | 0.014 | 2150/3569/5054 | 3 |
| BP | GO:0014706 | striated muscle tissue development | 0.009 | 0.026 | 0.014 | 3688/34/5318/5573/5465 | 5 |
| BP | GO:0015807 | L-amino acid transport | 0.01 | 0.027 | 0.014 | 3688/7124/7351 | 3 |
| BP | GO:0071867 | response to monoamine | 0.01 | 0.027 | 0.014 | 2778/3066/9370 | 3 |
| BP | GO:0071869 | response to catecholamine | 0.01 | 0.027 | 0.014 | 2778/3066/9370 | 3 |
| BP | GO:0034976 | response to endoplasmic reticulum stress | 0.01 | 0.027 | 0.014 | 857/5770/23411/26291/7466 | 5 |
| BP | GO:0014044 | Schwann cell development | 0.01 | 0.027 | 0.015 | 208/6647 | 2 |
| BP | GO:0032682 | negative regulation of chemokine production | 0.01 | 0.027 | 0.015 | 2150/3569 | 2 |
| BP | GO:0043507 | positive regulation of JUN kinase activity | 0.01 | 0.027 | 0.015 | 7124/5770 | 2 |
| BP | GO:0086011 | membrane repolarization during action potential | 0.01 | 0.027 | 0.015 | 857/481 | 2 |
| BP | GO:1900017 | itive regulation of cytokine production involved in inflammatory respo | 0.01 | 0.027 | 0.015 | 7124/3569 | 2 |
| BP | GO:1900027 | regulation of ruffle assembly | 0.01 | 0.027 | 0.015 | 857/3383 | 2 |
| BP | GO:1900745 | positive regulation of p38MAPK cascade | 0.01 | 0.027 | 0.015 | 3553/3952 | 2 |
| BP | GO:0031507 | heterochromatin formation | 0.01 | 0.027 | 0.015 | 3066/1786/23411 | 3 |
| BP | GO:0045824 | negative regulation of innate immune response | 0.01 | 0.027 | 0.015 | 5468/3630/2671 | 3 |
| BP | GO:0140747 | regulation of ncRNA transcription | 0.01 | 0.028 | 0.015 | 5468/7124/5465 | 3 |
| BP | GO:0010765 | positive regulation of sodium ion transport | 0.011 | 0.029 | 0.015 | 5318/481 | 2 |
| BP | GO:0016242 | negative regulation of macroautophagy | 0.011 | 0.029 | 0.015 | 3162/4864 | 2 |
| BP | GO:0030149 | sphingolipid catabolic process | 0.011 | 0.029 | 0.015 | 2717/4758 | 2 |
| BP | GO:0048710 | regulation of astrocyte differentiation | 0.011 | 0.029 | 0.015 | 3569/3949 | 2 |
| BP | GO:0070498 | interleukin-1-mediated signaling pathway | 0.011 | 0.029 | 0.015 | 3553/3569 | 2 |
| BP | GO:0098868 | bone growth | 0.011 | 0.029 | 0.015 | 3953/3952 | 2 |
| BP | GO:0031214 | biomineral tissue development | 0.011 | 0.029 | 0.015 | 5743/5465/3952/5167 | 4 |
| BP | GO:1902905 | positive regulation of supramolecular fiber organization | 0.011 | 0.029 | 0.015 | 2150/2876/348/335 | 4 |
| BP | GO:0039531 | on of viral-induced cytoplasmic pattern recognition receptor signaling | 0.011 | 0.029 | 0.015 | 2150/857/948 | 3 |
| BP | GO:0042475 | odontogenesis of dentin-containing tooth | 0.011 | 0.029 | 0.015 | 3066/5465/5054 | 3 |
| BP | GO:0043502 | regulation of muscle adaptation | 0.011 | 0.029 | 0.015 | 5468/7133/5465 | 3 |
| BP | GO:1900182 | positive regulation of protein localization to nucleus | 0.011 | 0.029 | 0.015 | 3630/5743/3952 | 3 |
| BP | GO:0002460 | sed on somatic recombination of immune receptors built from immuno | 0.011 | 0.029 | 0.015 | 7124/3553/7133/196/3383/3569 | 6 |
| BP | GO:0032411 | positive regulation of transporter activity | 0.011 | 0.029 | 0.016 | 6347/9370/481 | 3 |
| BP | GO:0071229 | cellular response to acid chemical | 0.011 | 0.029 | 0.016 | 1786/1291/7124 | 3 |
| BP | GO:0007611 | learning or memory | 0.011 | 0.03 | 0.016 | 5577/348/3688/5743/3949 | 5 |
| BP | GO:0006346 | DNA methylation-dependent heterochromatin formation | 0.011 | 0.03 | 0.016 | 1786/23411 | 2 |
| BP | GO:0033032 | regulation of myeloid cell apoptotic process | 0.011 | 0.03 | 0.016 | 23411/9370 | 2 |
| BP | GO:0045777 | positive regulation of blood pressure | 0.011 | 0.03 | 0.016 | 1636/153 | 2 |
| BP | GO:1903715 | regulation of aerobic respiration | 0.011 | 0.03 | 0.016 | 7124/3416 | 2 |
| BP | GO:1904893 | negative regulation of receptor signaling pathway via STAT | 0.011 | 0.03 | 0.016 | 5468/857 | 2 |
| BP | GO:0009267 | cellular response to starvation | 0.011 | 0.031 | 0.016 | 1312/23411/7351/5465 | 4 |
| BP | GO:0090101 | lation of transmembrane receptor protein serine/threonine kinase signa | 0.011 | 0.031 | 0.016 | 3066/5468/23411/5465 | 4 |
| BP | GO:0002042 | cell migration involved in sprouting angiogenesis | 0.012 | 0.031 | 0.016 | 3162/3688/5743 | 3 |
| BP | GO:0002690 | positive regulation of leukocyte chemotaxis | 0.012 | 0.031 | 0.016 | 2150/3569/5054 | 3 |
| BP | GO:0034109 | homotypic cell-cell adhesion | 0.012 | 0.031 | 0.016 | 2778/5318/3569 | 3 |
| BP | GO:0070664 | negative regulation of leukocyte proliferation | 0.012 | 0.031 | 0.016 | 7356/5573/1401 | 3 |
| BP | GO:0001894 | tissue homeostasis | 0.012 | 0.031 | 0.017 | 7879/3688/5743/6647/3569 | 5 |
| BP | GO:0060249 | anatomical structure homeostasis | 0.012 | 0.031 | 0.017 | 7879/3688/5743/6647/3569 | 5 |

| | | | | | | | |
|---|---|---|---|---|---|---|---|
| BP | GO:0051952 | regulation of amine transport | 0.012 | 0.032 | 0.017 | 3688/7124/3952 | 3 |
| BP | GO:0061337 | cardiac conduction | 0.012 | 0.032 | 0.017 | 857/5318/481 | 3 |
| BP | GO:0050954 | sensory perception of mechanical stimulus | 0.012 | 0.032 | 0.017 | 2876/7124/6647/7466 | 4 |
| BP | GO:1905475 | regulation of protein localization to membrane | 0.012 | 0.032 | 0.017 | 3688/3630/7124/208 | 4 |
| BP | GO:0010661 | positive regulation of muscle cell apoptotic process | 0.012 | 0.032 | 0.017 | 6648/5468 | 2 |
| BP | GO:0032743 | positive regulation of interleukin-2 production | 0.012 | 0.032 | 0.017 | 3553/3552 | 2 |
| BP | GO:0034405 | response to fluid shear stress | 0.012 | 0.032 | 0.017 | 1636/5743 | 2 |
| BP | GO:0035025 | positive regulation of Rho protein signal transduction | 0.012 | 0.032 | 0.017 | 2150/335 | 2 |
| BP | GO:0045939 | negative regulation of steroid metabolic process | 0.012 | 0.032 | 0.017 | 348/6647 | 2 |
| BP | GO:0090075 | relaxation of muscle | 0.012 | 0.032 | 0.017 | 6647/481 | 2 |
| BP | GO:0006835 | dicarboxylic acid transport | 0.012 | 0.032 | 0.017 | 3688/7124/7351 | 3 |
| BP | GO:0046578 | regulation of Ras protein signal transduction | 0.012 | 0.033 | 0.017 | 2150/348/3688/335 | 4 |
| BP | GO:0006942 | regulation of striated muscle contraction | 0.013 | 0.033 | 0.017 | 857/5318/481 | 3 |
| BP | GO:0010596 | negative regulation of endothelial cell migration | 0.013 | 0.033 | 0.017 | 5468/348/7124 | 3 |
| BP | GO:0002719 | egative regulation of cytokine production involved in immune respons | 0.013 | 0.033 | 0.017 | 7124/335 | 2 |
| BP | GO:0008340 | determination of adult lifespan | 0.013 | 0.033 | 0.017 | 6647/3952 | 2 |
| BP | GO:0030947 | gulation of vascular endothelial growth factor receptor signaling pathw | 0.013 | 0.033 | 0.017 | 3553/5770 | 2 |
| BP | GO:0048147 | negative regulation of fibroblast proliferation | 0.013 | 0.033 | 0.017 | 6648/857 | 2 |
| BP | GO:0051968 | positive regulation of synaptic transmission, glutamatergic | 0.013 | 0.033 | 0.017 | 5743/6347 | 2 |
| BP | GO:0061081 | gulation of myeloid leukocyte cytokine production involved in immun | 0.013 | 0.033 | 0.017 | 23411/948 | 2 |
| BP | GO:0086004 | regulation of cardiac muscle cell contraction | 0.013 | 0.033 | 0.017 | 857/5318 | 2 |
| BP | GO:0086005 | ventricular cardiac muscle cell action potential | 0.013 | 0.033 | 0.017 | 857/5318 | 2 |
| BP | GO:0090050 | positive regulation of cell migration involved in sprouting angiogenesi | 0.013 | 0.033 | 0.017 | 3162/5743 | 2 |
| BP | GO:0098810 | neurotransmitter reuptake | 0.013 | 0.033 | 0.017 | 3688/6581 | 2 |
| BP | GO:1902882 | regulation of response to oxidative stress | 0.013 | 0.033 | 0.017 | 10516/948 | 2 |
| BP | GO:0061448 | connective tissue development | 0.013 | 0.034 | 0.018 | 5468/4023/23411/79068/3952 | 5 |
| BP | GO:0070828 | heterochromatin organization | 0.013 | 0.034 | 0.018 | 3066/1786/23411 | 3 |
| BP | GO:0006941 | striated muscle contraction | 0.013 | 0.034 | 0.018 | 7124/857/5318/481 | 4 |
| BP | GO:0003333 | amino acid transmembrane transport | 0.013 | 0.034 | 0.018 | 3688/7124/7351 | 3 |
| BP | GO:0002474 | ntigen processing and presentation of peptide antigen via MHC class | 0.013 | 0.034 | 0.018 | 1636/3416 | 2 |
| BP | GO:0010984 | regulation of lipoprotein particle clearance | 0.013 | 0.034 | 0.018 | 26291/3949 | 2 |
| BP | GO:0032094 | response to food | 0.013 | 0.034 | 0.018 | 5465/3952 | 2 |
| BP | GO:0046164 | alcohol catabolic process | 0.013 | 0.034 | 0.018 | 348/949 | 2 |
| BP | GO:0050931 | pigment cell differentiation | 0.013 | 0.034 | 0.018 | 6648/5167 | 2 |
| BP | GO:0071711 | basement membrane organization | 0.013 | 0.034 | 0.018 | 3688/857 | 2 |
| BP | GO:0043266 | regulation of potassium ion transport | 0.014 | 0.035 | 0.019 | 3688/857/481 | 3 |
| BP | GO:0006304 | DNA modification | 0.014 | 0.036 | 0.019 | 1786/79068/4968 | 3 |
| BP | GO:0097306 | cellular response to alcohol | 0.014 | 0.036 | 0.019 | 1636/6648/196 | 3 |
| BP | GO:1901655 | cellular response to ketone | 0.014 | 0.036 | 0.019 | 1636/23411/196 | 3 |
| BP | GO:0010934 | macrophage cytokine production | 0.014 | 0.036 | 0.019 | 23411/948 | 2 |
| BP | GO:0010935 | regulation of macrophage cytokine production | 0.014 | 0.036 | 0.019 | 23411/948 | 2 |
| BP | GO:0043243 | positive regulation of protein-containing complex disassembly | 0.014 | 0.036 | 0.019 | 2150/7124 | 2 |
| BP | GO:0044058 | regulation of digestive system process | 0.014 | 0.036 | 0.019 | 335/3952 | 2 |
| BP | GO:1905477 | positive regulation of protein localization to membrane | 0.014 | 0.037 | 0.02 | 3688/7124/208 | 3 |
| BP | GO:0010586 | miRNA metabolic process | 0.015 | 0.038 | 0.02 | 5468/7124/5465 | 3 |
| BP | GO:0001990 | regulation of systemic arterial blood pressure by hormone | 0.015 | 0.038 | 0.02 | 1636/2150 | 2 |
| BP | GO:0007616 | long-term memory | 0.015 | 0.038 | 0.02 | 348/3949 | 2 |
| BP | GO:0010737 | protein kinase A signaling | 0.015 | 0.038 | 0.02 | 9370/2641 | 2 |
| BP | GO:0014912 | negative regulation of smooth muscle cell migration | 0.015 | 0.038 | 0.02 | 9370/5054 | 2 |
| BP | GO:0032965 | regulation of collagen biosynthetic process | 0.015 | 0.038 | 0.02 | 3066/3569 | 2 |
| BP | GO:0046466 | membrane lipid catabolic process | 0.015 | 0.038 | 0.02 | 2717/4758 | 2 |
| BP | GO:0060416 | response to growth hormone | 0.015 | 0.038 | 0.02 | 9518/5770 | 2 |
| BP | GO:0086091 | regulation of heart rate by cardiac conduction | 0.015 | 0.038 | 0.02 | 857/5318 | 2 |
| BP | GO:1990776 | response to angiotensin | 0.015 | 0.038 | 0.02 | 857/5743 | 2 |
| BP | GO:0032526 | response to retinoic acid | 0.015 | 0.038 | 0.02 | 3066/7124/3952 | 3 |
| BP | GO:0055001 | muscle cell development | 0.015 | 0.039 | 0.021 | 2876/3688/5573/5465 | 4 |
| BP | GO:0006650 | glycerophospholipid metabolic process | 0.015 | 0.039 | 0.021 | 335/1071/3990/949/3949 | 5 |
| BP | GO:0048144 | fibroblast proliferation | 0.015 | 0.039 | 0.021 | 6648/2876/857 | 3 |
| BP | GO:0014037 | Schwann cell differentiation | 0.015 | 0.039 | 0.021 | 208/6647 | 2 |
| BP | GO:0051938 | L-glutamate import | 0.015 | 0.039 | 0.021 | 3688/7124 | 2 |
| BP | GO:0061136 | regulation of proteasomal protein catabolic process | 0.016 | 0.039 | 0.021 | 2876/348/857/7466 | 4 |
| BP | GO:0015837 | amine transport | 0.016 | 0.04 | 0.021 | 3688/7124/3952 | 3 |
| BP | GO:1903076 | regulation of protein localization to plasma membrane | 0.016 | 0.04 | 0.021 | 3688/3630/7124 | 3 |
| BP | GO:0006939 | smooth muscle contraction | 0.016 | 0.041 | 0.022 | 857/5743/6647 | 3 |
| BP | GO:0035967 | cellular response to topologically incorrect protein | 0.016 | 0.041 | 0.022 | 5770/26291/7466 | 3 |
| BP | GO:0000266 | mitochondrial fission | 0.016 | 0.041 | 0.022 | 5468/7351 | 2 |
| BP | GO:0000731 | DNA synthesis involved in DNA repair | 0.016 | 0.041 | 0.022 | 5422/23411 | 2 |
| BP | GO:0031056 | regulation of histone modification | 0.016 | 0.041 | 0.022 | 23411/2641 | 2 |
| BP | GO:0035886 | vascular associated smooth muscle cell differentiation | 0.016 | 0.041 | 0.022 | 6648/1786 | 2 |
| BP | GO:0042307 | positive regulation of protein import into nucleus | 0.016 | 0.041 | 0.022 | 5743/3952 | 2 |
| BP | GO:0002444 | myeloid leukocyte mediated immunity | 0.017 | 0.042 | 0.022 | 1636/2150/3569 | 3 |
| BP | GO:0008637 | apoptotic mitochondrial changes | 0.017 | 0.042 | 0.022 | 6648/2876/4318 | 3 |
| BP | GO:0042116 | macrophage activation | 0.017 | 0.042 | 0.022 | 7124/3569/3949 | 3 |
| BP | GO:0045445 | myoblast differentiation | 0.017 | 0.042 | 0.022 | 2876/3688/7124 | 3 |
| BP | GO:0007626 | locomotory behavior | 0.017 | 0.042 | 0.022 | 6648/348/6647/4864 | 4 |
| BP | GO:0007229 | integrin-mediated signaling pathway | 0.017 | 0.042 | 0.023 | 3688/335/9510 | 3 |
| BP | GO:0010717 | regulation of epithelial to mesenchymal transition | 0.017 | 0.042 | 0.023 | 3066/3553/3569 | 3 |
| BP | GO:0035249 | synaptic transmission, glutamatergic | 0.017 | 0.042 | 0.023 | 7124/5743/6347 | 3 |
| BP | GO:0043280 | egulation of cysteine-type endopeptidase activity involved in apoptoti | 0.017 | 0.042 | 0.023 | 5468/7124/23411 | 3 |
| BP | GO:0060840 | artery development | 0.017 | 0.042 | 0.023 | 348/3952/3949 | 3 |
| BP | GO:0071347 | cellular response to interleukin-1 | 0.017 | 0.042 | 0.023 | 3553/6347/3569 | 3 |
| BP | GO:0043388 | positive regulation of DNA binding | 0.017 | 0.042 | 0.023 | 5468/4318 | 2 |
| BP | GO:1903115 | regulation of actin filament-based movement | 0.017 | 0.042 | 0.023 | 857/5318 | 2 |
| BP | GO:1904994 | regulation of leukocyte adhesion to vascular endothelial cell | 0.017 | 0.042 | 0.023 | 7124/3569 | 2 |
| BP | GO:0043087 | regulation of GTPase activity | 0.017 | 0.043 | 0.023 | 2150/3688/6347/6647/153 | 5 |
| BP | GO:0032733 | positive regulation of interleukin-10 production | 0.018 | 0.044 | 0.024 | 2150/3569 | 2 |
| BP | GO:0042119 | neutrophil activation | 0.018 | 0.044 | 0.024 | 2150/7124 | 2 |
| BP | GO:0045922 | negative regulation of fatty acid metabolic process | 0.018 | 0.044 | 0.024 | 3630/23411 | 2 |
| BP | GO:1901983 | regulation of protein acetylation | 0.018 | 0.044 | 0.024 | 3066/23411 | 2 |
| BP | GO:0034504 | protein localization to nucleus | 0.018 | 0.044 | 0.024 | 3630/4928/5743/948/3952 | 5 |
| BP | GO:0042752 | regulation of circadian rhythm | 0.018 | 0.045 | 0.024 | 5468/5465/153 | 3 |
| BP | GO:2000241 | regulation of reproductive process | 0.019 | 0.046 | 0.024 | 5327/3066/6647/3552 | 4 |
| BP | GO:0032735 | positive regulation of interleukin-12 production | 0.019 | 0.046 | 0.025 | 948/3952 | 2 |
| BP | GO:0090317 | negative regulation of intracellular protein transport | 0.019 | 0.046 | 0.025 | 9370/948 | 2 |
| BP | GO:0048640 | negative regulation of developmental growth | 0.019 | 0.046 | 0.025 | 9518/5465/153 | 3 |
| BP | GO:0001822 | kidney development | 0.019 | 0.046 | 0.025 | 1636/9510/9370/4318/7466 | 5 |
| BP | GO:0071383 | cellular response to steroid hormone stimulus | 0.019 | 0.047 | 0.025 | 1636/23411/5465/4864 | 4 |
| BP | GO:1901654 | response to ketone | 0.019 | 0.047 | 0.025 | 1636/857/23411/196 | 4 |
| BP | GO:0010633 | negative regulation of epithelial cell migration | 0.019 | 0.047 | 0.025 | 5468/348/7124 | 3 |
| BP | GO:0019882 | antigen processing and presentation | 0.019 | 0.047 | 0.025 | 1636/3416/3383 | 3 |
| BP | GO:0019722 | calcium-mediated signaling | 0.019 | 0.048 | 0.025 | 7124/283455/5443/481 | 4 |
| BP | GO:0006284 | base-excision repair | 0.019 | 0.048 | 0.025 | 4968/5428 | 2 |
| BP | GO:0043506 | regulation of JUN kinase activity | 0.019 | 0.048 | 0.025 | 7124/5770 | 2 |
| BP | GO:0097178 | ruffle assembly | 0.019 | 0.048 | 0.025 | 857/3383 | 2 |
| BP | GO:1903573 | negative regulation of response to endoplasmic reticulum stress | 0.019 | 0.048 | 0.025 | 5770/7466 | 2 |
| BP | GO:0042303 | molting cycle | 0.02 | 0.048 | 0.025 | 3066/7124/5743 | 3 |
| BP | GO:0042633 | hair cycle | 0.02 | 0.048 | 0.025 | 3066/7124/5743 | 3 |
| BP | GO:2000278 | regulation of DNA biosynthetic process | 0.02 | 0.048 | 0.025 | 7124/2671/9370 | 3 |
| BP | GO:0006090 | pyruvate metabolic process | 0.02 | 0.049 | 0.026 | 3630/7351/5465 | 3 |

| | | | | | | | |
|---|---|---|---|---|---|---|---|
| BP | GO:0001504 | neurotransmitter uptake | 0.02 | 0.049 | 0.026 | 3688/6581 | 2 |
| BP | GO:0030574 | collagen catabolic process | 0.02 | 0.049 | 0.026 | 3688/4318 | 2 |
| BP | GO:0031641 | regulation of myelination | 0.02 | 0.049 | 0.026 | 7124/7133 | 2 |
| BP | GO:0042596 | fear response | 0.02 | 0.049 | 0.026 | 348/153 | 2 |
| BP | GO:0045214 | sarcomere organization | 0.02 | 0.049 | 0.026 | 3688/5573 | 2 |
| BP | GO:0045687 | positive regulation of glial cell differentiation | 0.02 | 0.049 | 0.026 | 3066/7133 | 2 |
| BP | GO:0048713 | regulation of oligodendrocyte differentiation | 0.02 | 0.049 | 0.026 | 3066/7133 | 2 |
| BP | GO:0090199 | regulation of release of cytochrome c from mitochondria | 0.02 | 0.049 | 0.026 | 2876/4318 | 2 |
| BP | GO:1900744 | regulation of p38MAPK cascade | 0.02 | 0.049 | 0.026 | 3553/3952 | 2 |
| BP | GO:1901888 | regulation of cell junction assembly | 0.02 | 0.049 | 0.026 | 1636/7124/3553/857 | 4 |
| BP | GO:0009308 | amine metabolic process | 0.02 | 0.05 | 0.026 | 3630/1312/6581 | 3 |
| CC | GO:0034358 | plasma lipoprotein particle | 0 | 0 | 0 | 348/4023/335/1071/3990/3949 | 6 |
| CC | GO:1990777 | lipoprotein particle | 0 | 0 | 0 | 348/4023/335/1071/3990/3949 | 6 |
| CC | GO:0032994 | protein-lipid complex | 0 | 0 | 0 | 348/4023/335/1071/3990/3949 | 6 |
| CC | GO:0045121 | membrane raft | 0 | 0 | 0 | 3688/7124/857/5743/5573/7133/948/3383/3667/949/4864 | 11 |
| CC | GO:0098857 | membrane microdomain | 0 | 0 | 0 | 3688/7124/857/5743/5573/7133/948/3383/3667/949/4864 | 11 |
| CC | GO:0062023 | collagen-containing extracellular matrix | 0 | 0 | 0 | 9518/348/1291/10516/335/83729/9510/5265/9370/3383/5054/431[illegible] | 12 |
| CC | GO:0005788 | endoplasmic reticulum lumen | 0 | 0 | 0 | 348/1291/3630/335/3990/5265/5743/2641/3569/7466 | 10 |
| CC | GO:0034774 | secretory granule lumen | 0 | 0 | 0 | 2717/3630/4758/335/10577/5265/2641/5054/5443/1727 | 10 |
| CC | GO:0060205 | cytoplasmic vesicle lumen | 0 | 0 | 0 | 2717/3630/4758/335/10577/5265/2641/5054/5443/1727 | 10 |
| CC | GO:0031983 | vesicle lumen | 0 | 0 | 0 | 2717/3630/4758/335/10577/5265/2641/5054/5443/1727 | 10 |
| CC | GO:0034364 | high-density lipoprotein particle | 0 | 0 | 0 | 348/335/1071/3990 | 4 |
| CC | GO:0044853 | plasma membrane raft | 0 | 0 | 0 | 857/5743/5573/948/3667/949 | 6 |
| CC | GO:0005901 | caveola | 0 | 0.001 | 0 | 857/5743/948/3667/949 | 5 |
| CC | GO:0009925 | basal plasma membrane | 0 | 0.001 | 0.001 | 1636/3416/6581/3953/5167/4363/481/3949 | 8 |
| CC | GO:0045178 | basal part of cell | 0 | 0.001 | 0.001 | 1636/3416/6581/3953/5167/4363/481/3949 | 8 |
| CC | GO:0034361 | very-low-density lipoprotein particle | 0 | 0.002 | 0.001 | 348/4023/335 | 3 |
| CC | GO:0034385 | triglyceride-rich plasma lipoprotein particle | 0 | 0.002 | 0.001 | 348/4023/335 | 3 |
| CC | GO:0005811 | lipid droplet | 0 | 0.002 | 0.001 | 7879/10280/857/5346/1727 | 5 |
| CC | GO:0016323 | basolateral plasma membrane | 0 | 0.002 | 0.001 | 3416/6581/3953/5167/4363/481/3949 | 7 |
| CC | GO:0030666 | endocytic vesicle membrane | 0 | 0.004 | 0.003 | 7879/348/857/948/949/3949 | 6 |
| CC | GO:0005640 | nuclear outer membrane | 0 | 0.005 | 0.003 | 10280/6581/5743 | 3 |
| CC | GO:0009897 | external side of plasma membrane | 0 | 0.005 | 0.003 | 1636/3688/7124/3416/3953/948/3383/3949 | 8 |
| CC | GO:0005635 | nuclear envelope | 0 | 0.005 | 0.004 | 7356/5422/10280/4928/6581/129787/5743/23411/4864 | 9 |
| CC | GO:0005769 | early endosome | 0.001 | 0.009 | 0.006 | 2150/348/208/335/857/5770/153/3949 | 8 |
| CC | GO:1904090 | peptidase inhibitor complex | 0.001 | 0.011 | 0.008 | 5327/5054 | 2 |
| CC | GO:0030139 | endocytic vesicle | 0.001 | 0.011 | 0.008 | 7879/348/335/857/948/949/3949 | 7 |
| CC | GO:0034362 | low-density lipoprotein particle | 0.001 | 0.012 | 0.009 | 348/3949 | 2 |
| CC | GO:0042627 | chylomicron | 0.001 | 0.012 | 0.009 | 348/4023 | 2 |
| CC | GO:0014704 | intercalated disc | 0.002 | 0.014 | 0.01 | 3688/5318/481 | 3 |
| CC | GO:0005637 | nuclear inner membrane | 0.002 | 0.02 | 0.014 | 10280/5743/23411 | 3 |
| CC | GO:0031965 | nuclear membrane | 0.003 | 0.026 | 0.018 | 10280/4928/6581/129787/5743/23411 | 6 |
| CC | GO:0042575 | DNA polymerase complex | 0.003 | 0.027 | 0.019 | 5422/5428 | 2 |
| CC | GO:0042383 | sarcolemma | 0.004 | 0.031 | 0.022 | 3688/1291/857/481 | 4 |
| CC | GO:0005765 | lysosomal membrane | 0.004 | 0.031 | 0.022 | 7879/1291/4758/5167/949/4864/3949 | 7 |
| CC | GO:0098852 | lytic vacuole membrane | 0.004 | 0.031 | 0.022 | 7879/1291/4758/5167/949/4864/3949 | 7 |
| CC | GO:0044291 | cell-cell contact zone | 0.005 | 0.032 | 0.022 | 3688/5318/481 | 3 |
| CC | GO:0071682 | endocytic vesicle lumen | 0.005 | 0.034 | 0.024 | 348/335 | 2 |
| CC | GO:0031968 | organelle outer membrane | 0.006 | 0.037 | 0.026 | 3162/10280/6581/5743/1727 | 5 |
| CC | GO:0019867 | outer membrane | 0.006 | 0.037 | 0.026 | 3162/10280/6581/5743/1727 | 5 |
| CC | GO:0098992 | neuronal dense core vesicle | 0.006 | 0.037 | 0.026 | 5173/153 | 2 |
| CC | GO:0045177 | apical part of cell | 0.006 | 0.037 | 0.026 | 2778/5327/6581/948/4363/481/3949 | 7 |
| CC | GO:0005774 | vacuolar membrane | 0.007 | 0.043 | 0.03 | 7879/1291/4758/5167/949/4864/3949 | 7 |
| CC | GO:0005759 | mitochondrial matrix | 0.007 | 0.043 | 0.03 | 6648/2876/34/5770/6647/4968/5428 | 7 |
| CC | GO:0043025 | neuronal cell body | 0.008 | 0.043 | 0.03 | 5173/5577/348/7124/6581/6647/7133 | 7 |
| CC | GO:0031091 | platelet alpha granule | 0.009 | 0.047 | 0.033 | 5265/948/5054 | 3 |
| CC | GO:0035578 | azurophil granule lumen | 0.009 | 0.047 | 0.033 | 2717/10577/1727 | 3 |
| CC | GO:0098685 | Schaffer collateral - CA1 synapse | 0.009 | 0.047 | 0.033 | 5327/3688/153 | 3 |
| CC | GO:0005775 | vacuolar lumen | 0.009 | 0.047 | 0.033 | 2717/4758/10577/1727 | 4 |
| MF | GO:0071813 | lipoprotein particle binding | 0 | 0 | 0 | 348/4023/335/3990/948/1401/949/3949 | 8 |
| MF | GO:0071814 | protein-lipid complex binding | 0 | 0 | 0 | 348/4023/335/3990/948/1401/949/3949 | 8 |
| MF | GO:0030169 | low-density lipoprotein particle binding | 0 | 0 | 0 | 3990/948/1401/949/3949 | 5 |
| MF | GO:0002020 | protease binding | 0 | 0 | 0 | 2150/3688/3630/7124/5265/5054/5428/3949 | 8 |
| MF | GO:0015248 | sterol transporter activity | 0 | 0 | 0 | 348/335/1071/10577/4864 | 5 |
| MF | GO:0001618 | virus receptor activity | 0 | 0 | 0 | 3688/3416/3383/949/4864/3949 | 6 |
| MF | GO:0140272 | exogenous protein binding | 0 | 0 | 0 | 3688/3416/3383/949/4864/3949 | 6 |
| MF | GO:0005158 | insulin receptor binding | 0 | 0 | 0 | 3630/5770/3667/5167 | 4 |
| MF | GO:0120020 | cholesterol transfer activity | 0 | 0 | 0 | 348/335/1071/10577 | 4 |
| MF | GO:0120015 | sterol transfer activity | 0 | 0 | 0 | 348/335/1071/10577 | 4 |
| MF | GO:0015485 | cholesterol binding | 0 | 0 | 0 | 335/857/1071/10577/4864 | 5 |
| MF | GO:0005496 | steroid binding | 0 | 0 | 0 | 335/857/1071/10577/3290/4864 | 6 |
| MF | GO:0032934 | sterol binding | 0 | 0 | 0 | 335/857/1071/10577/4864 | 5 |
| MF | GO:0005125 | cytokine activity | 0 | 0.001 | 0 | 9518/7124/3553/83729/6347/9370/3569/3552 | 8 |
| MF | GO:0005319 | lipid transporter activity | 0 | 0.001 | 0 | 348/335/1071/10577/948/4363/4864 | 7 |
| MF | GO:0008035 | high-density lipoprotein particle binding | 0 | 0.001 | 0.001 | 335/948/949 | 3 |
| MF | GO:0005179 | hormone activity | 0 | 0.001 | 0.001 | 3630/83729/9370/2641/3952/5443 | 6 |
| MF | GO:0016209 | antioxidant activity | 0 | 0.001 | 0.001 | 6648/2876/348/5743/6647 | 5 |
| MF | GO:0005159 | insulin-like growth factor receptor binding | 0 | 0.001 | 0.001 | 2778/3630/3667 | 3 |
| MF | GO:0001540 | amyloid-beta binding | 0 | 0.001 | 0.001 | 348/335/948/949/3949 | 5 |
| MF | GO:0034185 | apolipoprotein binding | 0 | 0.001 | 0.001 | 4023/3990/949 | 3 |
| MF | GO:0050660 | flavin adenine dinucleotide binding | 0 | 0.001 | 0.001 | 34/2671/8540/5447/1727 | 5 |
| MF | GO:0043178 | alcohol binding | 0 | 0.001 | 0.001 | 335/857/1071/10577/4864 | 5 |
| MF | GO:0030228 | lipoprotein particle receptor activity | 0 | 0.002 | 0.001 | 948/949/3949 | 3 |
| MF | GO:0120013 | lipid transfer activity | 0 | 0.002 | 0.002 | 348/335/1071/10577 | 4 |
| MF | GO:0042277 | peptide binding | 0 | 0.003 | 0.002 | 5468/348/4928/335/3416/948/949/3949 | 8 |
| MF | GO:0038024 | cargo receptor activity | 0 | 0.004 | 0.003 | 948/5167/949/3949 | 4 |
| MF | GO:0005044 | scavenger receptor activity | 0 | 0.005 | 0.003 | 948/5167/949 | 3 |
| MF | GO:1990841 | promoter-specific chromatin binding | 0 | 0.005 | 0.003 | 3066/1786/4928/23411 | 4 |
| MF | GO:0070325 | lipoprotein particle receptor binding | 0 | 0.005 | 0.004 | 348/335/1401 | 3 |
| MF | GO:0001664 | G protein-coupled receptor binding | 0.001 | 0.008 | 0.006 | 1636/2778/5173/6347/2641/5443/153 | 7 |
| MF | GO:0033218 | amide binding | 0.001 | 0.012 | 0.009 | 5468/348/4928/335/3416/948/949/3949 | 8 |
| MF | GO:0004862 | cAMP-dependent protein kinase inhibitor activity | 0.001 | 0.012 | 0.009 | 5577/5573 | 2 |
| MF | GO:0016653 | oxidoreductase activity, acting on NAD(P)H, heme protein as accepto[illegible] | 0.001 | 0.012 | 0.009 | 5447/1727 | 2 |
| MF | GO:0052740 | 1-acyl-2-lysophosphatidylserine acylhydrolase activity | 0.001 | 0.012 | 0.009 | 4023/3990 | 2 |
| MF | GO:0008083 | growth factor activity | 0.001 | 0.014 | 0.01 | 9518/2671/83729/26291/3569 | 5 |
| MF | GO:0019215 | intermediate filament binding | 0.001 | 0.014 | 0.01 | 5318/23411 | 2 |
| MF | GO:0052739 | phosphatidylserine 1-acylhydrolase activity | 0.001 | 0.014 | 0.01 | 4023/3990 | 2 |
| MF | GO:0008603 | cAMP-dependent protein kinase regulator activity | 0.002 | 0.016 | 0.012 | 5577/5573 | 2 |
| MF | GO:0034236 | protein kinase A catalytic subunit binding | 0.002 | 0.019 | 0.014 | 5577/5573 | 2 |
| MF | GO:0004879 | nuclear receptor activity | 0.002 | 0.021 | 0.015 | 5468/5465/196 | 3 |
| MF | GO:0098531 | ligand-activated transcription factor activity | 0.002 | 0.021 | 0.015 | 5468/5465/196 | 3 |
| MF | GO:0004222 | metalloendopeptidase activity | 0.002 | 0.021 | 0.015 | 1636/9510/3416/4318 | 4 |
| MF | GO:0005041 | low-density lipoprotein particle receptor activity | 0.002 | 0.023 | 0.017 | 948/3949 | 2 |
| MF | GO:0005149 | interleukin-1 receptor binding | 0.003 | 0.028 | 0.021 | 3553/3552 | 2 |
| MF | GO:0046965 | nuclear retinoid X receptor binding | 0.003 | 0.028 | 0.021 | 5468/8431 | 2 |
| MF | GO:0008970 | phospholipase A1 activity | 0.004 | 0.029 | 0.022 | 4023/3990 | 2 |
| MF | GO:0031690 | adrenergic receptor binding | 0.004 | 0.029 | 0.022 | 2778/153 | 2 |
| MF | GO:0043395 | heparan sulfate proteoglycan binding | 0.004 | 0.029 | 0.022 | 348/4023 | 2 |
| MF | GO:0051428 | peptide hormone receptor binding | 0.004 | 0.032 | 0.024 | 2778/3952 | 2 |

| | | | | | | | |
|---|---|---|---|---|---|---|---|
| MF | GO:0005518 | collagen binding | 0.005 | 0.038 | 0.028 | 3688/1291/4318 | 3 |
| MF | GO:0004407 | histone deacetylase activity | 0.005 | 0.038 | 0.028 | 3066/23411 | 2 |
| MF | GO:0070851 | growth factor receptor binding | 0.005 | 0.038 | 0.028 | 3553/26291/3569/3552 | 4 |
| MF | GO:0033558 | protein lysine deacetylase activity | 0.005 | 0.04 | 0.029 | 3066/23411 | 2 |
| MF | GO:0030552 | cAMP binding | 0.006 | 0.043 | 0.032 | 5577/5573 | 2 |
| MF | GO:0019955 | cytokine binding | 0.006 | 0.043 | 0.032 | 3688/7133/3953/948 | 4 |
| MF | GO:0003887 | DNA-directed DNA polymerase activity | 0.006 | 0.043 | 0.032 | 5422/5428 | 2 |
| MF | GO:0004806 | triglyceride lipase activity | 0.006 | 0.043 | 0.032 | 4023/3990 | 2 |
| MF | GO:0050750 | low-density lipoprotein particle receptor binding | 0.006 | 0.043 | 0.032 | 348/1401 | 2 |
| MF | GO:0005178 | integrin binding | 0.007 | 0.048 | 0.036 | 3688/10516/3553/3383 | 4 |
| MF | GO:0042974 | nuclear retinoic acid receptor binding | 0.007 | 0.048 | 0.036 | 5468/8431 | 2 |
| MF | GO:0061629 | RNA polymerase II-specific DNA-binding transcription factor binding | 0.007 | 0.048 | 0.036 | 3066/5468/23411/5465/196/8431 | 6 |
