## Supplementary Table 7 for "Deciphering the Molecular Mechanism of Post-Acute Sequelae of COVID-19 through Comorbidity Network Analysis"

| Protein | Corr | RR | symbol | RR_idx | Corr_idx | mean_idx | idx |
|---|---|---|---|---|---|---|---|
| 1636 | 0.446 | 62.531 | ACE | 1 | 0 | 0.5 | 0 |
| 3162 | 0.427 | 83.609 | HMOX1 | 0 | 1 | 0.5 | 0 |
| 9518 | 0.287 | 37.007 | GDF15 | 3 | 4 | 3.5 | 2 |
| 2778 | 0.398 | 29.494 | GNAS | 6 | 2 | 4 | 3 |
| 5116 | 0.145 | 47.204 | PCNT | 2 | 7 | 4.5 | 4 |
| 5173 | 0.363 | 25.045 | PDYN | 8 | 3 | 5.5 | 5 |
| 5327 | 0.236 | 28.542 | PLAT | 7 | 6 | 6.5 | 6 |
| 3066 | 0.105 | 35.661 | HDAC2 | 4 | 10 | 7 | 7 |
| 7356 | 0.263 | 16.755 | SCGB1A1 | 16 | 5 | 10.5 | 8 |
| 1786 | 0.087 | 30.226 | DNMT1 | 5 | 16 | 10.5 | 8 |
| 2150 | 0.098 | 18.404 | F2RL1 | 12 | 12 | 12 | 10 |
| 7879 | 0.098 | 18.404 | RAB7A | 12 | 12 | 12 | 10 |
| 166 | 0.098 | 18.404 | TLE5 | 12 | 12 | 12 | 10 |
| 5577 | 0.133 | 11.128 | PRKAR2B | 17 | 8 | 12.5 | 13 |
| 2717 | 0.102 | 18.274 | GLA | 15 | 11 | 13 | 14 |
| 6648 | 0.105 | 7.678 | SOD2 | 20 | 9 | 14.5 | 15 |
| 2876 | 0.088 | 9.272 | GPX1 | 18 | 15 | 16.5 | 16 |
| 3688 | 0.08 | 7.552 | ITGB1 | 21 | 20 | 20.5 | 17 |
| 1291 | 0.08 | 7.552 | COL6A1 | 21 | 20 | 20.5 | 17 |
| 10516 | 0.08 | 7.552 | FBLN5 | 21 | 20 | 20.5 | 17 |
| 3553 | 0.063 | 8.04 | IL1B | 19 | 29 | 24 | 20 |
| 10280 | 0.064 | 6.975 | SIGMAR1 | 24 | 25 | 24.5 | 21 |
| 57534 | 0.064 | 6.975 | MIB1 | 24 | 25 | 24.5 | 21 |
| 5422 | 0.064 | 6.975 | POLA1 | 24 | 25 | 24.5 | 21 |
| 1312 | 0.064 | 6.975 | COMT | 24 | 25 | 24.5 | 21 |
| 55676 | 0.042 | 23.114 | SLC30A6 | 9 | 42 | 25.5 | 25 |
| 3416 | 0.042 | 23.114 | IDE | 9 | 42 | 25.5 | 25 |
| 9510 | 0.042 | 23.114 | ADAMTS1 | 9 | 42 | 25.5 | 25 |
| 3630 | 0.066 | 4.397 | INS | 31 | 23 | 27 | 28 |
| 7124 | 0.066 | 4.397 | TNF | 31 | 23 | 27 | 28 |
| 5468 | 0.086 | 3.71 | PPARG | 38 | 17 | 27.5 | 30 |
| 348 | 0.086 | 3.71 | APOE | 38 | 17 | 27.5 | 30 |
| 4023 | 0.086 | 3.71 | LPL | 38 | 17 | 27.5 | 30 |
| 4928 | 0.057 | 5.126 | NUP98 | 29 | 30 | 29.5 | 33 |
| 2671 | 0.046 | 5.977 | GFER | 28 | 33 | 30.5 | 34 |
| 34 | 0.054 | 3.576 | ACADM | 41 | 32 | 36.5 | 35 |
| 4758 | 0.054 | 2.705 | NEU1 | 45 | 31 | 38 | 36 |
| 5318 | 0.039 | 4.059 | PKP2 | 34 | 46 | 40 | 37 |
| 23177 | 0.039 | 4.059 | CEP68 | 34 | 46 | 40 | 37 |
| 10577 | 0.044 | 1.295 | NPC2 | 48 | 34 | 41 | 39 |
| 3990 | 0.044 | 1.295 | LIPC | 48 | 34 | 41 | 39 |
| 8540 | 0.044 | 1.295 | AGPS | 48 | 34 | 41 | 39 |
| 83729 | 0.044 | 1.295 | INHBE | 48 | 34 | 41 | 39 |
| 335 | 0.044 | 1.295 | APOA1 | 48 | 34 | 41 | 39 |
| 208 | 0.044 | 1.295 | AKT2 | 48 | 34 | 41 | 39 |
| 1071 | 0.044 | 1.295 | CETP | 48 | 34 | 41 | 39 |
| 857 | 0.044 | 1.295 | CAV1 | 48 | 34 | 41 | 39 |
| 5265 | 0.04 | 3.134 | SERPINA1 | 42 | 45 | 43.5 | 47 |
| 6581 | 0.026 | 1.264 | SLC22A3 | 56 | 48 | 52 | 48 |
| 5167 | 0.026 | 1.264 | ENPP1 | 56 | 48 | 52 | 48 |
| 153 | 0.026 | 1.264 | ADRB1 | 56 | 48 | 52 | 48 |
| 4318 | 0.026 | 1.264 | MMP9 | 56 | 48 | 52 | 48 |
| 5443 | 0.026 | 1.264 | POMC | 56 | 48 | 52 | 48 |
| 5054 | 0.026 | 1.264 | SERPINE1 | 56 | 48 | 52 | 48 |
| 283455 | 0.026 | 1.264 | KSR2 | 56 | 48 | 52 | 48 |
| 3952 | 0.026 | 1.264 | LEP | 56 | 48 | 52 | 48 |

| | | | | | | | |
|---|---|---|---|---|---|---|---|
| 3667 | 0.026 | 1.264 | IRS1 | 56 | 48 | 52 | 48 |
| 3569 | 0.026 | 1.264 | IL6 | 56 | 48 | 52 | 48 |
| 3383 | 0.026 | 1.264 | ICAM1 | 56 | 48 | 52 | 48 |
| 2784 | 0.026 | 1.264 | GNB3 | 56 | 48 | 52 | 48 |
| 2641 | 0.026 | 1.264 | GCG | 56 | 48 | 52 | 48 |
| 1401 | 0.026 | 1.264 | CRP | 56 | 48 | 52 | 48 |
| 948 | 0.026 | 1.264 | CD36 | 56 | 48 | 52 | 48 |
| 196 | 0.026 | 1.264 | AHR | 56 | 48 | 52 | 48 |
| 3290 | 0.026 | 1.264 | HSD11B1 | 56 | 48 | 52 | 48 |
| 5465 | 0.026 | 1.264 | PPARA | 56 | 48 | 52 | 48 |
| 129787 | 0.026 | 1.264 | TMEM18 | 56 | 48 | 52 | 48 |
| 3953 | 0.026 | 1.264 | LEPR | 56 | 48 | 52 | 48 |
| 5770 | 0.026 | 1.264 | PTPN1 | 56 | 48 | 52 | 48 |
| 6347 | 0.026 | 1.264 | CCL2 | 56 | 48 | 52 | 48 |
| 23411 | 0.026 | 1.264 | SIRT1 | 56 | 48 | 52 | 48 |
| 5573 | 0.026 | 1.264 | PRKAR1A | 56 | 48 | 52 | 48 |
| 6647 | 0.026 | 1.264 | SOD1 | 56 | 48 | 52 | 48 |
| 5743 | 0.026 | 1.264 | PTGS2 | 56 | 48 | 52 | 48 |
| 26291 | 0.026 | 1.264 | FGF21 | 56 | 48 | 52 | 48 |
| 9370 | 0.026 | 1.264 | ADIPOQ | 56 | 48 | 52 | 48 |
| 7351 | 0.026 | 1.264 | UCP2 | 56 | 48 | 52 | 48 |
| 7133 | 0.026 | 1.264 | TNFRSF1B | 56 | 48 | 52 | 48 |
| 4968 | 0.026 | 1.264 | OGG1 | 56 | 48 | 52 | 48 |
| 79068 | 0.026 | 1.264 | FTO | 56 | 48 | 52 | 48 |
| 949 | 0.023 | 5.016 | SCARB1 | 30 | 81 | 55.5 | 80 |
| 7466 | 0.024 | 4.286 | WFS1 | 33 | 80 | 56.5 | 81 |
| 5428 | 0.019 | 3.968 | POLG | 36 | 82 | 59 | 82 |
| 3552 | 0.019 | 3.968 | IL1A | 36 | 82 | 59 | 82 |
| 4363 | 0.018 | 2.765 | ABCC1 | 43 | 84 | 63.5 | 84 |
| 481 | 0.018 | 2.765 | ATP1B1 | 43 | 84 | 63.5 | 84 |
| 5447 | 0.01 | 1.612 | POR | 46 | 90 | 68 | 86 |
| 1727 | 0.007 | 1.482 | CYB5R3 | 47 | 91 | 69 | 87 |
| 8431 | 0.017 | 1.151 | NR0B2 | 88 | 86 | 87 | 88 |
| 5346 | 0.017 | 1.151 | PLIN1 | 88 | 86 | 87 | 88 |
| 4864 | 0.017 | 1.151 | NPC1 | 88 | 86 | 87 | 88 |
| 3949 | 0.017 | 1.151 | LDLR | 88 | 86 | 87 | 88 |
